# Synthesis of pyrazole-based macrocycles leads to a highly selective inhibitor for MST3

**DOI:** 10.1101/2023.10.20.563248

**Authors:** Jennifer A. Amrhein, Lena M. Berger, Dimitrios-Ilias Balourdas, Andreas C. Joerger, Amelie Menge, Andreas Krämer, Julia M. Frischkorn, Benedict-Tilman Berger, Lewis Elson, Astrid Kaiser, Manfred Schubert-Zsilavecz, Susanne Müller, Stefan Knapp, Thomas Hanke

## Abstract

MST1, MST2, MST3, MST4, and YSK1 are conserved members of the mammalian sterile 20 kinase (MST) family. MSTs regulate key cellular functions such as cell proliferation, cell migration, metabolic regulation, and cell polarity. The MST3 isozyme plays a role in regulation of cell growth, autophagy and apoptosis, and its dysregulation has been linked to the occurrence of high-grade tumors with poor survival prognosis. To date, there are no isoform-selective inhibitors available that could be used for validating the role of MST3 in tumorigenesis and to assess its potential as an anti-cancer target for drug development. To this end, we have designed a new series of 3-aminopyrazole-based macrocycles based on the structure of an acyclic promiscuous kinase inhibitor. By varying moieties targeting the solvent-exposed region and optimizing the linker, macrocycle JA310 (**21c**) was synthesized. JA310 exhibited high cellular potency for MST3 with an EC_50_ = 106 nM and excellent kinome-wide selectivity with significantly lower cellular activity on the closely related kinase MST4 (EC_50_ = 1.4 µM). The high-resolution crystal structure of the MST3-JA310 complex provided intriguing insights into the distinct binding mode of the macrocycle, which was associated with large-scale structural rearrangements, including concerted induced-fit movements of the glycine-rich loop, the αC helix, and the activation loop. In summary, the developed macrocyclic MST3 inhibitor, JA310, demonstrates the utility of macrocyclization for the design of highly selective inhibitors and presents a first chemical probe for MST3.

## Introduction

The mammalian sterile 20-like serine/threonine protein kinases (MSTs) comprise five related proteins, called MST1 (STK4), MST2 (STK3), MST3 (STK24), MST4 (STK26), and YSK1 (STK25), involved in cell proliferation, cell migration, and cell polarity as well as tissue homeostasis.^1^ The MSTs are further divided into two subgroups, called GckII (geminal center subgroup II) and GckIII. MST1 and MST2, members of the GckII subgroup, are key components of the Hippo signaling pathway.^2^ MST3, MST4, and YSK1 belong to the GckIII subgroup and are involved in the regulation of the cytoskeleton and the Golgi apparatus.^3^ MST3 and MST4 share about 90% sequence identity in the kinase domain but less than 20% in their diverse C-terminal domains.^4^ MST3 is mainly localized in the cytoplasm and is expressed in various tissues, including the brain.^5,6^ Increased kinase activity of MST3 may be a consequence of activation by autophosphorylation^7^, and overexpression of MST3 is involved in the development of epilepsy^8^ and various cancer types, such as breast cancer,^9^ gastric cancer,^10^ or CNS tumorigenesis^11^, and high MST3 expression levels have been frequently linked to a poor prognosis. It plays an important role in diverse physiological events, such as cell morphology and cell-cycle progression. For instance, MST3 phosphorylates NDR1/2, promoting G1 progression by stabilizing c-MYC, thereby preventing p21 accumulation.^12^ Moreover, high MST3 levels are responsible for apoptosis in trophoblasts triggered by oxidative stress.^13^ Ultanir et. al. identified TAO1/2 as a target of MST3. Phosphorylation of TAO1/2 at T440/T475 promotes recruitment of myosin Va in dendrites, which is necessary for its dendritic localization, synapse formation, and development of dendritic spine.^6^ In addition, MST3 can be phosphorylated at S79 by CDK5, inhibiting RhoA activity by phosphorylating S26, a key event regulating neuronal migration and actin dynamics in neurons.^11,14^ FAM40A, a component of the STRIPAK complex, negatively regulates MST3 by triggering its dephosphorylation. This complex is associated with breast cancer cell migration and metastasis, with FAM40B mutations preventing this interaction, resulting in a high MST3 activity enhancing cell migration.^15^

The closely related kinase MST4 is another possible interaction partner of the STRIPAK complex. MST4 is found in various tissues and is highly expressed in the brain, thymus, placenta, and peripheral blood leukocytes.^16,17^ It is responsible for many physiological functions, such as Golgi reorientation, epithelial cell brush border formation, and inflammatory response.^18^ Dysfunction of MST4 has been observed in various cancers such as breast cancer,^15^ pancreatic cancer,^19^ prostate cancer,^1^ and hepatocellular carcinoma cells (HCC). Hao *et. al.* for example have shown that MST4 suppresses cell proliferation in HCC by inactivating the PI3K/AKT pathway, leading to cell-cycle arrest in G1 phase. ^20^ In HCC, MST4 is downregulated, and expression levels negatively correlate with disease progression and poor prognosis.^20^ Interestingly, the transcription factor SNAIL1, which regulates epithelial to mesenchymal (EMT) transition during embryonic development, is an MST4 substrate, which is suppressed by MST4. Inhibited SNAIL1 expression leads to a reduced potential of EMT invasion and metastasis of HCC cells.^18^ Upon oxidative stress, MST4 is activated, then translocates and phosphorylates ERM family proteins to promote cell survival.^21^ Moreover, MST4 is involved in enhanced autophagy flux in glioblastomas triggered by phosphorylation of ATG4B at S383.^22^ It is also part of the non-canonical Hippo signaling pathway.^23^ Thus, the diverse functions of MST3 and MST4 as well as other MST family members that have not been introduced here suggest that chemical probes targeting this family should achieve isoform selectivity in order to functionally differentiate between the diverse roles of individual family members in regulating cellular signaling.

In recent years, macrocyclization of acyclic analogs has gained increasing interest in drug discovery. To date, analogs from natural products as well as synthetic macrocycles have been used.^24^ For the synthesis of such macrocycles, a linear ATP-mimetic pharmacophore is usually used and connected via a linker motif, resulting in a macrocycle that by definition contains 12 or more atoms. Here, the linker has a significant role in the orientation of the acyclic pharmacophore in the targeted binding pocket, so that by macrocyclization through the variation of different linkers, one can easily influence the biological and also physiochemical properties compared to the acyclic precursor.^25,26^ The potency of macrocycles is typically increased compared with their acyclic analogue due to the lower entropic cost of binding, and also the selectivity over related off-targets can be increased by limiting conformational flexibility.^27^ In addition, macrocyclization can also optimize other pharmacokinetic and physiochemical properties such as metabolic stability, solubility, or cell permeability.^28^ For the development of new ATP-mimetic inhibitors that target the highly conserved active site of the catalytic kinase domain, selectivity remains the main challenge. Macrocycles represent an opportunity for the design of a new class of type I/II inhibitors with optimal shape complementarity with the ATP binding site.

The promiscuous kinase inhibitors **1** and **2** (**Figure 1D**) have been published in 2008 by Statsuk *et. al.* A kinome-wide screen of **1** against 468 human protein kinases revealed a pronounced ligand promiscuity by targeting 262 kinases (>35% at 1 µM).^29^ The crystal structures of **1** and **2** in complex with VRK1 and c-SRC are shown in **Figure 1B** and **C**, respectively, suggesting at least two diverse binding modes also based on alternative H-bond donors and acceptors interacting with the kinase hinge region. The pyrimidine of **1** faced the hydrophobic back pocket, and the 3-aminopyrazole moiety interacted with D132 and F134 of the kinase hinge backbone. The crystal structure of **2** revealed a flipped binding mode in which the quinazoline moiety targeted the solvent-exposed region, while the 3-aminopyrazole moiety interacted with the hinge backbone, forming three hydrogen bonds.^29^ The flexibility of these molecules and the large number of potential hydrogen bond acceptors and donors explained that these ligands potently interact with diverse kinases. Our hypothesis was that fixing the bioactive conformation by macrocyclization would be an excellent strategy to achieve target selectivity.

**Figure 1:**
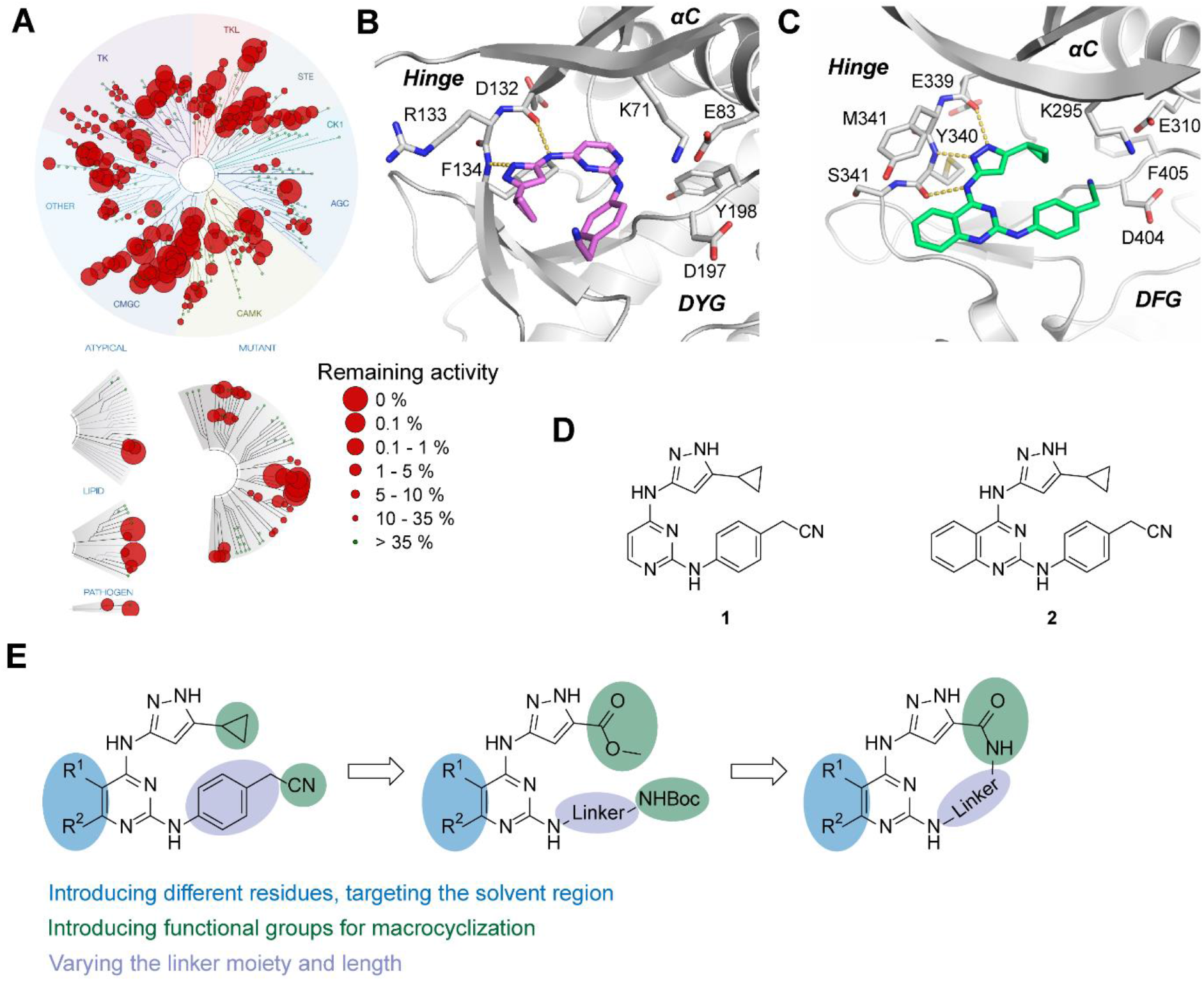
**A**. Selectivity profile of **1** assessed by screening against 468 recombinant human protein kinases at a concentration of 1 µM, using the KINOMEscan assay platform. **B.** Crystal structure of VRK1 in complex with **1** (PDB: 3OP5). **C.** Crystal structure of c-SRC in complex with **2** (PDB: 3F6X). **D.** Chemical structures of the previously published promiscuous kinase inhibitors **1** and **2**.^29^ **E.** Synthetic strategy for the design of macrocyclic inhibitors starting from the promiscuous kinase inhibitor **1**.

Here we report the design, synthesis, and characterization of a new chemical probe based on the N^4^-(*1H*-pyrazol-3-yl)pyrimidine-2,4-diamine core of the promiscuous inhibitors **1** and **2**. As outlined in **Figure 1E**, functional groups for macrocyclization were introduced together with diverse linker moieties, resulting in a comprehensive series of inhibitors for SAR (structure activity relationship) evaluation.

## Results and Discussion

### Synthesis of macrocyclic kinase inhibitors

Macrocycles **19** – **21** were synthesized in a 5-step synthetic route, as shown in Scheme 1. The commercially available methyl 3-amino-1*H*-pyrazole-5-carboxylate was reacted with the corresponding 2,4-dichloropyrimidine via a nucleophilic substitution under basic conditions to obtain compounds **7** – **9** with yields ranging from 7% to 84%. Different substituents at the pyrimidine on position 5 and 6 were explored to target the solvent region. A second nucleophilic substitution enabled derivatization by introducing diverse linker moieties. Aliphatic linkers were attached under basic conditions and microwave irradiation, while different aniline derivatives were introduced with a catalytic amount of 1M HCl. The yields ranged from 5% to 72%. The Boc-protecting group was cleaved with TFA in DCM, followed by a saponification with lithium hydroxide to afford the precursors **16** – **18**. Macrocyclization in the last step was carried out via amide coupling using HATU as a coupling reagent to obtain the macrocycles **19** – **21** with yields ranging from 2% to 42%.

**Scheme 1:**
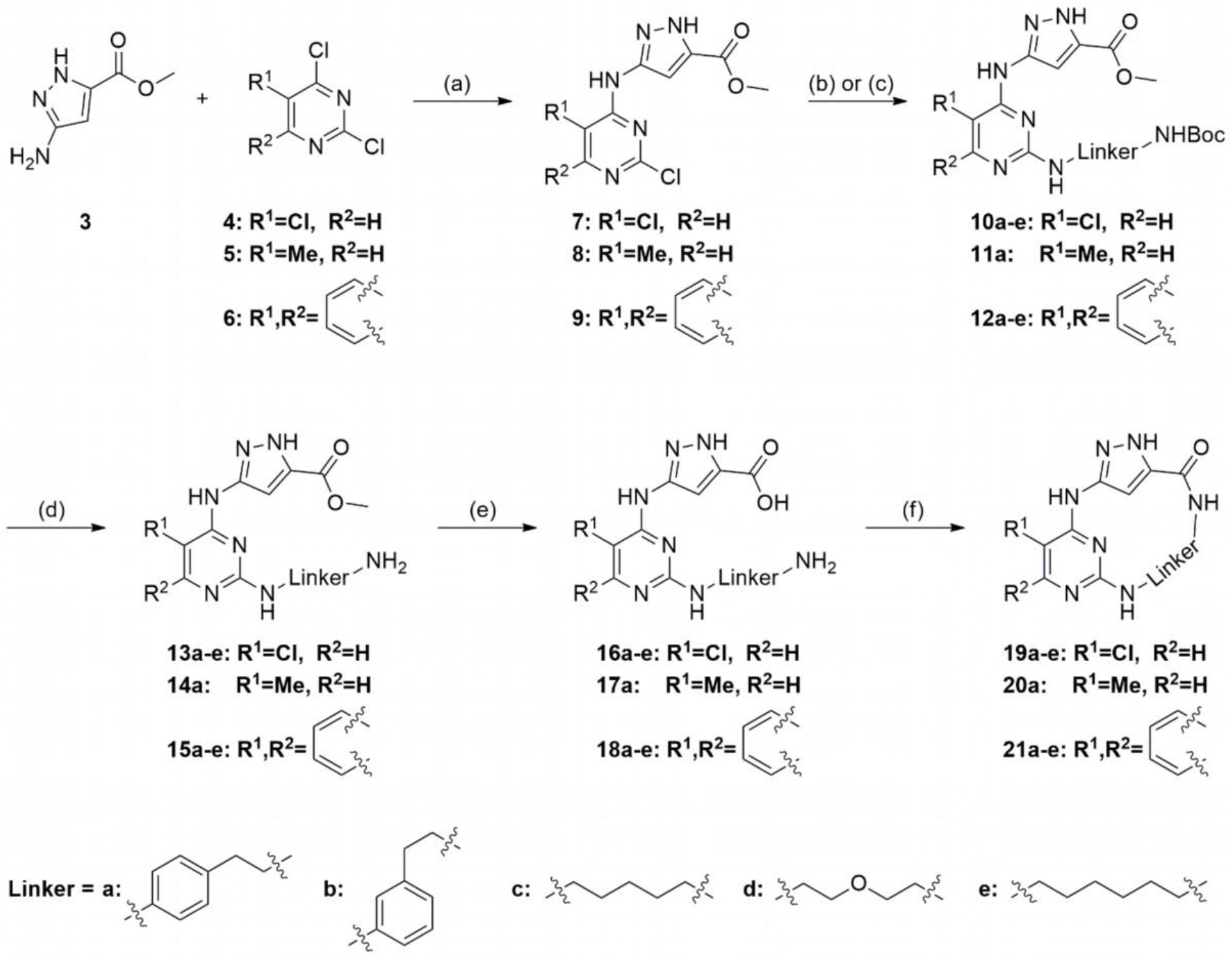
Synthesis of the macrocycles **19a-e**, **20a**, and **21a-e**.^a^ Reagents and conditions: (a) TEA and isopropanol, 24–72 h, 50–60 °C; (b) TEA and ethanol, microwave, 3–8 h, 80–120 °C; (c) HCl and ethanol, 70 °C–reflux, 18 h; (d) TFA and DCM, 0°C – rt, oN; (e) LiOH*H_2_O, THF and H_2_O, 18 h, 55 °C; (f) HATU, DIPEA and DMF, 18 h, rt–60°C.

### Structure-activity relationship of 3-aminopyrazole-based macrocycles

A differential scanning fluorimetry (DSF) assay was performed to assess the selectivity profile of the synthesized macrocycles. This assay represents a rapid and sensitive screening method that determines the melting temperature of a protein in the absence and presence of a compound.^30^ In this assay format, the difference between the melting temperature (Δ*T*_m_) in the absence and presence of the compound can be correlated with the potency of the ligand for a given kinase target. For screening all synthesized macrocycles, we used an established screening panel of 104 recombinant kinases, bromodomains, and typical off targets (Table S1 & S2). Lead structure (**1**) was resynthesized based on the synthetic route of Statsuk *et. al*.^29^ Using the DSF assay, we identified a highly potent molecule that showed high Δ*T*_m_ values for MST3 and MST4. In the publication by Statsuk et al., the lead structure (**1)** bound equally well to the MST family members MST1 to MST4. All four kinases were effectively targeted, with residual kinase activity ranging from 0.15% – 3.2% binding at 10 µM.^29^ The same trend was observed in our data presented here. MST1 and MST2 were stabilized by **1** with thermal shifts of 9.6 °C and 11.6 °C, while MST3 and MST4 showed Δ*T*_m_ values of 9.3 °C and 7.1 °C, respectively (**Figure 2A**).

**Figure 2.**
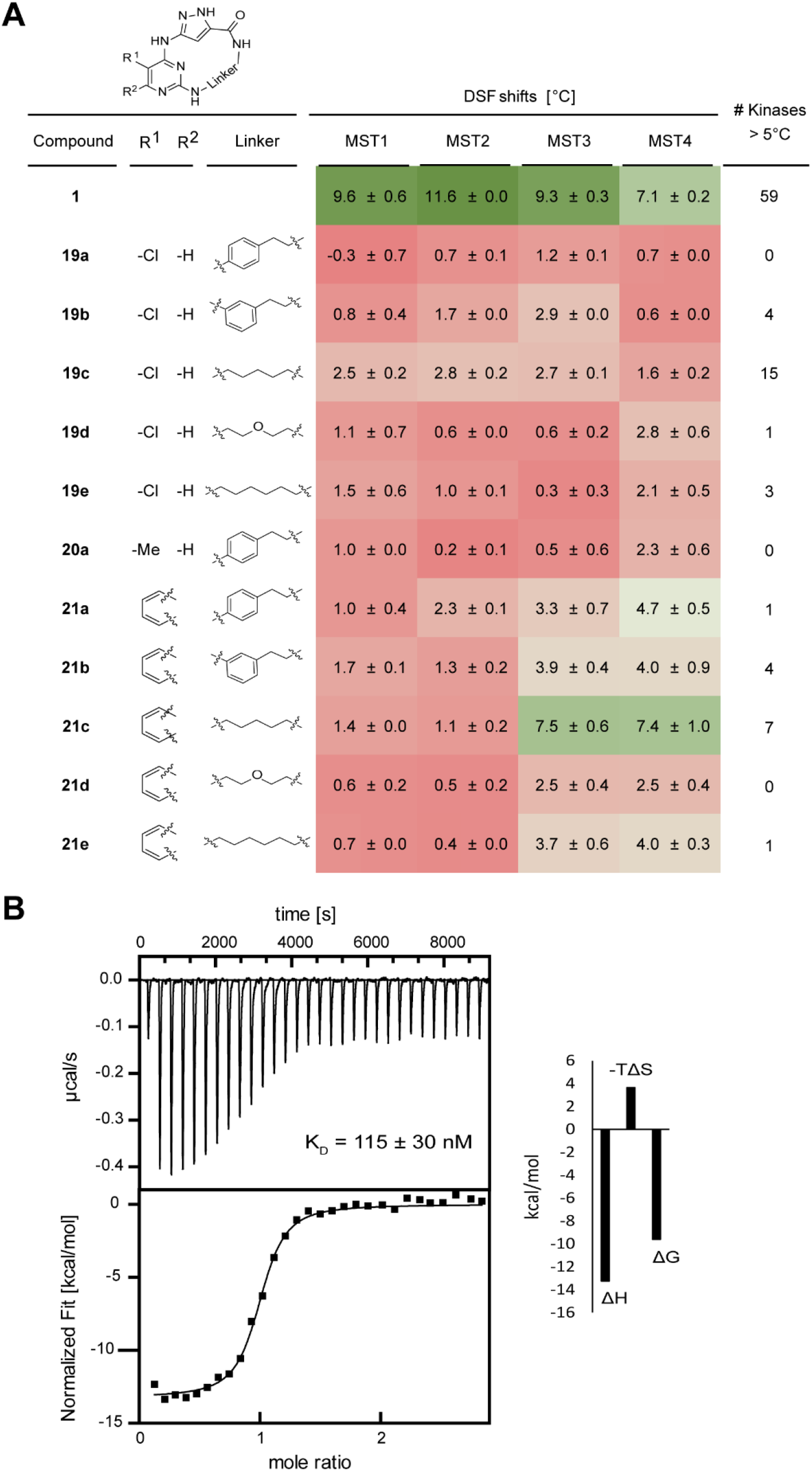
**A**. DSF thermal shift data for macrocycles **19** – **21** against MST1 – MST4. Red indicates a low stabilization of the targeted kinase and green a high stabilization. Measurements were performed in technical duplicates. **B.** ITC measurement of **21c** against MST4 revealed a high binding affinity with a K_D_ value of 115 nM.

Macrocycles **19a**-**e**, which harbor a chlorine moiety at the C5 position of the pyrimidine, showed no significant stabilization of the MSTs, with Δ*T*_m_ shifts smaller than 3 °C. Compounds **19a**, **19b**, **19d**, and **19e** stabilized only a few kinases with a thermal shift larger than 5 °C, which was negligible compared to the positive control staurosporine. **19c** appeared to be rather unselective, stabilizing 15 kinases > 5 °C. Also, the replacement of the chlorine moiety by a methyl group (**20a**) was not tolerated by any kinase of our screening panel. **20a** did not stabilize any kinase > 5 °C, and the shifts for MST3 and MST4 were low (0.5 °C and 2.3 °C, respectively). Introduction of the quinazoline moiety instead of the pyrimidine increased Δ*T*_m_ values for MST3 and MST4, while the shifts for MST1 and MST2 remained below 3 °C (**21a** vs **19a**). The macrocycles **21a** and **21b**, harboring an aromatic ring in the linker, led to a moderate stabilization, with Δ*T*_m_ values between 3.3 °C and 4.7 °C. Both compounds showed few Δ*T*_m_ shifts larger than 5 °C, however, these were relatively weak compared with those obtained with the non-cyclic control compound. **21c** led to a high stabilization of MST3 and MST4, with Δ*T*_m_ values of 7.5 °C and 7.4 °C, respectively, which was in the same range or even more potent than with staurosporine (7.3 °C and 6.1 °C). Reported K_D_ values for staurosporine are 120 nM and 140 nM.^31^ MST1 and MST2 were not significantly affected in DSF, with Δ*T*_m_ shifts of only 1.1 °C and 1.4 °C. Outside the MST family, **21c** stabilized five other kinases (CLK1, GSK3B, MELK, RIOK1, STK6) in a range of 5.1 °C – 6.5 °C. The replacement of the pentyl linker with an ethoxyethyl (**21d**) or hexyl (**21e**) linker decreased the shifts for MST3 and MST4. Both kinases were only mildly stabilized, with shifts of 2.5 °C – 3.7 °C and 2.5 °C – 4.0 °C, respectively. **21d** was completely inactive, while **21e** stabilized only GSK3B with a Δ*T*_m_ > 5 °C.

To verify the stabilization of MST4 by **21c**, an ITC measurement was performed, revealing a strong binding affinity with a low K_D_ value of 115 nM (**Figure 2B**). Binding of **21c** to the protein was driven by a favorable change in the binding enthalpy (ΔH), however, still off-set by a binding entropy change (-TΔS). Because of the high stabilization of MST3 and MST4 in the DSF assay and the good *in vitro* potency of **21c** in the ITC measurement, we decided to proceed characterizing this compound further in cellular assays.

### NanoBRET target engagement assays

The cellular target engagement potency of the 3-aminopyrazole-based macrocycles was determined against MST1 – MST4 using the NanoBRET assay.^32,33^ For MST3 and MST4, the measurements were also performed in the permeabilized mode for determining the *in vitro* potency. Lead structure (**1**) exhibited a high stabilization of all MSTs in DSF assays. However, it showed preferential binding of MST1 and MST2 in cellular NanoBRET assays, with EC_50_ values of 200 nM and 110 nM, respectively in intact cells. In the permeabilized mode, compound **1** showed EC_50_ values of 280 nM and 230 nM for MST3 and MST4, respectively. The EC_50_ values in intact cells, however, were in the micromolar range. The macrocycles bearing a chlorine moiety (**19a**-**e**) or a methyl group (**20a**) at the C5 position of the pyrimidine did not show stabilization of the MSTs in the DSF assay. These results agreed well with the NanoBRET data, which showed either weak binding in the micromolar range or complete loss of binding up to a concentration of 50 µM. The switch from a pyrimidine to a quinazoline moiety increased the potency of the macrocycles against MST3 and MST4, while the activity against MST1 and MST2 remained unaffected. Macrocycles **21a** and **21b**, which contained an aromatic moiety in their linker region, still showed weak potency in NanoBRET assays (4.5 µM – 17.5 µM) and a slightly increased potency against MST3 and MST4 in the permeabilized cells (0.9 µM – 2.8 µM), suggesting cell penetration is limiting the potency of these inhibitors. In contrast, further replacement of the linker to a pentyl linker in compound **21c** resulted in strong binding to MST3, with EC_50_ values of 106 nM and 76 nM for intact and permeabilized cells, respectively. Compound **21c** showed comparatively weaker activity for MST4, with EC_50_ values of 1.4 µM and 362 nM for intact and permeabilized cells, respectively. The two related family members MST1 and MST2 were not affected (EC_50_ values >50 µM). The ethoxy linker in **21d** decreased the potency against MST3 and MST4 (EC_50_ values 1.2 µM – 37.5 µM for intact and permeabilized cells). In addition, extension of the linker by one carbon atom (**21e**) caused attenuation of the efficacy, with EC_50_ values of 1.8 µM and 2.4 µM for intact cells and 450 nM and 1.3 µM for the permeabilized mode, respectively (Table 1).

**Table 1.**
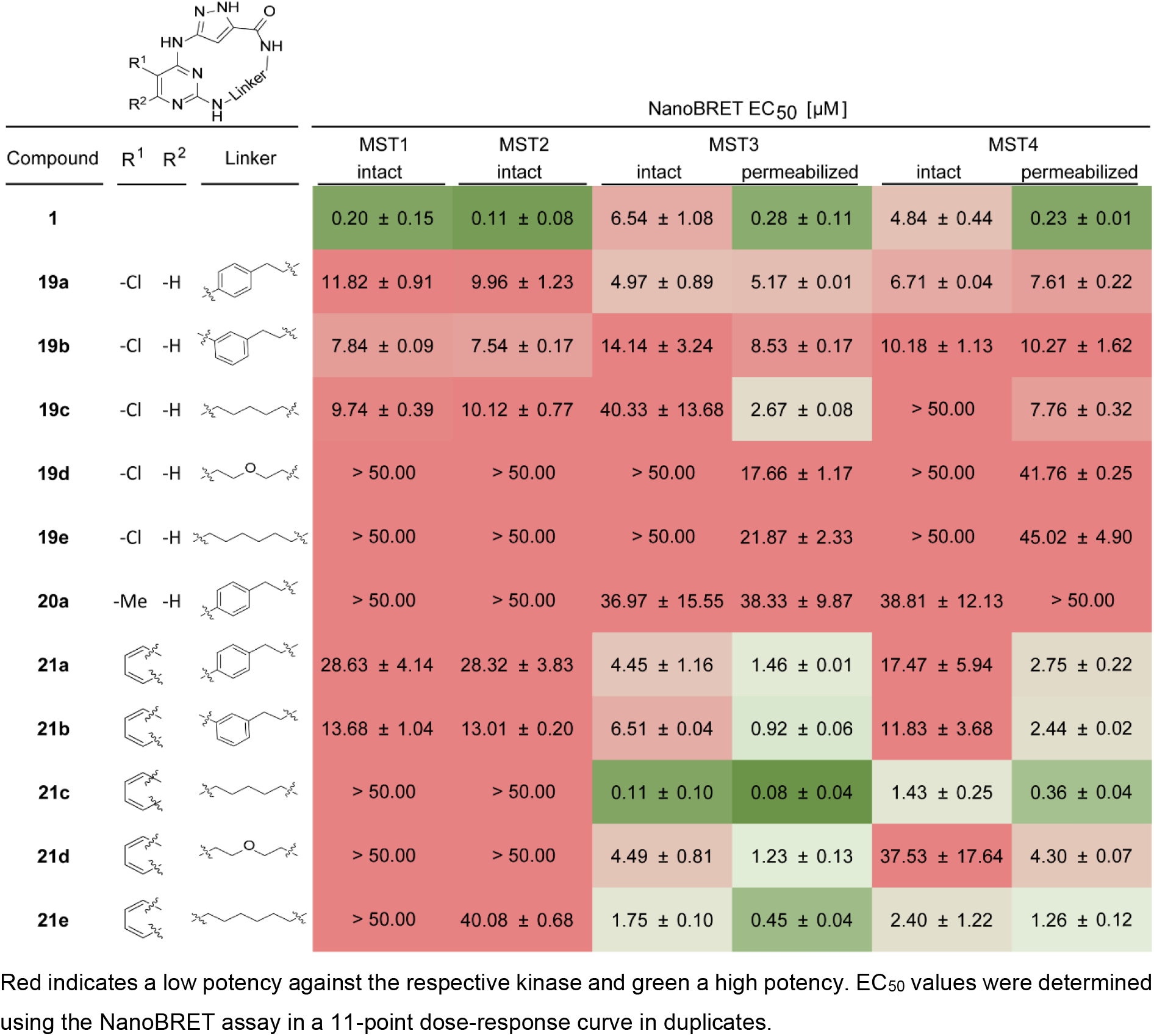
NanoBRET data for the 3-aminopyrazole-based macrocycles **19** – **21** against MST1 – MST4.

### Extended screening revealed kinome-wide selectivity of 21c

Based on the activity on MST3 and MST4 in the DSF assay, the excellent cellular potency for MST3 in the NanoBRET assay, and the clean selectivity profile in our DSF panel, we selected macrocycle **21c** as a chemical probe candidate, which we called JA310, and profiled this inhibitor further on a comprehensive screening panel of wild-type kinases (Reaction Biology). JA310 (**21c**) was tested against 340 wild-type kinases at a screening concentration of 1 µM (**Figure 3A**), revealing excellent selectivity with a selectivity score (S_40_) of 0.012 (**Figure 3B**). Gratifyingly, the most potently inhibited kinases were MST3 and MST4, with a significant selectivity window to the closest off-targets in this radiometric kinase screening panel. To validate the encouraging selectivity data of JA310 (**21c**), NanoBRET assays were performed in intact and in permeabilized cells to determine the EC_50_ values for all off-targets with percent of control values below 40% (**Figure 3C**). These results confirmed the excellent selectivity profile of JA310 (**21c**) as MST3 was the only kinase that was inhibited in the nanomolar range in cellular assays. MST4 was inhibited about 14 times less potently by macrocycle JA310 (**21c**). The next identified off-target outside the MST family was LIMK2, which was inhibited with EC_50_ values of 1.4 µM and 1.8 µM in intact and permeabilized cells. Surprisingly, LIMK1 was only weakly inhibited in cellular NanoBRET assays.

**Figure 3.**
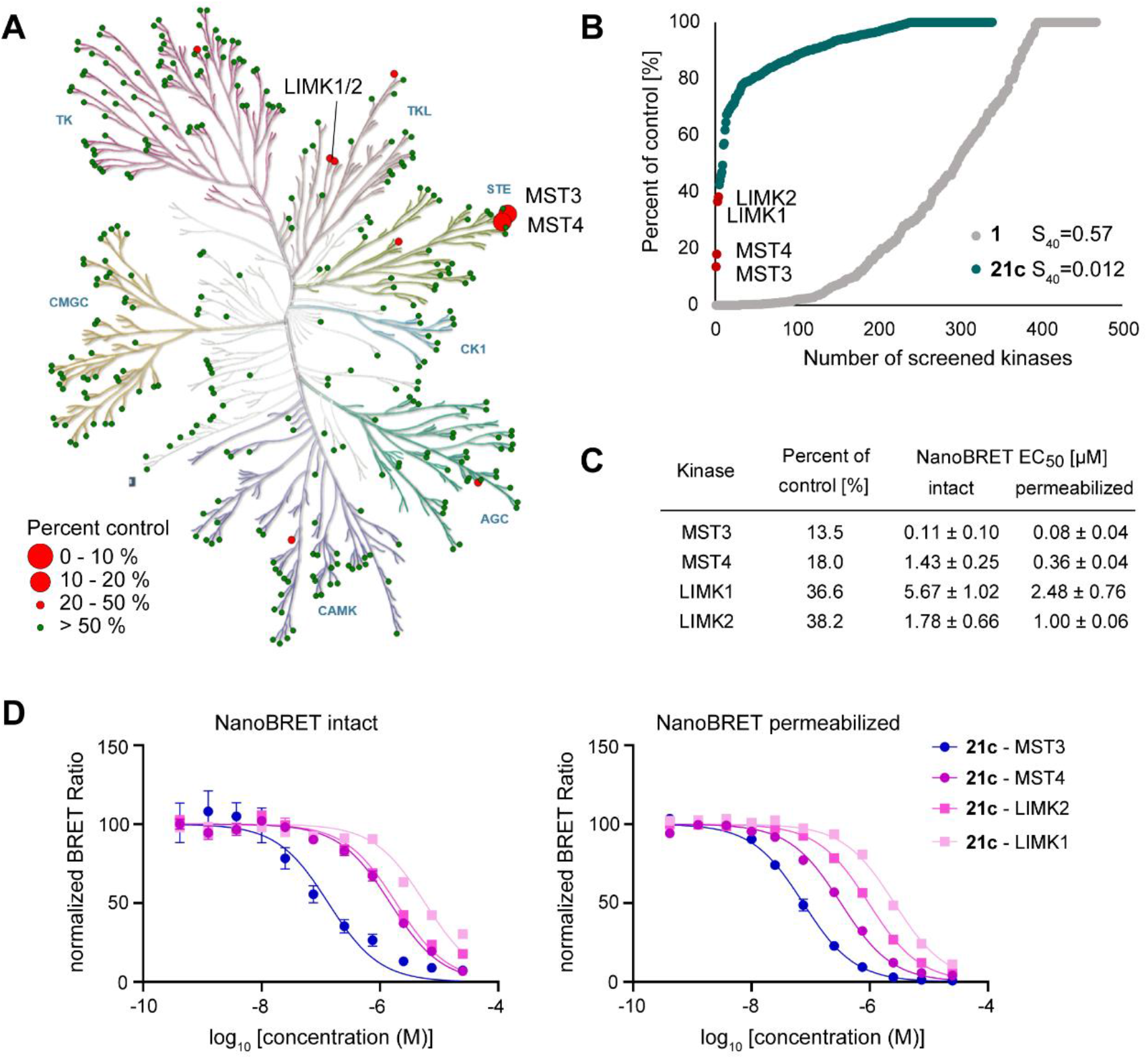
**A**. Selectivity profile of JA310 (**21c**) assessed by the wild-type kinase panel provided by Reaction Biology at a screening concentration of 1 μM. **B.** Waterfall plot of the selectivity data of **1** (gray) and **21c** (green). The remaining activity [%] was plotted against the number of screened kinases. **1** was screened against a panel of 468 kinases using the KINOMEscan (Eurofins/ DiscoverX at 1µM) platform. **21c** was screened against a panel of 340 kinases using the wild-type kinase panel provided by Reaction Biology. The selectivity score was determined with a cutoff value of 40% remaining activity. Kinases that showed larger than 40% inhibition for **21c** are highlighted as red dots. **C.** Table showing the top hits of the kinome profiling of **21c** with a remaining activity < 40% at 1 μM and the corresponding EC_50_ values determined by NanoBRET. **D.** Dose response titrations of NanoBRET data of **21c** for the kinases MST3, MST4, LIMK1, and LIMK2.

### Binding of JA310 (21c) induced large-scale structural rearrangements

To elucidate the binding mode of the new series of 3-aminopyrazole-based macrocycles, JA310 (**21c**) was co-crystallized with MST3, and the structure of the complex was determined at 1.64 Å resolution. Intriguingly, superimposition of this structure onto the structure of the MST3 AMP-PNP complex (PDB: 4QML),^34^ the apo structure (PDB: 3CKW), or previously published MST3-inhibitor complexes^34–36^ revealed a large-scale induced-fit movement upon macrocycle binding. This movement was characterized by a rotation and tilting of the upper lobe toward the C-lobe (**Figure 4A**), with large shifts of some of the β-strand backbone atoms by up to 8 Å and reorientation of the αC helix (including disruption of the K65-E82 salt bridge). This also drastically altered the shape and architecture of the binding site and adjacent structural motifs. The side chain of F47 in the glycine-rich loop, for example, which docks against the αC helix in the complex with the ATP analog, was shifted by almost 8 Å (CZ atom) and packed against L181 and T182 of the activation loop, resulting in a complete structural reorganization of the latter with associated changes in topology and shifts of backbone atoms of up to 20 Å compared with the conformation stabilized in its phosphorylated state (**Figure 4B**). The typically formed small β-sheet, which is seen in the ATP analog complex around G177, was disrupted in the complex with the macrocycle, and an additional short α-helical segment formed extending from residues 183-188. There was clear electron density for the whole activation loop, whereas this region was largely unresolved (indicating high conformational flexibility) in previously published inhibitor structures of unphosphorylated MST3 kinase domain.^35,36^ An animation of the structural rearrangements has been uploaded as a supporting video.

**Figure 4.**
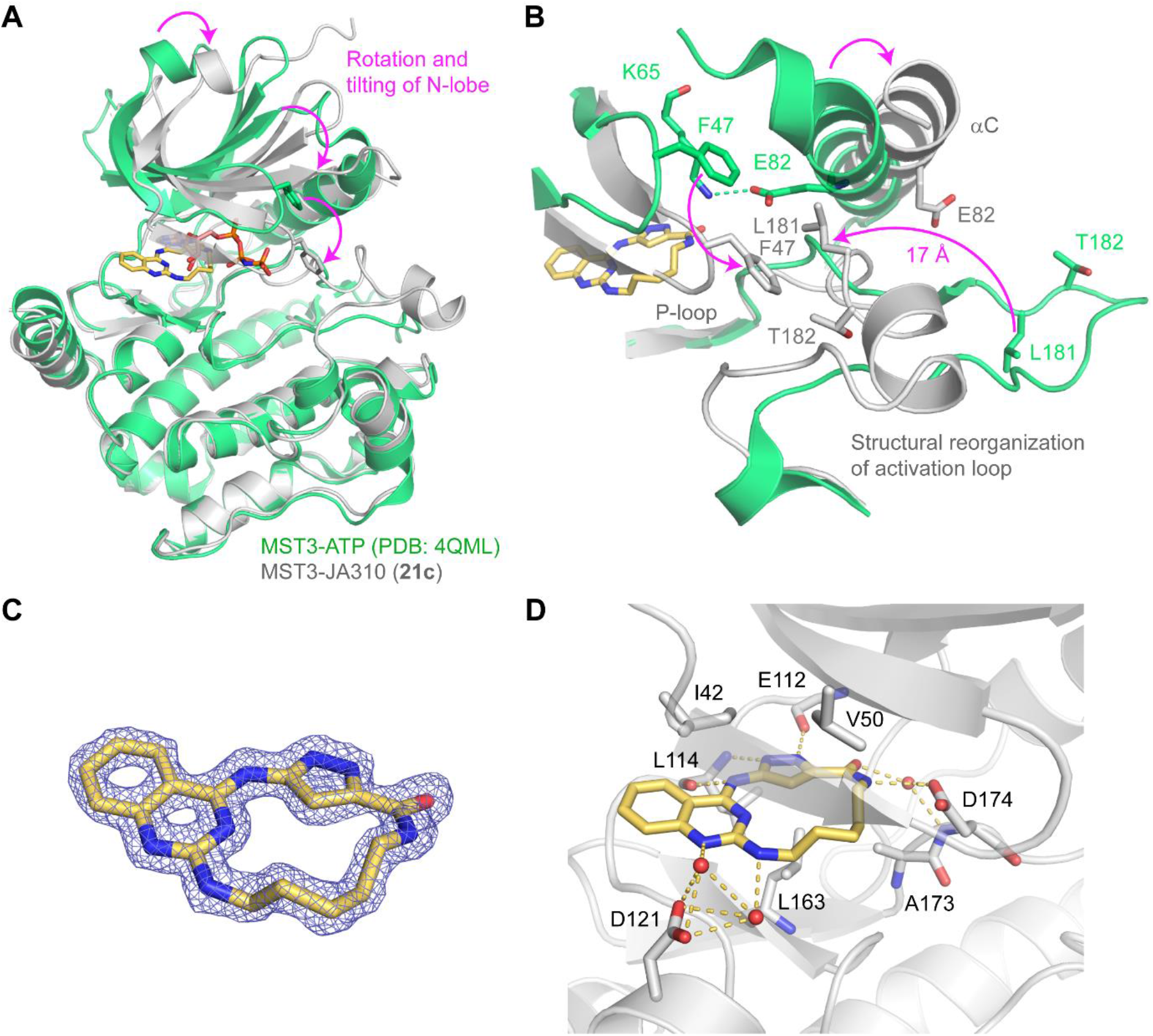
Crystal structure of the MST3 kinase domain with bound inhibitor JA310 (21c). **A.** Superimposition of the structure of the MST3-JA310 complex (green) and MST3 with a bound ATP analog (gray; PDB entry 4QML), revealing a substantial rotation and tilting of the N-lobe relative to the C-lobe upon binding of the inhibitor. **B.** Close-up view of the superimposition shown in A highlighting the concerted induced-fit movement involving the P-loop, the αC helix, and the activation loop, with a large-scale structural rearrangement and change in topology of the latter. Residue numbering is according to the canonical isoform in UniProt (Q9Y6E0-1), which is different from the numbering used in PDB entry 4QML, which is based on the shorter isoform 2 that is produced by alternative splicing. **C.** 2F_o_-F_c_ electron density map around the ligand shown at a contour level of 2.0 σ. **D.** Detailed binding mode of JA310 (yellow stick model). Key interacting residues in MST3 and structural water molecules interacting with the ligand are shown; inhibitor-mediated hydrogen bonds are highlighted as orange dashed lines.

The detailed binding mode of JA310 (**21c**) is shown in **Figure 4D**. As expected, the 3-aminopyrazole moiety interacted with the hinge region, forming three hydrogen bonds with the backbone of E112 and L114. The quinazoline moiety faced the entrance of the binding pocket, with the N1 nitrogen and the amine at the C2-position engaging in water-mediated hydrogen bonds with the carboxylate group of D121. The linker was fully resolved in the crystal structure (**Figure 4C**) and formed specific stabilizing interactions. Its amide group formed a hydrogen bond with the carboxylate group of D174 in the DFG motif (DFG-in conformation), which in turn formed a salt bridge with K65. The aliphatic moiety of the linker was stabilized by hydrophobic interactions on both sides of the ring, involving V50, L163, and V173, hence directly contributing to the potency of the inhibitor. The side chain of V50 was in direct contact with that of I42, which formed tight hydrophobic interactions with the quinazoline ring. Interestingly the latter residue is replaced by a leucine in family members MST1/2, which is bound to alter the surface complementary and may at least in part contribute to the selectivity of JA310 (**21c**) for MST3 over those family members. The exquisite kinome-wide selectivity overall may in fact also be linked to the large structural reorganization occurring in MST3 upon binding of the macrocycle, suggesting that residues beyond the immediate binding site that facilitate the concerted induced-fit movements may also be key for the observed selectivity.

### Metabolic stability of 21c

JA310 (**21c**) was tested against an activated liver microsome mix derived from Sprague-Dawley rats to determine the metabolic stability, as previously reported.^37^ The lead structure (**1**) was not included because of its toxicity based on its promiscuous kinase selectivity profile. JA310 (**21c**) was incubated at 37 °C for 60 min, and the amount of un-metabolized compound was determined every 15 min by high-performance liquid chromatography (HPLC). JA310 (**21c**) showed a good microsomal stability with 88% of un-metabolized compound after 60 min (**Figure 5A**).

**Figure 5.**
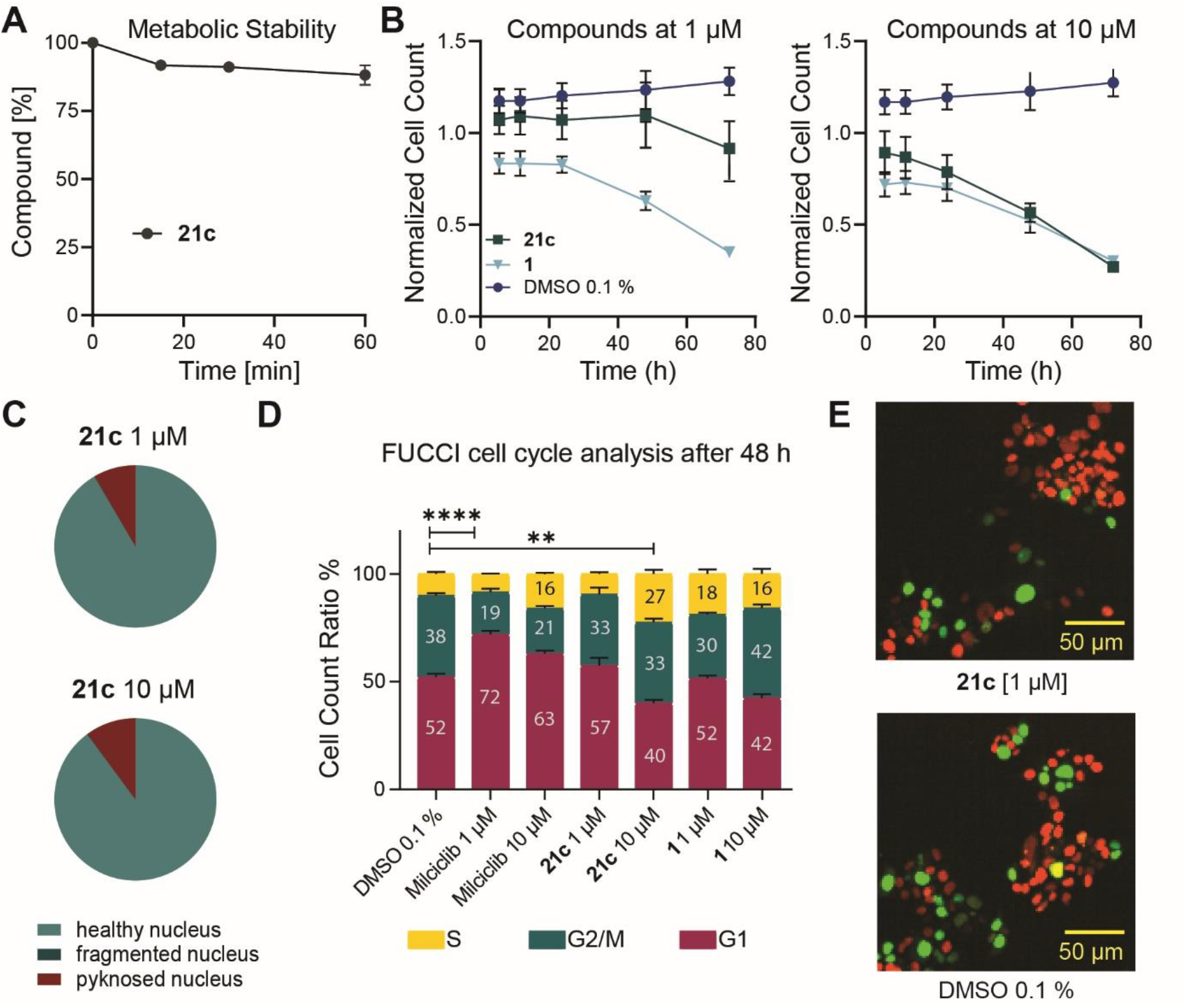
Metabolic stability, cell-cycle analysis (FUCCI), and viability assessment. **A.** Metabolic stability of JA310 (**21c**) [10µM] after treatment for 60 min. The residual amount of compound in percent was plotted against the time in min. **B.** Normalized cell count of HCT116-FUCCI cells after 6 h, 12 h, 24 h, 48 h and 72 h at 1 µM and 10 µM of compound exposure (**21c**, **1**) compared with the DMSO control. Error bars show SEM of two biological replicates. **C.** Fraction of healthy, fragmented and pyknosed nuclei after 24 h at 1 µM and 10 µM compound exposure (**21c**) in HCT116-FUCCI cells. Average data of two biological duplicates are shown. **D.** Fractions of red (G1), green (G2/M) or yellow (S) cells after 72 h of compound exposure (**21c**, **1**, milciclib). Numbers show percentages (%) of fractions. Error bars show SEM of two biological replicates. Significance was calculated using a two-way ANOVA analysis (****p<0.0001, **p<0.0021). **E.** Fluorescence Image of HCT116-FUCCI cells after 72 h exposure to 1 µM of compound (**21c**) compared with DMSO control.

### Toxicity and cell-cycle analysis

MST3 has been shown to play a pivotal role in mitotic cell-cycle progression through activation of NDR1/2 kinases, which can control G1/S progression.^38^ Downregulation or inhibition of MST3 and MST4 can lead to cell-cycle arrest. Given the link to cell-cycle control, we performed a live-cell assay using the fluorescent ubiquitination-based cell-cycle indicator (FUCCI) system to detect different cell-cycle states at the single-cell level.^39^ Human colorectal carcinoma cells (HCT116) stably expressing the FUCCI system were additionally stained with Hoechst33342 as a nuclear marker to assess viability as described by Tjaden *et. al.*^40^ Compound **21c** showed no effect on cell viability over 48 h at 1 µM (**Figure 5B**). At the highest concentration tested (10 µM), the normalized cell count decreased in a time-dependent manner to less than 30% after 72 h. 80% of the remaining cells showed healthy nuclei (**Figure 5B** and **5C**) after 24 h for both concentrations. In contrast, the lead structure (**1**) decreased the normalized cell count at all concentrations tested (1 µM, 5 µM, 10 µM) to less than 50% after 72 h. The cell-cycle analysis revealed that compound **21c** leads to a small increase of cells in G1 and reduction of S phase cells, consistent with the effect observed inhibiting MST3.^38^ However, at 10 µM this effect was no longer detectable and instead a significant number of cells were in the S phase compared with cells treated with the DMSO control (**Figure 5D**). The reduction of cells observed in G1 at 10 µM may be due to simultaneous inhibition of MST3 and MST4, which have opposing effects in G1, whereas at 1 µM only a minor effect on the cell cycle (increase in G1) was detectable in line with the effect of inhibition of MST3 (**Figure 5E**).^17,38^ In addition, the lead compound **1** increased the number of cells in G2/M at the highest concentration tested (10 µM), while no effect on the cell cycle was detected at 1 µM.

## Conclusion

In this study, we developed a new series of 3-amino-*1H*-pyrazole-based macrocycles, based on the scaffold of the highly promiscuous kinase inhibitor **1**. Macrocyclization has been used to minimize/eliminate off-targets through conformational constrains introduced by the linkers used for cyclization, which may lock the bioactive conformation of an inhibitor for a certain kinase, resulting often in improved selectivity. The developed kinase inhibitor JA310 (**21c**) was recently accepted as a chemical probe and shows an excellent kinome-wide selectivity and an *in vitro* potency of 76 nM and a cellular EC_50_ value of 106 nM for MST3. The highly similar family member MST4 was the next detected off-target (cellular EC_50_ = 1.4 μΜ). Also, LIMK1/2 were inhibited in the low micromolar range, therefore at higher compound concentrations (>2 μM), it is likely that these kinases will also be significantly inhibited. The detected G2M arrest at high compound concentration (10 μM), but not at 1 μM, may be due to inhibition of these off-targets. Macrocyclization is an excellent method achieving high kinome-wide selectivity which we and others demonstrated on several examples,^24,41–43^ including exclusive selectivity for oncogenic mutants over the wild-type protein.^44^ The kinome-wide selectivity we achieved for JA310 (**21c**) is remarkable compared with the promiscuous lead compound **1** used as starting point. Several factors may have contributed to this excellent selectivity profile. The insertion of a bulky quinazoline moiety substituted at the 2-position instead of the amino pyrimidine moiety in **1** most likely restrained the binding mode of JA310 (**21c**) to the 3-aminopyrazole moiety hinge interaction. Furthermore, large structural rearrangements were observed in the MST3-JA310 (**21c**) complex, locking the kinase in an inactive conformation characterized by rotation and tilting of the upper lobe and helix αC as well as structural rearrangement of the activation segment. JA310 (**21c**) induced a highly distorted catalytic domain conformation that may be exclusive to MST3 and the closely related MST4. LIMK1/2 may also assume the observed distorted kinase domain conformation as these kinases have been described to be highly dynamic, allowing also the development of back-pocket binding allosteric kinase inhibitors.^45^ In addition, JA310 (**21c**) showed good metabolic stability, and this chemical probe therefore provides a suitable chemical tool for studying the role of MST3 in cellular systems.

## Experimental Section

### Differential Scanning Fluorimetry Assay

Recombinant protein kinase domains with a concentration of 2 μM were mixed with a 10 μM compound solution in DMSO, 20 mM HEPES, pH 7.5, and 500 mM NaCl. SYPRO Orange (5000×, Invitrogen) was added as a fluorescence probe (1 µL per mL). Subsequently, temperature-dependent protein unfolding profiles were recorded using the QuantStudio™ 5 realtime PCR machine (Thermo Fisher). Excitation and emission filters were set to 465 nm and 590 nm. The temperature was increased at a heating rate of 3 °C per minute. Data points were analyzed with the Thermal Shift Software Version 1.4, (Thermo Fisher) using the Boltzmann equation to determine the inflection point of the transition curve. Differences in melting temperature are given as Δ*T*_m_ values in °C. Measurements were performed in technical duplicates.

### Kinome-Wide Selectivity Profile

Compound **21c** was tested at a concentration of 1 μM against a panel of 340 kinases in the wild-type kinase panel performed by Reaction Biology (Table S6).

### ITC Measurement

The K_D_ of the **21c** was determined by isothermal calorimetry. The sample cell, sample syringe, and injection syringe were equilibrated with gel filtration buffer. The sample cell was filled with compound solution (25 μM in gel filtration buffer), and the injection syringe was filled with protein solution (100 μM in gel filtration buffer). Measurements were performed using an Affinity ITC (TA-Instruments) at a temperature of 25 °C and a stirring rate of 250 rpm. MST4 expressed and purified as previously described^35^ was titrated into the compound solution (940 μL cell volume) with 8 μL per injection, except the first injection, which was 4 μL. The time between each injection was set to 300 s. The results were analyzed using the NanoAnalyze Software (TA instruments).

### NanoBRET^TM^ assay

The assay was performed as described previously.^32^ In brief: full-length kinases were obtained as plasmids cloned in frame with a terminal NanoLuc-fusion (*Promega*) as specified in Supplementary Table S3. Plasmids were transfected into HEK293T cells using FuGENE HD (*Promega*, E2312), and proteins were allowed to express for 20 h. Serially diluted inhibitor and NanoBRET™ Kinase Tracer (Supplementary Table S3) at a concentration determined previously as the Tracer K10 EC_50_ (Supplementary Table S3) were pipetted into white 384-well plates (*Greiner* 781207) using an Echo acoustic dispenser (*Labcyte*). The corresponding protein-transfected cells were added and reseeded at a density of 2 x 10^5^ cells/mL after trypsinization and resuspending in Opti-MEM without phenol red (*Life Technologies*). The system was allowed to equilibrate for 2 h at 37 °C/5% CO_2_ prior to BRET measurements. To measure BRET, NanoBRET™ NanoGlo Substrate + Extracellular NanoLuc Inhibitor (*Promega*, N2540) was added as per the manufacturer’s protocol, and filtered luminescence was measured on a PHERAstar plate reader (*BMG Labtech*) equipped with a luminescence filter pair (450 nm BP filter (donor) and 610 nm LP filter (acceptor)). Competitive displacement data were then graphed using GraphPad Prism 9 software using a normalized 3-parameter curve fit with the following equation: Y=100/(1+10^(X-LogIC_50_)).

### Protein Expression and Purification

MST3 (STK24) was expressed and purified as previously described.^35^ Briefly, MST3 (residues R4-D301) was cloned into vector pNIC-CH, resulting in the addition of a non-cleavable C-terminal 6xHis tag. The expression plasmid was transformed into BL21(D3)-R3-pRARE2 competent cells, and cell cultures were grown in TB media at 37 °C. After reducing the temperature to 18 °C, IPTG was added at a final concentration of 0.5 mM and cells were incubated overnight. The pelleted and resuspended cells were broken by sonication, and recombinant MST3 protein was purified using standard Ni-affinity chromatography protocols. The Ni-NTA fraction containing MST3 was concentrated using a 30 kDa cutoff ultrafiltration device and further purified by size exclusion chromatography (SEC; HiLoad 16/600 Superdex 200) pre-equilibrated with SEC buffer (25 mM HEPES pH 7.5, 200 mM NaCl, 0.5 mM TCEP, 5% glycerol). Quality control was performed by SDS-gel electrophoresis and ESI-MS (MST3: expected 34 507.7 Da, observed 34 506.9 Da).

### Crystallization

Co-crystallization trials were performed by incubating MST3 (11 mg/mL in 25 mM HEPES pH 7.5, 200 mM NaCl, 0.5 mM TCEP, 5% glycerol) with compound **21c** (final concentration of 3 mM) for 1h. MST3 crystals were then grown at 20 °C via the sitting drop vapor diffusion technique using a mosquito crystallization robot (TTP Labtech, Royston UK). Diffracting crystals were obtained at a roughly 1:2 ratio of protein:reservoir buffer (70 nL:130 nL; 14% PEG 6K, 0.1 M HEPES pH 7.2). Crystals were cryoprotected with reservoir buffer complemented with 23% ethylene glycol and flash frozen in liquid nitrogen. X-ray datasets were collected at 100 K at the Swiss Light Source (beamline X06SA/PXI). Data collection and refinement statistics are listed in Table S4. Diffraction data were integrated with the program XDS^46^ and scaled with AIMLESS,^47^ which is part of the CCP4 package.^48^ The structure was then solved by molecular replacement using PHASER^49^ with PDB entry 7B32 as a search model. Structure refinement was performed by iterative cycles of manual model building in COOT^50^ and refinement in PHENIX.^51^ The dictionary file for the macrocyclic inhibitor was generated using the Grade Web Server (http://grade.globalphasing.org). The geometry of the final model was validated using MolProbity.^52^

### Fucci cell-cycle assay

The influence of the compounds on the cell cycle was tested using a florescent ubiquitination-based cell-cycle indicator (FUCCI) assay as described previously.^40^ In Brief, HCT116-FUCCI cells, stably expressing the FUCCI system, introduced by the Sleeping Beauty transposon system,^40,53^ were seeded at a density of 1250 cells per well in a 384-well plate (Cell culture microplate, PS, f-bottom, µClear®, 781091, Greiner) in culture medium (50 µL per well) and stained additionally with 60 nM Hoechst33342 (Thermo Scientific). Fluorescence and cellular shape were measured before and after compound treatment for 72 h every 12 h using the CQ1 high-content confocal microscope (Yokogawa). Compounds were added directly to the cells at three different concentrations (1 µM, 5 µM, and 10 µM). The following parameters were used for image acquisition: Ex 405 nm/Em 447/60 nm, 500ms, 50%; Ex 561 nm/Em 617/73 nm, 100 ms, 40%; Ex 488 nm/Em 525/50 nm, 50 ms, 40%; Ex 640 nm/Em 685/40 nm, 50 ms, 20%; bright field, 300 ms, 100% transmission, one centered field per well, 7 z stacks per well with 55 µm spacing. Analysis of images was performed using the CellPathfinder software (Yokogawa). The cell count was normalized against the cell count of cells treated with 0.1% DMSO. Cells were gated in different categories as described previously by Tjaden et al.^54^ All normal gated cells were further classified in cells containing healthy, fragmented or pyknosed nuclei. Cells that showed a healthy nucleus were gated in red, green or yellow based on 11 features of the cell body and 4 features of the cell nuclei. Error bars show SEM of biological duplicates. Significance was calculated using a two-way ANOVA analysis in graph pad prism 8.4.3. All data can be found in Table S5.

### Microsomal stability assay

The solubilized test compound (5 μL, final concentration 10 µM) was preincubated at 37 °C in 432 μL of phosphate buffer (0.1 M, pH 7.4) together with 50 μL NADPH regenerating system (30 mM glucose 6-phosphate, 4 U/mL glucose 6-phosphate dehydrogenase, 10 mM NADP, 30 mM MgCl_2_). After 5 min, the reaction was started by the addition of 13 μL of microsome mix from the liver of Sprague−Dawley rats (Invitrogen; 20 mg protein/mL in 0.1 M phosphate buffer) in a shaking water bath at 37 °C. The reaction was stopped by adding 500 μL of ice cold methanol at 0, 15, 30 and 60 min. The samples were centrifuged at 5000 g for 5 min at 4 °C, the supernatants were analyzed, and the test compound was quantified by HPLC. The composition of the mobile phase was adapted to the test compound in a range of MeOH 40-90% and water (0.1% formic acid) 10-60%; flow rate: 1 mL/min; stationary phase: Purospher® STAR, RP18, 5 μm, 125×4, precolumn: Purospher® STAR, RP18, 5 μm, 4×4; detection wavelength: 254 and 280 nm; injection volume: 50 μL. Control samples were performed to check the test compound’s stability in the reaction mixture. The first control was without NADPH, which is needed for the enzymatic activity of the microsomes, the second control was with inactivated microsomes (incubated for 20 min at 90°C), and the third control was without test compound (to determine the baseline). The amounts of the test compound were quantified by an external calibration curve. Data are expressed as the mean ± SEM remaining compound from three independent experiments.

### Chemistry

The synthesis of compounds will be explained in the following and the analytical data for them can be found in the Supporting Information. All commercial chemicals were purchased from common suppliers with a purity ≥ 95% and were used without further purification. The solvents with an analytical grade were obtained from VWR Chemicals and Merck and all dry solvents from Acros Organics. All reactions were carried out under an argon atmosphere. The thin layer chromatography was done with silica gel on aluminum foils (60 Å pore diameter) obtained from Macherey-Nagel and visualized with ultraviolet light (λ = 254 and 365 nm). The purification of the compounds was done by flash chromatography. A puriFlash XS 420 device with a UV-VIS multiwave detector (200−400 nm) from Interchim was used with pre-packed normal-phase PF-SIHP silica columns with particle sizes of 15 and 30 μm (Interchim). Preparative purification by HPLC was carried out on an Agilent 1260 Infinity II device using an Eclipse XDB-C18 (Agilent, 21.2 x 250mm, 7µm) reversed phase column. A suitable gradient (flow rate 21 ml/min.) was used, with 0.1% TFA in water (A) and 0.1% TFA in acetonitrile (B), as a mobile phase. The nuclear magnetic resonance spectroscopy (NMR) was performed with DPX250, AV300, AV400 or AV500 MHz spectrometers from Bruker. Chemical shifts (δ) are reported in parts per million (ppm). DMSO-d6, chloroform-d and methylene chloride-d2 was used as a solvent, and the spectra were calibrated to the solvent signal: 2.50 ppm (1H NMR) or 39.52 ppm (13C NMR) for DMSO-d6, 7.26 ppm (1H NMR) or 77.16 ppm (13C NMR) for chloroform-d and 5.32 ppm (1H NMR) or 54.00 ppm (13C NMR) for methylene chloride-d2. Coupling constants (J) were reported in hertz (Hz) and multiplicities were designated as followed: s (singlet), d (doublet), dd (doublet of doublet), t (triplet), dt (doublet of triplets), td (triplet of doublets), ddd (doublet of doublet of doublet), q (quartet), m (multiplet). Mass spectra were measured on a Surveyor MSQ device from ThermoFisher measuring in the positive-or negative-ion mode. Final compounds were additionally characterized by HRMS using a MALDI LTQ Orbitrap XL from ThermoScientific. The purity of the final compounds was determined by HPLC using an Agilent 1260 Infinity II device with a 1260 DAD HS detector (G7117C; 254 nm, 280 nm, 310 nm) and a LC/MSD device (G6125B, ESI pos. 100-1000). The compounds were analyzed on a Poroshell 120 EC-C18 (Agilent, 3 x 150 mm, 2.7 µm) reversed phase column using 0.1% formic acid in water (A) and 0.1% formic acid in acetonitrile (B) as a mobile phase. The following gradient was used: 0 min 5% B – 2 min 5% B – 8 min 98% B – 10 min 98% B (flow rate of 0.5 mL/min). UV-detection was performed at 254, 280 and 310 nm and all compounds used for further biological characterization showed a purity ≥95%.

### Synthesis of methyl 5-((2,5-dichloropyrimidin-4-yl)amino)-1*H*-pyrazole-3-carboxylate (7)

3-amino-1*H*-pyrazole-5-carboxylate (500 mg, 3.54 mmol, 1.1 eq) and 2,4,5-trichloropyrimidine (591 mg, 3.22 mmol, 1.0 eq) were dissolved in 15 mL anhydrous isopropanol. TEA (978 mg, 9.66 mmol, 3.0 eq) was added and the mixture was stirred at 60 °C for 72 h. The solvent was evaporated under reduced pressure and the crude product was purified by flash chromatography using n-hexane/ethyl acetate as an eluent to obtain the product (781 mg, 84%) as a beige solid. ^1^H NMR (250 MHz, DMSO-d6) δ 13.87 (s, 1H), 10.10 (s, 1H), 8.40 (s, 1H), 7.06 (s, 1H), 3.86 (s, 3H). ^13^C NMR (126 MHz, DMSO) δ 159.22, 156.95, 156.77, 155.60, 146.39, 133.11, 113.43, 102.23, 52.12. MS-ESI m/z [M + H]^+^: calcd 287.1, found 286.0.

### Synthesis of methyl 5-((2-chloro-5-methylpyrimidin-4-yl)amino)-1*H*-pyrazole-3-carboxylate (8)

The title compound was prepared according to the procedure of **7**, using methyl 3-amino-1*H*-pyrazole-5-carboxylate (500 mg, 3.54 mmol) and 2,4-dichloro-5-methylpyrimidine (525 mg, 3.22 mmol). The mixture was stirred at 60 °C for 72 h to obtain the product (60 mg, 7%) as a white solid. ^1^H NMR (250 MHz, DMSO-d6) δ 13.73 (s, 1H, a), 9.68 (s, 1H, b), 8.05 (s, 1H, e), 7.14 (s, 1H, c), 3.86 (s, 3H, f), 2.16 (d, J = 0.9 Hz, 3H, d). ^13^C NMR (126 MHz, DMSO) δ 159.55, 159.31, 156.84, 156.30, 147.62, 132.81, 114.32, 101.26, 52.05, 13.36. MS-ESI m/z [M + H]^+^: calcd 266.7, found 266.0.

### Synthesis of methyl 5-((2-chloroquinazolin-4-yl)amino)-1*H*-pyrazole-3-carboxylate (9)

The title compound was prepared according to the procedure **7**, using methyl 3-amino-1*H*-pyrazole-5-carboxylate (500 mg, 3.54 mmol) and 2,4-dichloroquinazoline (641 mg, 3.22 mmol). The mixture was stirred at 60 °C for 72 h to obtain the product (823 mg, 84%) as a white solid. ^1^H NMR (300 MHz, DMSO-d6) δ 13.88 (s, 1H), 11.12 (s, 1H), 8.67 (d, J = 8.3 Hz, 1H), 7.88 (t, J = 7.6 Hz, 1H), 7.73 (d, J = 8.3 Hz, 1H), 7.61 (t, J = 7.6 Hz, 1H), 7.35 (s, 1H), 3.89 (s, 3H). ^13^C NMR (75 MHz, DMSO) δ 159.32, 158.58, 156.07, 150.82, 147.41, 134.16, 132.93, 126.87, 126.77, 123.63, 113.45, 101.83, 52.11. MS-ESI m/z [M + H]^+^: calcd 304.7, found 304.1.

### Synthesis of methyl 5-((2-((4-(2-((*tert*-butoxycarbonyl)amino)ethyl)phenyl)amino)-5-chloropyrimidin-4-yl)amino)-1*H*-pyrazole-3-carboxylate (10a)

**7** (170 mg, 0.59 mmol, 1.1 eq) and *tert*-butyl 4-aminophenethylcarbamate (127 mg, 0.54 mmol, 1.0 eq) were dissolved in 13 mL anhydrous ethanol. An catalytic amount of 1 M HCl was added and the mixture was stirred under reflux for 18 h. The solvent was evaporated under reduced pressure and the crude product was purified by flash chromatography using DCM/ methanol as an eluent to obtain the product (14 mg, 5%) as a light yellow solid. ^1^H NMR (250 MHz, DMSO-d6) δ 9.92 (s, 1H), 9.88 (s, 1H), 8.23 (s, 1H), 7.50 (d, J = 7.4 Hz, 2H), 7.09 (d, J = 7.5 Hz, 2H), 7.00 – 6.77 (m, 2H), 3.85 (s, 3H), 3.18 – 3.01 (m, 2H), 2.72 – 2.58 (m, 2H), 1.36 (s, 9H). ^13^C NMR (75 MHz, DMSO) δ 160.79, 155.83, 155.40, 155.23, 151.51, 143.10, 137.10, 133.41, 128.67, 119.73, 103.59, 77.37, 51.73, 41.55, 34.90, 28.16. MS-ESI m/z [M + H]^+^: calcd 489.0, found 488.2. HRMS m/z [M + H]^+^: calcd 488.1808, found 488.1799. HPLC: t_R_ = 9.01, purity ≥ 95% (UV: 254/ 280 nm).

### Synthesis of methyl 5-((2-((3-(2-((*tert*-butoxycarbonyl)amino)ethyl)phenyl)amino)-5-chloropyrimidin-4-yl)amino)-1*H*-pyrazole-3-carboxylate (10b)

The title compound was prepared according to the procedure of **10a**, using **7** (170 mg, 0.59 mmol) and *tert*-butyl 3-aminophenethylcarbamate (127 mg, 0.54 mmol). The mixture was stirred for 18 h under reflux to obtain the product (23 mg, 9%) as a light yellow solid. ^1^H NMR (300 MHz, DMSO-d6) δ 13.82 – 13.29 (m, 1H), 9.99 (s, 1H), 9.72 (s, 1H), 8.22 (s, 1H), 7.53 (d, J = 8.2 Hz, 1H), 7.42 (s, 1H), 7.17 (s, 1H), 6.88 – 6.73 (m, 2H), 6.62 (s, 1H), 3.83 (s, 3H), 3.22 – 3.04 (m, 2H), 2.70 – 2.56 (m, 2H), 1.37 (s, 9H). ^13^C NMR (75 MHz, DMSO) δ 157.65, 155.50, 154.62, 140.16, 139.83, 128.46, 126.57, 121.82, 118.74, 116.55, 103.68, 77.51, 51.59, 41.58, 35.77, 28.26. MS-ESI m/z [M + H]^+^: calcd 489.0, found 488.2. HRMS m/z [M + H]^+^: calcd 488.1808, found 488.1802. HPLC: t_R_ = 9.03, purity ≥ 95% (UV: 254/ 280 nm).

### Synthesis of methyl 5-((2-((5-((*tert*-butoxycarbonyl)amino)pentyl)amino)-5-chloropyrimidin-4-yl)amino)-1*H*-pyrazole-3-carboxylate (10c)

**7** (50 mg, 0.17 mmol, 1.0 eq) and *tert*-butyl (5-aminopentyl)carbamate (35 mg, 0.17 mmol, 1.0 eq) were dissolved in 12 mL anhydrous ethanol. TEA (215 mg, 2.13 mmol, 3.0 eq) was added and the mixture was stirred at 80 °C for 3 h under microwave irradiation. The solvent was evaporated under reduced pressure and the crude product was purified by flash chromatography using DCM/methanol as an eluent to obtain the product (28 mg, 35%) as a white solid. ^1^H NMR (300 MHz, DMSO-d6) δ 13.50 (d, J = 68.2 Hz, 1H), 9.36 (d, J = 271.2 Hz, 1H), 7.99 (d, J = 26.2 Hz, 1H), 7.63 – 6.31 (m, 3H), 3.80 (s, 3H), 3.29 – 3.12 (m, 2H), 2.99 –2.84 (m, 2H), 1.63 – 1.43 (m, 2H), 1.44 – 1.25 (m, 13H). ^13^C NMR (75 MHz, DMSO) δ 160.48, 160.03, 155.59, 155.20, 141.57, 140.72, 101.18, 95.10, 77.30, 51.49, 40.73, 29.27, 28.71, 28.25, 23.79. MS-ESI m/z [M + H]^+^: calcd 454.9, found 454.2. HRMS m/z [M + H]^+^: calcd 454.1964, found 454.1954. HPLC: t_R_ = 7.59, purity ≥ 95% (UV: 254/ 280 nm).

### Synthesis of methyl 5-((2-((2-(2-((*tert*-butoxycarbonyl)amino)ethoxy)ethyl)amino)-5-chloropyrimidin-4-yl)amino)-1*H*-pyrazole-3-carboxylate (10d)

The title compound was prepared according to the procedure of **10c**, using **7** (200 mg, 0.69 mmol) and *tert*-butyl (2-(2-aminoethoxy)ethyl)carbamate (142 mg, 0.69 mmol). The mixture was stirred for 8 h at 80 °C to obtain the product (147 mg, 46%) as a white solid. ^1^H NMR (300 MHz, DMSO-d6) δ 13.53 (d, J = 61.3 Hz, 1H), 9.38 (d, J = 280.8 Hz, 1H), 8.02 (d, J = 22.6 Hz, 1H), 7.17 (d, J = 40.1 Hz, 1H), 6.77 (t, J = 5.5 Hz, 1H), 6.57 (s, 1H), 3.92 – 3.75 (m, 3H), 3.58 – 3.48 (m, 2H), 3.49 – 3.35 (m, 4H), 3.17 – 3.01 (m, 2H), 1.38 (s, 9H). ^13^C NMR (75 MHz, DMSO) δ 163.17, 160.84, 156.09, 155.59, 154.53, 142.03, 141.04, 101.91, 95.75, 78.10, 69.55, 69.41, 52.46, 51.95, 41.18, 28.69. MS-ESI m/z [M + H]^+^: calcd 456.9, found 456.3. HRMS m/z [M + H]^+^: calcd 457.1790, found 457.1785. HPLC: t_R_ = 7.35, purity ≥ 95% (UV: 254/ 280 nm).

### Synthesis of methyl 5-((2-((6-((*tert*-butoxycarbonyl)amino)hexyl)amino)-5-chloropyrimidin-4-yl)amino)-1*H*-pyrazole-3-carboxylate (10e)

The title compound was prepared according to the procedure of **10c**, using **7** (200 mg, 0.69 mmol) and *tert*-butyl (6-aminohexyl)carbamate (150 mg, 0.69 mmol). The mixture was stirred for 8 h at 80 °C to obtain the product (177 mg, 54%) as a colorless oil. ^1^H NMR (300 MHz, DMSO-d6) δ 13.49 (d, J = 68.2 Hz, 1H), 9.33 (d, J = 284.3 Hz, 1H), 7.98 (d, J = 28.7 Hz, 1H), 7.17 (d, J = 43.4 Hz, 1H), 6.73 (t, J = 5.7 Hz, 1H), 6.65 – 6.41 (m, 1H), 3.90 – 3.76 (m, 3H), 3.29 – 3.14 (m, 2H), 2.95 – 2.83 (m, 2H), 1.60 – 1.44 (m, 2H), 1.39 – 1.20 (m, 15H). ^13^C NMR (75 MHz, DMSO) δ 162.58, 159.99, 155.54, 155.04, 153.78, 147.66, 132.63, 101.06, 95.10, 77.26, 51.91, 51.45, 40.72, 29.49, 29.00, 28.25, 26.23, 26.11. MS-ESI m/z [M + H]^+^: calcd 469.0, found 468.4. HRMS m/z [M + H]^+^: calcd 468.2121, found 468.2116. HPLC: t_R_ = 7.82, purity ≥ 95% (UV: 254/ 280 nm).

### Synthesis of methyl 5-((2-((4-(2-((*tert*-butoxycarbonyl)amino)ethyl)phenyl)amino)-5-methylpyrimidin-4-yl)amino)-1*H*-pyrazole-3-carboxylate (11a)

The title compound was prepared according to the procedure of **10a**, using **8** (91 mg, 0.34 mmol) and *tert*-butyl 4-aminophenethylcarbamate (73 mg, 0.31 mmol). The mixture was stirred for 18 h under reflux to obtain the product (178 mg, 49%) as a white solid with impurities. It was used without further purification. MS-ESI m/z [M + H]^+^: calcd 468.5, found 468.2.

### Synthesis of methyl 5-((2-((4-(2-((*tert*-butoxycarbonyl)amino)ethyl)phenyl)amino)-quinazolin-4-yl)amino)-1*H*-pyrazole-3-carboxylate (12a)

The title compound was prepared according to the procedure of **10a**, using **9** (300 mg, 0.99 mmol) and *tert*-butyl 4-aminophenethylcarbamate (212 mg, 0.90 mmol). The mixture was stirred for 18 h at 70 °C to obtain the product (325 mg, 72%) as a yellow solid. ^1^H NMR (300 MHz, DMSO-d6) δ 13.32 (s, 1H), 11.84 (s, 1H), 10.76 (s, 1H), 8.73 (d, J = 8.2 Hz, 1H), 7.87 (t, J = 7.7 Hz, 1H), 7.61 (d, J = 8.2 Hz, 1H), 7.55 – 7.41 (m, 3H), 7.23 (d, J = 8.0 Hz, 2H), 7.13 (s, 1H), 6.92 (t, J = 5.4 Hz, 1H), 3.88 (s, 3H), 3.24 – 3.09 (m, 2H), 2.81 – 2.67 (m, 2H), 1.37 (s, 9H). ^13^C NMR (75 MHz, DMSO) δ 159.64, 158.44, 155.56, 151.92, 145.61, 139.54, 136.79, 135.78, 134.22, 129.28, 124.88, 124.78, 122.92, 117.59, 110.25, 102.96, 77.55, 52.06, 41.49, 35.13, 28.25. MS-ESI m/z [M + H]^+^: calcd 505.6, found 505.2. HRMS m/z [M + H]^+^: calcd 504.2354, found 504.2348. HPLC: t_R_ = 7.77, purity ≥ 95% (UV: 254/ 280 nm).

### Synthesis of methyl 5-((2-((3-(2-((*tert*-butoxycarbonyl)amino)ethyl)phenyl)amino)-quinazolin-4-yl)amino)-1*H*-pyrazole-3-carboxylate (12b)

The title compound was prepared according to the procedure of **10a**, using **9** (300 mg, 0.99 mmol) and *tert*-butyl 3-aminophenethylcarbamate (233 mg, 0.99 mmol). The mixture was stirred for 18 h at 70 °C to obtain the product (305 mg, 61%) as a yellow solid. ^1^H NMR (300 MHz, DMSO-d6) δ 13.65 (d, J = 246.4 Hz, 1H), 11.82 (s, 1H), 10.70 (s, 1H), 8.72 (d, J = 8.3 Hz, 1H), 7.88 (t, J = 7.8 Hz, 1H), 7.65 (d, J = 8.3 Hz, 1H), 7.50 (t, J = 7.7 Hz, 1H), 7.45 – 7.29 (m, 3H), 7.13 (d, J = 7.3 Hz, 1H), 7.06 (s, 1H), 6.85 (t, J = 5.9 Hz, 1H), 3.86 (s, 3H), 3.20 – 3.06 (m, 2H), 2.74 – 2.64 (m, 2H), 1.34 (s, 9H). ^13^C NMR (75 MHz, DMSO) δ 159.75, 158.31, 155.56, 152.14, 140.75, 139.90, 136.16, 135.74, 129.18, 126.19, 124.95, 124.70, 123.51, 121.24, 117.69, 110.35, 102.58, 77.60, 51.95, 41.28, 35.38, 28.24. MS-ESI m/z [M + H]^+^: calcd 505.6, found 505.4. HRMS m/z [M + H]^+^: calcd 504.2354, found 504.2347. HPLC: t_R_ = 8.11, purity ≥ 95% (UV: 254/ 280 nm).

### Synthesis of methyl 5-((2-((5-((*tert*-butoxycarbonyl)amino)pentyl)amino)quinazolin-4-yl)amino)-1*H*-pyrazole-3-carboxylate (12c)

The title compound was prepared according to the procedure **10c**, using **9** (300 mg, 0.99 mmol) and *tert*-butyl (5-aminopentyl) carbamate (200 mg, 0.99 mmol). The mixture was stirred for 8 h at 90 °C to obtain the product (233 mg, 50%) as a yellow solid. ^1^H NMR (250 MHz, DMSO-d6) δ 14.04 (s, 1H), 11.60 (s, 1H), 8.59 (d, J = 8.1 Hz, 1H), 8.10 – 7.65 (m, 2H), 7.62 – 7.24 (m, 2H), 6.76 (s, 1H), 3.87 (s, 3H), 3.53 – 3.42 (m, 2H), 3.03 – 2.87 (m, 2H), 1.75 – 1.54 (m, 2H), 1.46 – 1.25 (m, 13H). ^13^C NMR (75 MHz, DMSO) δ 156.10, 136.42, 135.13, 125.12, 124.67, 117.61, 110.68, 102.74, 77.82, 52.56, 41.71, 40.17, 29.66, 29.02, 28.73, 24.05. MS-ESI m/z [M + H]^+^: calcd 470.6, found 470.5. HRMS m/z [M + H]^+^: calcd 470.2510, found 470.2505. HPLC: t_R_ = 7.91, purity ≥ 95% (UV: 254/ 280 nm).

### Synthesis of methyl 5-((2-((2-(2-((*tert*-butoxycarbonyl)amino)ethoxy)ethyl)amino)-quinazolin-4-yl)amino)-1*H*-pyrazole-3-carboxylate (12d)

The title compound was prepared according to the procedure of **10c**, using **9** (300 mg, 0.99 mmol) and *tert*-butyl (2-(2-aminoethoxy)ethyl)carbamate (202 mg, 0.99 mmol). The mixture was stirred for 10 h at 90 °C to obtain the product (151 mg, 32%) as a yellow solid with impurities. MS-ESI m/z [M + H]^+^: calcd 472.5, found 472.2.

### Synthesis of methyl 5-((2-((6-((*tert*-butoxycarbonyl)amino)hexyl)amino)quinazolin-4-yl)amino)-1*H*-pyrazole-3-carboxylate (12e)

The title compound was prepared according to the procedure of **10c**, using **9** (300 mg, 0.99 mmol) and *tert*-butyl (6-aminohexyl)carbamate (214 mg, 0.99 mmol). The mixture was stirred for 10 h at 90 °C to obtain the product (181 mg, 38%) as a yellow solid with impurities. MS-ESI m/z [M + H]^+^: calcd 484.6, found 484.3.

### Synthesis of methyl 5-((2-((4-(2-aminoethyl)phenyl)amino)-5-chloropyrimidin-4-yl)amino)-1*H*-pyrazole-3-carboxylate (13a)

**10a** (308 mg, 0.63 mmol, 1.0 eq) was dissolved in 25 mL anhydrous DCM. TFA (2.88 g, 25.3 mmol, 40.0 eq) was added at 0°C and the reaction mixture was allowed to warm up to rt overnight. The solvent was evaporated under reduced pressure. The residue was dissolved in methanol and neutralized with saturated K_2_CO_3_ solution. The solvent was again evaporated under reduced pressure, and the crude product was used without further purification to obtain the desired product as a white solid together with salts.

### Synthesis of methyl 5-((2-((3-(2-aminoethyl)phenyl)amino)-5-chloropyrimidin-4-yl)amino)-1*H*-pyrazole-3-carboxylate (13b)

The title compound was prepared according to the procedure of **13a**, using **10b** (293 mg, 0.60 mmol). The desired product was obtained as a white solid with salts.

### Synthesis of methyl 5-((2-((5-aminopentyl)amino)-5-chloropyrimidin-4-yl)amino)-1*H*-pyrazole-3-carboxylate (13c)

The title compound was prepared according to the procedure of **13a**, using **10c** (162 mg, 0.36 mmol). The desired product was obtained as a white solid with salts.

### Synthesis of methyl 5-((2-((2-(2-aminoethoxy)ethyl)amino)-5-chloropyrimidin-4-yl)amino)-1*H*-pyrazole-3-carboxylate (13d)

The title compound was prepared according to the procedure of **13a**, using **10d** (110 mg, 0.24 mmol). The desired product was obtained as a white solid with salts.

### Synthesis of methyl 5-((2-((6-aminohexyl)amino)-5-chloropyrimidin-4-yl)amino)-1*H*-pyrazole-3-carboxylate (13e)

The title compound was prepared according to the procedure of **13a**, using **10e** (141 mg, 0.30 mmol). The desired product was obtained as a white solid with salts.

### Synthesis of methyl 5-((2-((4-(2-aminoethyl)phenyl)amino)-5-methylpyrimidin-4-yl)amino)-1*H*-pyrazole-3-carboxylate (14a)

The title compound was prepared according to the procedure of **13a**, using **11a** (112 mg, 0.24 mmol). The desired product was obtained as a white solid with salts.

### Synthesis of methyl 5-((2-((4-(2-aminoethyl)phenyl)amino)quinazolin-4-yl)amino)-1*H*-pyrazole-3-carboxylate (15a)

The title compound was prepared according to the procedure of **13a**, using **12a** (282 mg, 0.56 mmol). The desired product was obtained as a white solid with salts.

### Synthesis of methyl 5-((2-((3-(2-aminoethyl)phenyl)amino)quinazolin-4-yl)amino)-1*H*-pyrazole-3-carboxylate (15b)

The title compound was prepared according to the procedure of **13a**, using **12b** (316 mg, 0.63 mmol). The desired product was obtained as a white solid with salts.

### Synthesis of methyl 5-((2-((5-aminopentyl)amino)quinazolin-4-yl)amino)-1*H*-pyrazole-3-carboxylate (15c)

The title compound was prepared according to the procedure of **13a**, using **12c** (266 mg, 0.57 mmol). The desired product was obtained as a white solid with salts.

### Synthesis of methyl 5-((2-((2-(2-aminoethoxy)ethyl)amino)quinazolin-4-yl)amino)-1*H*-pyrazole-3-carboxylate (15d)

The title compound was prepared according to the procedure of **13a**, using **12d** (147 mg, 0.31 mmol). The desired product was obtained as a white solid with salts.

### Synthesis of methyl 5-((2-((6-aminohexyl)amino)quinazolin-4-yl)amino)-1*H*-pyrazole-3-carboxylate (15e)

The title compound was prepared according to the procedure of **13a**, using **12e** (181 mg, 0.37 mmol). The desired product was obtained as a white solid with salts.

### Synthesis of 5-((2-((4-(2-aminoethyl)phenyl)amino)-5-chloropyrimidin-4-yl)amino)-1*H*-pyrazole-3-carboxylic acid (16a)

**13a** (244 mg, 0.63 mmol, 1.0 eq) and lithium hydroxide monohydrate (132 mg, 3.15 mmol, 5.0 eq) were dissolved in 12.9 mL THF and 3.3 mL H_2_O. The resulting mixture was strirred at 50 °C for 16 h. The solvent was removed under reduced pressure, the residue was dissolved in H_2_O and it was neutralized with a 10% HCl. The solvent was removed under reduced pressure and the crude product was purified by flash chromatography using acetonitile/ water as an eluent to obtain the desired product as a white solid with salts. MS-ESI m/z [M + H]^+^: calcd 374.8, found 374.1.

### Synthesis of 5-((2-((3-(2-aminoethyl)phenyl)amino)-5-chloropyrimidin-4-yl)amino)-1*H*-pyrazole-3-carboxylic acid (16b)

The title compound was prepared according to the procedure of **16a**, using **13b** (233 mg, 0.60 mmol). The desired product was obtained as a white solid with salts. MS-ESI m/z [M + H]^+^: calcd 374.8, found 374.1.

### Synthesis of 5-((2-((5-aminopentyl)amino)-5-chloropyrimidin-4-yl)amino)-1*H*-pyrazole-3-carboxylic acid (16c)

The title compound was prepared according to the procedure of **16a**, using **13c** (162 mg, 0.46 mmol). The desired product was obtained as a white solid with salts. MS-ESI m/z [M + H]^+^: calcd 340.8, found 340.1.

### Synthesis of 5-((2-((2-(2-aminoethoxy)ethyl)amino)-5-chloropyrimidin-4-yl)amino)-1*H*-pyrazole-3-carboxylic acid (16d)

The title compound was prepared according to the procedure of **16a**, using **13d** (86 mg, 0.24 mmol). The desired product was obtained as a white solid with salts. MS-ESI m/z [M + H]^+^: calcd 342.8, found 342.3.

### Synthesis of 5-((2-((6-aminohexyl)amino)-5-chloropyrimidin-4-yl)amino)-1*H*-pyrazole-3-carboxylic acid (16e)

The title compound was prepared according to the procedure of **16a**, using **13e** (110 mg, 0.30 mmol). The desired product was obtained as a white solid with salts. MS-ESI m/z [M + H]^+^: calcd 354.8, found 354.3.

### Synthesis of 5-((2-((4-(2-aminoethyl)phenyl)amino)-5-methylpyrimidin-4-yl)amino)-1*H*-pyrazole-3-carboxylic acid (17a)

The title compound was prepared according to the procedure of **16a**, using **14a** (88 mg, 0.24 mmol). The desired product was obtained as a white solid with salts. MS-ESI m/z [M + H]^+^: calcd 354.4, found 354.2.

### Synthesis of 5-((2-((4-(2-aminoethyl)phenyl)amino)quinazolin-4-yl)amino)-1*H*-pyrazole-3-carboxylic acid (18a)

The title compound was prepared according to the procedure of **16a**, using **15a** (225 mg, 0.56 mmol). The desired product was obtained as a white solid with salts. MS-ESI m/z [M + H]^+^: calcd 390.4, found 390.3.

### Synthesis of 5-((2-((3-(2-aminoethyl)phenyl)amino)quinazolin-4-yl)amino)-1*H*-pyrazole-3-carboxylic acid (18b)

The title compound was prepared according to the procedure of **16a**, using **15b** (250 mg, 0.62 mmol). The desired product was obtained as a white solid with salts. MS-ESI m/z [M + H]^+^: calcd 390.4, found 390.2.

### Synthesis of 5-((2-((5-aminopentyl)amino)quinazolin-4-yl)amino)-1*H*-pyrazole-3-carboxylic acid (18c)

The title compound was prepared according to the procedure of **16a**, using **15c** (209 mg, 0.57 mmol). The desired product was obtained as a white solid with salts. MS-ESI m/z [M + H]^+^: calcd 356.4, found 356.3.

### Synthesis of 5-((2-((2-(2-aminoethoxy)ethyl)amino)quinazolin-4-yl)amino)-1*H*-pyrazole-3-carboxylic acid (18d)

The title compound was prepared according to the procedure of **16a**, using **15d** (115 mg, 0.31 mmol). The desired product was obtained as a white solid with salts. MS-ESI m/z [M + H]^+^: calcd 358.4, found 358.2.

### Synthesis of 5-((2-((6-aminohexyl)amino)quinazolin-4-yl)amino)-1*H*-pyrazole-3-carboxylic acid (18e)

The title compound was prepared according to the procedure of **16a**, using **15e** (143 mg, 0.37 mmol). The desired product was obtained as a white solid with salts. MS-ESI m/z [M + H]^+^: calcd 370.4, found 370.2.

### Synthesis of (Z)-3^5^-chloro-1^1^H-2,4,8-triaza-3(4,2)-pyrimidina-1(3,5)-pyrazola-5(1,4)-benzena-cyclononaphan-9-one (19a)

**16a** (58 mg, 0.16 mmol, 1.0 eq) and HATU (71 mg, 0.18 mmol, 1.2 eq) were dissolved in 60 mL anhydrous DMF. DIPEA (52 mg, 0.40 mmol, 2.6 eq) was added to the resulting mixture and it was stirred at 70 °C for 16 h. The solvent was removed under reduced pressure and the crude product was purified by flash chromatography using acetonitrile/ water as an eluent and by preparative HPLC to obtain the product (5 mg, 9%) as a white solid. ^1^H NMR (300 MHz, DMSO-d6) δ 12.87 (s, 1H), 9.17 (s, 1H), 8.91 (s, 1H), 8.01 (s, 1H), 7.10 – 6.99 (m, 4H), 6.93 (s, 1H), 5.84 (s, 1H), 3.47 (q, J = 6.5 Hz, 2H), 2.91 (t, J = 6.6 Hz, 2H). ^13^C NMR (75 MHz, DMSO) δ 160.63, 159.63, 155.14, 154.62, 146.26, 138.10, 137.04, 135.46, 129.92, 128.22, 103.00, 102.67, 38.09, 32.41. MS-ESI m/z [M + H]^+^: calcd 356.8, found 356.1. HRMS m/z [M + H]^+^: calcd 356.1021, found 356.1023. HPLC: t_R_ = 6.23, purity ≥ 95% (UV: 254/ 280 nm).

### Synthesis of (Z)-3^5^-chloro-1^1^H-2,4,8-triaza-3(4,2)-pyrimidina-1(3,5)-pyrazola-5(1,3)-benzena-cyclononaphan-9-one (19b)

The title compound was prepared according to the procedure of **19a**, using **16b** (100 mg, 0.27 mmol). The mixture was stirred for 16 h at 70 °C to obtain the product (5 mg, 5%) as a white solid. ^1^H NMR (300 MHz, DMSO-d6) δ 9.51 (s, 1H), 9.23 (s, 1H), 8.37 (s, 1H), 8.14 (s, 2H), 7.19 – 7.05 (m, 2H), 7.01 – 6.93 (m, 1H), 6.77 (s, 1H), 3.15 – 3.05 (m, 2H), 2.75 – 2.68 (m, 2H). MS-ESI m/z [M + H]^+^: calcd 356.8, found 356.3. HRMS m/z [M + H]^+^: calcd 356.1021, found 356.1021. HPLC: t_R_ = 6.76, purity ≥ 95% (UV: 254/ 280 nm).

### Synthesis of (Z)-1^5^-chloro-3^1^H-2,5,11-triaza-1(4,2)-pyrimidina-3(3,5)-pyrazolacyclo-undecaphan-4-one (19c)

The title compound was prepared according to the procedure of **19a**, using **16c** (46 mg, 0.14 mmol). The mixture was stirred for 16 h at 50 °C to obtain the product (3 mg, 7%) as a white solid. ^1^H NMR (300 MHz, DMSO-d6) δ 13.10 (s, 1H), 9.00 (s, 1H), 7.99 (t, J = 7.2 Hz, 1H), 7.91 (s, 1H), 7.34 (s, 1H), 7.24 (t, J = 5.6 Hz, 1H), 3.28 – 3.26 (m, 2H), 3.25 – 3.16 (m, 2H), 1.71 – 1.54 (m, 4H), 1.48 – 1.36 (m, 2H). MS-ESI m/z [M + H]^+^: calcd 322.8, found 322.2. HRMS m/z [M + H]^+^: calcd 322.1178, found 322.1179. HPLC: t_R_ = 6.01, purity ≥ 95% (UV: 254/ 280 nm).

### Synthesis of (Z)-1^5^-chloro-3^1^H-8-oxa-2,5,11-triaza-1(4,2)-pyrimidina-3(3,5)-pyrazolacyclo-undecaphan-4-one (19d)

The title compound was prepared according to the procedure of **19a**, using **16d** (70 mg, 0.20 mmol). The mixture was stirred for 16 h at 50 °C to obtain the product (3 mg, 5%) as a beige solid. ^1^H NMR (300 MHz, DMSO-d6) δ 10.14 (s, 1H), 8.13 (s, 1H), 7.90 (t, J = 7.4 Hz, 1H), 7.76 (s, 1H), 7.43 (s, 1H), 3.48 – 3.36 (m, 8H). MS-ESI m/z [M + H]^+^: calcd 324.7, found 324.2. HRMS m/z [M + H]^+^: calcd 324.0970, found 324.0971. HPLC: t_R_ = 5.72, purity ≥ 95% (UV: 254/ 280 nm).

### Synthesis of (Z)-1^5^-chloro-3^1^H-2,5,12-triaza-1(4,2)-pyrimidina-3(3,5)-pyrazolacyclo-dodecaphan-4-one (19e)

The title compound was prepared according to the procedure of **19a**, using **16e** (58 mg, 0.16 mmol). The mixture was stirred for 16 h at 50 °C to obtain the product (22 mg, 40%) as a white solid. ^1^H NMR (300 MHz, DMSO-d6) δ 13.09 (s, 1H), 9.23 (s, 1H), 8.18 (t, J = 5.7 Hz, 1H), 7.92 (s, 1H), 7.23 – 7.03 (m, 2H), 3.11 – 3.01 (m, 2H), 2.91 – 2.81 (m, 2H), 1.62 – 1.43 (m, 6H), 1.43 – 1.29 (m, 2H). ^13^C NMR (75 MHz, DMSO) δ 161.41, 160.86, 159.62, 155.97, 154.11, 102.66, 100.92, 44.28, 41.58, 29.61, 27.56, 27.09, 24.45. MS-ESI m/z [M + H]^+^: calcd 336.8, found 336.2. HRMS m/z [M + H]^+^: calcd 336.1334, found 336.1336. HPLC: t_R_ = 6.09, purity ≥ 95% (UV: 254/ 280 nm).

### Synthesis of (Z)-3^5^-methyl-1^1^H-2,4,8-triaza-3(4,2)-pyrimidina-1(3,5)-pyrazola-5(1,4)-benzena-cyclononaphan-9-one (20a)

The title compound was prepared according to the procedure of **19a**, using **17a** (80 mg, 0.23 mmol). The mixture was stirred for 16 h at 50 °C to obtain the product (9 mg, 12%) as a white solid. ^1^H NMR (500 MHz, DMSO-d6) δ 12.73 (s, 1H), 8.88 (s, 1H), 8.49 (s, 1H), 7.76 (s, 1H), 7.07 (d, J = 8.4 Hz, 2H), 7.01 (d, J = 8.3 Hz, 2H), 6.76 (s, 1H), 5.92 (s, 1H), 3.46 (q, J = 6.6 Hz, 2H), 2.91 (t, J = 6.7 Hz, 2H), 1.99 (d, J = 0.8 Hz, 3H). MS-ESI m/z [M + H]^+^: calcd 336.4, found 336.2. HRMS m/z [M + H]^+^: calcd 336.1567, found 336.1572. HPLC: t_R_ = 6.04, purity ≥ 95% (UV: 254/ 280 nm).

### Synthesis of (Z)-1^1^H-2,4,8-triaza-3(4,2)-quinazolina-1(3,5)-pyrazola-5(1,4)-benzenacyclo-nonaphan-9-one (21a)

The title compound was prepared according to the procedure of **19a**, using **18a** (210 mg, 0.54 mmol). The mixture was stirred for 16 h at 50 °C to obtain the product (7 mg, 4%) as a yellow solid. ^1^H NMR (500 MHz, DMSO-d6) δ 13.32 (s, 1H), 11.64 (s, 1H), 10.47 (s, 1H), 8.57 (d, J = 8.2 Hz, 1H), 7.88 (t, J = 7.8 Hz, 1H), 7.61 (d, J = 8.4 Hz, 1H), 7.48 (t, J = 7.7 Hz, 1H), 7.30 – 7.18 (m, 5H), 5.85 (s, 1H), 3.55 (q, J = 6.5 Hz, 2H), 2.99 (t, J = 6.7 Hz, 2H). ^13^C NMR (126 MHz, DMSO) δ 159.49, 157.68, 153.82, 153.77, 145.34, 140.65, 137.88, 135.59, 135.03, 130.59, 128.47, 124.75, 124.47, 117.89, 110.15, 103.84, 37.91, 32.44. MS-ESI m/z [M + H]^+^: calcd 372.4, found 372.6. HRMS m/z [M + H]^+^: calcd 372.1567, found 372.1569. HPLC: t_R_ = 6.36, purity ≥ 95% (UV: 254/ 280 nm).

### Synthesis of (Z)-1^1^H-2,4,8-triaza-3(4,2)-quinazolina-1(3,5)-pyrazola-5(1,3)-benzenacyclo-nonaphan-9-one (21b)

The title compound was prepared according to the procedure of **19a**, using **18b** (240 mg, 0.62 mmol). The mixture was stirred for 16 h at 50 °C to obtain the product (10 mg, 4%) as a yellow solid. ^1^H NMR (400 MHz, DMSO-d6) δ 13.44 (s, 1H), 11.81 (s, 1H), 10.88 (s, 1H), 8.65 (d, J = 8.8 Hz, 1H), 8.10 (s, 1H), 7.97 – 7.86 (m, 2H), 7.61 (d, J = 8.4 Hz, 1H), 7.50 (t, J = 7.7 Hz, 1H), 7.36 (t, J = 7.7 Hz, 1H), 7.09 (d, J = 8.3 Hz, 2H), 6.92 (s, 1H), 3.74 – 3.63 (m, 2H), 2.87 – 2.80 (m, 2H). ^13^C NMR (101 MHz, DMSO) δ 162.36, 159.20, 158.86, 158.16, 152.60, 139.91, 137.20, 135.87, 129.26, 127.50, 126.28, 124.99, 124.64, 117.99, 117.45, 115.06, 110.19, 100.73, 44.37, 34.34. MS-ESI m/z [M + H]^+^: calcd 372.4, found 372.6. HRMS m/z [M + H]^+^: calcd 372.1567, found 372.1572. HPLC: t_R_ = 6.40, purity ≥ 95% (UV: 254/ 280 nm).

### Synthesis of (Z)-3^1^H-2,5,11-triaza-1(4,2)-quinazolina-3(3,5)-pyrazolacycloundecaphan-4-one (21c)

The title compound was prepared according to the procedure of **19a**, using **18c** (180 mg, 0.51 mmol). The mixture was stirred for 16 h at 50 °C to obtain the product (5 mg, 3%) as a light yellow solid. ^1^H NMR (500 MHz, DMSO-d6) δ 13.20 (s, 1H), 10.59 (s, 1H), 8.40 (d, J = 8.2 Hz, 1H), 8.03 (t, J = 7.2 Hz, 1H), 7.59 – 7.48 (m, 2H), 7.26 (dd, J = 8.4, 1.2 Hz, 1H), 7.19 (t, J = 1071.7, 5.4 Hz, 1H), 7.14 – 7.05 (m, 1H), 3.43 – 3.33 (m, 4H), 1.77 – 1.59 (m, 4H), 1.57 – 1.41 (m, 2H). ^13^C NMR (126 MHz, DMSO) δ 160.68, 158.41, 157.50, 151.31, 148.19, 134.21, 133.02, 123.52, 123.26, 120.56, 110.12, 98.36, 41.90, 41.07, 28.27, 27.26, 22.57. MS-ESI m/z [M + H]^+^: calcd 338.4, found 338.3. HRMS m/z [M + H]^+^: calcd 338.1724, found 338.1730. HPLC: t_R_ = 6.34, purity ≥ 95% (UV: 254/ 280 nm).

### Synthesis of (Z)-3^1^H-8-oxa-2,5,11-triaza-1(4,2)-quinazolina-3(3,5)-pyrazolacyclo-undecaphan-4-one (21d)

The title compound was prepared according to the procedure of **19a**, using **18d** (100 mg, 0.28 mmol). The mixture was stirred for 16 h at 50 °C to obtain the product (21 mg, 22%) as a white solid. ^1^H NMR (400 MHz, DMSO-d6) δ 13.55 (s, 1H), 11.87 (s, 1H), 8.83 (t, J = 5.7 Hz, 1H), 8.62 (d, J = 8.3 Hz, 1H), 7.99 (t, J = 7.1 Hz, 1H), 7.86 – 7.80 (m, 1H), 7.58 (s, 1H), 7.55 – 7.35 (m, 2H), 3.76 – 3.58 (m, 8H). ^13^C NMR (101 MHz, DMSO) δ 161.43, 159.33, 158.02, 153.32, 146.39, 139.75, 135.73, 134.84, 124.62, 116.94, 109.76, 100.71, 69.14, 66.96, 41.70, 34.33. MS-ESI m/z [M + H]^+^: calcd 340.4, found 340.2. HRMS m/z [M + H]^+^: calcd 340.1517, found 340.1513. HPLC: t_R_ = 6.05, purity ≥ 95% (UV: 254/ 280 nm).

### Synthesis of (Z)-3^1^H-2,5,12-triaza-1(4,2)-quinazolina-3(3,5)-pyrazolacyclododecaphan-4-one (21e)

The title compound was prepared according to the procedure of **19a**, using **18e** (90 mg, 0.24 mmol). The mixture was stirred for 16 h at 50 °C to obtain the product (17 mg, 20%) as a yellow solid. ^1^H NMR (400 MHz, DMSO-d6) δ 13.57 (s, 1H), 11.64 (s, 1H), 8.67 (t, J = 5.9 Hz, 1H), 8.54 (d, J = 8.2 Hz, 1H), 8.36 (t, J = 6.1 Hz, 1H), 7.85 (t, J = 7.7 Hz, 1H), 7.53 – 7.42 (m, 2H), 7.27 (s, 1H), 3.34 – 3.23 (m, 4H), 1.68 – 1.53 (m, 2H), 1.56 – 1.47 (m, 2H), 1.46 – 1.34 (m, 2H), 1.31 – 1.19 (m, 2H). ^13^C NMR (101 MHz, DMSO) δ 159.26, 158.89, 154.01, 152.79, 140.15, 135.77, 124.50, 124.42, 118.66, 117.06, 109.57, 103.48, 41.55, 38.50, 29.29, 27.44, 26.96, 24.27. MS-ESI m/z [M + H]^+^: calcd 352.4, found 352.7. HRMS m/z [M + H]^+^: calcd 352.1880, found 352.1882. HPLC: t_R_ = 6.41, purity ≥ 95% (UV: 254/ 280 nm).

## Associated content

Supporting Information Available: The supporting Information contains Tables S1-S6 and the analytical data of compounds **7** – **21**.

## Data availability

The coordinates and structure factors of the MST3-**21c** complex have been deposited in the Protein Data Bank (PDB) under accession number 8QLQ.

## Author Information

### Corresponding Authors

**Thomas Hanke** – Institute for Pharmaceutical Chemistry, Johann Wolfgang Goethe-University, Max-von-Laue-Str. 9, D-60438 Frankfurt am Main, Germany; Structure Genomics Consortium Buchmann Institute for Molecular Life Sciences, Johann Wolfgang Goethe-University, Max-von-Laue-Str. 15, D-60438 Frankfurt am Main, Germany; Phone: (+49)69 798 29313

**Stefan Knapp** – Institute for Pharmaceutical Chemistry, Johann Wolfgang Goethe-University, Max-von-Laue-Str. 9, D-60438 Frankfurt am Main, Germany; Structure Genomics Consortium Buchmann Institute for Molecular Life Sciences, Johann Wolfgang Goethe-University, Max-von-Laue-Str. 15, D-60438 Frankfurt am Main, Germany; Phone: (+49)69 798 29871

### Author Contributions

J.A.A., S.K. and T.H. designed the project; J.A.A. synthesized the compounds; L.M.B., J.F. and B.T.B. performed NanoBRET measurements; A.Kr. performed ITC measurement and provided the proteins for the DSF assay; D.I.B. and A.C.J. performed the X-ray crystallography and structural analyses; A.M. performed the cell cycle experiment; L.E. did the DSF measurements; A.Ka. performed the microsomal stability assay; S.M. and S.K. supervised the research; the manuscript was written by J.A.A., A.C.J., S.K. and T.H. with contributions from all coauthors.

### Notes

The authors declare the following competing financial interest(s): L.M.B. is a cofounder and B.-T.B. is a cofounder and the CEO of the Contract Research Organization CELLinib GmbH, Frankfurt, Germany.

## Acknowledgements

The SGC is a registered charity (no: 1097737) that receives funds from; AbbVie, Bayer AG, Boehringer Ingelheim, Canada Foundation for Innovation, Eshelman Institute for Innovation, Genentech, Genome Canada through Ontario Genomics Institute [OGI-196], EU/EFPIA/OICR/McGill/KTH/Diamond, Innovative Medicines Initiative 2 Joint Undertaking [EUbOPEN grant 875510], Janssen, Merck KGaA (aka EMD in Canada and US), Merck & Co (aka MSD outside Canada and US), Pfizer, São Paulo Research Foundation-FAPESP, Takeda and Wellcome [106169/ZZ14/Z]. D.I.B. and A.C.J. were supported by the German Research Foundation (DFG) grant JO 1473/1-3 (A.C.J.). We thank the stuff at the Swiss Light Source for assistance during x-ray data collection. A.M. is supported by the SFB 1177 ‘Molecular and Functional Characterization of Selective Autophagy. The CQ1 microscope was funded by FUGG (INST 161/920-1 FUGG).

## Abbreviations

ATP, adenosine triphosphate; DCM, dichloromethane; DSF, differential scanning fluorimetry; EMT, epithelial–mesenchymal transition; ERM, ezrin-radixin-moesin; FUCCI, florescent ubiquitination-based cell cycle indicator; Gck, geminal center subgroup; HATU, hexafluorophosphate azabenzotriazole tetramethyl uronium; HCC, hepatocellular carcinoma cells; MST, mammalian sterile 20-like serine/threonin; oN, over night; rt, room temperature; STK, serine/threonine protein kinase; TEA, triethylamine; TFA, trifluoroacetic acid; YSK, yeast Sps1-Ste20-related kinase;

