## Supplementary Material for "Synthesis of pyrazole-based macrocycles leads to a highly selective inhibitor for MST3"

#### Table of Content:

Table S1-S6 S2

Analytical data of compounds **7 – 21** S31

**Table S1.** DSF results of the macrocycles **19a** – **20a** and the lead structure **1** against an *in-house* panel of 104 kinases. Measurements were performed in duplicates.

| Kinase | ΔTm [°C] | | | | | | | | | | | | |
|---|---|---|---|---|---|---|---|---|---|---|---|---|---|
| | 1 | | 19a | | 19b | | 19c | | 19d | | 19e | | 20a |
| BMP2K | 18.8 | BMPR2 | 4.8 | BMP2K | 11.1 | RIOK1 | 10.7 | RIOK1 | 5.3 | RIOK1 | 6.0 | RIOK1 | 2.8 |
| MAPK1 | 17.4 | BMP2K | 4.4 | STK6 | 8.1 | BMP2K | 8.8 | BMP2K | 4.6 | STK6 | 5.5 | RPS6KA5 | 2.6 |
| ULK1 | 15.5 | STK6 | 4.2 | CK2A2 | 6.5 | MELK | 7.2 | RPS6KA5 | 4.4 | RPS6KA5 | 5.4 | BMPR2 | 2.4 |
| CDK2 | 15.2 | MERTK | 3.7 | CAMKK2B | 6.0 | STK6 | 7.0 | CLK1 | 4.0 | BMP2K | 5.0 | MST4 | 2.3 |
| STK6 | 14.3 | PHKG2 | 3.7 | AAK1 | 4.8 | CLK1 | 6.6 | STK6 | 3.4 | MAPK8B | 4.1 | NEK7 | 2.2 |
| HIPK2 | 14.0 | PIM1 | 3.7 | CLK1 | 4.8 | CHEK2 | 6.5 | MELK | 3.2 | AAK1 | 3.8 | MAPK8B | 2.2 |
| AAK1 | 13.8 | MAP2K4 | 3.5 | GSK3B | 4.7 | MAP3K5 | 6.5 | FGFR3 | 3.2 | MAPK10 | 3.7 | MERTK | 2.0 |
| CLK3 | 12.2 | MAPK8B | 3.3 | MAPK8B | 4.7 | UKL3 | 6.3 | AAK1 | 3.2 | GSK3B | 3.6 | BRD4 | 1.9 |
| CAMKK2B | 12.0 | TTK | 3.3 | BMPR2 | 4.3 | MAPK15 | 6.2 | DYRK2 | 3.1 | MAPK9 | 3.6 | FLT1 | 1.9 |
| MAP3K5 | 12.0 | CLK1 | 2.8 | PAK4 | 4.0 | MAPK8B | 5.8 | MAP3K5 | 3.1 | MAPK15 | 3.5 | MAPK9 | 1.8 |
| STK3 | 11.6 | CAMKK2B | 2.6 | CSNK2A1 | 3.9 | FGFR3 | 5.7 | CK2A2 | 3.0 | PIM3 | 3.5 | TLK1 | 1.8 |
| CSNK1D | 11.3 | DYRK2 | 2.5 | GAK | 3.9 | RPS6KA6 | 5.6 | MST4 | 2.8 | MAP3K5 | 3.4 | MAPK10 | 1.8 |
| PLK4 | 11.1 | FLT1 | 2.3 | UKL3 | 3.9 | GSK3B | 5.4 | GSG2 | 2.7 | PLK4 | 3.3 | STK6 | 1.7 |
| EPHA2 | 11.0 | GSK3B | 2.3 | CDK2 | 3.7 | PIM1 | 5.1 | UKL3 | 2.7 | CLK1 | 3.3 | STK17A | 1.5 |
| GAK | 10.6 | MAPK15 | 2.3 | CHEK2 | 3.6 | PIM3 | 5.1 | MAPK15 | 2.6 | MELK | 3.2 | PKMYT1 | 1.5 |
| MAP2K7 | 10.4 | CK2A2 | 2.1 | FGFR3 | 3.4 | NEK2 | 4.9 | GSK3B | 2.4 | PHKG2 | 3.0 | NEK2 | 1.5 |
| CDKL1 | 10.3 | MAPK9 | 2.1 | PLK4 | 3.4 | DYRK2 | 4.8 | MAPK9 | 2.4 | CK2A2 | 3.0 | PIM3 | 1.5 |
| STK10 | 10.1 | CDK2 | 2.0 | DYRK2 | 3.2 | PHKG2 | 4.8 | PLK4 | 2.4 | DYRK2 | 3.0 | CAMKK2B | 1.3 |
| MAP2K6 | 10.0 | CHEK2 | 2.0 | MAP2K4 | 3.1 | BRAF | 4.6 | CSNK2A1 | 2.3 | CAMKK2B | 2.7 | MAPK14 | 1.3 |
| MERTK | 9.8 | RIOK1 | 2.0 | MARK4 | 3.0 | CK2A2 | 4.6 | FGFR2 | 2.2 | CAMK1D | 2.7 | PHKG2 | 1.3 |
| FLT1 | 9.8 | FGFR3 | 1.9 | MELK | 2.9 | AAK1 | 4.5 | MAPK8B | 2.2 | PIM1 | 2.7 | CLK1 | 1.3 |
| STK17B | 9.7 | PIM3 | 1.8 | MST3 | 2.9 | RIOK2 | 4.5 | PIM3 | 2.2 | RIOK2 | 2.5 | PIM1 | 1.2 |
| DCAMKL1 | 9.7 | MELK | 1.7 | MAPK15 | 2.8 | ULK1 | 4.5 | FGFR1B | 2.2 | GAK | 2.5 | TTK | 1.2 |
| STK17A | 9.6 | CSNK1D | 1.6 | MERTK | 2.8 | CAMKK2B | 4.4 | TLK1 | 2.2 | STK17A | 2.4 | EPHB1 | 1.2 |
| STK4 | 9.6 | PAK4 | 1.6 | PHKG2 | 2.7 | MARK3 | 4.4 | DAPK3 | 2.0 | UKL3 | 2.3 | FES | 1.1 |
| MST4 | 9.3 | AAK1 | 1.5 | DMPK1 | 2.6 | TTK | 4.4 | PHKG2 | 2.0 | MST4 | 2.1 | CDK2 | 1.1 |
| BMPR2 | 9.2 | MAP3K5 | 1.5 | FES | 2.5 | CDK2 | 4.3 | MERTK | 2.0 | TTK | 2.1 | AAK1 | 1.1 |
| DMPK1 | 8.9 | MARK4 | 1.5 | FLT1 | 2.5 | GSG2 | 4.3 | BRAF | 1.9 | MERTK | 2.0 | BMP2K | 1.0 |
| PHKG2 | 8.8 | MARK3 | 1.4 | ABL1 | 2.2 | STK38L | 4.1 | ULK1 | 1.9 | CDK2 | 1.9 | STK4 | 1.0 |
| TTK | 8.8 | NEK2 | 1.4 | NEK2 | 2.2 | PLK4 | 4.0 | MAPK10 | 1.9 | MAP2K4 | 1.9 | PLK4 | 1.0 |
| SLK | 8.8 | BRAF | 1.3 | CLK3 | 2.1 | FGFR2 | 3.9 | CSNK1D | 1.9 | CSNK1D | 1.8 | DYRK1A | 1.0 |
| STK38L | 8.7 | DYRK1A | 1.3 | MAP3K5 | 2.1 | MAPK9 | 3.9 | FLT1 | 1.8 | FGFR3 | 1.8 | CLK3 | 1.0 |
| ABL1 | 8.7 | PLK4 | 1.3 | BRD4 | 2.0 | CLK3 | 3.8 | OSR1 | 1.8 | TLK1 | 1.8 | MAP2K7 | 0.9 |
| CLK1 | 8.6 | UKL3 | 1.3 | FGFR2 | 2.0 | STK17A | 3.8 | MARK4 | 1.7 | FLT1 | 1.7 | DAPK3 | 0.9 |
| MAPK8B | 8.5 | MAPK10 | 1.2 | GSG2 | 2.0 | CSNK1D | 3.6 | CDK2 | 1.7 | BMPR2 | 1.6 | CAMK2D | 0.9 |
| DAPK1 | 8.5 | MST3 | 1.2 | RIOK1 | 2.0 | FLT1 | 3.6 | PIM1 | 1.7 | OSR1 | 1.6 | GAK | 0.8 |
| EPHA4 | 8.4 | PCTK1 | 1.2 | STK10 | 2.0 | GAK | 3.5 | STK17A | 1.6 | GSG2 | 1.6 | CHEK2 | 0.8 |
| UKL3 | 8.3 | BRD4 | 1.1 | DAPK3 | 1.9 | MERTK | 3.5 | FES | 1.6 | STK38L | 1.6 | VRK1 | 0.8 |
| MELK | 8.2 | MAP2K7 | 1.1 | MAP2K6 | 1.9 | MAP2K4 | 3.4 | BRD4 | 1.6 | MARK4 | 1.5 | PAK4 | 0.8 |
| DYRK2 | 8.2 | STK10 | 1.1 | PIM1 | 1.9 | BRD4 | 3.3 | CHEK2 | 1.5 | STK4 | 1.5 | FGFR1B | 0.8 |
| MARK4 | 8.1 | STK38L | 1.1 | STK38L | 1.9 | DYRK1A | 3.3 | CAMK1D | 1.5 | DYRK1A | 1.4 | FGFR3 | 0.8 |
| MAPK14 | 8.0 | CAMK2B | 1.0 | TTK | 1.8 | CSNK2A1 | 3.2 | STK38L | 1.3 | CHEK2 | 1.4 | GSK3B | 0.7 |
| GPRK5 | 8.0 | CAMK2D | 1.0 | EPHA7 | 1.7 | STK39 | 3.1 | GAK | 1.3 | PCTK1 | 1.4 | MAP2K4 | 0.7 |
| PCTK1 | 7.7 | DMPK1 | 1.0 | HIPK2 | 1.7 | DAPK3 | 2.9 | RPS6KA6 | 1.3 | CAMK2D | 1.4 | CAMK1D | 0.7 |
| STK39 | 7.6 | CSNK2A1 | 0.9 | MAPK9 | 1.7 | DCAMKL1 | 2.9 | STK17B | 1.3 | FGFR1B | 1.4 | EPHA4 | 0.7 |
| MAPK13 | 7.2 | FES | 0.9 | MARK3 | 1.7 | FGFR1B | 2.8 | BMPR2 | 1.2 | FGFR2 | 1.3 | HIPK2 | 0.6 |
| NEK1 | 7.1 | GPRK5 | 0.9 | PIM3 | 1.7 | PCTK1 | 2.8 | CAMKK2B | 1.1 | DAPK3 | 1.3 | ULK1 | 0.6 |
| CHEK2 | 6.8 | MAPK1 | 0.9 | SLK | 1.7 | RPS6KA1 | 2.8 | STK4 | 1.1 | FES | 1.3 | TOPK | 0.6 |
| MAPKAPK2 | 6.5 | ULK1 | 0.9 | STK3 | 1.7 | STK3 | 2.8 | DYRK1A | 1.1 | CSNK2A1 | 1.2 | CK2A2 | 0.6 |
| EPHA7 | 6.3 | FGFR2 | 0.8 | RIOK2 | 1.5 | ABL1 | 2.7 | RIOK2 | 1.1 | BRAF | 1.2 | MST3 | 0.5 |
| VRK1 | 5.9 | GSG2 | 0.8 | ULK1 | 1.5 | MST3 | 2.7 | MAP2K7 | 1.1 | MARK3 | 1.1 | RPS6KA6 | 0.5 |
| FGFR3 | 5.9 | CLK3 | 0.7 | FGFR1B | 1.4 | BMPR2 | 2.5 | CLK3 | 1.0 | HIPK2 | 1.1 | MELK | 0.5 |
| PAK1 | 5.6 | MAP2K6 | 0.7 | RPS6KA6 | 1.4 | FES | 2.5 | NEK2 | 1.0 | MAP2K7 | 1.0 | DCAMKL1 | 0.5 |
| CAMK1G | 5.5 | MAPK13 | 0.7 | STK39 | 1.4 | STK4 | 2.5 | MARK3 | 1.0 | BRD4 | 1.0 | GSG2 | 0.5 |
| CAMK1D | 5.4 | MST4 | 0.7 | BRAF | 1.3 | DMPK1 | 2.3 | CAMK2D | 1.0 | STK3 | 1.0 | STK38L | 0.5 |
| CAMK4 | 5.4 | STK3 | 0.7 | PCTK1 | 1.2 | EPHA7 | 2.3 | DCAMKL1 | 1.0 | DCAMKL1 | 0.9 | WNK1 | 0.5 |
| GSG2 | 5.2 | FGFR1B | 0.6 | EPHA2 | 1.1 | MAP2K6 | 2.1 | PCTK1 | 1.0 | EPHB1 | 0.7 | MAPKAPK2 | 0.5 |
| RPS6KA1 | 5.1 | OSR1 | 0.6 | EPHA5 | 1.1 | PAK4 | 2.1 | MAP2K4 | 1.0 | RPS6KA6 | 0.7 | PAK1 | 0.5 |
| FES | 5.1 | STK39 | 0.6 | GPRK5 | 1.1 | GPRK5 | 2.0 | STK10 | 0.8 | MAP2K6 | 0.7 | FGFR2 | 0.4 |
| DAPK3 | 4.9 | TOPK | 0.6 | BMX | 1.0 | MAPK13 | 2.0 | DMPK1 | 0.8 | ABL1 | 0.7 | CAMK1G | 0.4 |
| EPHB1 | 4.9 | ABL1 | 0.5 | CAMK1D | 1.0 | OSR1 | 2.0 | PKMYT1 | 0.8 | CASK | 0.7 | CAMK4 | 0.4 |
| RIOK1 | 4.9 | DCAMKL1 | 0.5 | CAMK4 | 1.0 | MAPK10 | 1.9 | EPHB1 | 0.8 | MAPK13 | 0.7 | PCTK1 | 0.4 |
| BRAF | 4.6 | EPHA7 | 0.5 | CSNK1D | 1.0 | SLK | 1.8 | TTK | 0.7 | CAMK2B | 0.7 | CASK | 0.4 |
| FGFR2 | 4.5 | TLK1 | 0.5 | MAPK13 | 1.0 | AURKB | 1.7 | HIPK2 | 0.7 | CDC42BPA | 0.7 | MAP3K5 | 0.4 |
| FGFR1B | 4.5 | DAPK3 | 0.4 | MAPK10 | 0.9 | TLK1 | 1.7 | TIF1 | 0.7 | PKMYT1 | 0.6 | MARK4 | 0.4 |
| CAMK2D | 4.5 | NEK7 | 0.4 | NEK7 | 0.9 | HIPK2 | 1.6 | EPHA4 | 0.7 | EPHA4 | 0.6 | AKT3 | 0.4 |
| SRC | 4.4 | RIOK2 | 0.4 | OSR1 | 0.9 | MST4 | 1.6 | MST3 | 0.6 | STK17B | 0.6 | ABL1 | 0.4 |
| BMX | 4.4 | NEK1 | 0.3 | SRC | 0.9 | CAMK2B | 1.5 | NEK1 | 0.6 | DMPK1 | 0.6 | CSNK2A1 | 0.3 |
| CAMK2B | 4.4 | TIF1 | 0.3 | CAMK1G | 0.8 | MAPK1 | 1.5 | CASK | 0.6 | TIF1 | 0.6 | UKL3 | 0.3 |
| MST3 | 4.4 | WNK1 | 0.3 | STK4 | 0.8 | CAMK2D | 1.3 | STK3 | 0.6 | RPS6KA1 | 0.6 | STK17B | 0.3 |
| PIM3 | 3.8 | BMX | 0.2 | AURKB | 0.7 | STK10 | 1.3 | CDC42BPA | 0.6 | CAMK4 | 0.6 | DAPK1 | 0.3 |
| PAK4 | 3.3 | CAMK4 | 0.2 | DCAMKL1 | 0.7 | EPHA2 | 1.2 | CAMK4 | 0.6 | DAPK1 | 0.6 | SPRK1 | 0.3 |
| PIM1 | 3.3 | CASK | 0.2 | EPHA4 | 0.7 | NEK7 | 1.2 | ABL1 | 0.6 | MAPK1 | 0.5 | GPRK5 | 0.3 |
| DYRK1A | 3.2 | CDC42BPA | 0.2 | NEK1 | 0.7 | CAMK4 | 1.1 | CAMK2B | 0.6 | NEK2 | 0.5 | BRAF | 0.2 |
| CSNK2A1 | 3.1 | EPHA2 | 0.2 | WNK1 | 0.7 | NEK1 | 1.1 | DAPK1 | 0.6 | PAK4 | 0.4 | RPS6KA1 | 0.2 |
| GSK3B | 2.9 | EPHA4 | 0.2 | MAPK14 | 0.6 | BMX | 1.0 | AKT3 | 0.5 | MAPK14 | 0.4 | MAPK13 | 0.2 |
| MAPK15 | 2.5 | EPHA5 | 0.2 | MST4 | 0.6 | SPRK1 | 1.0 | MAPK14 | 0.5 | CAMK1G | 0.4 | OSR1 | 0.2 |
| CASK | 2.0 | GAK | 0.2 | TIF1 | 0.6 | WNK1 | 0.9 | MAP2K6 | 0.5 | GPRK5 | 0.4 | CDKL1 | 0.2 |
| MAP2K4 | 1.4 | HIPK2 | 0.2 | RPS6KA1 | 0.5 | CASK | 0.8 | RPS6KA1 | 0.5 | CLK3 | 0.4 | CSNK1D | 0.2 |
| AKT3 | 1.4 | MAP2K1 | 0.2 | MAP2K7 | 0.4 | DAPK1 | 0.6 | CAMK1G | 0.5 | SRC | 0.4 | DYRK2 | 0.2 |
| BRD4 | 1.3 | MAPK14 | 0.2 | DAPK1 | 0.3 | MARK4 | 0.6 | MAPK1 | 0.5 | STK39 | 0.4 | NEK1 | 0.2 |
| PDK4 | 1.2 | PKMYT1 | 0.2 | MAPK1 | 0.3 | CAMK1G | 0.5 | PAK4 | 0.5 | NEK1 | 0.4 | CDC42BPA | 0.2 |
| NEK2 | 1.1 | RPS6KA1 | 0.2 | STK17A | 0.3 | TOPK | 0.5 | GPRK5 | 0.4 | SPRK1 | 0.4 | MAP2K6 | 0.2 |
| WNK1 | 1.1 | SRC | 0.2 | TLK1 | 0.3 | BRPF1B | 0.4 | VRK1 | 0.4 | VRK1 | 0.4 | MAPK1 | 0.2 |
| PKMYT1 | 1.0 | DAPK1 | 0.1 | CDKL1 | 0.2 | EPHA4 | 0.4 | MAPK13 | 0.4 | CDKL1 | 0.3 | MAPK15 | 0.2 |
| RPS6KA6 | 0.4 | VRK1 | 0.1 | DYRK1A | 0.2 | EPHA5 | 0.4 | CDKL1 | 0.3 | WNK1 | 0.3 | STK3 | 0.2 |
| TAF1 | 0.1 | MAPKAPK2 | 0.0 | EPHB3 | 0.2 | EPHB1 | 0.4 | TAF1 | 0.3 | MST3 | 0.3 | CAMK2B | 0.1 |
| CDC42BPA | -0.1 | SPRK1 | 0.0 | MAP2K1 | 0.2 | MAP2K7 | 0.4 | SPRK1 | 0.3 | TOPK | 0.3 | STK39 | 0.1 |
| AURKB | n.d. | AKT3 | -0.1 | CAMK2B | 0.1 | STK17B | 0.4 | WNK1 | 0.3 | STK10 | 0.3 | EPHA7 | 0.1 |
| BRPF1B | n.d. | CDKL1 | -0.1 | CAMK2D | 0.1 | MAPK14 | 0.3 | NEK7 | 0.2 | BMX | 0.3 | EPHA2 | 0.1 |
| CK2A2 | n.d. | PAK1 | -0.1 | SPRK1 | 0.1 | VRK1 | 0.3 | TOPK | 0.2 | EPHA2 | 0.3 | RIOK2 | 0.0 |
| EPHA5 | n.d. | PDK4 | -0.1 | TAF1 | 0.1 | EPHB3 | 0.2 | EPHA2 | 0.2 | ULK1 | 0.3 | EPHB3 | -0.1 |
| EPHB3 | n.d. | SLK | -0.1 | TOPK | 0.1 | PKMYT1 | 0.2 | MAPKAPK2 | 0.2 | AKT3 | 0.2 | PDK4 | -0.1 |
| MAP2K1 | n.d. | BRPF1B | -0.2 | VRK1 | 0.1 | SRC | 0.2 | SRC | 0.2 | EPHA7 | 0.2 | EPHA5 | -0.1 |
| MAPK10 | n.d. | CAMK1G | -0.2 | CASK | 0.0 | TIF1 | 0.2 | BMX | 0.1 | NEK7 | 0.0 | DMPK1 | -0.1 |
| MAPK9 | n.d. | STK4 | -0.3 | PAK1 | 0.0 | MAP2K1 | 0.1 | EPHB3 | 0.1 | TAF1 | -0.1 | TIF1 | -0.2 |
| MARK3 | n.d. | STK17A | -0.4 | AKT3 | -0.1 | CDKL1 | 0.0 | EPHA5 | -0.1 | MAPKAPK2 | -0.1 | STK10 | -0.2 |
| NEK7 | n.d. | EPHB1 | -0.5 | CDC42BPA | -0.2 | PAK1 | 0.0 | PDK4 | -0.2 | PDK4 | -0.2 | BMX | -0.2 |
| OSR1 | n.d. | EPHB3 | -0.5 | EPHB1 | -0.2 | AKT3 | -0.1 | EPHA7 | -0.2 | PAK1 | -0.4 | BRPF1B | -0.3 |
| RIOK2 | n.d. | CAMK1D | -0.8 | PDK4 | -0.2 | MAPKAPK2 | -0.2 | PAK1 | -0.3 | EPHA5 | -0.5 | TAF1 | -0.3 |
| RPS6KA5 | n.d. | RPS6KA6 | -0.8 | PKMYT1 | -0.2 | PDK4 | -0.2 | STK39 | -0.9 | EPHB3 | -0.7 | MARK3 | -0.4 |
| SPRK1 | n.d. | STK17B | -1.0 | MAPKAPK2 | -0.3 | TAF1 | -0.3 | AURKB | -1.1 | BRPF1B | -1.0 | SRC | -0.7 |
| TIF1 | n.d. | AURKB | -1.3 | BRPF1B | -0.6 | CAMK1D | -0.4 | BRPF1B | -1.2 | AURKB | -2.3 | AURKB | -1.6 |
| TLK1 | n.d. | TAF1 | -1.8 | STK17B | -0.9 | CDC42BPA | -0.6 | MAP2K1 | n.d. | MAP2K1 | n.d. | MAP2K1 | n.d. |
| No. kinases >5°C | 59 | No. kinases >5°C | 0 | No. kinases >5°C | 4 | No. kinases >5°C | 15 | No. kinases >5°C | 1 | No. kinases >5°C | 3 | No. kinases >5°C | 0 |

**Table S2.** DSF results of the macrocycles **21a** – **21e** and the reference staurosporine against an *in-house* panel of 104 kinases. Measurements were performed in duplicates.

| Kinase | ΔTm [°C] | | | | | | | | | | |
|---|---|---|---|---|---|---|---|---|---|---|---|
| | 21a | | 21b | | 21c | | 21d | | 21e | | Staurosporine |
| PIM3 | 6.1 | CAMKK2B | 5.9 | MST3 | 7.5 | GSK3B | 4.8 | GSK3B | 5.5 | CAMKK2B | 24.6 |
| GSK3B | 4.9 | FLT1 | 5.7 | MST4 | 7.4 | PIM3 | 4.4 | MST4 | 4.0 | STK10 | 23.3 |
| MST4 | 4.7 | RIOK1 | 5.3 | MELK | 6.5 | FLT1 | 4.2 | MST3 | 3.7 | PHKG2 | 21.2 |
| STK6 | 4.3 | FGFR3 | 5.1 | RIOK1 | 5.9 | MELK | 4.0 | CK2A2 | 3.5 | PIM3 | 19.7 |
| DYRK1A | 4.2 | BMP2K | 4.9 | GSK3B | 5.5 | STK6 | 3.8 | PIM3 | 3.4 | UKL3 | 19.6 |
| AAK1 | 4.1 | GSK3B | 4.8 | CLK1 | 5.1 | RIOK1 | 3.5 | CAMKK2B | 3.2 | BMP2K | 19.1 |
| PIM1 | 3.9 | STK6 | 4.4 | STK6 | 5.1 | FGFR3 | 3.2 | STK6 | 2.7 | MARK3 | 19.0 |
| TTK | 3.9 | MELK | 4.1 | PIM3 | 5.0 | CLK1 | 3.1 | PLK4 | 2.7 | MAP3K5 | 18.5 |
| MAPK8B | 3.7 | MST4 | 4.0 | PLK4 | 4.4 | BRD4 | 3.1 | CSNK1D | 2.6 | SLK | 18.0 |
| BMP2K | 3.6 | ABL1 | 4.0 | BMP2K | 4.4 | CK2A2 | 3.0 | CDK2 | 2.4 | STK6 | 17.1 |
| FGFR3 | 3.6 | PIM3 | 4.0 | CK2A2 | 4.2 | PLK4 | 2.7 | MELK | 2.4 | CHEK2 | 17.1 |
| FLT1 | 3.5 | MST3 | 3.9 | ABL1 | 4.0 | MST4 | 2.5 | RPS6KA5 | 2.4 | STK3 | 16.6 |
| CK2A2 | 3.3 | BRD4 | 3.9 | AAK1 | 4.0 | MST3 | 2.5 | STK38L | 2.2 | MARK4 | 16.4 |
| MST3 | 3.3 | FGFR2 | 3.8 | TLK1 | 3.7 | OSR1 | 2.5 | AAK1 | 2.2 | CAMK2D | 16.2 |
| BRD4 | 3.2 | MARK4 | 3.7 | CAMKK2B | 3.5 | BMP2K | 2.4 | CLK1 | 2.0 | PLK4 | 16.0 |
| FGFR2 | 3.0 | CLK1 | 3.7 | STK39 | 3.5 | DAPK3 | 2.3 | BRD4 | 1.9 | DAPK3 | 15.8 |
| CAMKK2B | 2.9 | CK2A2 | 3.6 | UKL3 | 3.4 | AAK1 | 2.3 | FLT1 | 1.8 | AAK1 | 15.6 |
| CSNK1D | 2.8 | STK17A | 3.6 | CSNK1D | 3.3 | MAPK8B | 2.2 | PIM1 | 1.8 | CDK2 | 15.5 |
| MARK4 | 2.7 | AAK1 | 3.4 | BRD4 | 3.3 | UKL3 | 2.1 | CAMK1D | 1.8 | STK4 | 14.9 |
| PLK4 | 2.7 | UKL3 | 3.2 | CLK3 | 3.1 | NEK7 | 2.1 | TTK | 1.7 | MAPK15 | 14.7 |
| STK17A | 2.6 | STK10 | 3.2 | OSR1 | 3.1 | CAMKK2B | 2.0 | RIOK1 | 1.6 | MELK | 13.8 |
| UKL3 | 2.6 | DMPK1 | 3.2 | MAPK8B | 3.1 | TLK1 | 2.0 | PCTK1 | 1.6 | RPS6KA5 | 13.7 |
| MERTK | 2.4 | PLK4 | 3.1 | RPS6KA5 | 3.0 | PIM1 | 1.8 | OSR1 | 1.6 | FLT1 | 13.6 |
| PAK4 | 2.4 | MAPK8B | 3.0 | STK17A | 2.9 | CSNK1D | 1.8 | UKL3 | 1.6 | CAMK2B | 13.3 |
| STK3 | 2.3 | FGFR1B | 2.9 | DAPK3 | 2.9 | MAPK10 | 1.7 | MAPK9 | 1.6 | FGFR3 | 13.3 |
| MAPK9 | 2.3 | OSR1 | 2.9 | FLT1 | 2.8 | CLK3 | 1.6 | MAPK8B | 1.5 | STK38L | 12.6 |
| DAPK3 | 2.3 | MAPK10 | 2.7 | DMPK1 | 2.7 | FGFR2 | 1.6 | STK39 | 1.5 | PIM1 | 12.5 |
| TLK1 | 2.2 | GAK | 2.7 | FGFR3 | 2.6 | CHEK2 | 1.6 | BMP2K | 1.5 | DCAMKL1 | 12.4 |
| FGFR1B | 2.2 | CDK2 | 2.7 | PIM1 | 2.6 | ABL1 | 1.6 | TLK1 | 1.4 | MAP2K6 | 12.3 |
| MAPK10 | 2.1 | CSNK1D | 2.7 | GAK | 2.4 | FGFR1B | 1.6 | ABL1 | 1.3 | PAK4 | 12.2 |
| CDK2 | 2.0 | TTK | 2.5 | GPRK5 | 2.4 | MERTK | 1.5 | STK17A | 1.3 | MAP2K4 | 12.0 |
| CLK1 | 2.0 | STK38L | 2.4 | MAPK10 | 2.3 | RPS6KA5 | 1.5 | DMPK1 | 1.3 | CLK1 | 11.9 |
| OSR1 | 2.0 | GPRK5 | 2.4 | NEK7 | 2.3 | CAMK1D | 1.3 | MARK4 | 1.2 | STK17A | 11.8 |
| NEK7 | 1.9 | TLK1 | 2.3 | CDK2 | 2.2 | EPHA4 | 1.3 | DAPK3 | 1.2 | GSK3B | 11.8 |
| RIOK1 | 1.8 | CAMK1G | 2.2 | CAMK1G | 2.2 | GPRK5 | 1.3 | EPHA4 | 1.2 | STK17B | 11.5 |
| STK38L | 1.7 | STK39 | 2.2 | EPHB1 | 2.1 | CSNK2A1 | 1.2 | TIF1 | 1.1 | ULK1 | 11.3 |
| CAMK2D | 1.6 | CLK3 | 2.2 | STK38L | 2.1 | GAK | 1.2 | GAK | 1.0 | TTK | 11.2 |
| ABL1 | 1.5 | MARK3 | 2.2 | MERTK | 2.0 | TTK | 1.2 | GPRK5 | 1.0 | CAMK1G | 10.9 |
| GSG2 | 1.5 | PIM1 | 2.1 | MAPK9 | 1.9 | DMPK1 | 1.2 | MAP2K4 | 1.0 | EPHA7 | 10.3 |
| BMPR2 | 1.5 | DYRK2 | 2.0 | BMX | 1.8 | MAPK14 | 1.1 | MAPK15 | 1.0 | ABL1 | 10.3 |
| CAMK2B | 1.4 | MAPK9 | 1.9 | FGFR1B | 1.8 | MAP2K4 | 1.0 | CHEK2 | 1.0 | CAMK1D | 10.0 |
| PHKG2 | 1.4 | SRC | 1.9 | SPRK1 | 1.8 | CAMK2D | 0.9 | CLK3 | 1.0 | DAPK1 | 9.8 |
| PAK1 | 1.4 | NEK7 | 1.8 | EPHA4 | 1.8 | BMPR2 | 0.9 | EPHB1 | 1.0 | STK39 | 9.6 |
| CAMK1D | 1.4 | EPHB1 | 1.8 | CAMK1D | 1.8 | DYRK2 | 0.9 | FGFR2 | 0.9 | TLK1 | 9.4 |
| RPS6KA5 | 1.4 | CAMK1D | 1.8 | MARK4 | 1.7 | NEK2 | 0.9 | FGFR3 | 0.9 | PCTK1 | 9.1 |
| TIF1 | 1.3 | DAPK3 | 1.8 | EPHA2 | 1.7 | STK39 | 0.9 | PAK4 | 0.9 | FGFR2 | 9.0 |
| VRK1 | 1.3 | STK4 | 1.7 | PHKG2 | 1.7 | CDK2 | 0.9 | RPS6KA6 | 0.8 | GAK | 9.0 |
| CLK3 | 1.3 | MERTK | 1.7 | BMPR2 | 1.7 | ULK1 | 0.8 | BMPR2 | 0.8 | DYRK1A | 8.7 |
| STK10 | 1.3 | EPHA2 | 1.6 | SRC | 1.5 | MAPK9 | 0.8 | NEK7 | 0.7 | DMPK1 | 8.7 |
| PKMYT1 | 1.2 | SPRK1 | 1.6 | FGFR2 | 1.5 | EPHA5 | 0.8 | MAPK14 | 0.7 | CAMK4 | 8.6 |
| GAK | 1.2 | GSG2 | 1.6 | DYRK2 | 1.5 | HIPK2 | 0.8 | PKMYT1 | 0.7 | MAP2K7 | 8.4 |
| PCTK1 | 1.2 | HIPK2 | 1.6 | EPHA5 | 1.5 | GSG2 | 0.8 | STK4 | 0.7 | EPHA2 | 8.3 |
| SPRK1 | 1.2 | RPS6KA5 | 1.5 | CSNK2A1 | 1.5 | STK17A | 0.8 | PHKG2 | 0.7 | AURKB | 8.3 |
| EPHB1 | 1.1 | EPHA4 | 1.5 | TTK | 1.4 | PKMYT1 | 0.8 | MERTK | 0.7 | EPHA5 | 7.9 |
| MELK | 1.1 | BMX | 1.3 | STK4 | 1.4 | EPHB1 | 0.8 | HIPK2 | 0.7 | MAPK8B | 7.8 |
| FES | 1.1 | STK3 | 1.3 | PCTK1 | 1.4 | STK17B | 0.7 | NEK2 | 0.6 | GPRK5 | 7.7 |
| NEK2 | 1.1 | CAMK4 | 1.3 | CHEK2 | 1.4 | FES | 0.7 | CAMK2B | 0.6 | OSR1 | 7.7 |
| BRPF1B | 1.0 | PCTK1 | 1.2 | HIPK2 | 1.2 | SRC | 0.7 | MAPK10 | 0.6 | MAPK13 | 7.5 |
| MARK3 | 1.0 | STK17B | 1.2 | ULK1 | 1.2 | EPHA2 | 0.7 | FGFR1B | 0.6 | MAPK10 | 7.4 |
| STK4 | 1.0 | ULK1 | 1.2 | DYRK1A | 1.2 | CDKL1 | 0.7 | BMX | 0.6 | MST3 | 7.3 |
| CSNK2A1 | 1.0 | BMPR2 | 1.1 | DCAMKL1 | 1.2 | STK10 | 0.7 | CDKL1 | 0.6 | GSG2 | 7.2 |
| CAMK1G | 0.9 | EPHA7 | 1.0 | CAMK4 | 1.2 | CAMK4 | 0.7 | SPRK1 | 0.6 | BMX | 7.1 |
| CHEK2 | 0.9 | CSNK2A1 | 1.0 | CAMK2D | 1.2 | MARK4 | 0.7 | CAMK4 | 0.6 | DYRK2 | 7.0 |
| DAPK1 | 0.9 | FES | 1.0 | CAMK2B | 1.2 | CAMK2B | 0.7 | CAMK2D | 0.6 | SPRK1 | 6.9 |
| MAPK14 | 0.9 | CAMK2B | 1.0 | STK3 | 1.1 | BRAF | 0.7 | DAPK1 | 0.5 | PAK1 | 6.6 |
| EPHA7 | 0.8 | PHKG2 | 1.0 | GSG2 | 1.1 | BMX | 0.6 | ULK1 | 0.5 | MERTK | 6.6 |
| DCAMKL1 | 0.8 | CHEK2 | 0.9 | STK17B | 1.1 | STK38L | 0.6 | DYRK2 | 0.5 | AKT3 | 6.6 |
| RPS6KA1 | 0.8 | EPHA5 | 0.8 | RPS6KA1 | 1.1 | DCAMKL1 | 0.6 | FES | 0.5 | EPHB1 | 6.4 |
| EPHA5 | 0.8 | EPHB3 | 0.8 | STK10 | 1.0 | BRPF1B | 0.6 | CAMK1G | 0.5 | EPHA4 | 6.2 |
| CDKL1 | 0.7 | PAK4 | 0.8 | MARK3 | 1.0 | MARK3 | 0.6 | EPHA5 | 0.5 | FES | 6.2 |
| WNK1 | 0.7 | CAMK2D | 0.7 | MAPK15 | 0.9 | SPRK1 | 0.6 | MARK3 | 0.5 | MST4 | 6.1 |
| CAMK4 | 0.7 | DCAMKL1 | 0.7 | FES | 0.9 | STK4 | 0.6 | STK3 | 0.4 | FGFR1B | 5.9 |
| MAP2K4 | 0.6 | BRAF | 0.7 | EPHA7 | 0.9 | PCTK1 | 0.6 | SRC | 0.4 | SRC | 5.9 |
| RPS6KA6 | 0.6 | MAP2K4 | 0.7 | CDKL1 | 0.8 | DYRK1A | 0.6 | GSG2 | 0.4 | CK2A2 | 5.6 |
| ULK1 | 0.6 | WNK1 | 0.6 | BRAF | 0.8 | PHKG2 | 0.5 | CASK | 0.4 | CLK3 | 5.6 |
| MAPK15 | 0.6 | RPS6KA6 | 0.6 | PKMYT1 | 0.8 | PAK4 | 0.5 | VRK1 | 0.4 | EPHB3 | 5.0 |
| CASK | 0.6 | MAPK15 | 0.6 | NEK2 | 0.8 | RPS6KA6 | 0.5 | MAP3K5 | 0.4 | CASK | 4.8 |
| BMX | 0.5 | DAPK1 | 0.6 | MAP2K4 | 0.7 | MAP2K7 | 0.5 | TOPK | 0.3 | HIPK2 | 4.4 |
| EPHA2 | 0.5 | MAPK1 | 0.5 | MAPK14 | 0.7 | STK3 | 0.5 | STK17B | 0.3 | RPS6KA1 | 3.9 |
| AKT3 | 0.5 | MAP3K5 | 0.5 | EPHB3 | 0.7 | MAP3K5 | 0.4 | DCAMKL1 | 0.3 | MAPK9 | 3.7 |
| DMPK1 | 0.4 | TIF1 | 0.5 | VRK1 | 0.7 | TIF1 | 0.4 | PAK1 | 0.3 | NEK2 | 3.3 |
| STK39 | 0.4 | CASK | 0.5 | PAK4 | 0.6 | PAK1 | 0.4 | STK10 | 0.3 | MAPKAPK2 | 3.2 |
| MAPK1 | 0.4 | NEK1 | 0.4 | PAK1 | 0.6 | MAPK15 | 0.4 | TAF1 | 0.2 | CDKL1 | 3.1 |
| HIPK2 | 0.4 | VRK1 | 0.4 | DAPK1 | 0.6 | WNK1 | 0.4 | AKT3 | 0.2 | CDC42BPA | 2.8 |
| DYRK2 | 0.4 | RPS6KA1 | 0.3 | MAP3K5 | 0.5 | VRK1 | 0.3 | BRAF | 0.2 | VRK1 | 2.8 |
| SRC | 0.4 | PAK1 | 0.3 | CASK | 0.5 | TOPK | 0.3 | CSNK2A1 | 0.2 | BMPR2 | 2.6 |
| GPRK5 | 0.4 | MAP2K6 | 0.2 | TIF1 | 0.5 | CASK | 0.3 | MAP2K7 | 0.2 | CSNK2A1 | 2.1 |
| TOPK | 0.4 | TOPK | 0.2 | NEK1 | 0.5 | EPHA7 | 0.2 | WNK1 | 0.2 | CSNK1D | 1.8 |
| STK17B | 0.3 | MAP2K7 | 0.1 | RPS6KA6 | 0.5 | EPHB3 | 0.2 | EPHB3 | 0.2 | RIOK2 | 1.5 |
| MAP2K7 | 0.3 | MAPKAPK2 | 0.1 | WNK1 | 0.4 | MAP2K6 | 0.2 | EPHA2 | 0.2 | MAPK1 | 1.4 |
| MAPKAPK2 | 0.3 | PDK4 | 0.1 | MAPKAPK2 | 0.3 | MAPKAPK2 | 0.2 | MAPKAPK2 | 0.1 | MAP2K1 | 1.3 |
| MAP2K6 | 0.2 | AKT3 | 0.0 | RIOK2 | 0.3 | NEK1 | 0.1 | EPHA7 | 0.1 | BRD4 | 1.1 |
| MAP3K5 | 0.0 | MAPK14 | 0.0 | MAP2K7 | 0.3 | MAPK1 | 0.1 | BRPF1B | 0.1 | NEK7 | 1.0 |
| CDC42BPA | 0.0 | CDKL1 | -0.1 | AKT3 | 0.2 | RPS6KA1 | 0.1 | RPS6KA1 | 0.1 | WNK1 | 0.9 |
| BRAF | 0.0 | PKMYT1 | -0.1 | TOPK | 0.2 | AKT3 | 0.1 | CDC42BPA | 0.0 | BRAF | 0.8 |
| PDK4 | 0.0 | NEK2 | -0.1 | MAP2K6 | 0.1 | RIOK2 | 0.0 | MAP2K6 | 0.0 | TIF1 | 0.8 |
| TAF1 | -0.2 | RIOK2 | -0.1 | MAPK1 | 0.1 | CDC42BPA | 0.0 | PDK4 | 0.0 | MAPK14 | 0.3 |
| NEK1 | -0.2 | MAPK13 | -0.2 | PDK4 | -0.1 | PDK4 | -0.1 | NEK1 | 0.0 | PDK4 | 0.2 |
| EPHB3 | -0.3 | CDC42BPA | -0.2 | MAPK13 | -0.1 | MAPK13 | -0.1 | MAPK13 | 0.0 | PKMYT1 | 0.2 |
| RIOK2 | -0.3 | TAF1 | -0.4 | TAF1 | -0.2 | CAMK1G | -0.2 | RIOK2 | -0.1 | TAF1 | 0.2 |
| MAPK13 | -0.4 | AURKB | -0.5 | CDC42BPA | -0.4 | DAPK1 | -0.2 | MAPK1 | -0.1 | RIOK1 | 0.0 |
| AURKB | -1.7 | BRPF1B | -0.7 | BRPF1B | -0.8 | TAF1 | -0.3 | AURKB | -0.9 | BRPF1B | -0.1 |
| EPHA4 | -1.8 | DYRK1A | -1.8 | AURKB | -1.4 | AURKB | -2.7 | DYRK1A | -1.6 | RPS6KA6 | -0.5 |
| MAP2K1 | n.d. | MAP2K1 | n.d. | MAP2K1 | n.d. | MAP2K1 | n.d. | MAP2K1 | n.d. | NEK1 | -0.6 |
| No. kinases >5°C | 1 | No. kinases >5°C | 4 | No. kinases >5°C | 7 | No. kinases >5°C | 0 | No. kinases >5°C | 1 | No. kinases >5°C | 75 |

**Table S3.** NanoBRET™ assay information. Tracer K9 and K10 have the following ordering numbers at Promega: N2632 and N2642, respectively).

| Protein Kinase | Alias | Catalog #/ CAS # | NanoLuc orientation | Tracer, used | Tracer $K_{D,app}$ [nM] | Tracer, used [nM] |
|---|---|---|---|---|---|---|
| STK4 | MST1 | NV4351 | N | K10 | 165 | 170 |
| STK3 | MST2 | NV4301 | N | K10 | 270 | 300 |
| STK24 | MST3 | NV4281 | C | K9 | 150 | 150 |
| STK26 | MST4 | NV4291 | C | K9 | 480 | 500 |

**Table S4.** X-ray data collection and refinement statistics for the MST3-JA310 complex

| | MST3-JA310 (**21c**) |
|---|---|
| **Data collection** | |
| Space group | $P2_12_12_1$ |
| Molecules asymmetric unit | 1 |
| *a, b, c* (Å) | 38.13, 79.75, 100.54 |
| α, β, γ (°) | 90.0, 90.0, 90.0 |
| Resolution (Å)[a] | 42.5-1.64 (1.67-1.64) |
| Unique reflections[a] | 37,428 (1,815) |
| Completeness (%)[a] | 97.7 (97.2) |
| Multiplicity[a] | 6.4 (6.3) |
| Mean $I/\sigma(I)$[a] | 15.5 (2.0) |
| $R_{meas}$ | 0.073 (0.785) |
| CC1/2[a] | 0.998 (0.853) |
| **Refinement** | |
| $R_{work}$, (%)[b] | 17.9 |
| $R_{free}$, (%)[b] | 22.7 |
| No. of atoms | |
| Protein[c] | 2197 |
| Inhibitor JA310 | 25 |
| Ethylene glycol | 4 |
| Water | 236 |
| Overall B-factor (Å$^2$) | 27.3 |
| RMSD bond lengths (Å) | 0.006 |
| RMSD bond angles (°) | 0.82 |
| Ramachandran favored (%)[d] | 97.1 |
| Ramachandran outliers (%)[d] | 0.0 |
| **Protein Data Bank entry** | 8QLQ |

[a]Values in parentheses are for the highest-resolution shell.
[b]$R_{work}$ and $R_{free}$ = $\sum||F_{obs}| - |F_{calc}||/\sum|F_{obs}|$, where $R_{free}$ was calculated with 5% of the reflections chosen at random and not used in the refinement.
[c]Number includes alternative conformations.
[d]MolProbity statistics

**Table S5.** FUCCI cell-cycle assay data of **1**, **21c**, and **21d**. Milciclib was used as a reference.

| Compound name | Concen-tration [μM] | Timepoint (h) | [Cell] Count | Ratio Hoechst High Cell count | Ratio Normal Cells | Ratio Healthy [Cell] Count | Ratio Fragmented [Cell] Count | Ratio Pyknosed [Cell] Count | Ratio Red [Cell] Count | Ratio Green [Cell] Count | Ratio Yellow [Cell] Count | Normalized normal cell count |
|---|---|---|---|---|---|---|---|---|---|---|---|---|
| DMSO | 10 | 0 | 105 | 0% | 100% | 97% | 0% | 3% | 74% | 13% | 14% | 0.44 |
| DMSO | 10 | 0 | 129 | 1% | 99% | 97% | 3% | 0% | 77% | 12% | 11% | 0.54 |
| DMSO | 10 | 0 | 164 | 0% | 100% | 95% | 2% | 4% | 75% | 16% | 9% | 0.69 |
| DMSO | 10 | 0 | 261 | 0% | 100% | 95% | 2% | 3% | 78% | 12% | 10% | 1.10 |
| DMSO | 10 | 0 | 275 | 0% | 100% | 97% | 2% | 1% | 78% | 11% | 11% | 1.16 |
| DMSO | 10 | 0 | 265 | 0% | 100% | 97% | 2% | 2% | 77% | 12% | 11% | 1.12 |
| DMSO | 10 | 0 | 206 | 0% | 100% | 93% | 3% | 4% | 71% | 18% | 10% | 0.87 |
| DMSO | 10 | 0 | 275 | 0% | 100% | 96% | 1% | 3% | 70% | 13% | 17% | 1.16 |
| DMSO | 10 | 0 | 229 | 0% | 100% | 96% | 1% | 3% | 79% | 12% | 10% | 0.97 |
| DMSO | 10 | 0 | 260 | 0% | 100% | 98% | 1% | 2% | 72% | 11% | 16% | 1.10 |
| DMSO | 10 | 0 | 261 | 0% | 100% | 97% | 2% | 2% | 71% | 14% | 15% | 1.10 |
| DMSO | 10 | 0 | 300 | 0% | 100% | 97% | 2% | 1% | 70% | 16% | 14% | 1.27 |
| DMSO | 10 | 0 | 328 | 0% | 100% | 96% | 2% | 2% | 73% | 13% | 14% | 1.38 |
| DMSO | 10 | 0 | 256 | 0% | 100% | 95% | 4% | 2% | 67% | 18% | 14% | 1.08 |
| DMSO | 10 | 6 | 102 | 5% | 95% | 96% | 0% | 4% | 11% | 66% | 24% | 0.48 |
| DMSO | 10 | 6 | 119 | 2% | 98% | 95% | 0% | 5% | 11% | 63% | 26% | 0.57 |
| DMSO | 10 | 6 | 143 | 3% | 97% | 94% | 0% | 6% | 10% | 67% | 23% | 0.68 |
| DMSO | 10 | 6 | 232 | 2% | 98% | 94% | 0% | 6% | 8% | 66% | 26% | 1.12 |
| DMSO | 10 | 6 | 251 | 3% | 97% | 96% | 0% | 4% | 8% | 71% | 22% | 1.20 |
| DMSO | 10 | 6 | 223 | 4% | 96% | 93% | 0% | 7% | 6% | 74% | 20% | 1.05 |
| DMSO | 10 | 6 | 196 | 5% | 95% | 96% | 0% | 4% | 7% | 73% | 21% | 0.92 |

| | | | | | | | | | | | | |
|---|---|---|---|---|---|---|---|---|---|---|---|---|
| DMSO | 10 | 6 | 236 | 2% | 98% | 96% | 0% | 4% | 6% | 65% | 29% | 1.14 |
| DMSO | 10 | 6 | 218 | 4% | 96% | 98% | 0% | 2% | 7% | 71% | 22% | 1.03 |
| DMSO | 10 | 6 | 235 | 2% | 98% | 96% | 0% | 4% | 5% | 74% | 21% | 1.13 |
| DMSO | 10 | 6 | 231 | 5% | 95% | 98% | 0% | 2% | 5% | 67% | 28% | 1.08 |
| DMSO | 10 | 6 | 260 | 2% | 98% | 95% | 0% | 5% | 11% | 71% | 18% | 1.25 |
| DMSO | 10 | 6 | 278 | 3% | 97% | 94% | 0% | 6% | 6% | 73% | 20% | 1.32 |
| DMSO | 10 | 6 | 219 | 4% | 96% | 94% | 0% | 6% | 9% | 70% | 21% | 1.04 |
| DMSO | 10 | 12 | 110 | 4% | 96% | 95% | 0% | 5% | 6% | 82% | 12% | 0.49 |
| DMSO | 10 | 12 | 129 | 1% | 99% | 96% | 0% | 4% | 12% | 73% | 15% | 0.59 |
| DMSO | 10 | 12 | 155 | 1% | 99% | 94% | 0% | 6% | 10% | 78% | 12% | 0.71 |
| DMSO | 10 | 12 | 233 | 2% | 98% | 94% | 0% | 6% | 7% | 74% | 19% | 1.06 |
| DMSO | 10 | 12 | 259 | 1% | 99% | 95% | 0% | 5% | 7% | 79% | 14% | 1.19 |
| DMSO | 10 | 12 | 243 | 4% | 96% | 94% | 0% | 6% | 10% | 84% | 7% | 1.08 |
| DMSO | 10 | 12 | 205 | 3% | 97% | 94% | 0% | 6% | 7% | 74% | 18% | 0.92 |
| DMSO | 10 | 12 | 250 | 2% | 98% | 95% | 0% | 5% | 6% | 78% | 16% | 1.14 |
| DMSO | 10 | 12 | 224 | 3% | 97% | 96% | 0% | 4% | 9% | 80% | 11% | 1.01 |
| DMSO | 10 | 12 | 242 | 1% | 99% | 96% | 0% | 4% | 7% | 81% | 12% | 1.11 |
| DMSO | 10 | 12 | 243 | 2% | 98% | 94% | 0% | 6% | 9% | 73% | 18% | 1.11 |
| DMSO | 10 | 12 | 268 | 1% | 99% | 97% | 0% | 3% | 10% | 78% | 12% | 1.23 |
| DMSO | 10 | 12 | 292 | 3% | 97% | 94% | 0% | 6% | 8% | 80% | 12% | 1.32 |
| DMSO | 10 | 12 | 228 | 3% | 97% | 94% | 0% | 6% | 10% | 78% | 12% | 1.03 |
| DMSO | 10 | 24 | 121 | 1% | 99% | 93% | 0% | 7% | 38% | 56% | 6% | 0.50 |
| DMSO | 10 | 24 | 133 | 1% | 99% | 95% | 0% | 5% | 45% | 41% | 13% | 0.56 |

| | | | | | | | | | | | | |
|---|---|---|---|---|---|---|---|---|---|---|---|---|
| DMSO | 10 | 24 | 167 | 3% | 97% | 92% | 0% | 8% | 44% | 48% | 8% | 0.68 |
| DMSO | 10 | 24 | 253 | 3% | 97% | 93% | 0% | 7% | 38% | 46% | 16% | 1.03 |
| DMSO | 10 | 24 | 283 | 2% | 98% | 94% | 0% | 6% | 35% | 52% | 13% | 1.16 |
| DMSO | 10 | 24 | 262 | 3% | 97% | 94% | 0% | 6% | 45% | 42% | 13% | 1.06 |
| DMSO | 10 | 24 | 229 | 4% | 96% | 95% | 0% | 5% | 40% | 48% | 12% | 0.93 |
| DMSO | 10 | 24 | 288 | 2% | 98% | 95% | 0% | 5% | 45% | 39% | 16% | 1.19 |
| DMSO | 10 | 24 | 248 | 1% | 99% | 93% | 0% | 7% | 42% | 48% | 10% | 1.03 |
| DMSO | 10 | 24 | 273 | 2% | 98% | 97% | 0% | 3% | 41% | 48% | 11% | 1.13 |
| DMSO | 10 | 24 | 260 | 3% | 97% | 91% | 0% | 9% | 38% | 46% | 16% | 1.06 |
| DMSO | 10 | 24 | 302 | 2% | 98% | 96% | 0% | 4% | 40% | 46% | 15% | 1.24 |
| DMSO | 10 | 24 | 335 | 3% | 97% | 94% | 0% | 6% | 45% | 47% | 8% | 1.37 |
| DMSO | 10 | 24 | 255 | 2% | 98% | 93% | 0% | 7% | 46% | 43% | 10% | 1.06 |
| DMSO | 10 | 48 | 151 | 1% | 99% | 98% | 0% | 2% | 54% | 38% | 8% | 0.49 |
| DMSO | 10 | 48 | 179 | 1% | 99% | 98% | 0% | 2% | 52% | 39% | 9% | 0.58 |
| DMSO | 10 | 48 | 199 | 2% | 98% | 97% | 0% | 3% | 50% | 38% | 12% | 0.64 |
| DMSO | 10 | 48 | 329 | 1% | 99% | 96% | 0% | 4% | 49% | 39% | 12% | 1.07 |
| DMSO | 10 | 48 | 353 | 2% | 98% | 97% | 0% | 3% | 46% | 41% | 13% | 1.13 |
| DMSO | 10 | 48 | 326 | 1% | 99% | 97% | 0% | 3% | 53% | 36% | 11% | 1.06 |
| DMSO | 10 | 48 | 268 | 1% | 99% | 97% | 0% | 3% | 50% | 41% | 9% | 0.87 |
| DMSO | 10 | 48 | 359 | 0% | 100% | 96% | 0% | 4% | 56% | 36% | 8% | 1.17 |
| DMSO | 10 | 48 | 300 | 0% | 100% | 96% | 0% | 4% | 49% | 38% | 13% | 0.98 |
| DMSO | 10 | 48 | 333 | 2% | 98% | 98% | 0% | 2% | 51% | 37% | 12% | 1.07 |
| DMSO | 10 | 48 | 320 | 2% | 98% | 96% | 0% | 4% | 48% | 39% | 12% | 1.02 |

| | | | | | | | | | | | | |
|---|---|---|---|---|---|---|---|---|---|---|---|---|
| DMSO | 10 | 48 | 364 | 1% | 99% | 96% | 0% | 4% | 47% | 40% | 13% | 1.17 |
| DMSO | 10 | 48 | 467 | 1% | 99% | 97% | 0% | 3% | 46% | 41% | 14% | 1.51 |
| DMSO | 10 | 48 | 376 | 0% | 100% | 97% | 0% | 3% | 45% | 44% | 10% | 1.23 |
| DMSO | 10 | 72 | 268 | 0% | 100% | 98% | 0% | 2% | 44% | 43% | 12% | 0.52 |
| DMSO | 10 | 72 | 240 | 0% | 100% | 99% | 0% | 1% | 53% | 29% | 18% | 0.46 |
| DMSO | 10 | 72 | 349 | 0% | 100% | 98% | 0% | 2% | 52% | 30% | 18% | 0.68 |
| DMSO | 10 | 72 | 568 | 1% | 99% | 98% | 0% | 2% | 55% | 35% | 11% | 1.10 |
| DMSO | 10 | 72 | 550 | 0% | 100% | 99% | 0% | 1% | 48% | 36% | 15% | 1.07 |
| DMSO | 10 | 72 | 553 | 0% | 100% | 100% | 0% | 0% | 49% | 40% | 12% | 1.07 |
| DMSO | 10 | 72 | 421 | 0% | 100% | 99% | 0% | 1% | 56% | 31% | 13% | 0.82 |
| DMSO | 10 | 72 | 585 | 1% | 99% | 99% | 0% | 1% | 56% | 33% | 11% | 1.13 |
| DMSO | 10 | 72 | 531 | 1% | 99% | 98% | 0% | 2% | 58% | 29% | 14% | 1.02 |
| DMSO | 10 | 72 | 534 | 1% | 99% | 99% | 0% | 1% | 45% | 41% | 13% | 1.03 |
| DMSO | 10 | 72 | 578 | 1% | 99% | 99% | 0% | 1% | 50% | 36% | 15% | 1.11 |
| DMSO | 10 | 72 | 631 | 0% | 100% | 99% | 0% | 1% | 54% | 33% | 13% | 1.22 |
| DMSO | 10 | 72 | 759 | 1% | 99% | 100% | 0% | 0% | 43% | 43% | 15% | 1.46 |
| DMSO | 10 | 72 | 682 | 1% | 99% | 99% | 0% | 1% | 48% | 39% | 13% | 1.32 |
| 21c | 1 | 0 | 223 | 0% | 100% | 96% | 1% | 2% | 77% | 10% | 13% | 0.94 |
| 21c | 1 | 0 | 264 | 0% | 100% | 96% | 1% | 3% | 76% | 14% | 10% | 1.12 |
| 21c | 1 | 0 | 347 | 0% | 100% | 92% | 4% | 4% | 67% | 18% | 15% | 1.47 |
| 21c | 1 | 0 | 303 | 0% | 100% | 94% | 2% | 3% | 71% | 17% | 12% | 1.28 |
| 21c | 10 | 0 | 193 | 0% | 100% | 95% | 2% | 3% | 83% | 7% | 10% | 0.82 |
| 21c | 10 | 0 | 166 | 0% | 100% | 92% | 4% | 4% | 73% | 13% | 14% | 0.70 |

| | | | | | | | | | | | | |
|---|---|---|---|---|---|---|---|---|---|---|---|---|
| 21c | 10 | 0 | 295 | 0% | 100% | 97% | 2% | 2% | 73% | 16% | 11% | 1.25 |
| 21c | 10 | 0 | 231 | 0% | 100% | 97% | 2% | 2% | 74% | 14% | 12% | 0.97 |
| 21c | 5 | 0 | 141 | 0% | 100% | 96% | 2% | 1% | 71% | 18% | 12% | 0.60 |
| 21c | 5 | 0 | 202 | 0% | 100% | 99% | 0% | 1% | 73% | 12% | 16% | 0.85 |
| 21c | 5 | 0 | 262 | 1% | 99% | 98% | 1% | 2% | 77% | 11% | 12% | 1.10 |
| 21c | 5 | 0 | 242 | 1% | 99% | 95% | 1% | 4% | 73% | 14% | 12% | 1.01 |
| 21c | 1 | 6 | 187 | 2% | 98% | 91% | 1% | 8% | 8% | 73% | 19% | 0.90 |
| 21c | 1 | 6 | 231 | 3% | 97% | 94% | 0% | 6% | 9% | 62% | 29% | 1.10 |
| 21c | 1 | 6 | 294 | 5% | 95% | 91% | 0% | 8% | 8% | 67% | 25% | 1.37 |
| 21c | 1 | 6 | 248 | 4% | 96% | 91% | 0% | 8% | 10% | 71% | 20% | 1.17 |
| 21c | 10 | 6 | 172 | 2% | 98% | 95% | 0% | 5% | 7% | 70% | 23% | 0.82 |
| 21c | 10 | 6 | 142 | 4% | 96% | 96% | 0% | 4% | 6% | 62% | 32% | 0.67 |
| 21c | 10 | 6 | 255 | 2% | 98% | 96% | 0% | 4% | 6% | 69% | 25% | 1.22 |
| 21c | 10 | 6 | 191 | 6% | 94% | 92% | 0% | 8% | 4% | 70% | 25% | 0.88 |
| 21c | 5 | 6 | 117 | 3% | 97% | 90% | 0% | 10% | 9% | 68% | 23% | 0.56 |
| 21c | 5 | 6 | 157 | 4% | 96% | 92% | 0% | 8% | 9% | 70% | 21% | 0.74 |
| 21c | 5 | 6 | 223 | 2% | 98% | 93% | 0% | 7% | 10% | 62% | 29% | 1.07 |
| 21c | 5 | 6 | 207 | 5% | 95% | 91% | 0% | 9% | 12% | 65% | 23% | 0.96 |
| 21c | 1 | 12 | 207 | 2% | 98% | 93% | 0% | 7% | 5% | 85% | 10% | 0.94 |
| 21c | 1 | 12 | 236 | 2% | 98% | 93% | 0% | 7% | 11% | 73% | 17% | 1.07 |
| 21c | 1 | 12 | 313 | 3% | 97% | 93% | 0% | 7% | 8% | 77% | 16% | 1.41 |
| 21c | 1 | 12 | 273 | 3% | 97% | 92% | 0% | 8% | 9% | 80% | 12% | 1.24 |
| 21c | 10 | 12 | 177 | 2% | 98% | 94% | 0% | 6% | 8% | 75% | 17% | 0.81 |

| | | | | | | | | | | | | |
|---|---|---|---|---|---|---|---|---|---|---|---|---|
| 21c | 10 | 12 | 145 | 6% | 94% | 94% | 0% | 6% | 12% | 59% | 29% | 0.63 |
| 21c | 10 | 12 | 263 | 3% | 97% | 93% | 0% | 7% | 9% | 69% | 22% | 1.18 |
| 21c | 10 | 12 | 196 | 4% | 96% | 90% | 1% | 9% | 8% | 77% | 15% | 0.87 |
| 21c | 5 | 12 | 120 | 6% | 94% | 92% | 1% | 7% | 11% | 68% | 21% | 0.52 |
| 21c | 5 | 12 | 171 | 4% | 96% | 94% | 1% | 5% | 10% | 74% | 17% | 0.77 |
| 21c | 5 | 12 | 225 | 6% | 94% | 91% | 0% | 9% | 8% | 75% | 17% | 0.98 |
| 21c | 5 | 12 | 204 | 5% | 95% | 88% | 0% | 12% | 12% | 66% | 22% | 0.90 |
| 21c | 1 | 24 | 233 | 3% | 97% | 91% | 0% | 9% | 40% | 49% | 11% | 0.95 |
| 21c | 1 | 24 | 263 | 2% | 98% | 92% | 0% | 8% | 42% | 49% | 9% | 1.08 |
| 21c | 1 | 24 | 342 | 4% | 96% | 90% | 0% | 10% | 40% | 51% | 8% | 1.39 |
| 21c | 1 | 24 | 307 | 4% | 96% | 93% | 0% | 7% | 44% | 43% | 13% | 1.24 |
| 21c | 10 | 24 | 177 | 3% | 97% | 88% | 0% | 12% | 12% | 71% | 17% | 0.72 |
| 21c | 10 | 24 | 159 | 4% | 96% | 90% | 0% | 10% | 9% | 70% | 22% | 0.64 |
| 21c | 10 | 24 | 270 | 6% | 94% | 90% | 0% | 10% | 16% | 64% | 20% | 1.06 |
| 21c | 10 | 24 | 194 | 8% | 92% | 90% | 1% | 9% | 10% | 63% | 27% | 0.75 |
| 21c | 5 | 24 | 128 | 2% | 98% | 89% | 2% | 10% | 23% | 61% | 16% | 0.53 |
| 21c | 5 | 24 | 178 | 2% | 98% | 90% | 0% | 10% | 15% | 72% | 13% | 0.73 |
| 21c | 5 | 24 | 232 | 3% | 97% | 88% | 0% | 11% | 19% | 67% | 14% | 0.94 |
| 21c | 5 | 24 | 224 | 3% | 97% | 85% | 0% | 15% | 18% | 68% | 14% | 0.91 |
| 21c | 1 | 48 | 248 | 0% | 100% | 94% | 0% | 6% | 55% | 34% | 12% | 0.81 |
| 21c | 1 | 48 | 293 | 1% | 99% | 95% | 0% | 5% | 68% | 24% | 8% | 0.95 |
| 21c | 1 | 48 | 344 | 2% | 98% | 92% | 0% | 8% | 56% | 36% | 8% | 1.10 |
| 21c | 1 | 48 | 340 | 1% | 99% | 93% | 0% | 6% | 52% | 38% | 10% | 1.10 |

| | | | | | | | | | | | | |
|---|---|---|---|---|---|---|---|---|---|---|---|---|
| 21c | 10 | 48 | 183 | 7% | 93% | 80% | 1% | 20% | 36% | 40% | 24% | 0.56 |
| 21c | 10 | 48 | 160 | 9% | 91% | 81% | 0% | 19% | 39% | 35% | 26% | 0.47 |
| 21c | 10 | 48 | 248 | 12% | 88% | 83% | 0% | 17% | 42% | 34% | 24% | 0.71 |
| 21c | 10 | 48 | 199 | 16% | 84% | 85% | 1% | 14% | 43% | 41% | 17% | 0.55 |
| 21c | 5 | 48 | 131 | 8% | 92% | 83% | 2% | 15% | 46% | 41% | 13% | 0.39 |
| 21c | 5 | 48 | 173 | 6% | 94% | 83% | 0% | 17% | 39% | 45% | 16% | 0.53 |
| 21c | 5 | 48 | 242 | 7% | 93% | 80% | 0% | 20% | 54% | 38% | 8% | 0.74 |
| 21c | 5 | 48 | 219 | 8% | 92% | 81% | 0% | 18% | 48% | 33% | 18% | 0.66 |
| 21c | 1 | 72 | 323 | 1% | 99% | 97% | 0% | 3% | 72% | 19% | 9% | 0.62 |
| 21c | 1 | 72 | 305 | 1% | 99% | 98% | 0% | 2% | 73% | 17% | 10% | 0.59 |
| 21c | 1 | 72 | 458 | 1% | 99% | 96% | 0% | 4% | 66% | 22% | 12% | 0.88 |
| 21c | 1 | 72 | 467 | 1% | 99% | 97% | 0% | 3% | 67% | 21% | 12% | 0.90 |
| 21c | 10 | 72 | 158 | 14% | 86% | 66% | 4% | 30% | 37% | 47% | 17% | 0.26 |
| 21c | 10 | 72 | 146 | 18% | 82% | 69% | 0% | 31% | 35% | 37% | 28% | 0.23 |
| 21c | 10 | 72 | 217 | 15% | 85% | 64% | 2% | 34% | 38% | 45% | 18% | 0.36 |
| 21c | 10 | 72 | 169 | 17% | 83% | 66% | 4% | 29% | 39% | 45% | 16% | 0.27 |
| 21c | 5 | 72 | 136 | 15% | 85% | 66% | 2% | 32% | 46% | 46% | 8% | 0.22 |
| 21c | 5 | 72 | 187 | 4% | 96% | 76% | 1% | 23% | 43% | 42% | 15% | 0.35 |
| 21c | 5 | 72 | 264 | 6% | 94% | 88% | 0% | 11% | 52% | 32% | 16% | 0.48 |
| 21c | 5 | 72 | 243 | 5% | 95% | 84% | 0% | 16% | 54% | 32% | 14% | 0.45 |
| 21d | 1 | 0 | 232 | 0% | 100% | 94% | 1% | 5% | 72% | 13% | 15% | 0.98 |
| 21d | 1 | 0 | 214 | 0% | 100% | 96% | 1% | 2% | 76% | 12% | 12% | 0.90 |
| 21d | 1 | 0 | 263 | 0% | 100% | 92% | 5% | 3% | 75% | 12% | 14% | 1.11 |

| | | | | | | | | | | | | |
|---|---|---|---|---|---|---|---|---|---|---|---|---|
| 21d | 1 | 0 | 285 | 0% | 100% | 98% | 1% | 1% | 71% | 13% | 17% | 1.20 |
| 21d | 10 | 0 | 217 | 0% | 100% | 97% | 0% | 2% | 75% | 14% | 11% | 0.92 |
| 21d | 10 | 0 | 218 | 0% | 100% | 94% | 2% | 4% | 81% | 12% | 7% | 0.92 |
| 21d | 10 | 0 | 251 | 0% | 100% | 97% | 2% | 1% | 70% | 20% | 10% | 1.06 |
| 21d | 10 | 0 | 266 | 0% | 100% | 95% | 1% | 4% | 76% | 14% | 10% | 1.12 |
| 21d | 5 | 0 | 135 | 0% | 100% | 95% | 3% | 2% | 75% | 16% | 9% | 0.57 |
| 21d | 5 | 0 | 202 | 0% | 100% | 97% | 2% | 1% | 80% | 11% | 9% | 0.85 |
| 21d | 5 | 0 | 318 | 0% | 100% | 96% | 2% | 3% | 75% | 16% | 10% | 1.34 |
| 21d | 5 | 0 | 228 | 0% | 100% | 96% | 1% | 3% | 70% | 16% | 15% | 0.96 |
| 21d | 1 | 6 | 205 | 1% | 99% | 95% | 0% | 5% | 7% | 70% | 23% | 1.00 |
| 21d | 1 | 6 | 187 | 3% | 97% | 96% | 0% | 4% | 8% | 58% | 34% | 0.89 |
| 21d | 1 | 6 | 249 | 3% | 97% | 94% | 0% | 6% | 10% | 63% | 27% | 1.19 |
| 21d | 1 | 6 | 251 | 2% | 98% | 95% | 0% | 5% | 9% | 68% | 23% | 1.21 |
| 21d | 10 | 6 | 200 | 2% | 98% | 92% | 0% | 8% | 8% | 69% | 22% | 0.96 |
| 21d | 10 | 6 | 187 | 5% | 95% | 94% | 0% | 6% | 14% | 66% | 20% | 0.87 |
| 21d | 10 | 6 | 232 | 2% | 98% | 94% | 0% | 6% | 5% | 79% | 16% | 1.12 |
| 21d | 10 | 6 | 234 | 3% | 97% | 94% | 0% | 6% | 7% | 66% | 27% | 1.12 |
| 21d | 5 | 6 | 116 | 5% | 95% | 95% | 0% | 5% | 5% | 77% | 18% | 0.54 |
| 21d | 5 | 6 | 187 | 1% | 99% | 92% | 0% | 8% | 6% | 71% | 22% | 0.91 |
| 21d | 5 | 6 | 278 | 4% | 96% | 93% | 0% | 7% | 10% | 73% | 16% | 1.31 |
| 21d | 5 | 6 | 204 | 4% | 96% | 94% | 0% | 6% | 11% | 65% | 24% | 0.96 |
| 21d | 1 | 12 | 212 | 1% | 99% | 93% | 0% | 7% | 9% | 80% | 11% | 0.97 |
| 21d | 1 | 12 | 191 | 1% | 99% | 95% | 0% | 5% | 7% | 72% | 21% | 0.88 |

| | | | | | | | | | | | | |
|---|---|---|---|---|---|---|---|---|---|---|---|---|
| 21d | 1 | 12 | 293 | 4% | 96% | 98% | 0% | 2% | 13% | 72% | 16% | 1.31 |
| 21d | 1 | 12 | 269 | 3% | 97% | 97% | 0% | 3% | 11% | 79% | 10% | 1.21 |
| 21d | 10 | 12 | 216 | 3% | 97% | 93% | 0% | 7% | 10% | 81% | 9% | 0.98 |
| 21d | 10 | 12 | 209 | 5% | 95% | 94% | 0% | 6% | 8% | 77% | 15% | 0.92 |
| 21d | 10 | 12 | 245 | 2% | 98% | 96% | 0% | 4% | 10% | 81% | 10% | 1.12 |
| 21d | 10 | 12 | 254 | 4% | 96% | 93% | 0% | 7% | 7% | 79% | 14% | 1.13 |
| 21d | 5 | 12 | 128 | 5% | 95% | 96% | 0% | 4% | 5% | 84% | 10% | 0.56 |
| 21d | 5 | 12 | 193 | 1% | 99% | 94% | 0% | 6% | 10% | 76% | 14% | 0.89 |
| 21d | 5 | 12 | 293 | 3% | 97% | 94% | 0% | 6% | 10% | 77% | 13% | 1.32 |
| 21d | 5 | 12 | 223 | 4% | 96% | 96% | 0% | 4% | 13% | 75% | 12% | 1.00 |
| 21d | 1 | 24 | 230 | 1% | 99% | 93% | 0% | 7% | 40% | 50% | 10% | 0.95 |
| 21d | 1 | 24 | 203 | 1% | 99% | 93% | 0% | 8% | 35% | 49% | 16% | 0.84 |
| 21d | 1 | 24 | 317 | 3% | 97% | 96% | 0% | 4% | 49% | 36% | 16% | 1.29 |
| 21d | 1 | 24 | 292 | 3% | 97% | 94% | 0% | 6% | 44% | 45% | 11% | 1.19 |
| 21d | 10 | 24 | 233 | 1% | 99% | 89% | 0% | 11% | 45% | 46% | 9% | 0.97 |
| 21d | 10 | 24 | 232 | 1% | 99% | 88% | 0% | 12% | 38% | 57% | 5% | 0.97 |
| 21d | 10 | 24 | 268 | 2% | 98% | 93% | 0% | 7% | 45% | 45% | 11% | 1.11 |
| 21d | 10 | 24 | 282 | 3% | 97% | 91% | 0% | 9% | 43% | 45% | 12% | 1.15 |
| 21d | 5 | 24 | 142 | 3% | 97% | 93% | 0% | 7% | 39% | 53% | 8% | 0.58 |
| 21d | 5 | 24 | 212 | 4% | 96% | 92% | 0% | 8% | 41% | 49% | 10% | 0.85 |
| 21d | 5 | 24 | 330 | 3% | 97% | 92% | 0% | 8% | 45% | 46% | 9% | 1.35 |
| 21d | 5 | 24 | 247 | 2% | 98% | 93% | 0% | 7% | 43% | 42% | 15% | 1.02 |
| 21d | 1 | 48 | 260 | 2% | 98% | 95% | 0% | 5% | 58% | 33% | 9% | 0.84 |

| | | | | | | | | | | | | |
|---|---|---|---|---|---|---|---|---|---|---|---|---|
| 21d | 1 | 48 | 240 | 1% | 99% | 95% | 0% | 5% | 59% | 33% | 8% | 0.78 |
| 21d | 1 | 48 | 466 | 1% | 99% | 100% | 0% | 0% | 57% | 29% | 14% | 1.51 |
| 21d | 1 | 48 | 388 | 1% | 99% | 96% | 0% | 4% | 54% | 35% | 11% | 1.26 |
| 21d | 10 | 48 | 242 | 1% | 99% | 92% | 0% | 8% | 59% | 30% | 10% | 0.78 |
| 21d | 10 | 48 | 242 | 1% | 99% | 91% | 0% | 9% | 64% | 29% | 7% | 0.78 |
| 21d | 10 | 48 | 276 | 0% | 100% | 93% | 0% | 7% | 54% | 37% | 9% | 0.90 |
| 21d | 10 | 48 | 284 | 1% | 99% | 90% | 0% | 10% | 54% | 35% | 11% | 0.92 |
| 21d | 5 | 48 | 176 | 2% | 98% | 98% | 0% | 2% | 60% | 36% | 5% | 0.57 |
| 21d | 5 | 48 | 229 | 2% | 98% | 94% | 0% | 6% | 49% | 42% | 9% | 0.73 |
| 21d | 5 | 48 | 350 | 2% | 98% | 94% | 0% | 6% | 63% | 31% | 6% | 1.13 |
| 21d | 5 | 48 | 250 | 3% | 97% | 92% | 0% | 8% | 57% | 36% | 7% | 0.80 |
| 21d | 1 | 72 | 365 | 2% | 98% | 99% | 0% | 1% | 60% | 28% | 12% | 0.69 |
| 21d | 1 | 72 | 325 | 1% | 99% | 98% | 0% | 2% | 66% | 22% | 12% | 0.63 |
| 21d | 1 | 72 | 612 | 1% | 99% | 100% | 0% | 0% | 58% | 29% | 12% | 1.18 |
| 21d | 1 | 72 | 603 | 0% | 100% | 99% | 0% | 1% | 53% | 31% | 16% | 1.17 |
| 21d | 10 | 72 | 284 | 1% | 99% | 95% | 0% | 5% | 66% | 22% | 13% | 0.55 |
| 21d | 10 | 72 | 235 | 0% | 100% | 93% | 0% | 7% | 65% | 22% | 13% | 0.45 |
| 21d | 10 | 72 | 338 | 1% | 99% | 95% | 0% | 5% | 60% | 29% | 11% | 0.65 |
| 21d | 10 | 72 | 394 | 0% | 100% | 97% | 0% | 3% | 61% | 26% | 13% | 0.76 |
| 21d | 5 | 72 | 207 | 0% | 100% | 98% | 0% | 2% | 61% | 25% | 13% | 0.40 |
| 21d | 5 | 72 | 296 | 2% | 98% | 97% | 0% | 3% | 61% | 27% | 11% | 0.56 |
| 21d | 5 | 72 | 486 | 0% | 100% | 98% | 0% | 2% | 64% | 24% | 12% | 0.94 |
| 21d | 5 | 72 | 321 | 1% | 99% | 97% | 0% | 3% | 67% | 21% | 12% | 0.62 |

| | | | | | | | | | | | | |
|---|---|---|---|---|---|---|---|---|---|---|---|---|
| 1 | 1 | 0 | 189 | 1% | 99% | 96% | 2% | 3% | 76% | 14% | 10% | 0.79 |
| 1 | 1 | 0 | 185 | 0% | 100% | 96% | 3% | 1% | 76% | 14% | 11% | 0.78 |
| 1 | 1 | 0 | 234 | 0% | 100% | 97% | 0% | 2% | 74% | 14% | 12% | 0.99 |
| 1 | 1 | 0 | 251 | 0% | 100% | 96% | 2% | 2% | 73% | 15% | 11% | 1.06 |
| 1 | 10 | 0 | 160 | 0% | 100% | 96% | 1% | 3% | 73% | 15% | 12% | 0.68 |
| 1 | 10 | 0 | 168 | 0% | 100% | 99% | 1% | 1% | 76% | 14% | 10% | 0.71 |
| 1 | 10 | 0 | 201 | 0% | 100% | 99% | 0% | 1% | 70% | 16% | 14% | 0.85 |
| 1 | 10 | 0 | 255 | 1% | 99% | 95% | 4% | 0% | 73% | 16% | 11% | 1.07 |
| 1 | 5 | 0 | 132 | 0% | 100% | 97% | 2% | 1% | 76% | 13% | 11% | 0.56 |
| 1 | 5 | 0 | 150 | 0% | 100% | 95% | 3% | 2% | 66% | 17% | 17% | 0.63 |
| 1 | 5 | 0 | 229 | 0% | 100% | 97% | 1% | 1% | 78% | 13% | 9% | 0.97 |
| 1 | 5 | 0 | 235 | 0% | 100% | 94% | 2% | 4% | 75% | 14% | 12% | 0.99 |
| 1 | 1 | 6 | 155 | 3% | 97% | 89% | 0% | 11% | 18% | 51% | 31% | 0.74 |
| 1 | 1 | 6 | 157 | 3% | 97% | 92% | 0% | 8% | 18% | 53% | 29% | 0.75 |
| 1 | 1 | 6 | 188 | 4% | 96% | 91% | 1% | 8% | 23% | 47% | 30% | 0.89 |
| 1 | 1 | 6 | 209 | 5% | 95% | 92% | 0% | 8% | 18% | 51% | 31% | 0.97 |
| 1 | 10 | 6 | 130 | 2% | 98% | 88% | 0% | 12% | 14% | 59% | 27% | 0.62 |
| 1 | 10 | 6 | 140 | 4% | 96% | 93% | 0% | 7% | 16% | 59% | 25% | 0.66 |
| 1 | 10 | 6 | 151 | 4% | 96% | 86% | 0% | 14% | 17% | 46% | 37% | 0.71 |
| 1 | 10 | 6 | 198 | 6% | 94% | 88% | 0% | 12% | 12% | 57% | 30% | 0.92 |
| 1 | 5 | 6 | 115 | 4% | 96% | 88% | 1% | 11% | 19% | 61% | 21% | 0.54 |
| 1 | 5 | 6 | 133 | 5% | 95% | 90% | 0% | 10% | 12% | 68% | 20% | 0.62 |
| 1 | 5 | 6 | 168 | 4% | 96% | 85% | 0% | 15% | 12% | 55% | 33% | 0.79 |

| | | | | | | | | | | | | |
|---|---|---|---|---|---|---|---|---|---|---|---|---|
| 1 | 5 | 6 | 190 | 5% | 95% | 83% | 1% | 16% | 20% | 58% | 23% | 0.89 |
| 1 | 1 | 12 | 165 | 5% | 95% | 84% | 0% | 16% | 45% | 37% | 18% | 0.72 |
| 1 | 1 | 12 | 163 | 3% | 97% | 85% | 0% | 15% | 37% | 43% | 21% | 0.73 |
| 1 | 1 | 12 | 203 | 3% | 97% | 90% | 0% | 10% | 46% | 36% | 18% | 0.91 |
| 1 | 1 | 12 | 220 | 3% | 97% | 87% | 0% | 13% | 42% | 43% | 15% | 0.99 |
| 1 | 10 | 12 | 144 | 3% | 97% | 68% | 0% | 32% | 27% | 48% | 24% | 0.65 |
| 1 | 10 | 12 | 151 | 5% | 95% | 76% | 0% | 24% | 30% | 42% | 28% | 0.66 |
| 1 | 10 | 12 | 167 | 7% | 93% | 68% | 0% | 32% | 32% | 41% | 27% | 0.72 |
| 1 | 10 | 12 | 215 | 7% | 93% | 73% | 0% | 27% | 35% | 46% | 19% | 0.92 |
| 1 | 5 | 12 | 126 | 2% | 98% | 81% | 1% | 18% | 38% | 47% | 15% | 0.57 |
| 1 | 5 | 12 | 138 | 9% | 91% | 79% | 0% | 21% | 33% | 48% | 19% | 0.59 |
| 1 | 5 | 12 | 194 | 4% | 96% | 78% | 0% | 22% | 39% | 44% | 17% | 0.87 |
| 1 | 5 | 12 | 204 | 6% | 94% | 76% | 0% | 24% | 41% | 41% | 18% | 0.89 |
| 1 | 1 | 24 | 191 | 3% | 97% | 70% | 0% | 30% | 52% | 34% | 15% | 0.78 |
| 1 | 1 | 24 | 182 | 5% | 95% | 76% | 0% | 24% | 50% | 34% | 15% | 0.73 |
| 1 | 1 | 24 | 223 | 5% | 95% | 77% | 0% | 23% | 64% | 23% | 12% | 0.89 |
| 1 | 1 | 24 | 238 | 8% | 92% | 75% | 0% | 25% | 53% | 34% | 13% | 0.92 |
| 1 | 10 | 24 | 152 | 6% | 94% | 58% | 0% | 42% | 41% | 39% | 20% | 0.60 |
| 1 | 10 | 24 | 158 | 10% | 90% | 56% | 0% | 44% | 45% | 33% | 23% | 0.60 |
| 1 | 10 | 24 | 186 | 7% | 93% | 60% | 0% | 40% | 50% | 35% | 15% | 0.73 |
| 1 | 10 | 24 | 232 | 7% | 93% | 60% | 0% | 40% | 46% | 34% | 20% | 0.90 |
| 1 | 5 | 24 | 132 | 7% | 93% | 72% | 0% | 28% | 42% | 40% | 18% | 0.52 |
| 1 | 5 | 24 | 135 | 8% | 92% | 65% | 0% | 35% | 50% | 31% | 19% | 0.52 |

| | | | | | | | | | | | | |
|---|---|---|---|---|---|---|---|---|---|---|---|---|
| 1 | 5 | 24 | 196 | 5% | 95% | 68% | 0% | 32% | 48% | 35% | 17% | 0.78 |
| 1 | 5 | 24 | 225 | 9% | 91% | 58% | 0% | 42% | 44% | 37% | 19% | 0.86 |
| 1 | 1 | 48 | 189 | 6% | 94% | 67% | 0% | 33% | 55% | 31% | 14% | 0.58 |
| 1 | 1 | 48 | 174 | 6% | 94% | 64% | 0% | 36% | 49% | 28% | 24% | 0.53 |
| 1 | 1 | 48 | 229 | 10% | 90% | 77% | 0% | 23% | 53% | 31% | 16% | 0.67 |
| 1 | 1 | 48 | 244 | 5% | 95% | 70% | 0% | 30% | 51% | 29% | 21% | 0.76 |
| 1 | 10 | 48 | 149 | 15% | 85% | 45% | 0% | 55% | 47% | 42% | 11% | 0.42 |
| 1 | 10 | 48 | 156 | 12% | 88% | 50% | 0% | 50% | 39% | 45% | 16% | 0.45 |
| 1 | 10 | 48 | 181 | 7% | 93% | 40% | 0% | 60% | 43% | 43% | 15% | 0.55 |
| 1 | 10 | 48 | 239 | 9% | 91% | 52% | 0% | 48% | 41% | 37% | 22% | 0.71 |
| 1 | 5 | 48 | 129 | 9% | 91% | 65% | 0% | 35% | 43% | 38% | 18% | 0.38 |
| 1 | 5 | 48 | 136 | 10% | 90% | 57% | 1% | 42% | 47% | 36% | 17% | 0.40 |
| 1 | 5 | 48 | 195 | 8% | 92% | 62% | 0% | 38% | 51% | 41% | 8% | 0.59 |
| 1 | 5 | 48 | 213 | 9% | 91% | 52% | 1% | 47% | 39% | 44% | 18% | 0.63 |
| 1 | 1 | 72 | 188 | 6% | 94% | 65% | 0% | 35% | 51% | 33% | 16% | 0.34 |
| 1 | 1 | 72 | 175 | 7% | 93% | 68% | 0% | 32% | 60% | 27% | 13% | 0.31 |
| 1 | 1 | 72 | 215 | 8% | 92% | 73% | 0% | 27% | 56% | 31% | 13% | 0.38 |
| 1 | 1 | 72 | 230 | 8% | 92% | 62% | 0% | 38% | 48% | 34% | 17% | 0.41 |
| 1 | 10 | 72 | 151 | 11% | 89% | 44% | 0% | 56% | 29% | 56% | 15% | 0.26 |
| 1 | 10 | 72 | 157 | 7% | 93% | 42% | 0% | 58% | 38% | 52% | 10% | 0.28 |
| 1 | 10 | 72 | 182 | 8% | 92% | 41% | 0% | 59% | 43% | 46% | 10% | 0.32 |
| 1 | 10 | 72 | 222 | 9% | 91% | 35% | 0% | 65% | 39% | 41% | 20% | 0.39 |
| 1 | 5 | 72 | 135 | 10% | 90% | 57% | 0% | 43% | 39% | 43% | 17% | 0.24 |

| | | | | | | | | | | | | |
|---|---|---|---|---|---|---|---|---|---|---|---|---|
| 1 | 5 | 72 | 132 | 8% | 92% | 54% | 1% | 45% | 50% | 30% | 20% | 0.24 |
| 1 | 5 | 72 | 188 | 9% | 91% | 53% | 0% | 47% | 49% | 37% | 14% | 0.33 |
| 1 | 5 | 72 | 222 | 9% | 91% | 48% | 0% | 52% | 32% | 49% | 19% | 0.39 |
| milciclib | 1 | 0 | 201 | 0% | 100% | 96% | 1% | 3% | 74% | 16% | 11% | 0.85 |
| milciclib | 1 | 0 | 261 | 0% | 100% | 95% | 2% | 3% | 77% | 11% | 13% | 1.10 |
| milciclib | 1 | 0 | 302 | 0% | 100% | 96% | 2% | 2% | 73% | 16% | 11% | 1.28 |
| milciclib | 10 | 0 | 237 | 0% | 100% | 96% | 1% | 3% | 77% | 11% | 12% | 1.00 |
| milciclib | 10 | 0 | 297 | 0% | 100% | 94% | 2% | 4% | 74% | 15% | 11% | 1.26 |
| milciclib | 10 | 0 | 258 | 0% | 100% | 96% | 2% | 2% | 71% | 17% | 12% | 1.09 |
| milciclib | 1 | 6 | 181 | 2% | 98% | 93% | 0% | 7% | 9% | 74% | 18% | 0.87 |
| milciclib | 1 | 6 | 214 | 2% | 98% | 92% | 0% | 8% | 8% | 71% | 21% | 1.03 |
| milciclib | 1 | 6 | 251 | 2% | 98% | 93% | 0% | 7% | 9% | 83% | 7% | 1.20 |
| milciclib | 10 | 6 | 179 | 2% | 98% | 94% | 0% | 6% | 11% | 73% | 16% | 0.86 |
| milciclib | 10 | 6 | 233 | 3% | 97% | 89% | 0% | 11% | 19% | 69% | 12% | 1.11 |
| milciclib | 10 | 6 | 195 | 3% | 97% | 91% | 1% | 8% | 21% | 67% | 12% | 0.93 |
| milciclib | 1 | 12 | 191 | 3% | 97% | 93% | 0% | 7% | 13% | 75% | 12% | 0.86 |
| milciclib | 1 | 12 | 236 | 4% | 96% | 91% | 0% | 9% | 12% | 76% | 12% | 1.05 |
| milciclib | 1 | 12 | 267 | 1% | 99% | 90% | 0% | 10% | 16% | 76% | 8% | 1.23 |
| milciclib | 10 | 12 | 187 | 3% | 97% | 90% | 0% | 10% | 38% | 50% | 12% | 0.84 |
| milciclib | 10 | 12 | 226 | 3% | 97% | 84% | 0% | 16% | 36% | 52% | 13% | 1.02 |
| milciclib | 10 | 12 | 198 | 3% | 97% | 86% | 0% | 14% | 39% | 47% | 13% | 0.89 |
| milciclib | 1 | 24 | 183 | 4% | 96% | 89% | 0% | 11% | 70% | 26% | 4% | 0.74 |
| milciclib | 1 | 24 | 228 | 4% | 96% | 87% | 0% | 13% | 65% | 30% | 5% | 0.93 |

| | | | | | | | | | | | | |
|---|---|---|---|---|---|---|---|---|---|---|---|---|
| milciclib | 1 | 24 | 254 | 3% | 97% | 90% | 0% | 10% | 68% | 27% | 5% | 1.04 |
| milciclib | 10 | 24 | 197 | 7% | 93% | 79% | 0% | 21% | 59% | 31% | 10% | 0.77 |
| milciclib | 10 | 24 | 231 | 3% | 97% | 72% | 0% | 28% | 59% | 32% | 9% | 0.94 |
| milciclib | 10 | 24 | 191 | 4% | 96% | 80% | 1% | 20% | 60% | 27% | 14% | 0.77 |
| milciclib | 1 | 48 | 160 | 1% | 99% | 90% | 0% | 10% | 71% | 20% | 9% | 0.52 |
| milciclib | 1 | 48 | 178 | 2% | 98% | 91% | 1% | 8% | 77% | 15% | 8% | 0.57 |
| milciclib | 1 | 48 | 184 | 3% | 97% | 94% | 0% | 6% | 70% | 22% | 8% | 0.59 |
| milciclib | 10 | 48 | 190 | 5% | 95% | 73% | 0% | 27% | 60% | 24% | 16% | 0.59 |
| milciclib | 10 | 48 | 231 | 7% | 93% | 75% | 0% | 25% | 61% | 22% | 17% | 0.70 |
| milciclib | 10 | 48 | 199 | 3% | 97% | 75% | 1% | 24% | 66% | 19% | 15% | 0.63 |
| milciclib | 1 | 72 | 150 | 1% | 99% | 93% | 0% | 7% | 65% | 26% | 9% | 0.29 |
| milciclib | 1 | 72 | 164 | 2% | 98% | 93% | 0% | 7% | 66% | 22% | 12% | 0.31 |
| milciclib | 1 | 72 | 170 | 1% | 99% | 98% | 0% | 2% | 64% | 24% | 12% | 0.33 |
| milciclib | 10 | 72 | 180 | 6% | 94% | 62% | 0% | 38% | 54% | 30% | 16% | 0.33 |
| milciclib | 10 | 72 | 244 | 6% | 94% | 68% | 0% | 32% | 54% | 30% | 17% | 0.44 |
| milciclib | 10 | 72 | 199 | 5% | 95% | 70% | 0% | 30% | 53% | 32% | 16% | 0.37 |

**Table S6.** Results of the wild-type kinase profiling of **21c** provided by Reaction Biology at a screening concentration of 1 µM.

| **21c** | |
|---|---|
| **Kinase** | **Remaining activity, percent of control [%]** |
| MST3 | 13.5 |
| MST4 | 18.0 |
| LIMK1 | 36.6 |
| LIMK2 | 38.2 |
| PKCt | 42.6 |
| MERTK | 44.2 |
| MAP3K11 | 45.8 |
| MELK | 47.0 |
| STK39 | 49.5 |
| FYN | 56.6 |
| STK25 | 57.2 |
| SLK | 61.9 |
| CLK2 | 64.6 |
| GSK3A | 67.3 |
| EPHA4 | 67.5 |
| NIK | 68.5 |
| MYLK | 68.9 |
| KIT | 69.4 |
| MINK1 | 70.0 |
| MEK5 | 70.1 |
| CDK17 | 70.9 |
| CDK12 | 71.1 |
| PDGFRb | 71.2 |
| SIK3 | 72.1 |
| RPS6KA6 | 73.5 |
| CSK | 73.5 |
| AMPKa1 | 74.0 |

| | |
|---|---|
| HIPK2 | 74.8 |
| EIF2AK2 | 75.1 |
| FGR | 75.3 |
| CLK4 | 76.4 |
| EPHB1 | 77.1 |
| TYRO3 | 77.8 |
| PKCnu | 78.1 |
| SNK | 78.3 |
| MATK | 78.6 |
| MAP4K5 | 78.8 |
| TRKB | 79.0 |
| CLK1 | 79.0 |
| MYLK2 | 79.0 |
| NEK1 | 79.1 |
| PKCg | 79.4 |
| FGFR2 | 79.5 |
| DYRK1B | 79.7 |
| ACK1 | 79.8 |
| BMPR1B | 80.1 |
| AXL | 80.2 |
| STK23 | 80.7 |
| IKKa | 80.7 |
| PYK2 | 80.8 |
| FGFR1 | 80.9 |
| MAP4K4 | 81.0 |
| ACVR1 | 81.0 |
| YES | 81.2 |
| GSK3B | 81.4 |
| SRMS | 81.5 |
| TGFBR2 | 81.8 |
| PRKD2 | 82.0 |
| DYRK1A | 82.2 |
| PKCb | 82.5 |

| | |
|---|---|
| EPHA2 | 82.5 |
| ZAK | 82.9 |
| MUSK | 83.0 |
| CDK9 | 83.3 |
| PKCi | 83.5 |
| AurA | 83.6 |
| AurB | 83.9 |
| CHK2 | 84.0 |
| JNK2 | 84.0 |
| RPS6KA2 | 84.0 |
| ABL2 | 84.2 |
| CDK20 | 84.4 |
| CDK4 | 84.5 |
| BMX | 84.8 |
| JNK3 | 84.9 |
| CDK10 | 85.1 |
| ACVR1B | 85.2 |
| CDK6/CycD3 | 85.3 |
| CK2a2 | 85.6 |
| CDK9/CycT1 | 85.7 |
| CDK6/CycD2 | 85.8 |
| ERK1 | 85.8 |
| EPHB4 | 86.1 |
| RPS6KB1 | 86.2 |
| SIK1 | 86.3 |
| ARK5 | 86.3 |
| MYLK3 | 86.4 |
| PKCe | 86.5 |
| SAK | 86.5 |
| NDR1 | 86.7 |
| CDK3/CycC | 87.0 |
| PAK7 | 87.1 |
| CDK1/CycB1 | 87.2 |

| | |
|---|---|
| IRAK4 | 87.3 |
| RPS6KA3 | 87.3 |
| GRK6 | 87.6 |
| EIF2AK3 | 87.8 |
| WEE1 | 88.0 |
| JAK1 | 88.0 |
| CDK8 | 88.5 |
| p38g | 88.6 |
| CLK3 | 88.6 |
| STK33 | 88.7 |
| CAMK2B | 88.8 |
| PIM3 | 88.9 |
| CDK20/CycT1 | 89.1 |
| GRK7 | 89.2 |
| PLK1 | 89.3 |
| NEK6 | 89.5 |
| PRK2 | 89.5 |
| CDK1 | 89.7 |
| BRAF | 89.7 |
| ROCK2 | 89.7 |
| SGK1 | 90.0 |
| AurC | 90.2 |
| MEK2 | 90.2 |
| ERK7 | 90.2 |
| LRRK2 | 90.3 |
| VEGFR2 | 90.3 |
| AKT2 | 90.3 |
| RPS6KA1 | 90.5 |
| ERBB4 | 90.7 |
| ERK2 | 90.7 |
| HIPK3 | 90.8 |
| TRKA | 90.8 |
| CAMKK2 | 90.8 |

| | |
|---|---|
| CSF1R | 90.9 |
| ACVRL1 | 91.0 |
| CDK11B | 91.2 |
| DYRK2 | 91.4 |
| PKCd | 91.4 |
| TEC | 91.4 |
| FAK | 91.4 |
| VRK1 | 91.5 |
| PRK1 | 91.8 |
| CDK1/CycA2 | 91.9 |
| LTK | 92.0 |
| LCK | 92.0 |
| AKT1 | 92.1 |
| TLK1 | 92.4 |
| RPS6KA4 | 92.5 |
| PKN3 | 92.6 |
| p38a | 92.7 |
| CDK19 | 92.8 |
| p38b | 92.9 |
| NEK11 | 93.3 |
| ERK5 | 93.4 |
| NEK2 | 93.5 |
| PRKG2 | 93.5 |
| TAOK2 | 93.6 |
| TSK2 | 93.8 |
| PRKX | 93.9 |
| NDR2 | 93.9 |
| FLT4 | 93.9 |
| DAPK1 | 94.0 |
| HCK | 94.0 |
| TSF1 | 94.0 |
| IKKb | 94.0 |
| ERBB2 | 94.1 |

| | |
|---|---|
| CDK16 | 94.1 |
| NEK3 | 94.1 |
| DMPK | 94.3 |
| MAP3K10 | 94.4 |
| MAP3K7 | 94.4 |
| RON | 94.5 |
| CAMKK1 | 94.6 |
| WNK2 | 94.8 |
| PRKG1 | 94.8 |
| SRC | 94.8 |
| p38d | 94.8 |
| PAK1 | 94.9 |
| WNK1 | 94.9 |
| CAMK2D | 95.0 |
| PAK6 | 95.0 |
| PAK4 | 95.1 |
| ACVR2A | 95.3 |
| TLK2 | 95.4 |
| CK2a1 | 95.5 |
| DYRK4 | 95.6 |
| SNARK | 95.6 |
| MST2 | 95.6 |
| JNK1 | 95.6 |
| MAPKAPK3 | 95.7 |
| MAP3K9 | 95.7 |
| TSSK1 | 95.8 |
| CDK3 | 95.9 |
| ULK2 | 96.0 |
| MARK1 | 96.0 |
| PKCmu | 96.1 |
| SGK3 | 96.1 |
| JAK2 | 96.2 |
| PKMzeta | 96.2 |

| | |
|---|---|
| TNK1 | 96.2 |
| PKCz | 96.3 |
| PKA | 96.3 |
| DYRK3 | 96.4 |
| INSRR | 96.5 |
| BTK | 96.6 |
| CDK5/p35NCK | 96.6 |
| GRK2 | 96.7 |
| CDK18 | 96.7 |
| FGFR4 | 96.9 |
| VRK2 | 96.9 |
| SRPK2 | 97.0 |
| HIPK1 | 97.2 |
| MKNK1 | 97.3 |
| EPHA6 | 97.4 |
| GRK3 | 97.4 |
| SIK2 | 97.5 |
| TTK | 97.7 |
| ASK1 | 97.7 |
| EPHA3 | 97.7 |
| BRSK2 | 97.8 |
| CAMK4 | 98.0 |
| INSR | 98.0 |
| PHKG1 | 98.0 |
| PHKG2 | 98.0 |
| PKCh | 98.2 |
| HRI | 98.2 |
| TXK | 98.2 |
| ROCK1 | 98.4 |
| EPHA5 | 98.4 |
| WNK3 | 98.5 |
| BLK | 98.6 |
| CK1d | 98.8 |

| | |
|---|---|
| PBK | 98.8 |
| NEK7 | 98.8 |
| MARK2 | 98.8 |
| RPS6KB2 | 99.0 |
| MLK4 | 99.1 |
| MTOR | 99.2 |
| BRK | 99.3 |
| EPHA1 | 99.3 |
| DCAMKL2 | 99.3 |
| PDK1 | 99.5 |
| SYK | 99.7 |
| ITK | 99.7 |
| FGFR3 | 99.8 |
| EGFR | 100.0 |
| MARK4 | 100.1 |
| ALK | 100.1 |
| MAP3K1 | 100.1 |
| EPHB2 | 100.1 |
| DAPK2 | 100.3 |
| MKK7 | 100.3 |
| VEGFR1 | 100.3 |
| ACVR2B | 100.5 |
| FES | 100.7 |
| DNAPK | 100.7 |
| PKCbeta2 | 100.8 |
| CK1g3 | 100.8 |
| PASK | 100.8 |
| MARK3 | 101.1 |
| RIPK4 | 101.1 |
| CDK13 | 101.2 |
| PIM1 | 101.3 |
| CDK5 | 101.3 |
| CDC7 | 101.4 |

| | |
|---|---|
| SRPK1 | 101.4 |
| BUB1B | 101.6 |
| FER | 101.7 |
| IRAK1 | 101.8 |
| TRKC | 102.0 |
| CK1a | 102.0 |
| PDGFRa | 102.0 |
| CK1g1 | 102.2 |
| TAOK3 | 102.4 |
| AKT3 | 102.5 |
| PLK3 | 102.7 |
| IGF1R | 102.9 |
| DDR2 | 103.0 |
| CDK2 | 103.0 |
| CAMK1D | 103.0 |
| CK1e | 103.1 |
| PIM2 | 103.1 |
| PAK3 | 103.3 |
| FLT3 | 103.4 |
| STK17A | 103.5 |
| MEKK2 | 103.5 |
| IKKe | 103.6 |
| EPHA7 | 103.7 |
| GRK5 | 103.8 |
| NEK4 | 103.8 |
| FRK | 104.0 |
| RIPK2 | 105.0 |
| LKB1/MO25a/STRADa | 105.1 |
| CDK6 | 105.2 |
| EPHB3 | 105.4 |
| SGK2 | 105.5 |
| MST1 | 105.6 |
| MKNK2 | 105.8 |

| | |
|---|---|
| MEKK3 | 105.8 |
| MAPKAPK2 | 106.1 |
| CDK2/CycA2 | 106.1 |
| ABL1 | 106.1 |
| CDC42BPB | 106.6 |
| CDK4/CycD2 | 106.8 |
| DAPK3 | 107.0 |
| CDC42BPA | 107.0 |
| MET | 107.2 |
| PKCa | 107.7 |
| GSG2 | 108.1 |
| BMPR1A | 108.3 |
| ZAP70 | 108.6 |
| RPS6KA5 | 108.9 |
| ROS | 109.6 |
| CK1g2 | 109.6 |
| NEK9 | 109.8 |
| TTBK2 | 109.9 |
| MAPKAPK5 | 110.4 |
| CAMK2A | 110.6 |
| CDK4/CycD3 | 111.1 |
| TBK1 | 111.6 |
| TTBK1 | 111.9 |
| NLK | 111.9 |
| CHK1 | 112.0 |
| GRK4 | 112.1 |
| EPHA8 | 112.4 |
| CDK7 | 112.5 |
| MAP4K2 | 112.8 |
| BRSK1 | 112.9 |
| PAK2 | 113.2 |
| EEF2K | 114.2 |
| COT | 115.1 |

| | |
|---|---|
| HIPK4 | 115.2 |
| JAK3 | 115.4 |
| TYK2 | 116.8 |
| TIE2 | 116.9 |
| ARAF YDYD | 117.0 |
| MASTL | 118.0 |
| MEK1 | 118.7 |
| CDK2/CycD1 | 121.3 |
| RIPK5 | 122.0 |
| MKK6 SDTD | 122.2 |
| RET | 122.5 |
| TGFBR1 | 124.6 |
| MKK4 | 127.3 |
| RAF1 YDYD | 130.3 |
| CAMK2G | 132.6 |
| LYN | 139.3 |

Analytical data of compounds **7** – **21**.

ESI, $^1$H and $^{13}$C NMR data of compound **7**.

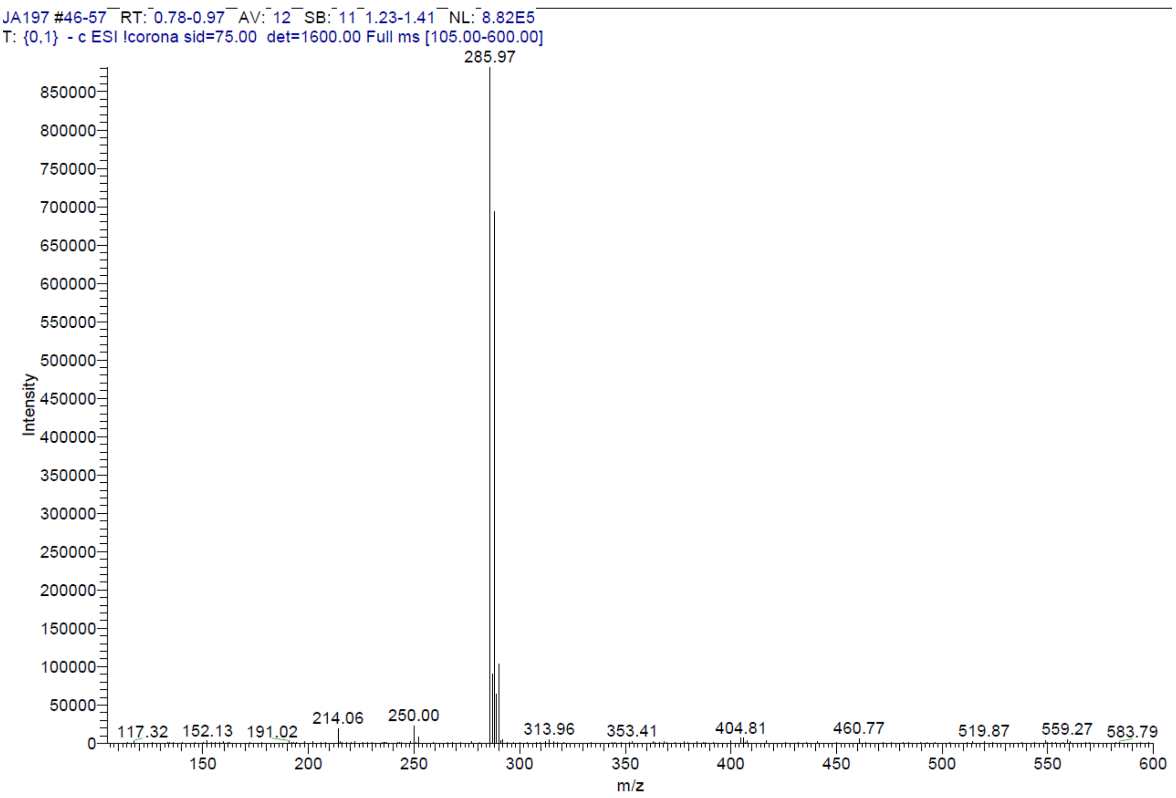


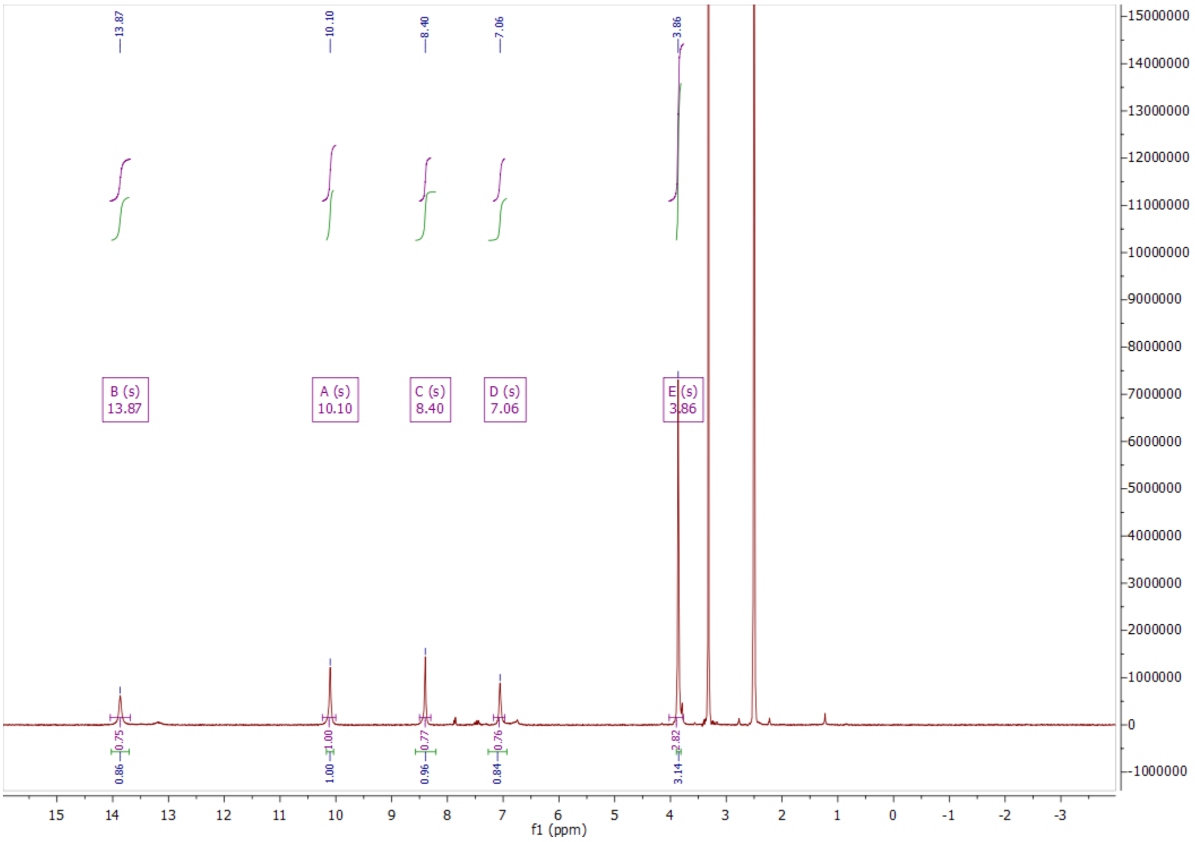

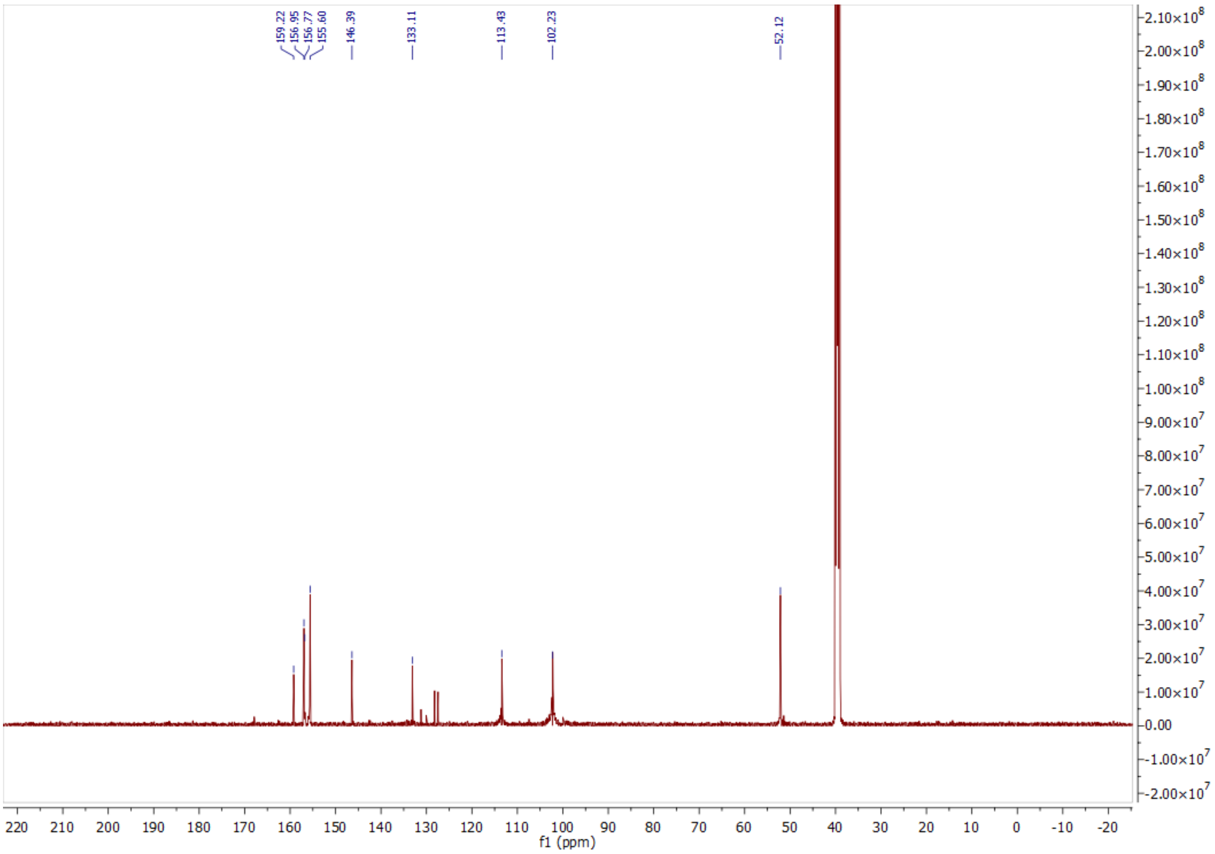


ESI, [1]H and [13]C NMR data of compound **8**.

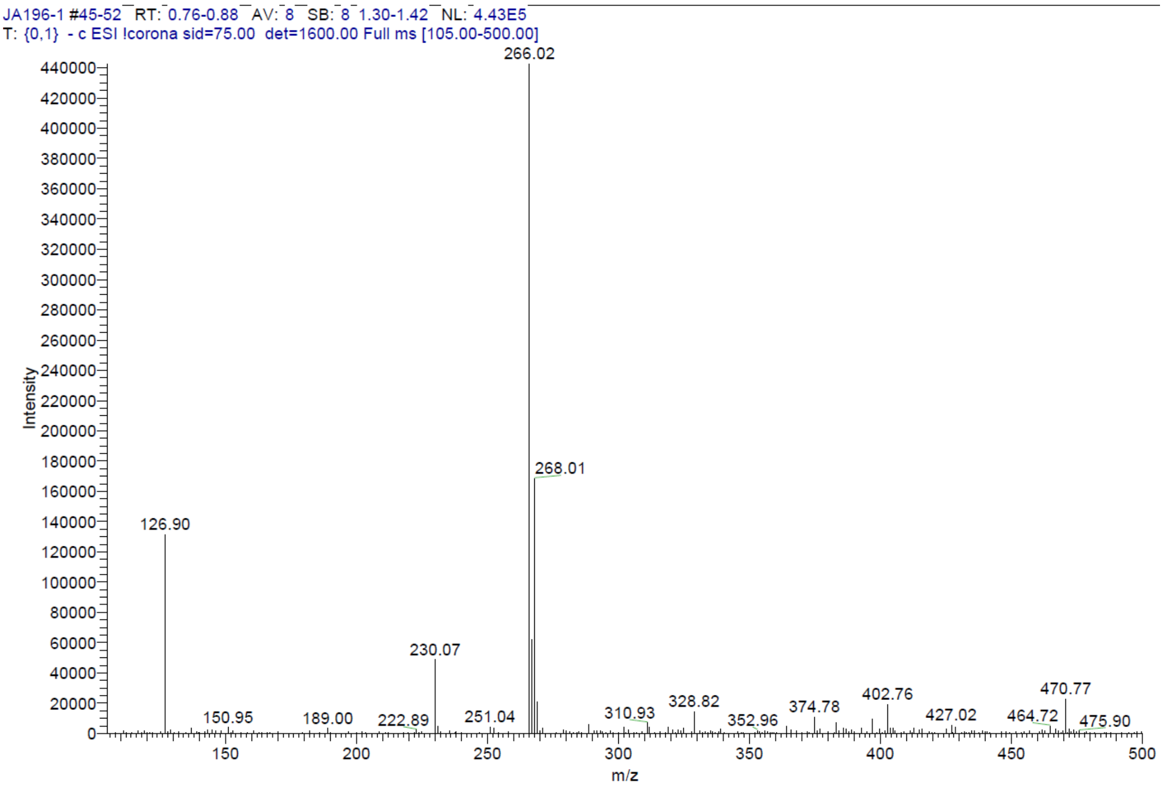

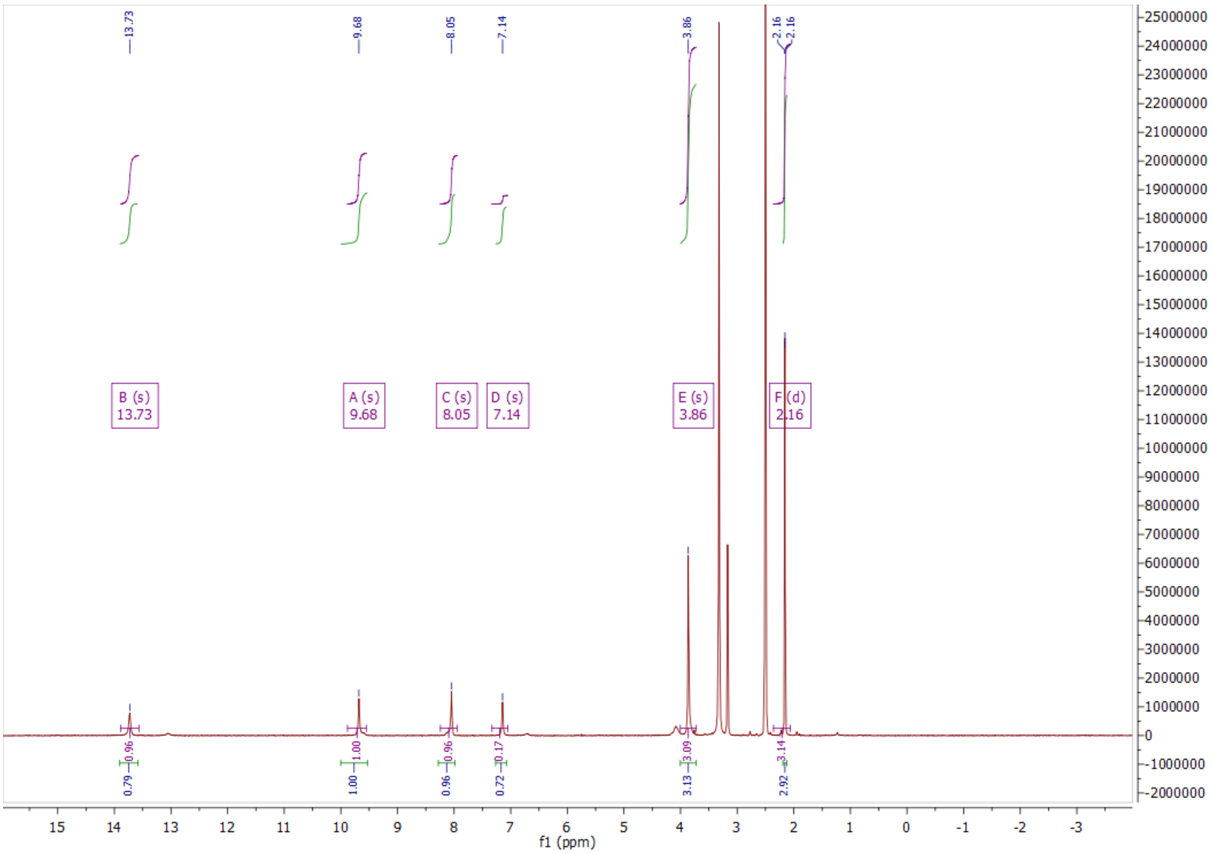
13.73
9.68
8.05
7.14
3.86
2.16
2.16
B (s)
13.73
A (s)
9.68
C (s)
8.05
D (s)
7.14
E (s)
3.86
F (d)
2.16
0.79
0.96
1.00
1.00
0.96
0.96
0.72
0.17
3.13
3.09
2.92
3.14
f1 (ppm)

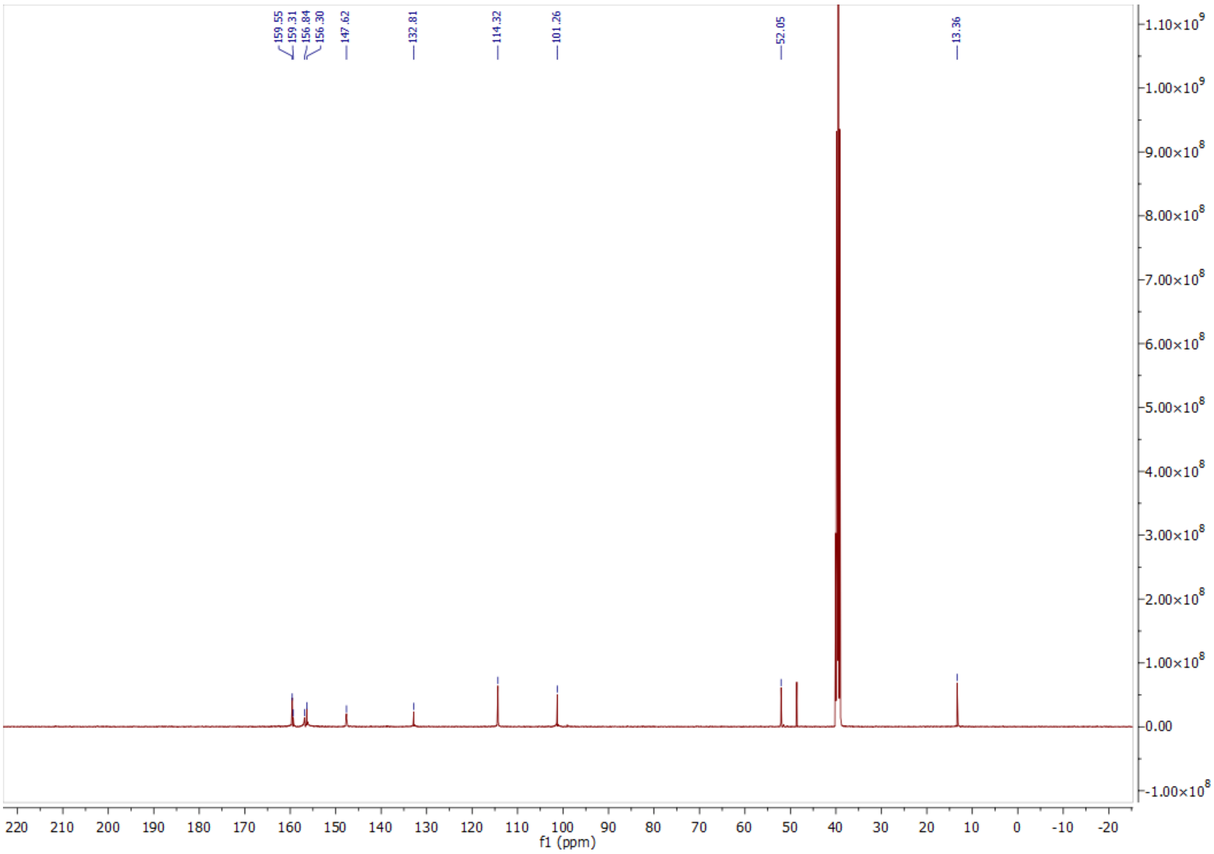
159.55
159.31
156.84
156.30
147.62
132.81
114.32
101.26
52.05
13.36
f1 (ppm)

ESI, $^{1}$H and $^{13}$C NMR data of compound **9**.

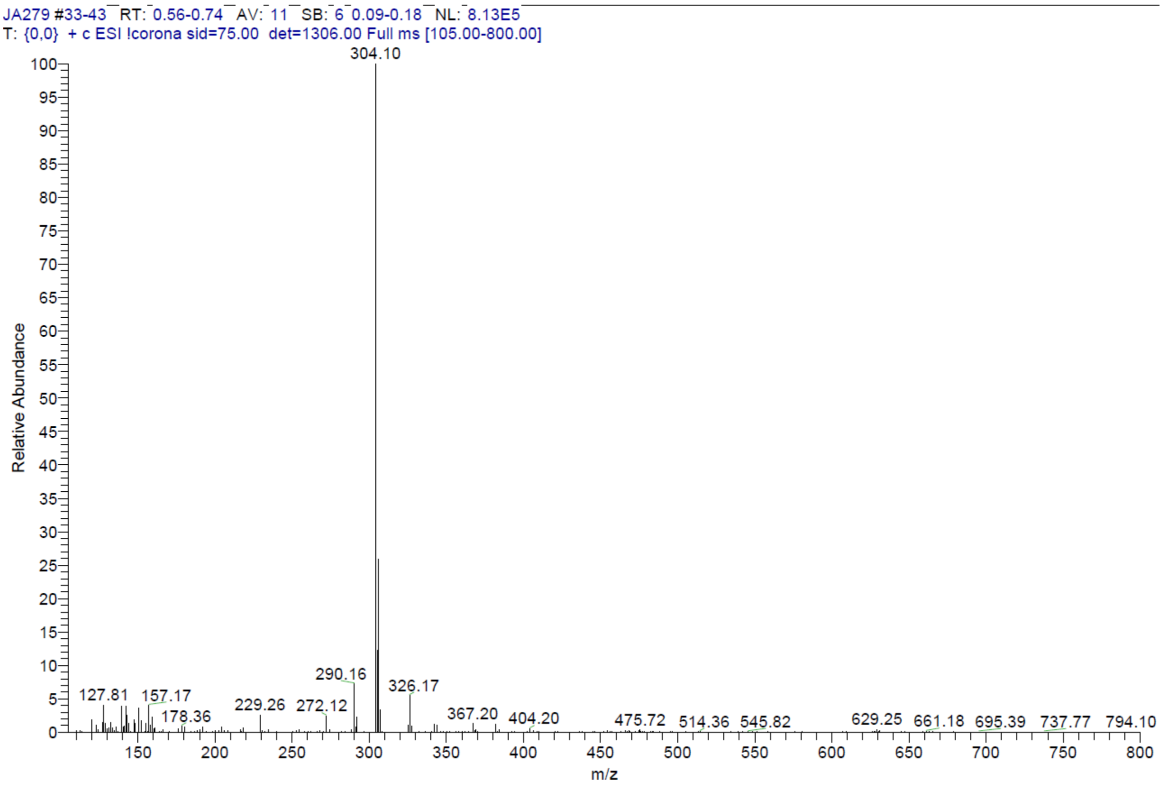


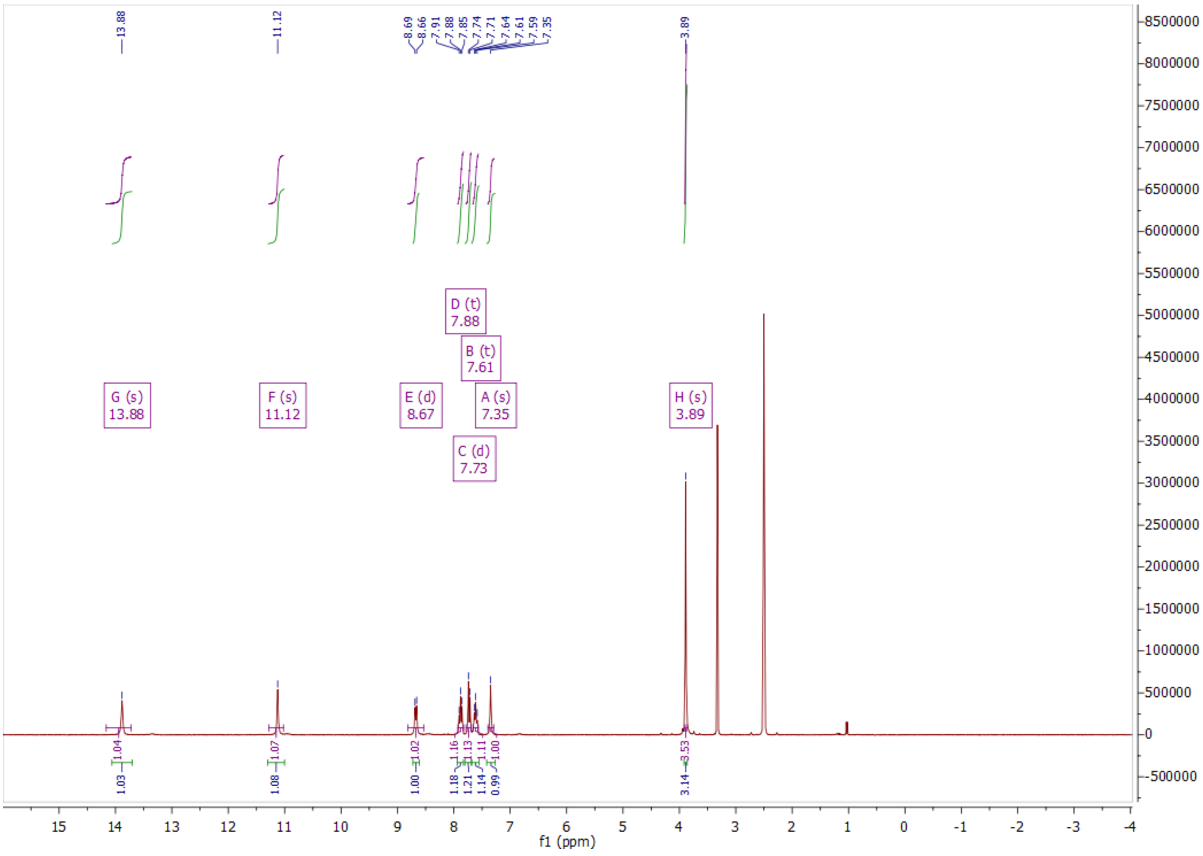

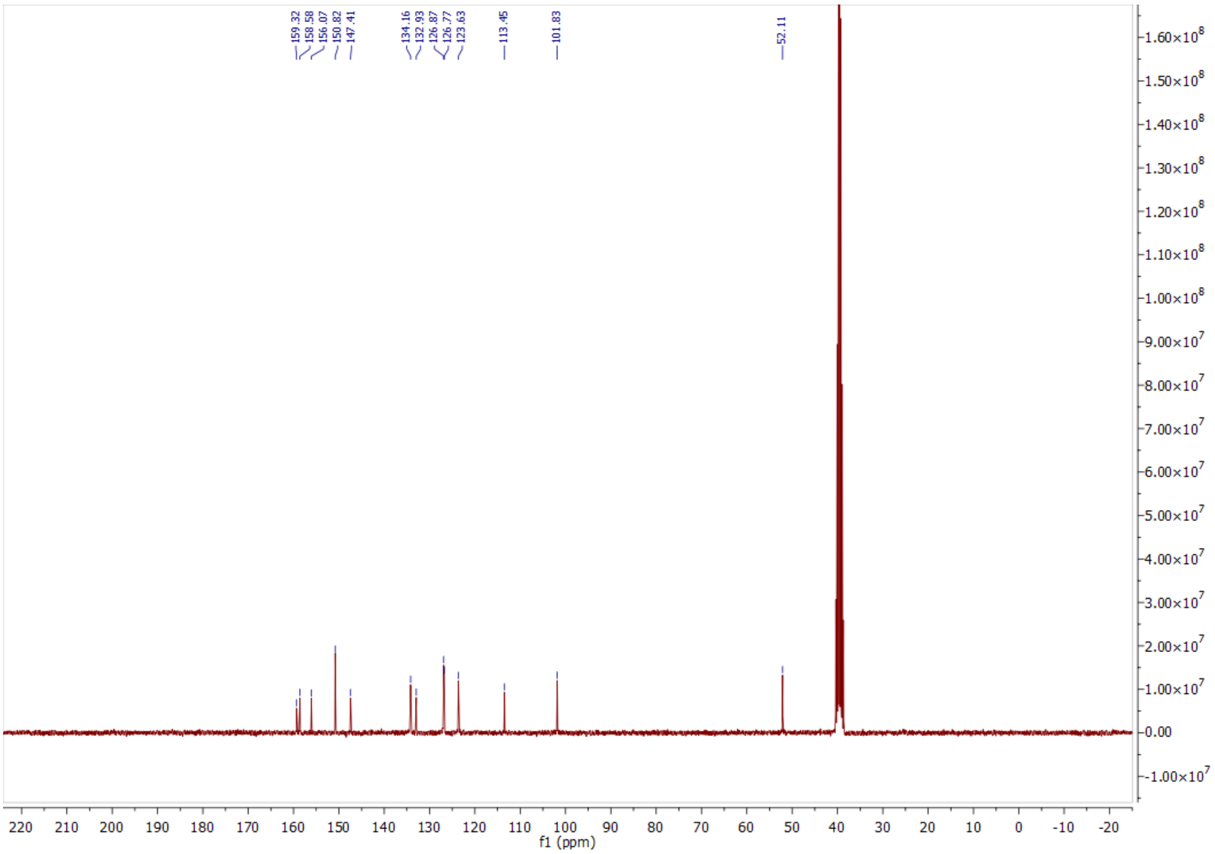


ESI, $^{1}H$, $^{13}C$ NMR and HPLC data of compound **10a**.

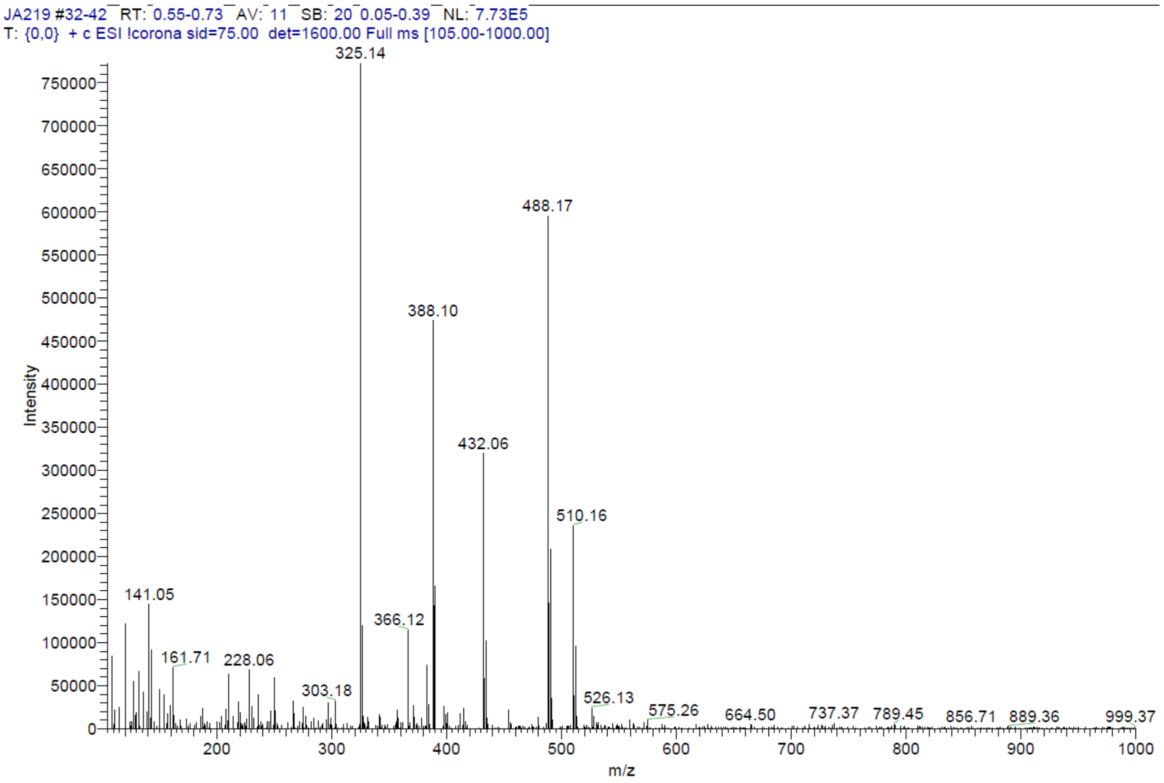

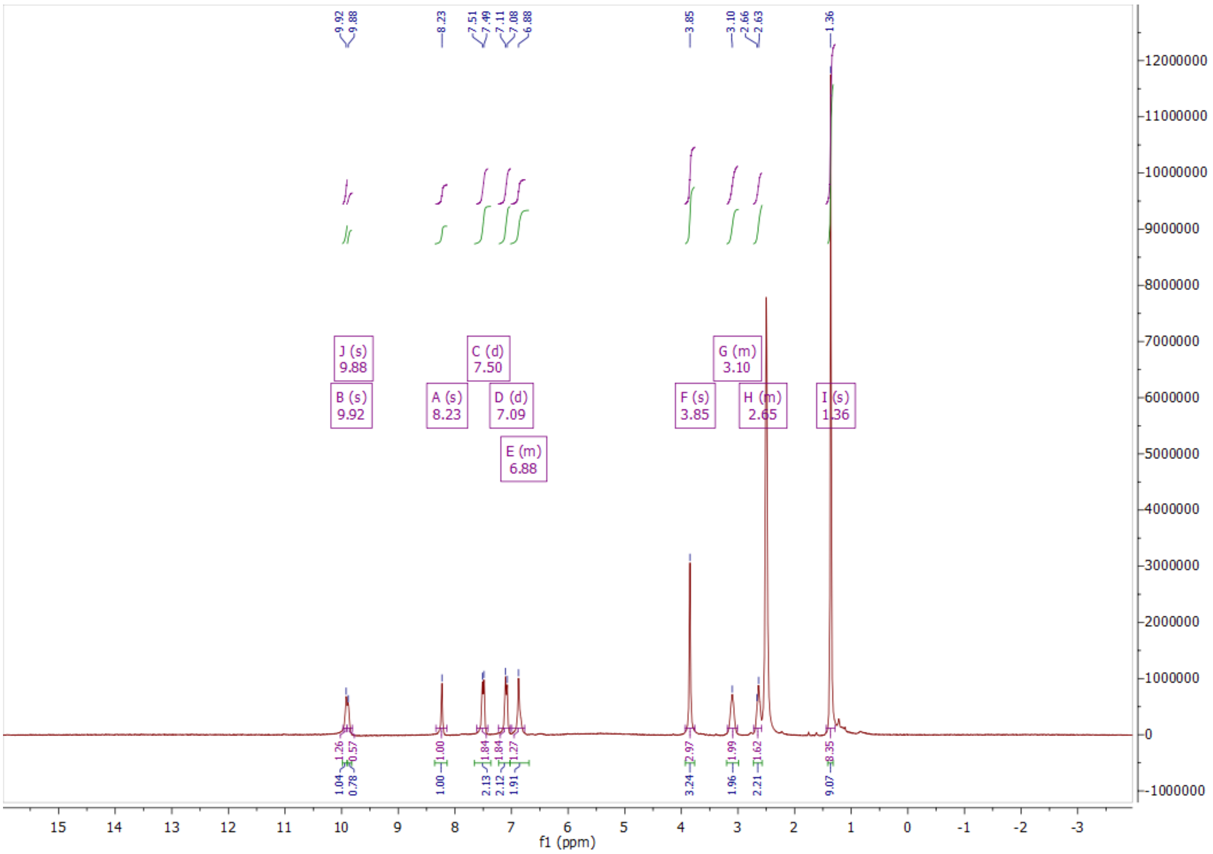
9.92
9.88
8.23
7.51
7.49
7.11
7.08
6.88
3.85
3.10
2.66
2.63
1.36
J (s) 9.88
B (s) 9.92
A (s) 8.23
C (d) 7.50
D (d) 7.09
E (m) 6.88
F (s) 3.85
G (m) 3.10
H (m) 2.65
I (s) 1.36
f1 (ppm)

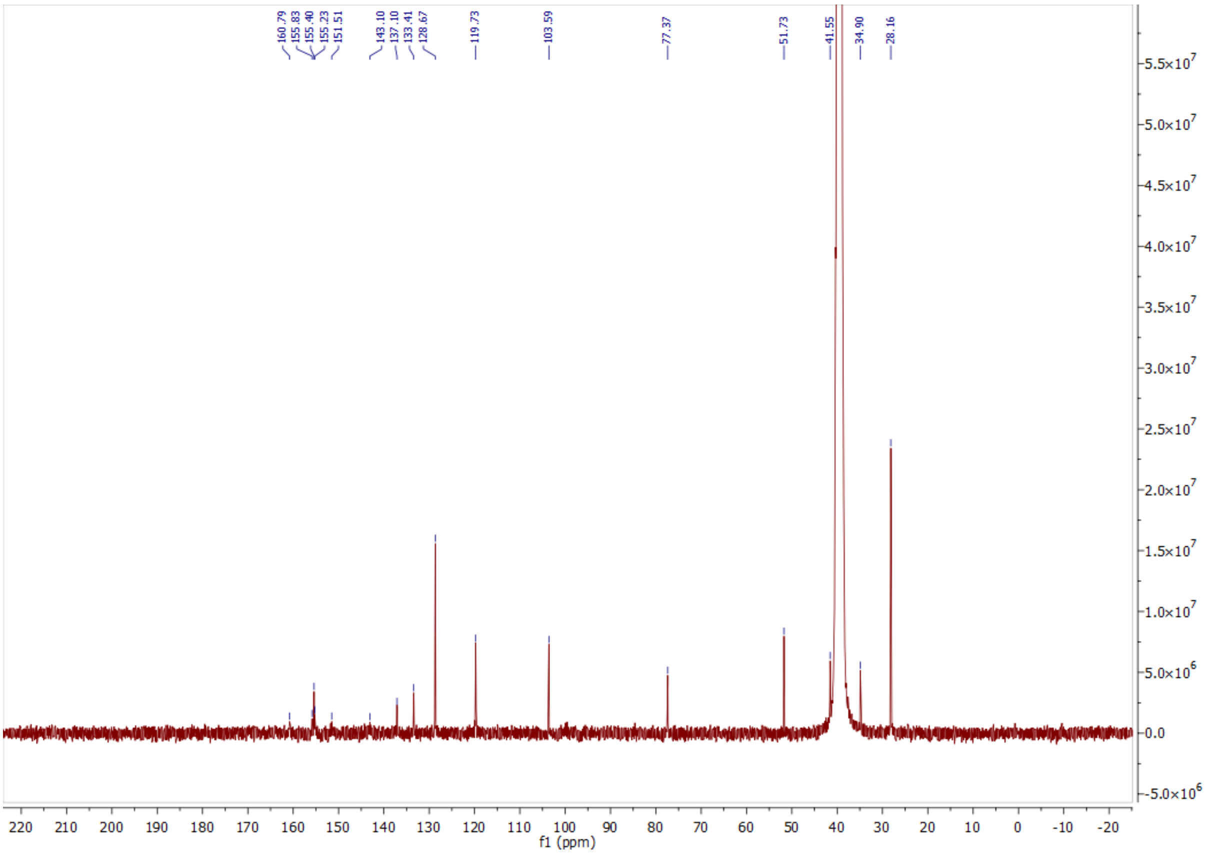
160.79
155.83
155.40
155.23
151.51
143.10
137.10
133.41
128.67
119.73
103.59
77.37
51.73
41.55
34.90
28.16
f1 (ppm)

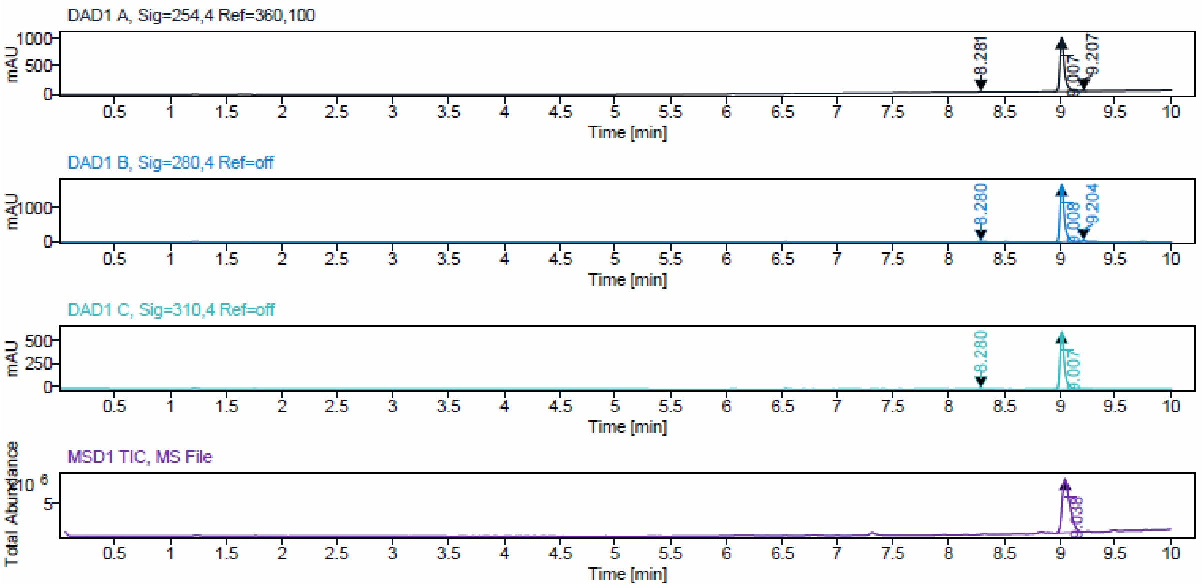


### Sample Purity

Signal Description DAD1 A, Sig=254,4 Ref=360,100

| Sample Name | Name | RT | Width | Area | Area% | Height |
|---|---|---|---|---|---|---|
| JA219_F17 | | 8.281 | 0.046 | 26.9441 | 0.98 | 12.2955 |
| JA219_F17 | | 9.007 | 0.039 | 2707.9041 | 98.28 | 987.3286 |
| JA219_F17 | | 9.207 | 0.045 | 20.5357 | 0.75 | 8.6404 |

Max Area% 98.277

UV Signal Purity>95% Pass

Signal Description DAD1 B, Sig=280,4 Ref=off

| Sample Name | Name | RT | Width | Area | Area% | Height |
|---|---|---|---|---|---|---|
| JA219_F17 | | 8.280 | 0.036 | 43.1173 | 0.88 | 16.4284 |
| JA219_F17 | | 9.008 | 0.039 | 4741.7402 | 97.06 | 1734.8192 |
| JA219_F17 | | 9.204 | 0.041 | 100.4228 | 2.06 | 36.5106 |

Max Area% 97.062

UV Signal Purity>95% Pass

Signal Description DAD1 C, Sig=310,4 Ref=off

| Sample Name | Name | RT | Width | Area | Area% | Height |
|---|---|---|---|---|---|---|
| JA219_F17 | | 8.280 | 0.035 | 16.3519 | 0.96 | 6.4223 |
| JA219_F17 | | 9.007 | 0.038 | 1693.8707 | 99.04 | 617.0402 |

Max Area% 99.044

UV Signal Purity>95% Pass

ESI, $^{1}$H, $^{13}$C NMR and HPLC data of compound **10b**.

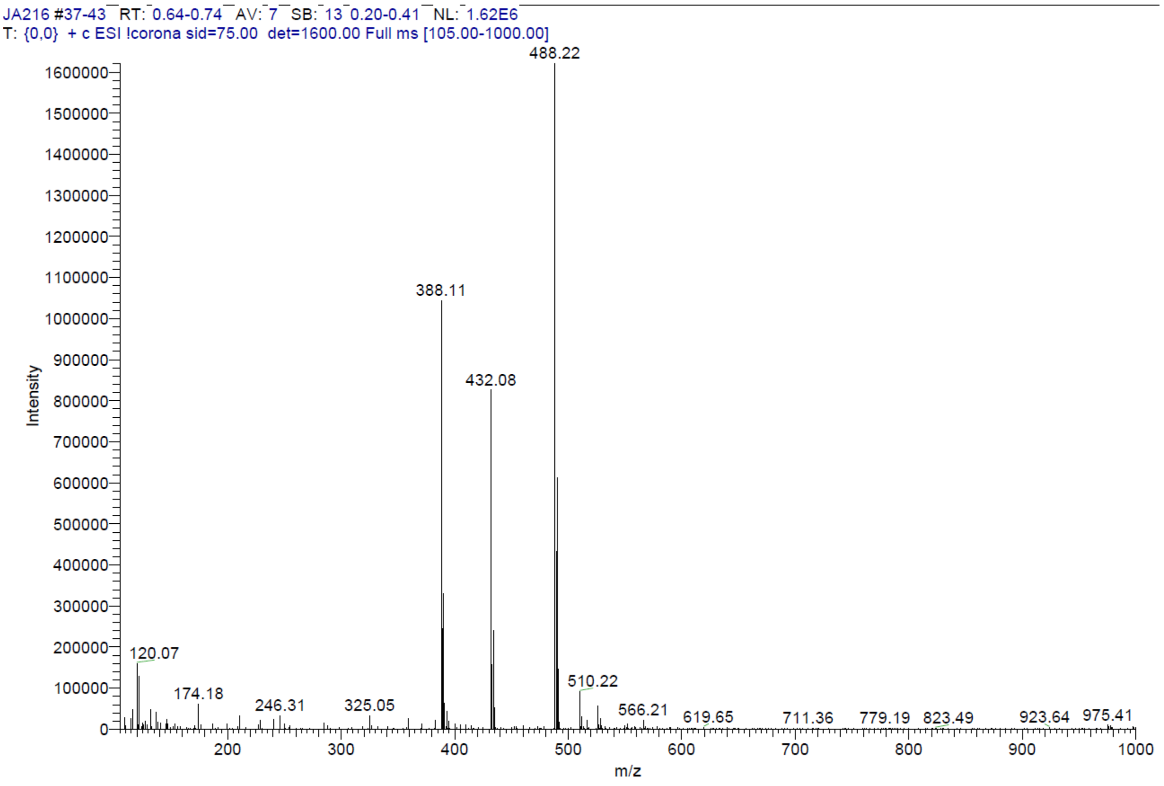


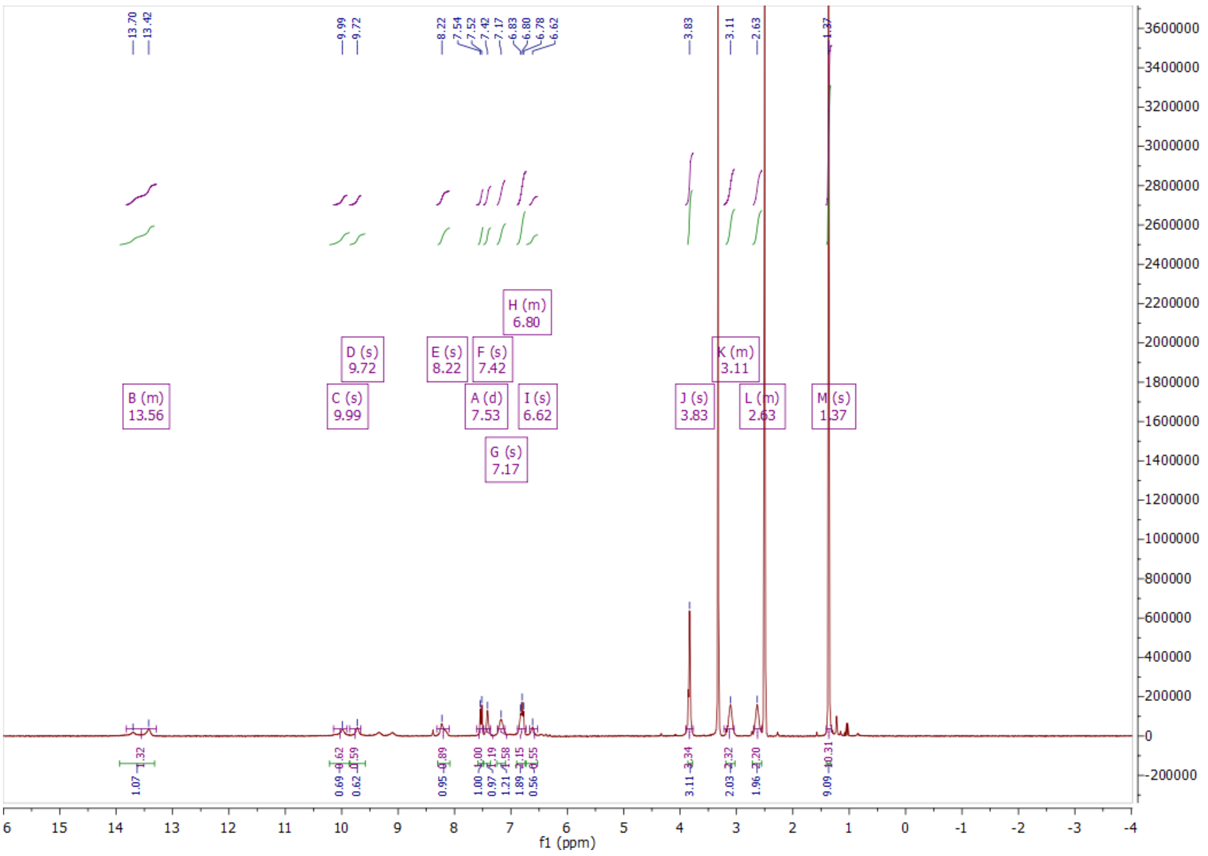

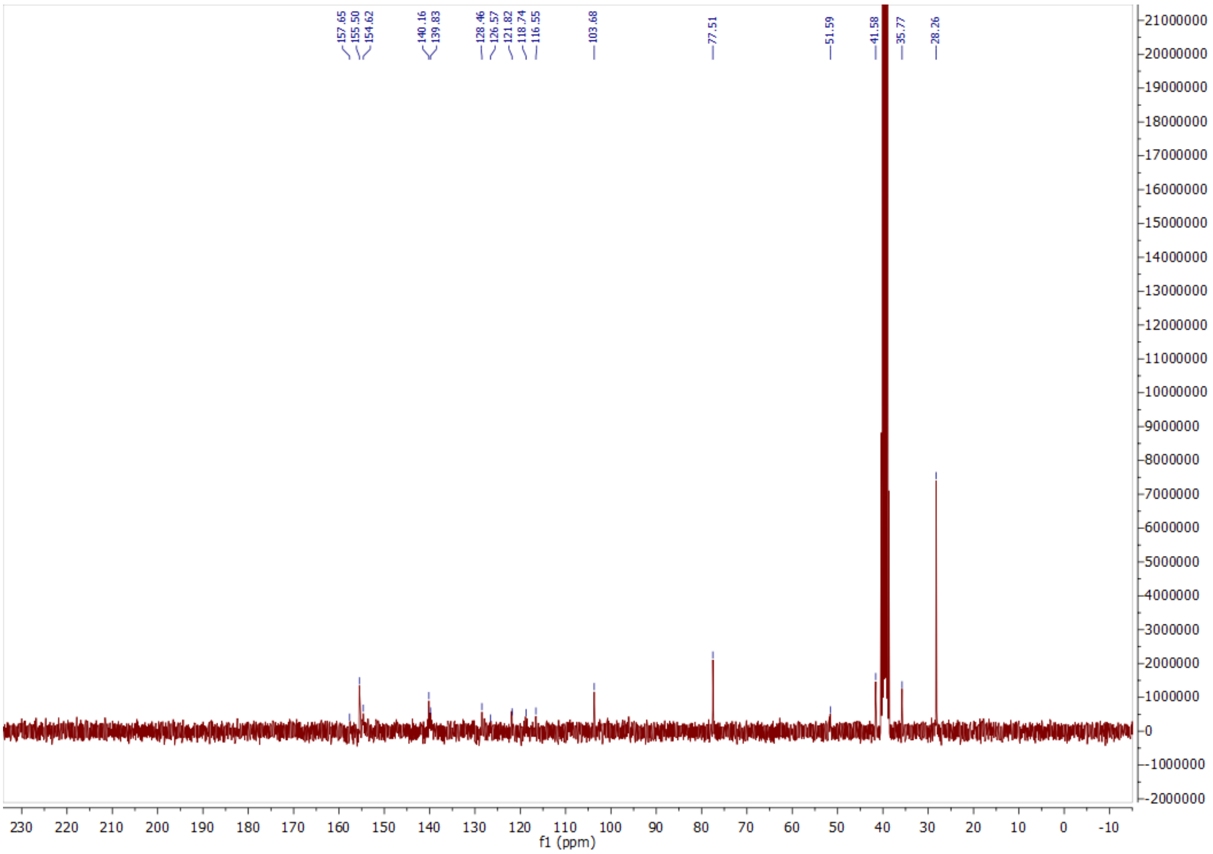
157.65
155.50
154.62
140.16
139.83
128.46
126.57
121.82
118.74
116.55
103.68
77.51
51.59
41.58
35.77
28.26
f1 (ppm)

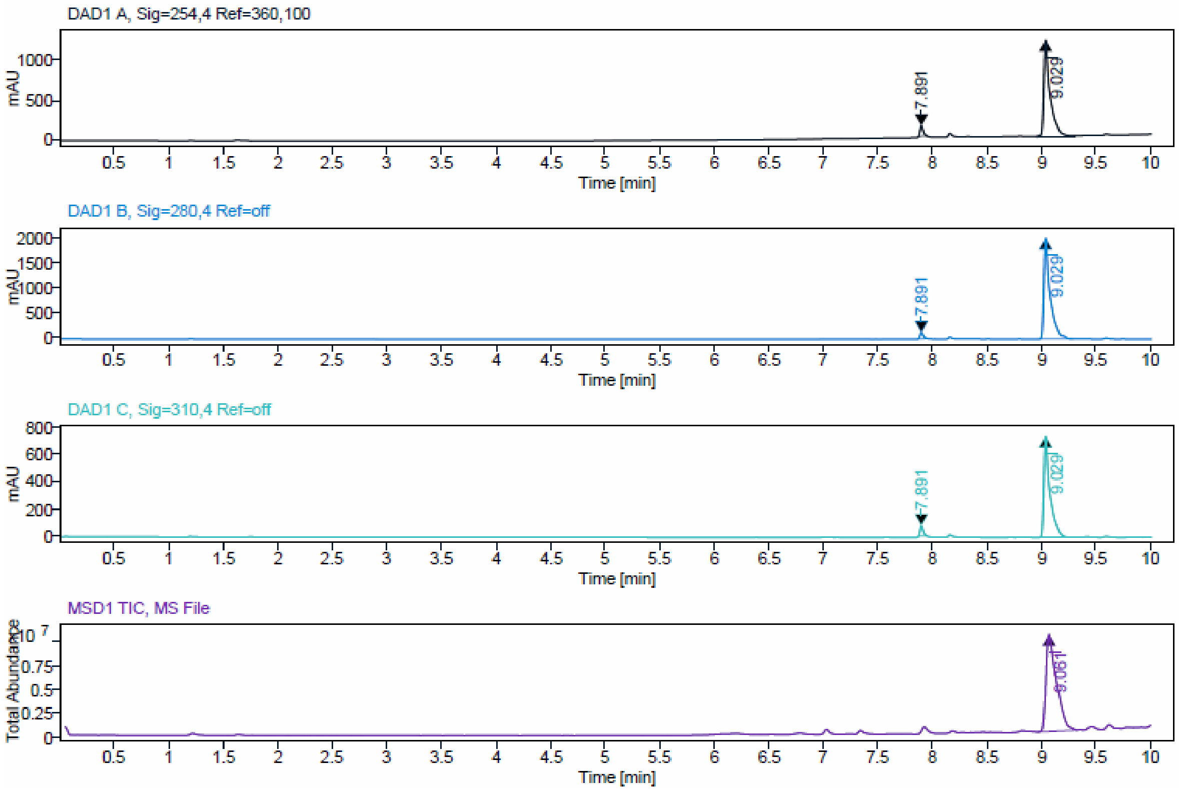
DAD1 A, Sig=254,4 Ref=360,100
DAD1 B, Sig=280,4 Ref=off
DAD1 C, Sig=310,4 Ref=off
MSD1 TIC, MS File
7.891
9.029
9.061
mAU
Total Abundance
Time [min]

**Sample Purity**

| | |
|---|---|
| Signal Description | DAD1 A, Sig=254,4 Ref=360,100 |

| Sample Name | Name | RT | Width | Area | Area% | Height |
|---|---|---|---|---|---|---|
| JA216_2_F5 | | 7.891 | 0.032 | 257.2323 | 5.00 | 128.6893 |
| JA216_2_F5 | | 9.029 | 0.048 | 4891.3472 | 95.00 | 1214.5306 |

| | |
|---|---|
| Max Area% | 95.004 |
| UV Signal Purity>95% | Pass |
| Signal Description | DAD1 B, Sig=280,4 Ref=off |

| Sample Name | Name | RT | Width | Area | Area% | Height |
|---|---|---|---|---|---|---|
| JA216_2_F5 | | 7.891 | 0.033 | 258.4486 | 3.17 | 122.4907 |
| JA216_2_F5 | | 9.029 | 0.048 | 7897.1875 | 96.83 | 2018.6160 |

| | |
|---|---|
| Max Area% | 96.831 |
| UV Signal Purity>95% | Pass |
| Signal Description | DAD1 C, Sig=310,4 Ref=off |

| Sample Name | Name | RT | Width | Area | Area% | Height |
|---|---|---|---|---|---|---|
| JA216_2_F5 | | 7.891 | 0.031 | 152.4315 | 4.97 | 76.1894 |
| JA216_2_F5 | | 9.029 | 0.047 | 2912.3542 | 95.03 | 752.7323 |

| | |
|---|---|
| Max Area% | 95.026 |
| UV Signal Purity>95% | Pass |

ESI, $^{1}H$, $^{13}C$ NMR and HPLC data of compound **10c**.

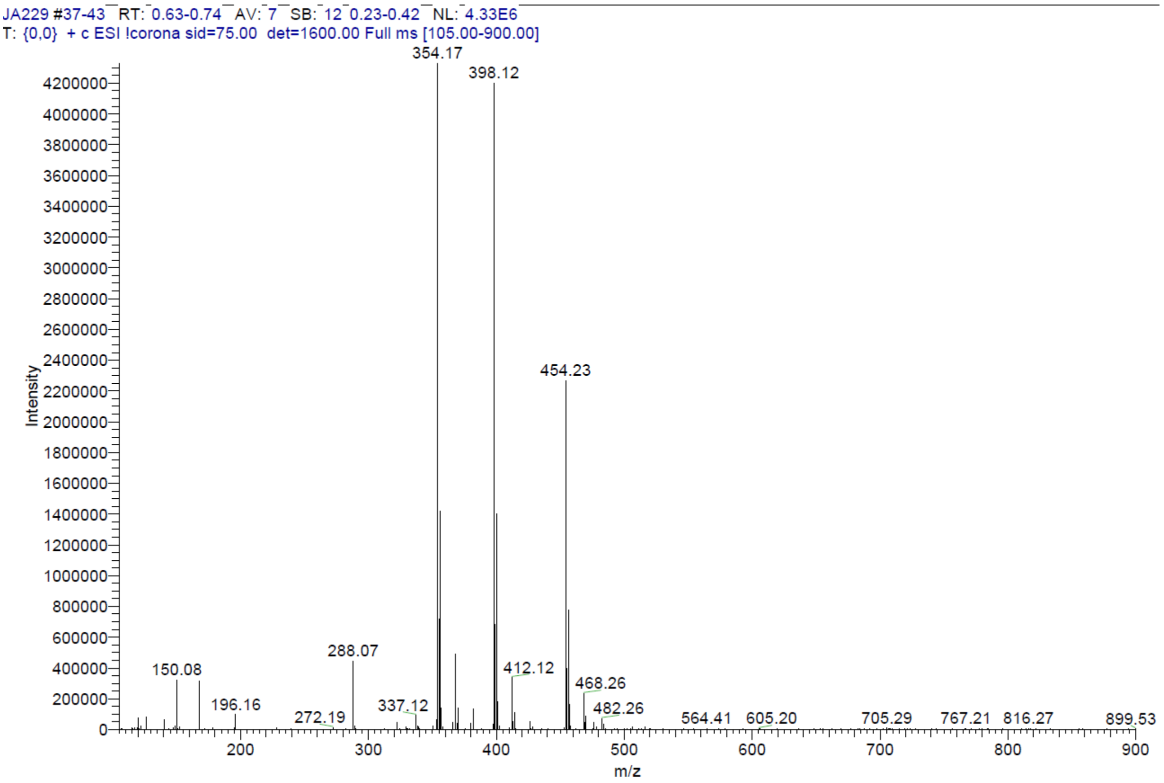

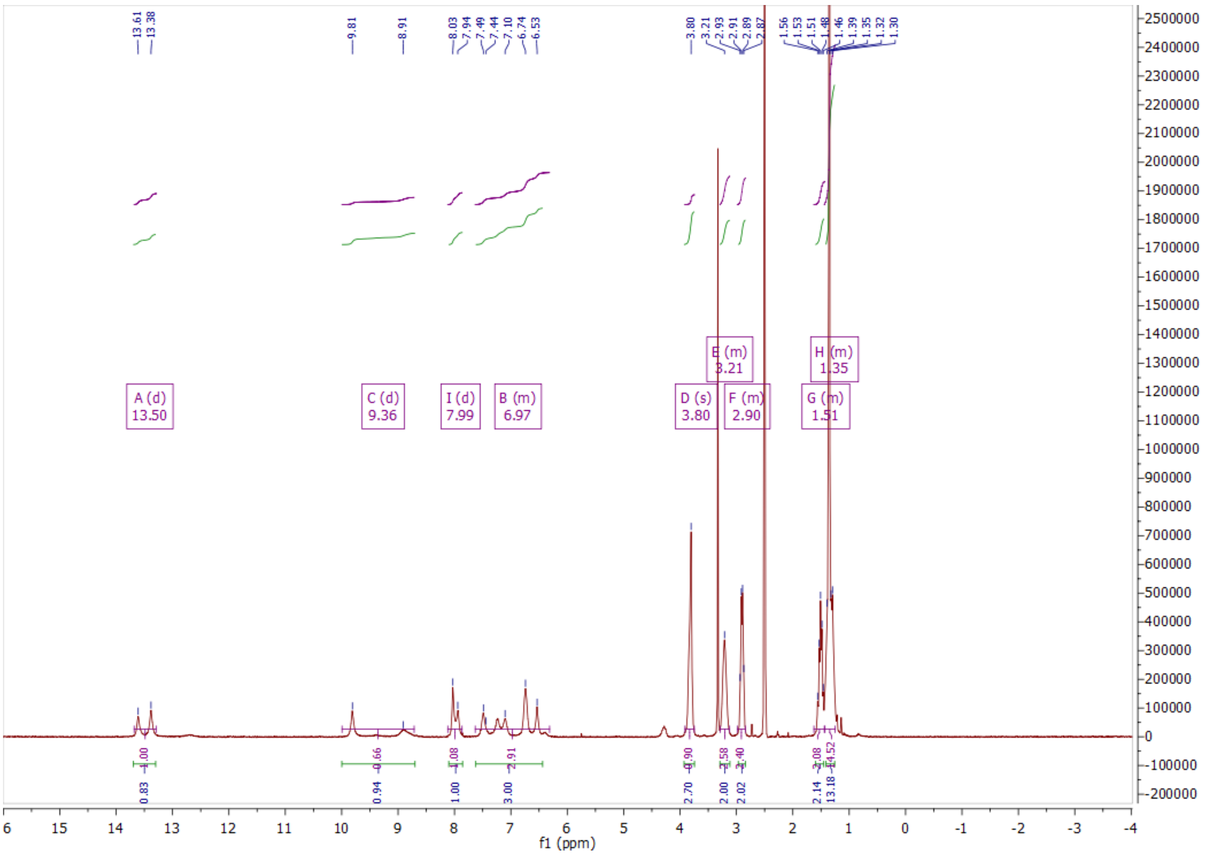

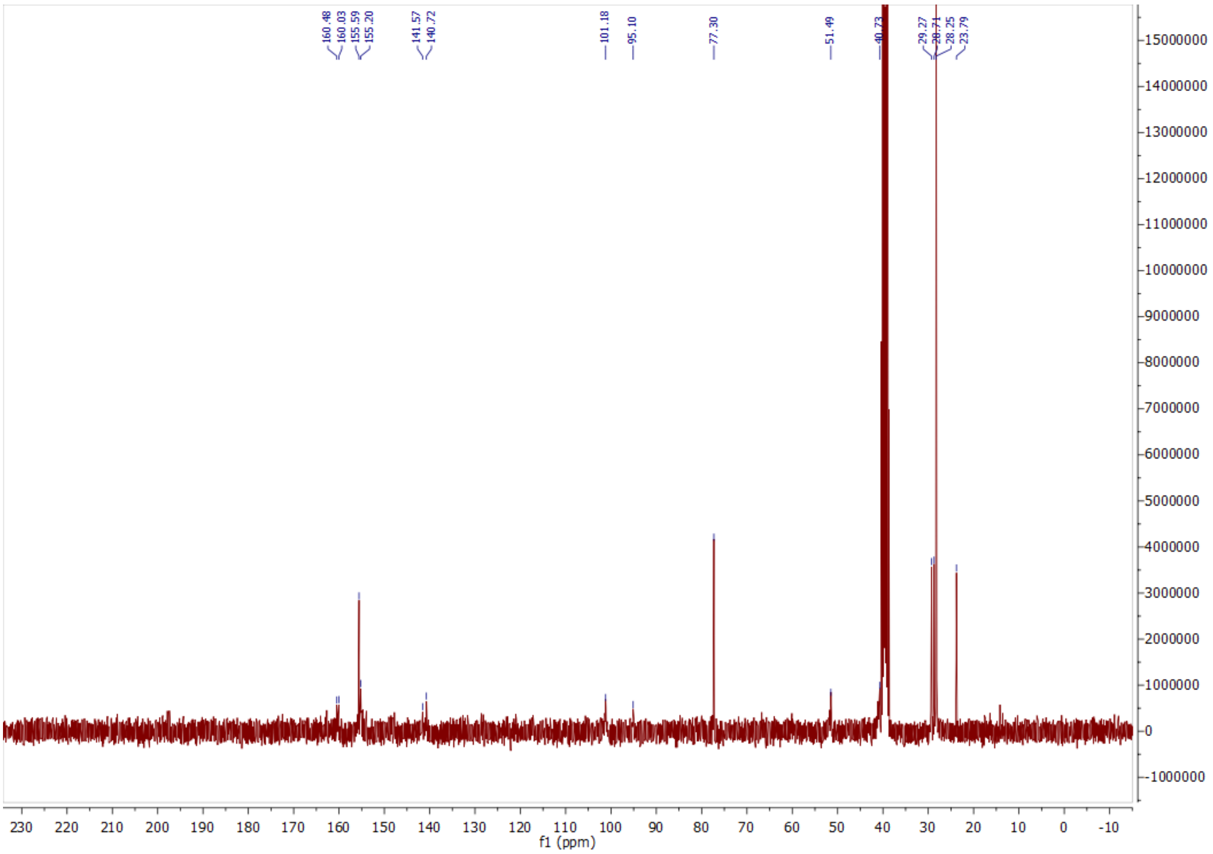

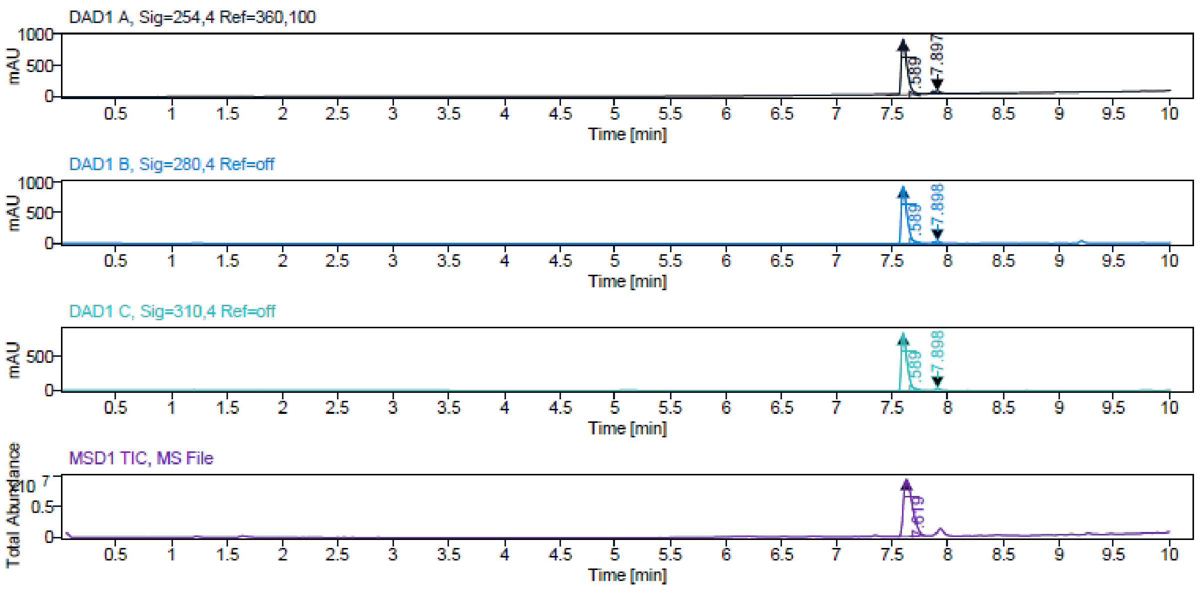


**Sample Purity**

Signal Description DAD1 A, Sig=254,4 Ref=360,100

| Sample Name | Name | RT | Width | Area | Area% | Height |
|---|---|---|---|---|---|---|
| JA229_F12 | | 7.589 | 0.054 | 3321.4458 | 95.04 | 905.7442 |
| JA229_F12 | | 7.897 | 0.071 | 173.3794 | 4.96 | 43.8780 |

Max Area% 95.039

UV Signal Purity>95% Pass

Signal Description DAD1 B, Sig=280,4 Ref=off

| Sample Name | Name | RT | Width | Area | Area% | Height |
|---|---|---|---|---|---|---|
| JA229_F12 | | 7.589 | 0.053 | 3382.2988 | 97.46 | 951.0269 |
| JA229_F12 | | 7.898 | 0.035 | 88.3266 | 2.54 | 35.7450 |

Max Area% 97.455

UV Signal Purity>95% Pass

Signal Description DAD1 C, Sig=310,4 Ref=off

| Sample Name | Name | RT | Width | Area | Area% | Height |
|---|---|---|---|---|---|---|
| JA229_F12 | | 7.589 | 0.053 | 3105.7893 | 98.19 | 871.3784 |
| JA229_F12 | | 7.898 | 0.032 | 57.3088 | 1.81 | 30.3489 |

Max Area% 98.188

UV Signal Purity>95% Pass

ESI, $^{1}$H, $^{13}$C NMR and HPLC data of compound **10d**.

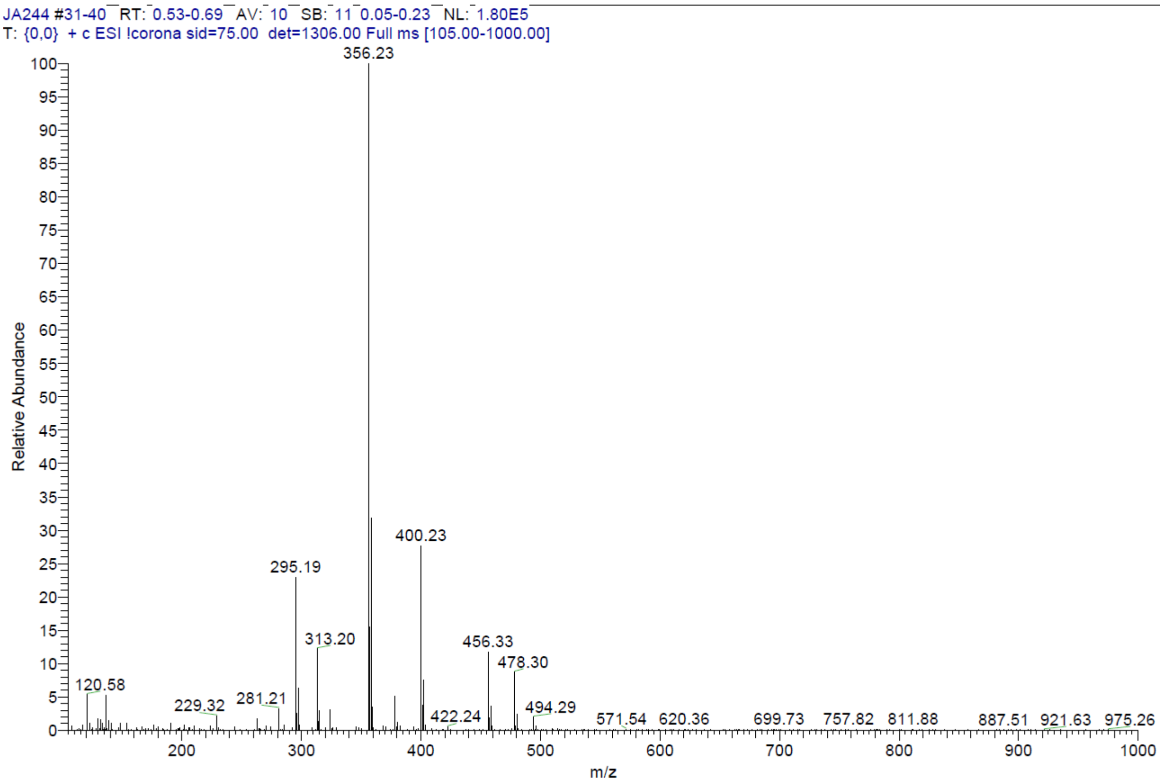


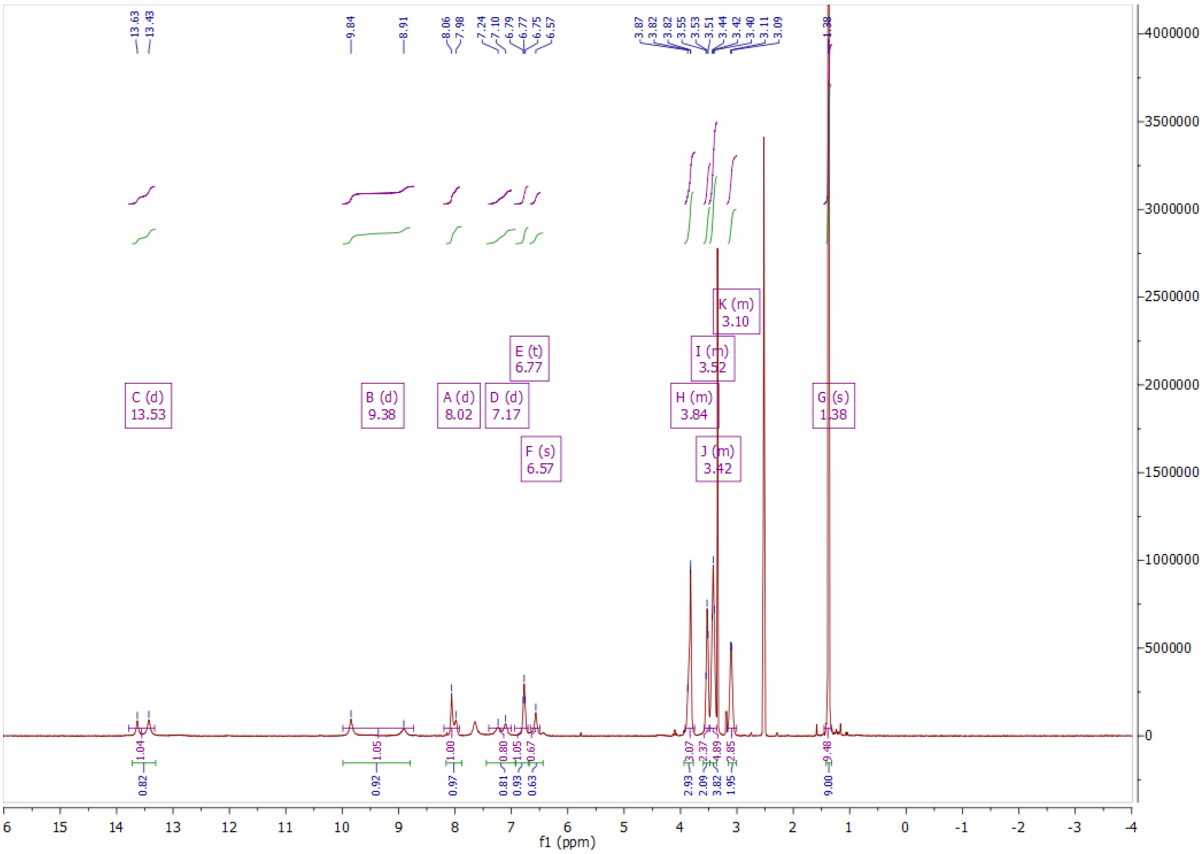

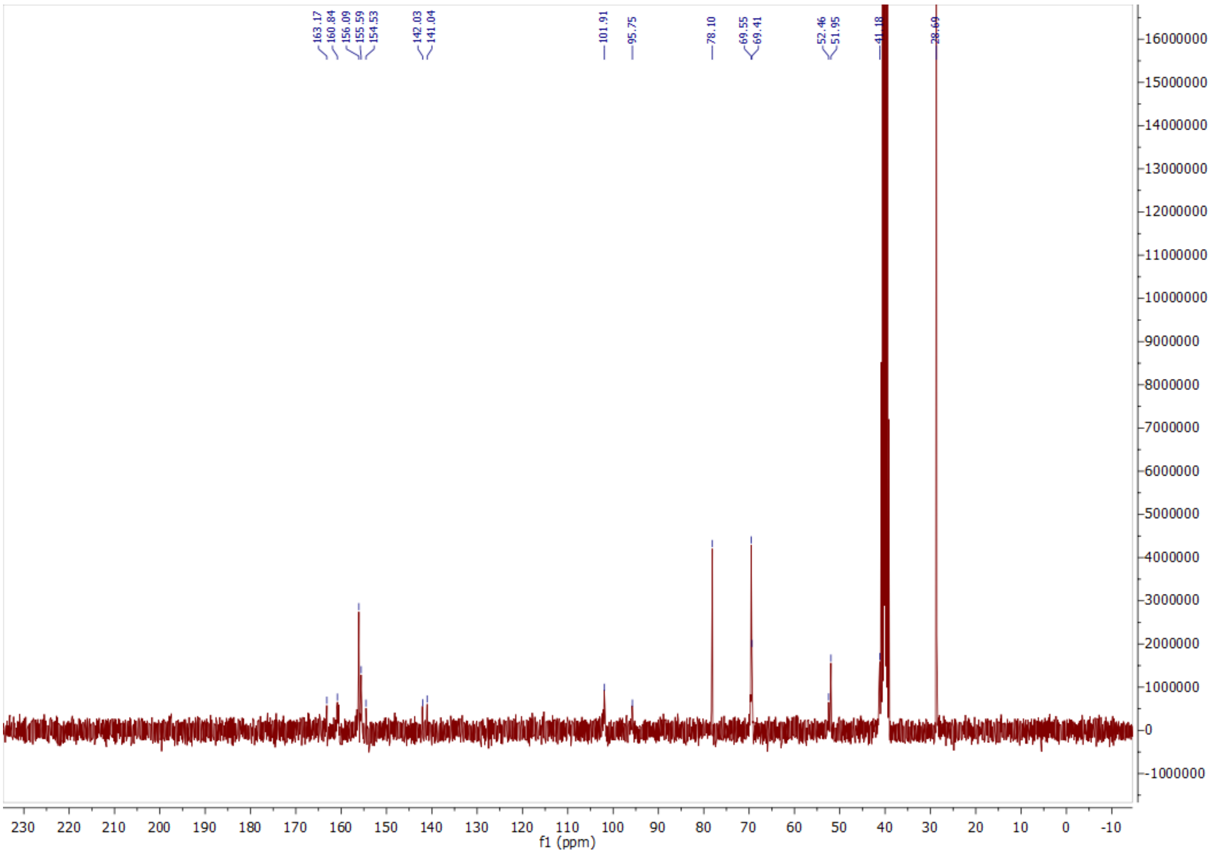
163.17
160.84
156.09
155.59
154.53
142.03
141.04
101.91
95.75
78.10
69.55
69.41
52.46
51.95
41.18
28.69
f1 (ppm)

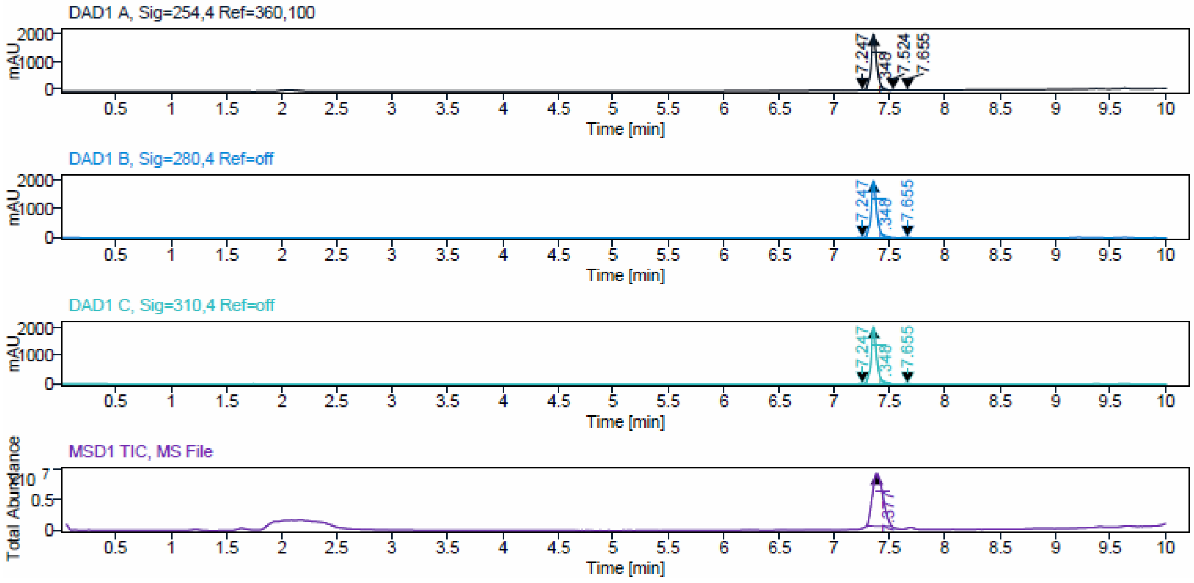
DAD1 A, Sig=254,4 Ref=360,100
DAD1 B, Sig=280,4 Ref=off
DAD1 C, Sig=310,4 Ref=off
MSD1 TIC, MS File
mAU
Total Abundance
Time [min]

**Sample Purity**

Signal Description DAD1 A, Sig=254,4 Ref=360,100

| Sample Name | Name | RT | Width | Area | Area% | Height |
|---|---|---|---|---|---|---|
| JA244_product | | 7.247 | 0.018 | 6.6651 | 0.10 | 6.2215 |
| JA244_product | | 7.348 | 0.049 | 6960.0249 | 99.43 | 1960.5637 |
| JA244_product | | 7.524 | 0.033 | 15.4476 | 0.22 | 7.8229 |
| JA244_product | | 7.655 | 0.031 | 17.9934 | 0.26 | 9.7974 |

Max Area% 99.427

UV Signal Purity>95% Pass

Signal Description DAD1 B, Sig=280,4 Ref=off

| Sample Name | Name | RT | Width | Area | Area% | Height |
|---|---|---|---|---|---|---|
| JA244_product | | 7.247 | 0.041 | 55.2211 | 0.75 | 18.9209 |
| JA244_product | | 7.348 | 0.047 | 7226.4063 | 98.36 | 2043.6461 |
| JA244_product | | 7.655 | 0.045 | 65.2849 | 0.89 | 14.6890 |

Max Area% 98.360

UV Signal Purity>95% Pass

Signal Description DAD1 C, Sig=310,4 Ref=off

| Sample Name | Name | RT | Width | Area | Area% | Height |
|---|---|---|---|---|---|---|
| JA244_product | | 7.247 | 0.040 | 47.7897 | 0.64 | 17.4729 |
| JA244_product | | 7.348 | 0.047 | 7309.6895 | 98.63 | 2069.8733 |
| JA244_product | | 7.655 | 0.042 | 53.9669 | 0.73 | 14.0873 |

Max Area% 98.627

UV Signal Purity>95% Pass

ESI, $^{1}H$, $^{13}C$ NMR and HPLC data of compound **10e**.

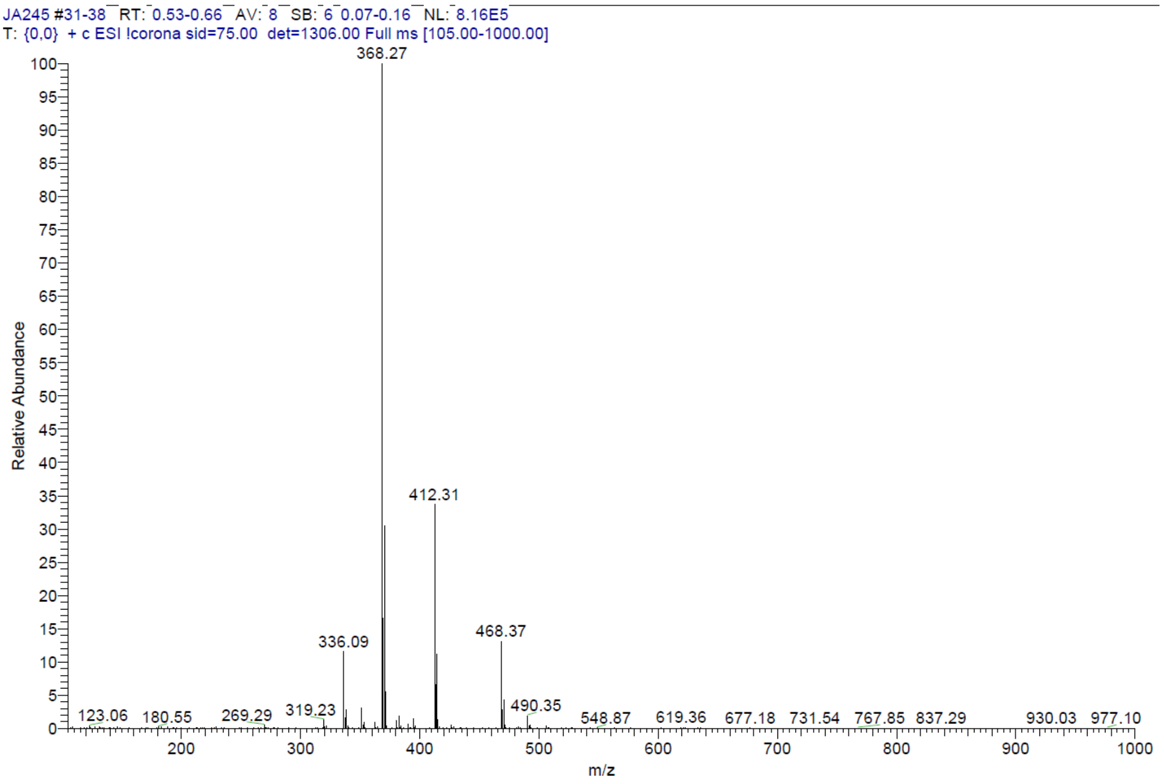


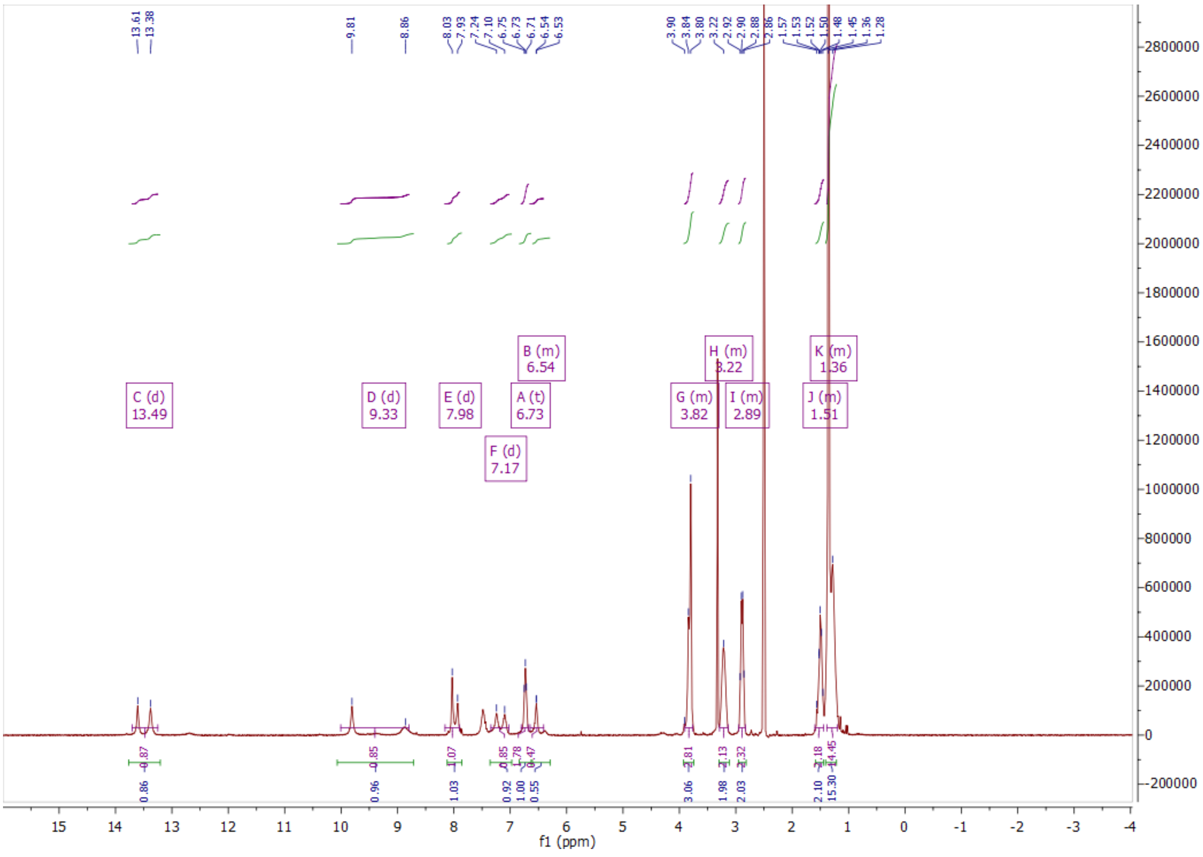

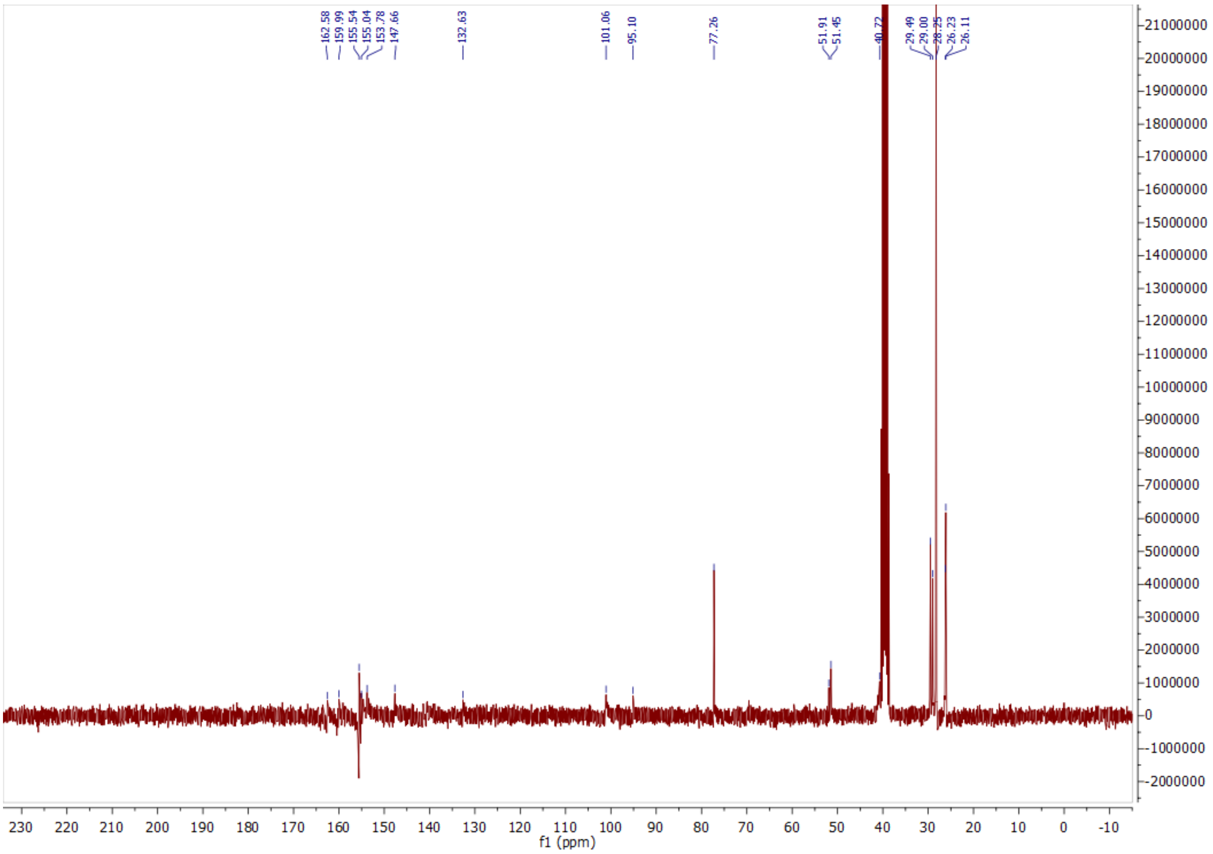

162.58
159.99
155.54
155.04
153.78
147.66
132.63
101.06
95.10
77.26
51.91
51.45
40.72
29.49
29.00
26.23
26.11
f1 (ppm)


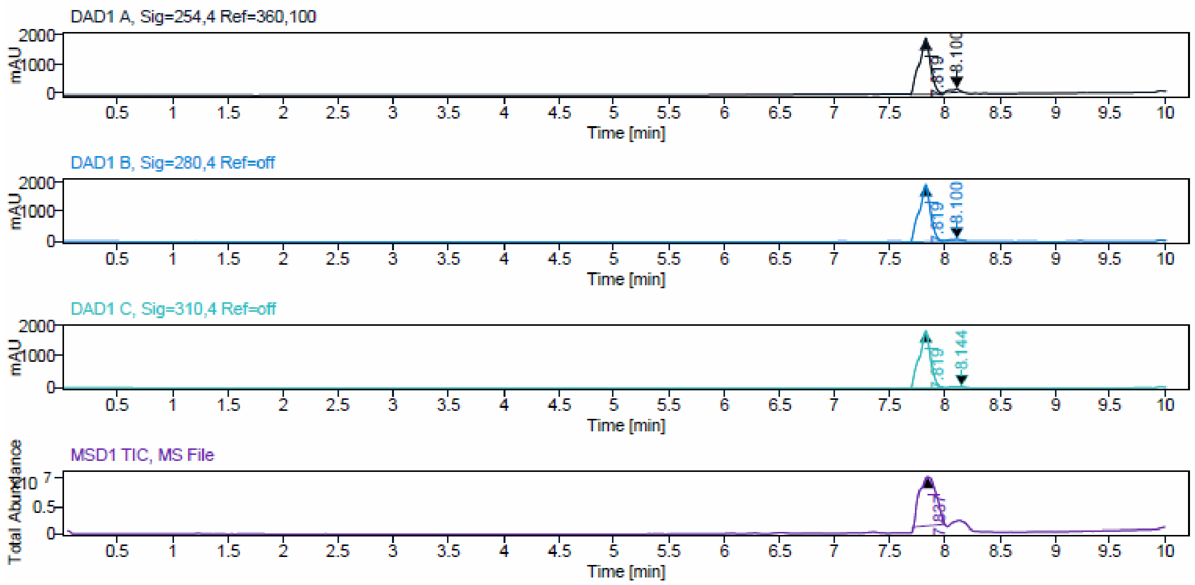

DAD1 A, Sig=254,4 Ref=360,100
DAD1 B, Sig=280,4 Ref=off
DAD1 C, Sig=310,4 Ref=off
MSD1 TIC, MS File
mAU
Time [min]
Total Abundance

**Sample Purity**

Signal Description DAD1 A, Sig=254,4 Ref=360,100

| Sample Name | Name | RT | Width | Area | Area% | Height |
|---|---|---|---|---|---|---|
| JA242_F10 | | 7.819 | 0.115 | 13908.8906 | 95.12 | 1891.8716 |
| JA242_F10 | | 8.100 | 0.102 | 713.2922 | 4.88 | 125.9100 |

Max Area% 95.122

UV Signal Purity>95% Pass

Signal Description DAD1 B, Sig=280,4 Ref=off

| Sample Name | Name | RT | Width | Area | Area% | Height |
|---|---|---|---|---|---|---|
| JA242_F10 | | 7.819 | 0.110 | 14103.9756 | 96.49 | 1974.7263 |
| JA242_F10 | | 8.100 | 0.139 | 513.7810 | 3.51 | 70.9866 |

Max Area% 96.485

UV Signal Purity>95% Pass

Signal Description DAD1 C, Sig=310,4 Ref=off

| Sample Name | Name | RT | Width | Area | Area% | Height |
|---|---|---|---|---|---|---|
| JA242_F10 | | 7.819 | 0.110 | 13242.5020 | 98.30 | 1838.7095 |
| JA242_F10 | | 8.144 | 0.137 | 228.7905 | 1.70 | 33.6280 |

Max Area% 98.302

UV Signal Purity>95% Pass

ESI data of compound **11a**.

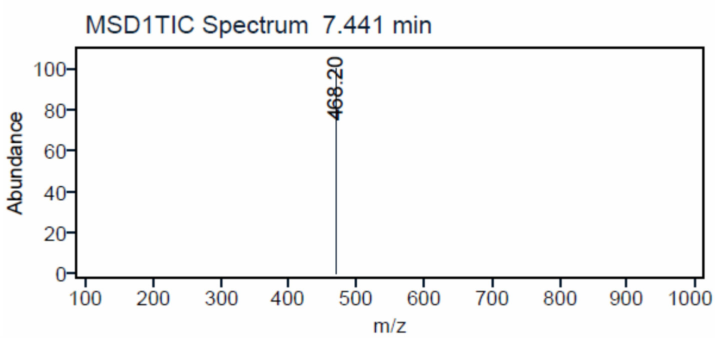

ESI, $^{1}$H, $^{13}$C NMR and HPLC data of compound **12a**.

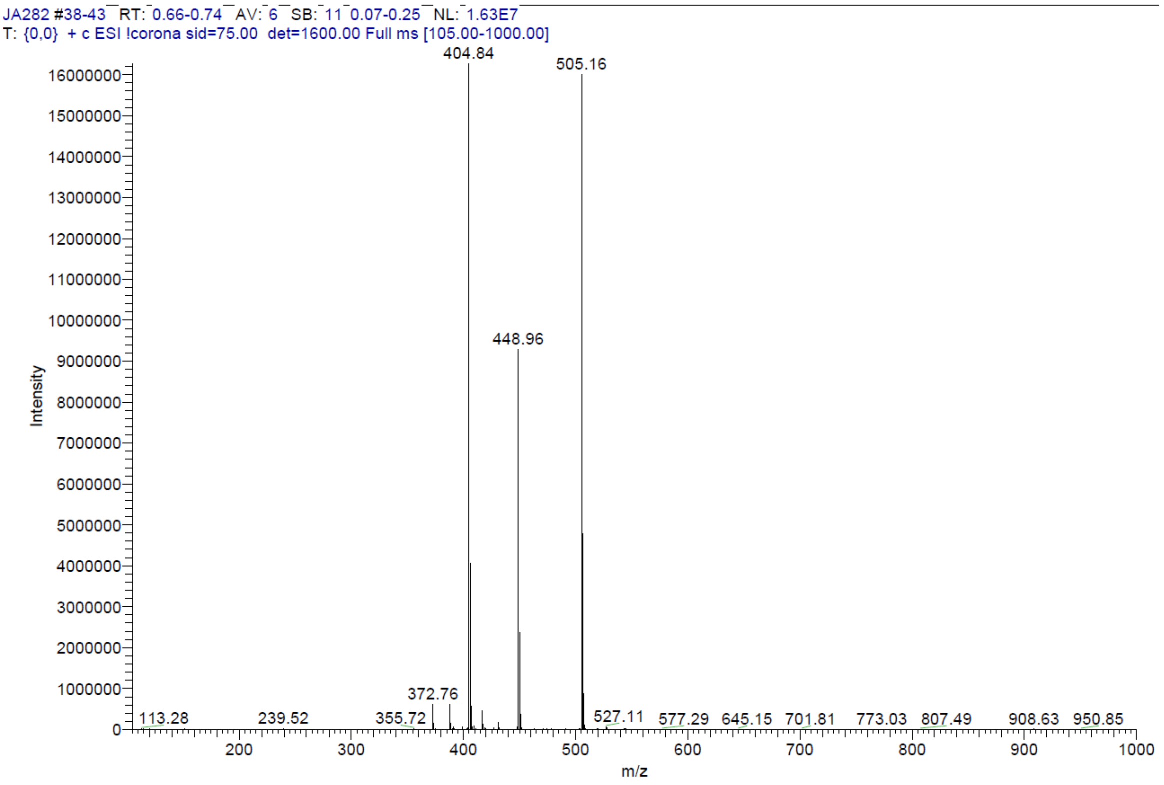


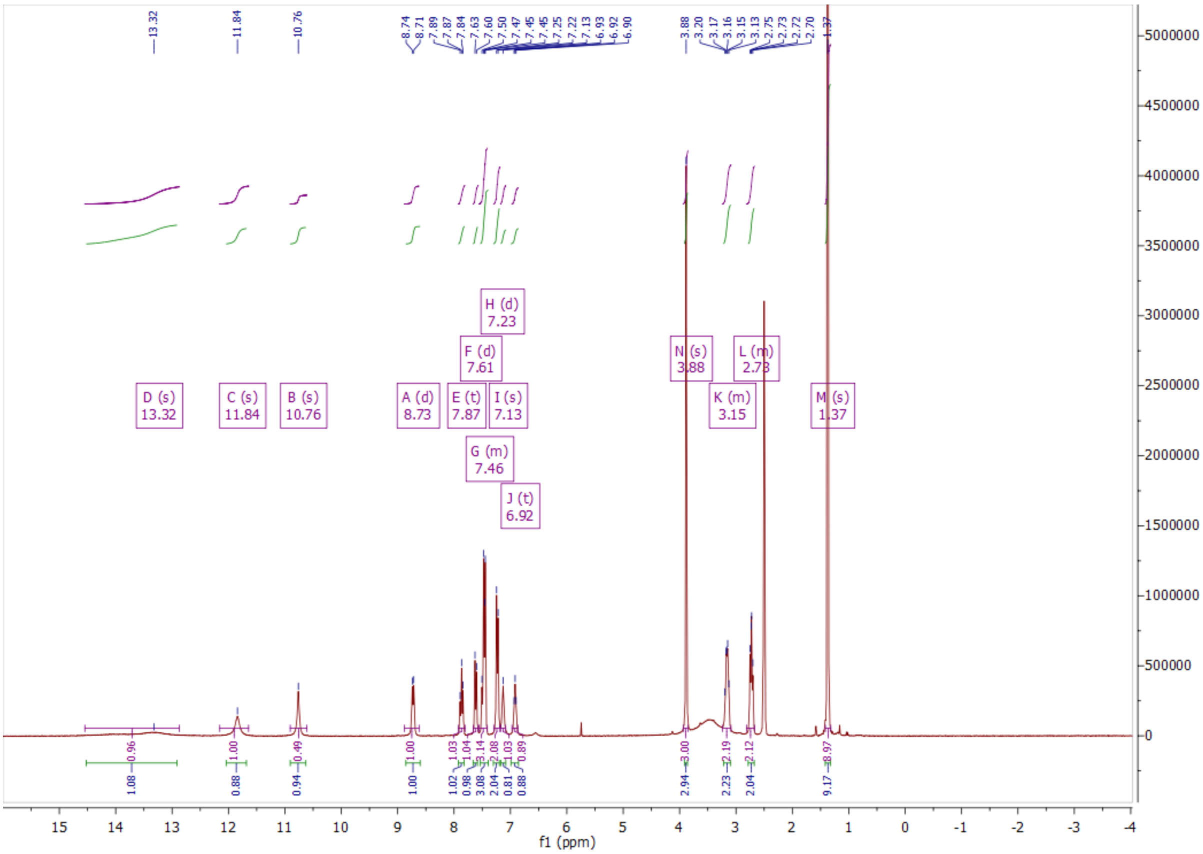

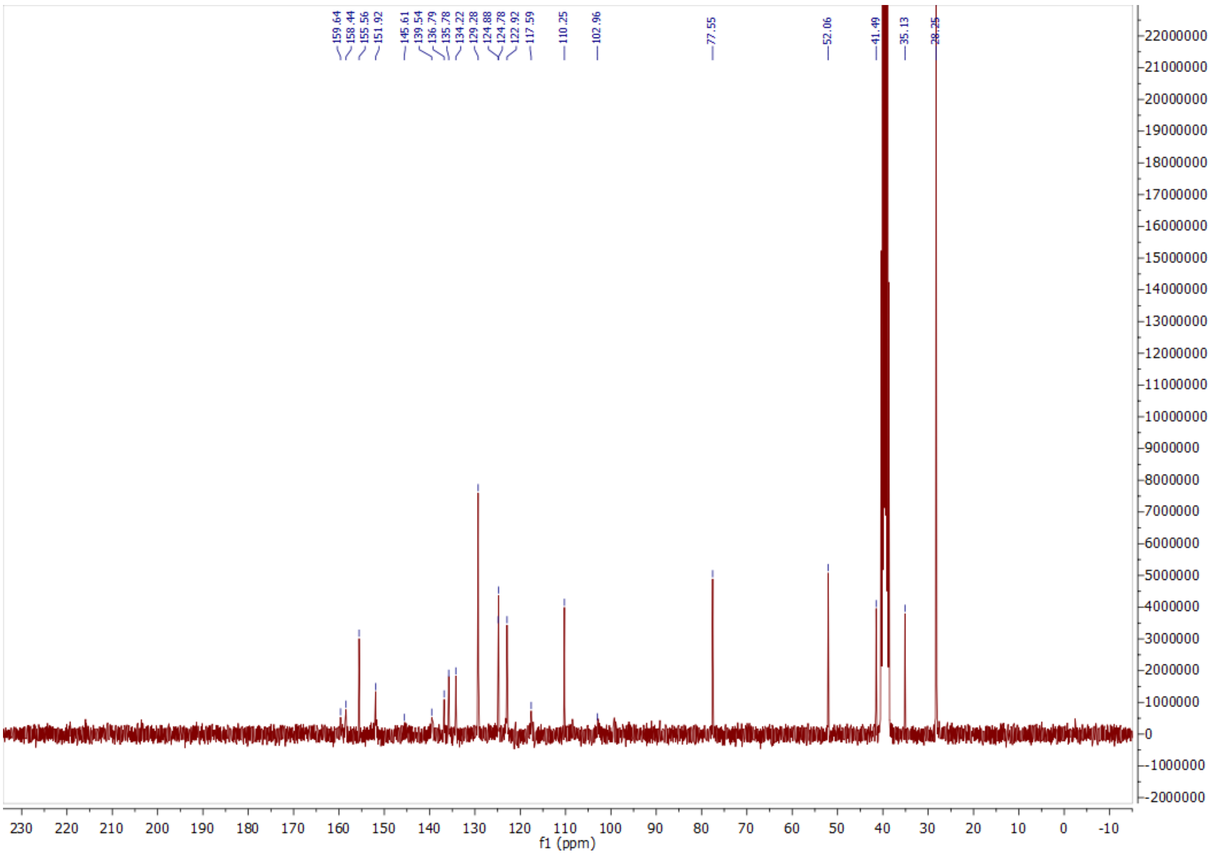
159.64
158.44
155.56
151.92
145.61
139.54
136.79
135.78
134.22
129.28
124.88
124.78
122.92
117.59
110.25
102.96
77.55
52.06
41.49
35.13
28.25
f1 (ppm)

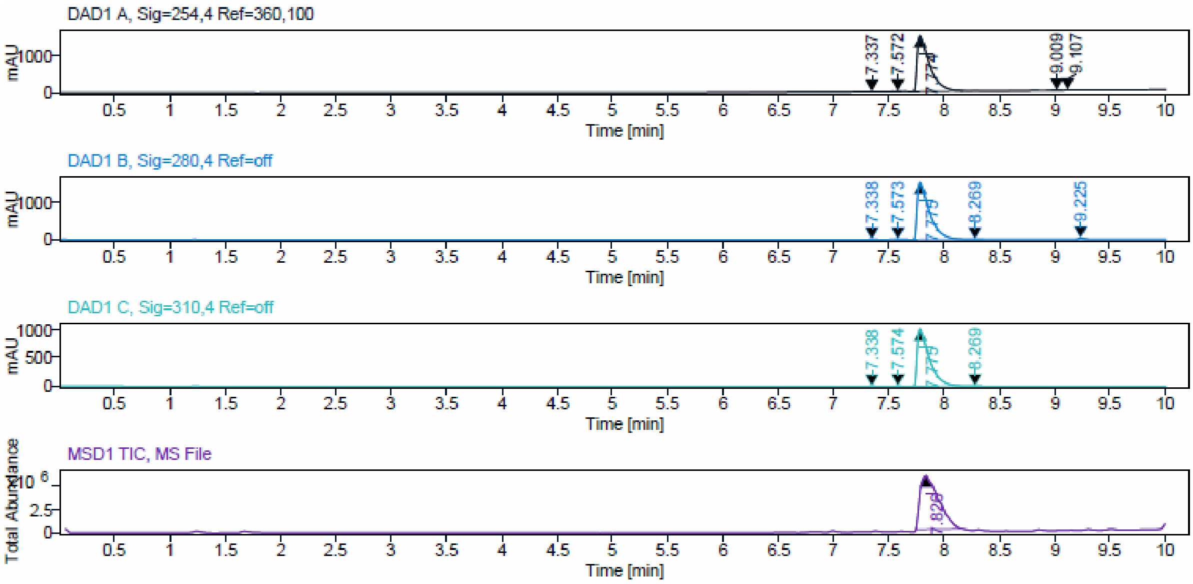
DAD1 A, Sig=254,4 Ref=360,100
DAD1 B, Sig=280,4 Ref=off
DAD1 C, Sig=310,4 Ref=off
MSD1 TIC, MS File
Time [min]
mAU
Total Abundance

**Sample Purity**

Signal Description DAD1 A, Sig=254,4 Ref=360,100

| Sample Name | Name | RT | Width | Area | Area% | Height |
|---|---|---|---|---|---|---|
| JA282_F15 | | 7.337 | 0.054 | 18.5845 | 0.16 | 6.4313 |
| JA282_F15 | | 7.572 | 0.053 | 43.8318 | 0.38 | 12.9010 |
| JA282_F15 | | 7.774 | 0.113 | 11492.2168 | 98.66 | 1489.1780 |
| JA282_F15 | | 9.009 | 0.069 | 46.4684 | 0.40 | 12.3748 |
| JA282_F15 | | 9.107 | 0.074 | 47.3371 | 0.41 | 13.1516 |

Max Area% 98.659

UV Signal Purity>95% Pass

Signal Description DAD1 B, Sig=280,4 Ref=off

| Sample Name | Name | RT | Width | Area | Area% | Height |
|---|---|---|---|---|---|---|
| JA282_F15 | | 7.338 | 0.045 | 23.2447 | 0.20 | 6.7086 |
| JA282_F15 | | 7.573 | 0.037 | 22.7132 | 0.19 | 8.0980 |
| JA282_F15 | | 7.775 | 0.111 | 11714.4355 | 98.69 | 1518.0446 |
| JA282_F15 | | 8.269 | 0.046 | 18.0723 | 0.15 | 5.9514 |
| JA282_F15 | | 9.225 | 0.047 | 91.2506 | 0.77 | 32.6501 |

Max Area% 98.692

UV Signal Purity>95% Pass

Signal Description DAD1 C, Sig=310,4 Ref=off

| Sample Name | Name | RT | Width | Area | Area% | Height |
|---|---|---|---|---|---|---|
| JA282_F15 | | 7.338 | 0.045 | 16.0702 | 0.20 | 4.6109 |
| JA282_F15 | | 7.574 | 0.039 | 17.9199 | 0.22 | 5.7650 |
| JA282_F15 | | 7.775 | 0.111 | 8134.1978 | 99.42 | 1057.8473 |
| JA282_F15 | | 8.269 | 0.047 | 13.2664 | 0.16 | 4.1943 |

Max Area% 99.422

UV Signal Purity>95% Pass

ESI, $^{1}$H, $^{13}$C NMR and HPLC data of compound **12b**.

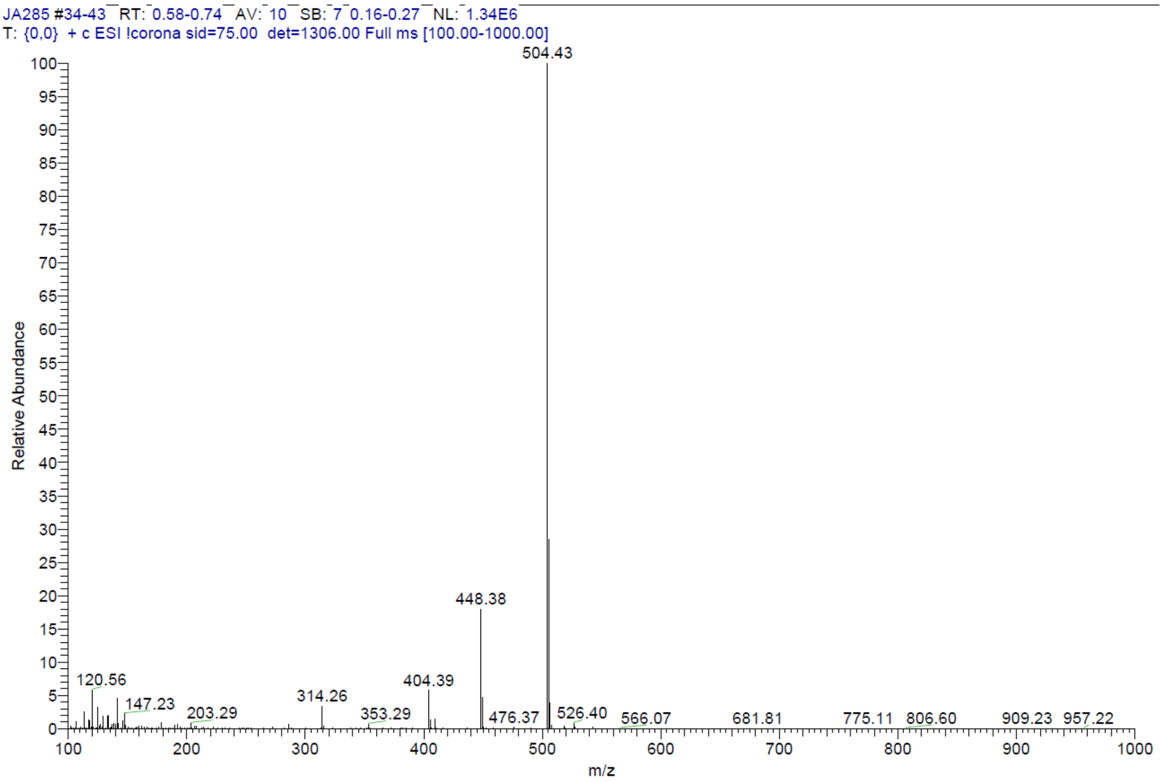


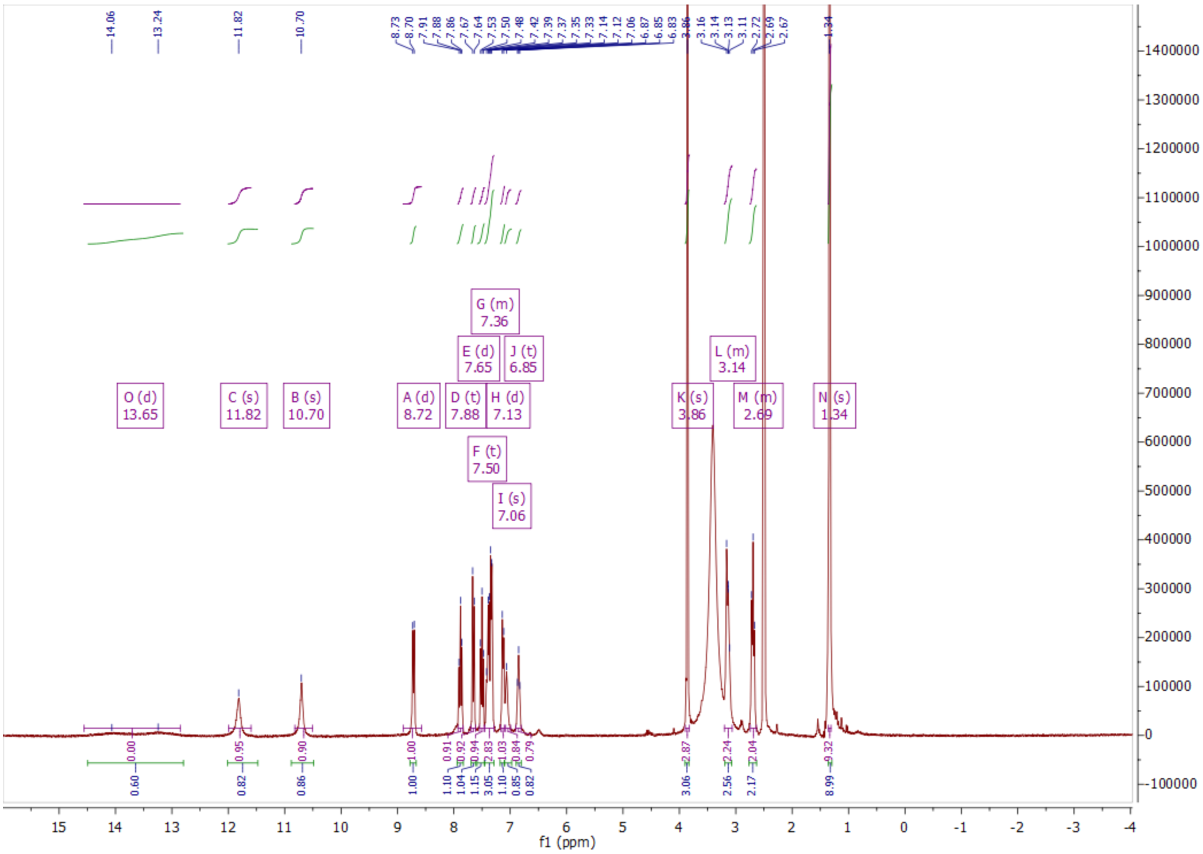

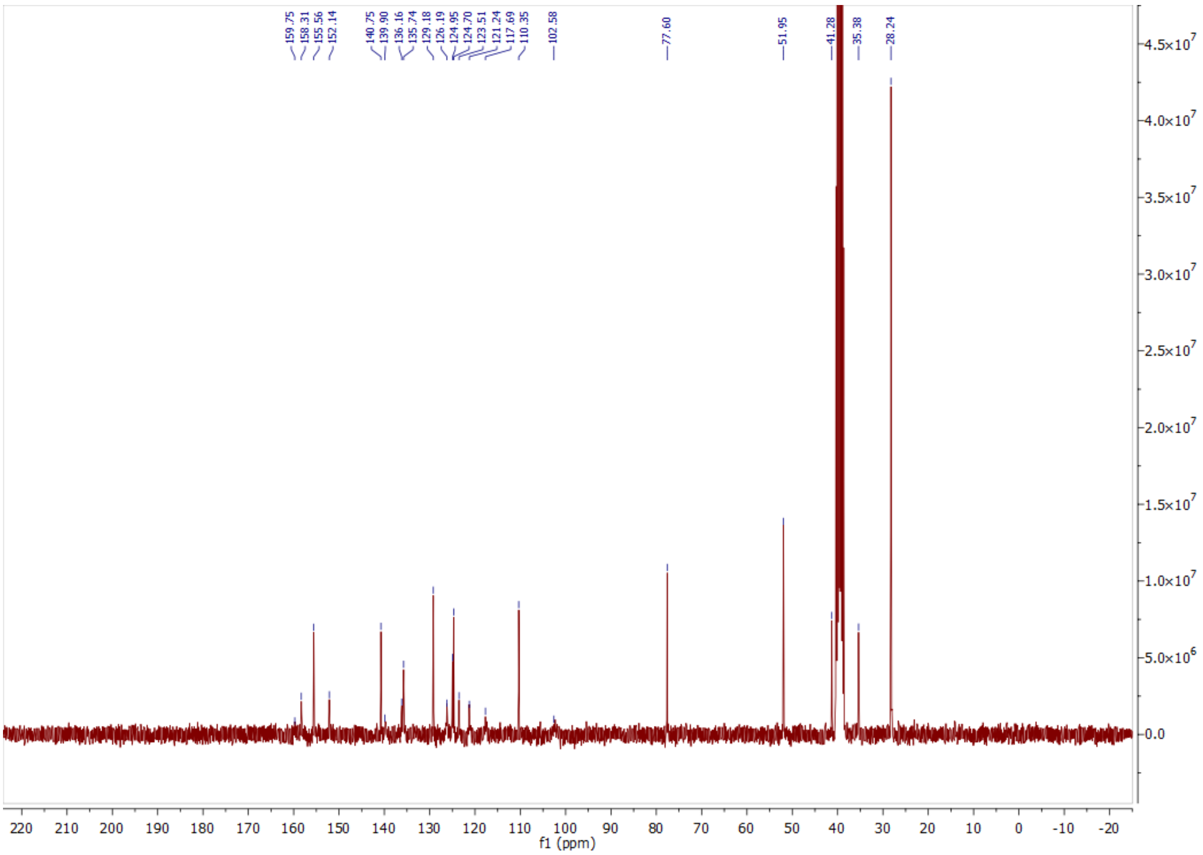
159.75
158.31
155.56
152.14
140.75
139.90
136.16
135.74
129.18
126.19
124.95
124.70
123.51
121.24
117.69
110.35
102.58
77.60
51.95
41.28
35.38
28.24
f1 (ppm)

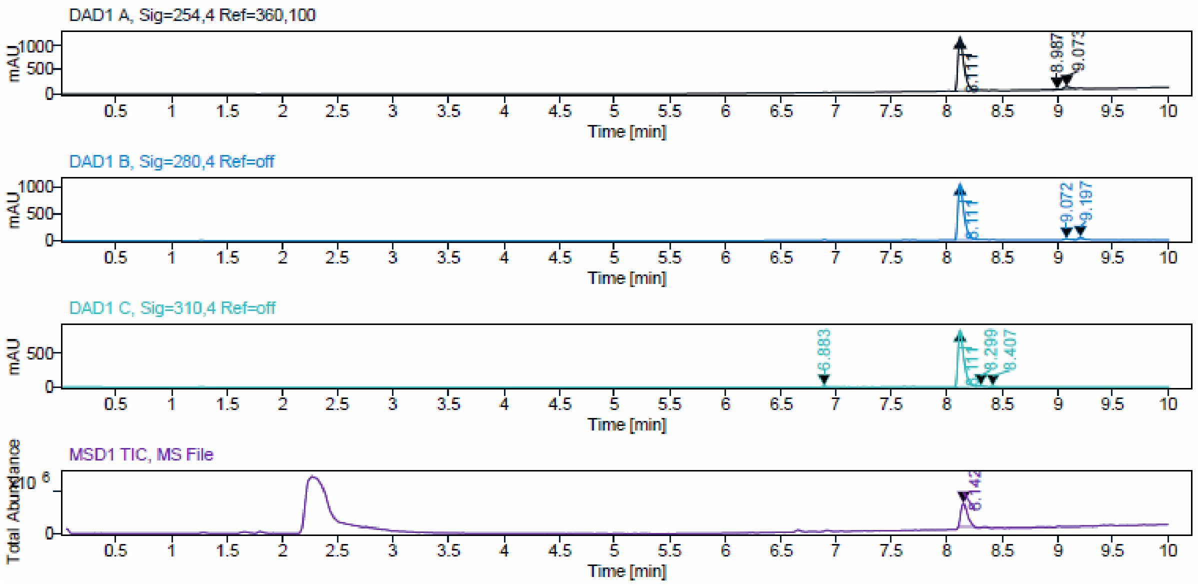
DAD1 A, Sig=254,4 Ref=360,100
mAU
Time [min]
DAD1 B, Sig=280,4 Ref=off
DAD1 C, Sig=310,4 Ref=off
MSD1 TIC, MS File
Total Abundance

**Sample Purity**

Signal Description DAD1 A, Sig=254,4 Ref=360,100

| Sample Name | Name | RT | Width | Area | Area% | Height |
|---|---|---|---|---|---|---|
| JA285_F7 | | 8.111 | 0.057 | 4212.3657 | 95.26 | 1127.8860 |
| JA285_F7 | | 8.987 | 0.046 | 22.7012 | 0.51 | 8.2231 |
| JA285_F7 | | 9.073 | 0.065 | 186.7537 | 4.22 | 48.2658 |

Max Area% 95.263

UV Signal Purity>95% Pass

Signal Description DAD1 B, Sig=280,4 Ref=off

| Sample Name | Name | RT | Width | Area | Area% | Height |
|---|---|---|---|---|---|---|
| JA285_F7 | | 8.111 | 0.057 | 3988.7537 | 95.70 | 1060.9926 |
| JA285_F7 | | 9.072 | 0.064 | 54.3462 | 1.30 | 14.3521 |
| JA285_F7 | | 9.197 | 0.050 | 124.8755 | 3.00 | 41.3984 |

Max Area% 95.700

UV Signal Purity>95% Pass

Signal Description DAD1 C, Sig=310,4 Ref=off

| Sample Name | Name | RT | Width | Area | Area% | Height |
|---|---|---|---|---|---|---|
| JA285_F7 | | 6.883 | 0.033 | 28.6834 | 0.89 | 13.9921 |
| JA285_F7 | | 8.111 | 0.057 | 3146.4600 | 97.92 | 836.7382 |
| JA285_F7 | | 8.299 | 0.046 | 17.4013 | 0.54 | 5.8589 |
| JA285_F7 | | 8.407 | 0.044 | 20.7846 | 0.65 | 7.1484 |

Max Area% 97.919

UV Signal Purity>95% Pass

ESI, [1]H, [13]C NMR and HPLC data of compound **12c**.

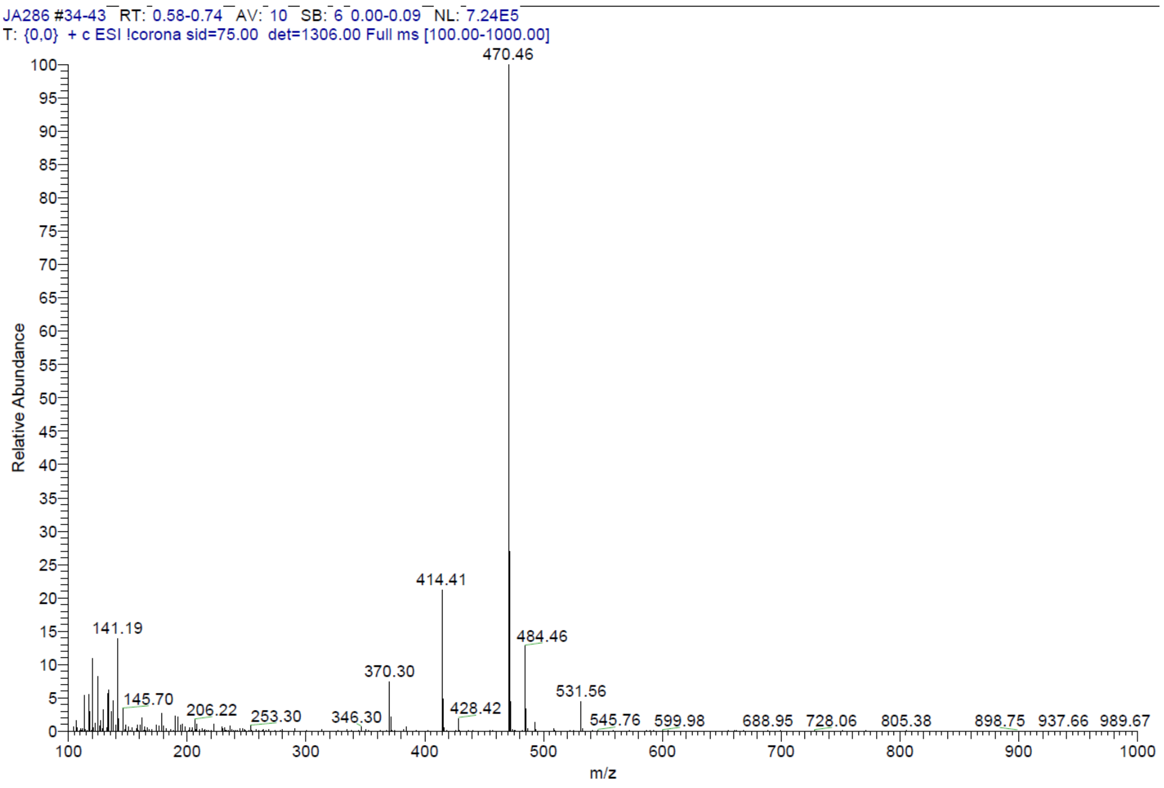


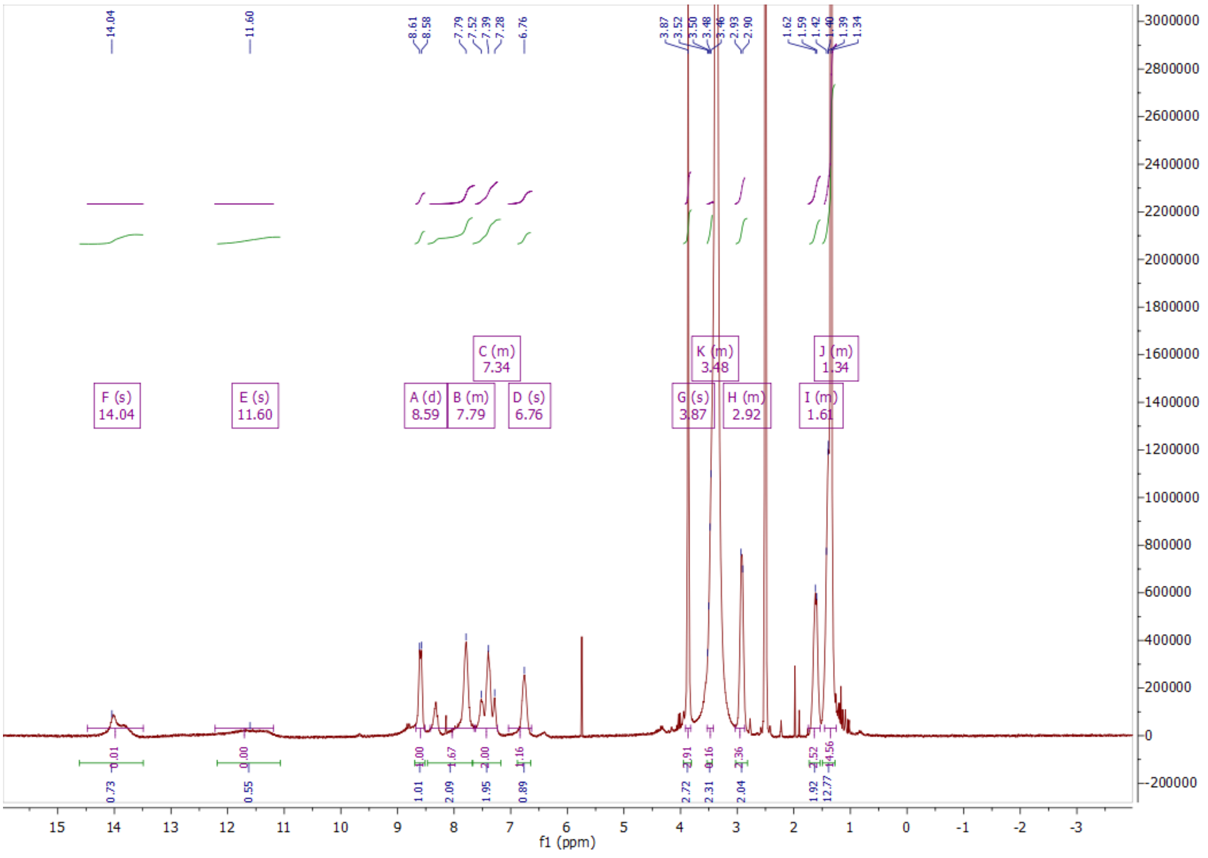

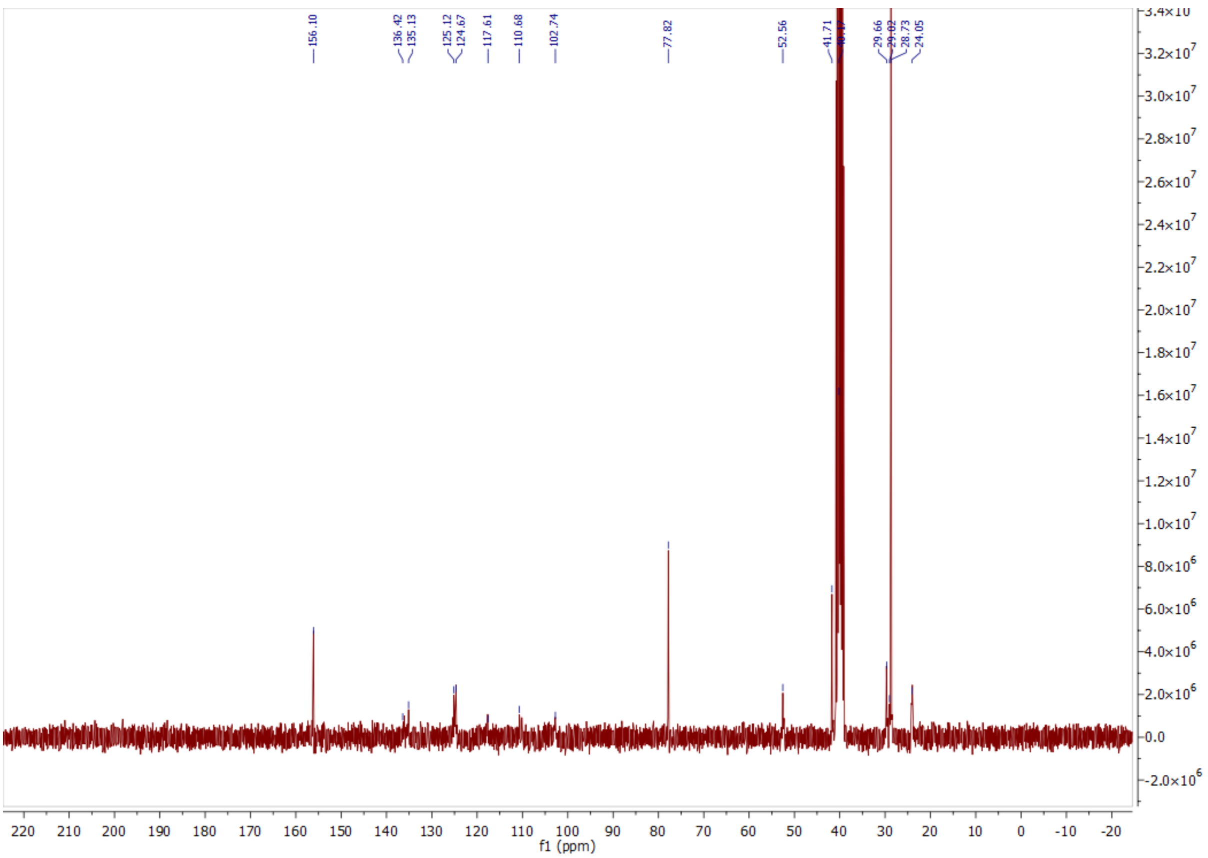
156.10
136.42
135.13
125.12
124.67
117.61
110.68
102.74
77.82
52.56
41.71
29.66
28.73
24.05
f1 (ppm)

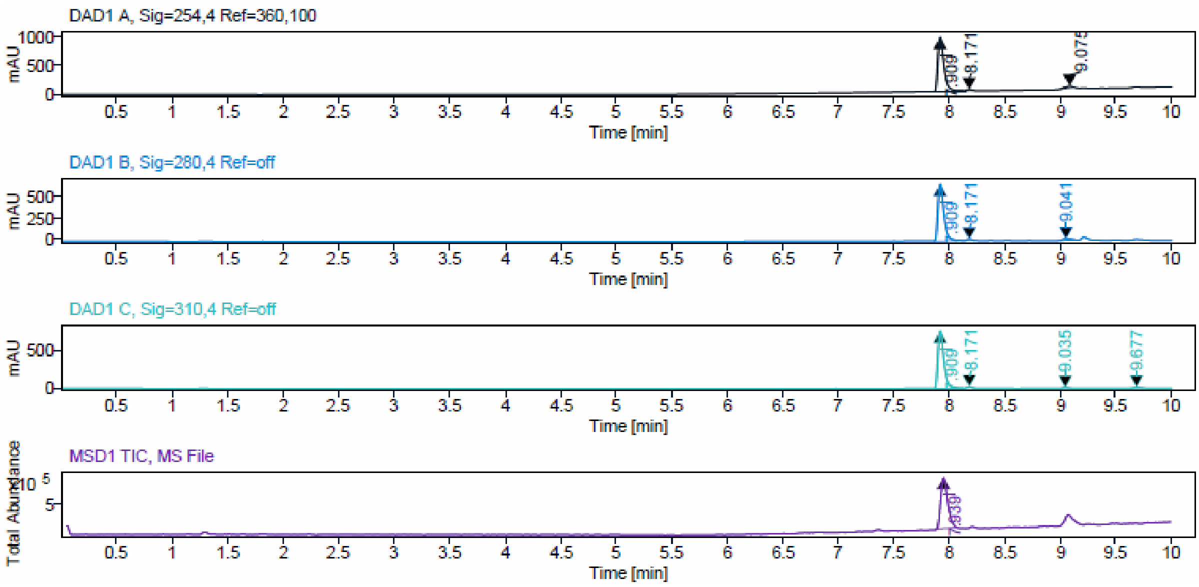
DAD1 A, Sig=254,4 Ref=360,100
mAU
8.171
Time [min]
DAD1 B, Sig=280,4 Ref=off
mAU
8.171
9.041
Time [min]
DAD1 C, Sig=310,4 Ref=off
mAU
8.171
9.035
9.677
Time [min]
MSD1 TIC, MS File
Total Abundance
Time [min]

**Sample Purity**

Signal Description DAD1 A, Sig=254,4 Ref=360,100

| Sample Name | Name | RT | Width | Area | Area% | Height |
|---|---|---|---|---|---|---|
| JA286_2_F6 | | 7.909 | 0.056 | 3739.5625 | 95.11 | 977.5157 |
| JA286_2_F6 | | 8.171 | 0.035 | 36.5721 | 0.93 | 17.3514 |
| JA286_2_F6 | | 9.075 | 0.069 | 155.7726 | 3.96 | 34.1628 |

Max Area% 95.108

UV Signal Purity>95% Pass

Signal Description DAD1 B, Sig=280,4 Ref=off

| Sample Name | Name | RT | Width | Area | Area% | Height |
|---|---|---|---|---|---|---|
| JA286_2_F6 | | 7.909 | 0.054 | 2451.2275 | 95.22 | 661.6134 |
| JA286_2_F6 | | 8.171 | 0.042 | 38.3186 | 1.49 | 15.1009 |
| JA286_2_F6 | | 9.041 | 0.092 | 84.6018 | 3.29 | 17.6666 |

Max Area% 95.225

UV Signal Purity>95% Pass

Signal Description DAD1 C, Sig=310,4 Ref=off

| Sample Name | Name | RT | Width | Area | Area% | Height |
|---|---|---|---|---|---|---|
| JA286_2_F6 | | 7.909 | 0.054 | 2791.7505 | 95.86 | 764.8190 |
| JA286_2_F6 | | 8.171 | 0.041 | 43.9157 | 1.51 | 17.2489 |
| JA286_2_F6 | | 9.035 | 0.053 | 34.8448 | 1.20 | 11.1154 |
| JA286_2_F6 | | 9.677 | 0.067 | 41.8919 | 1.44 | 11.1434 |

Max Area% 95.857

UV Signal Purity>95% Pass

ESI data of compound **12d**.

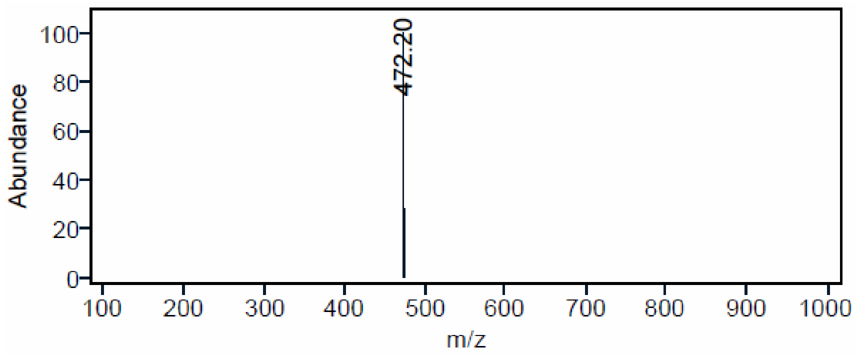

ESI data of compound **12e**.

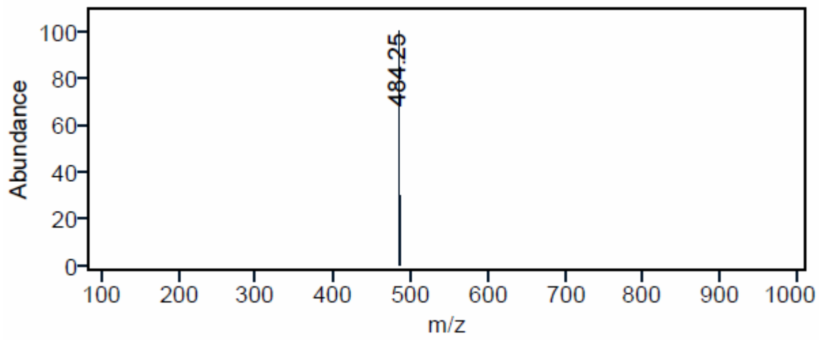


ESI data of compound **16a**.

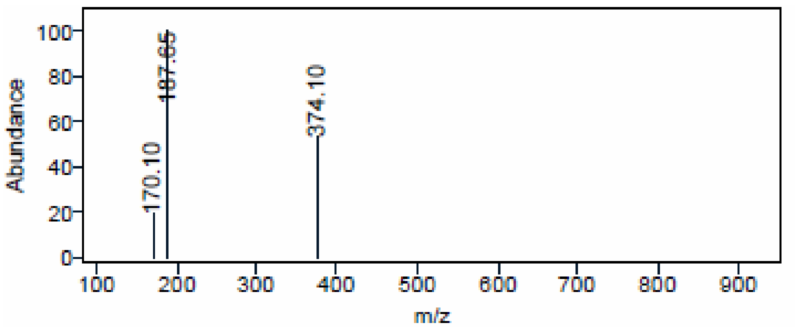


ESI data of compound **16b**.

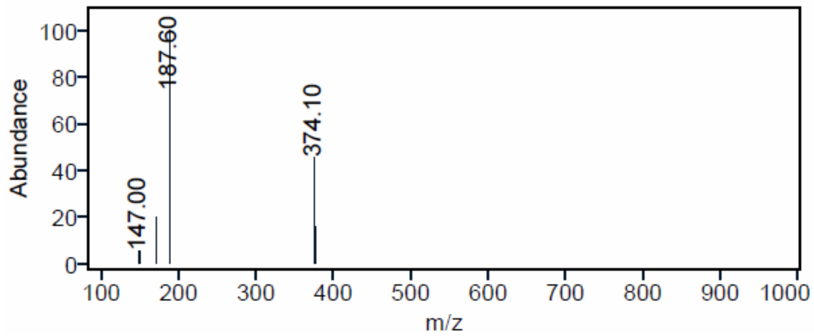


ESI data of compound **16c**.

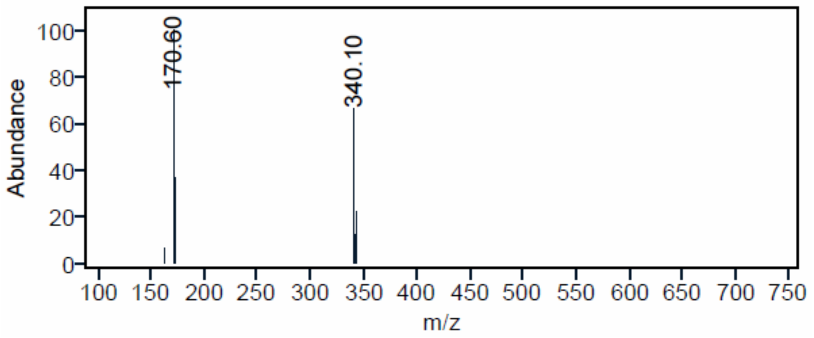

ESI data of compound **16d**.

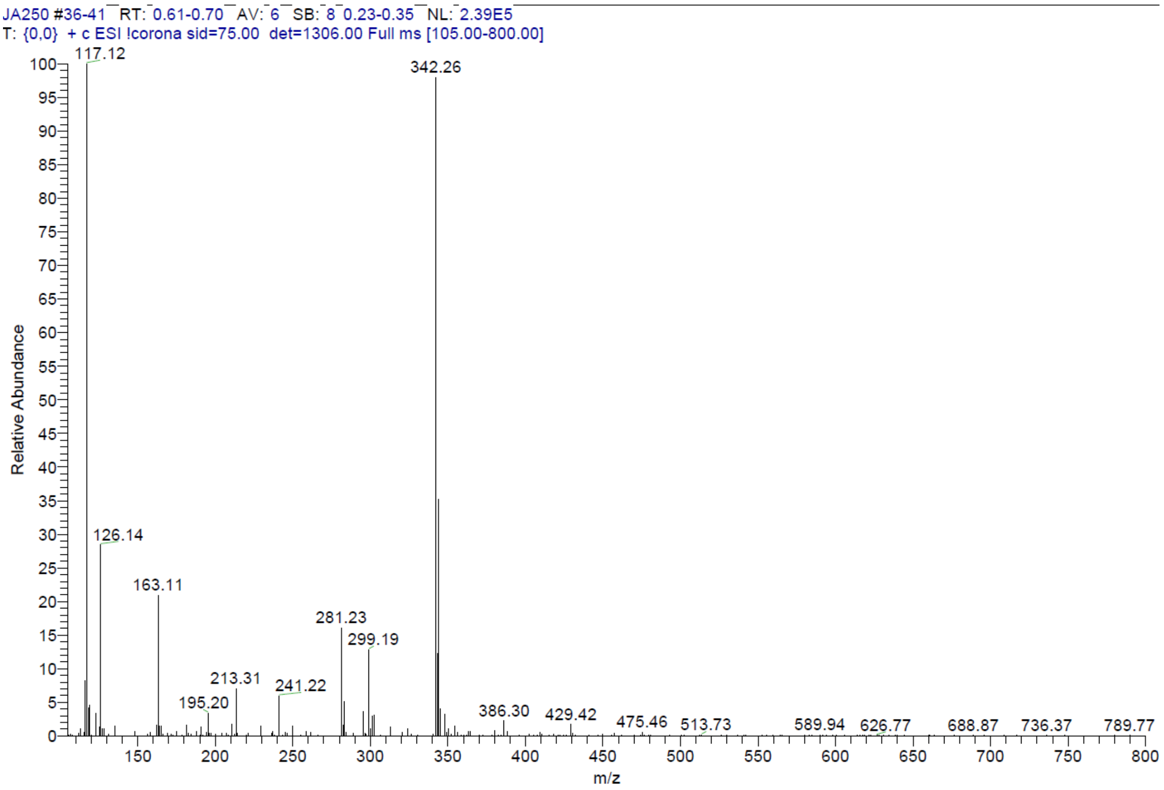


ESI data of compound **16e**.

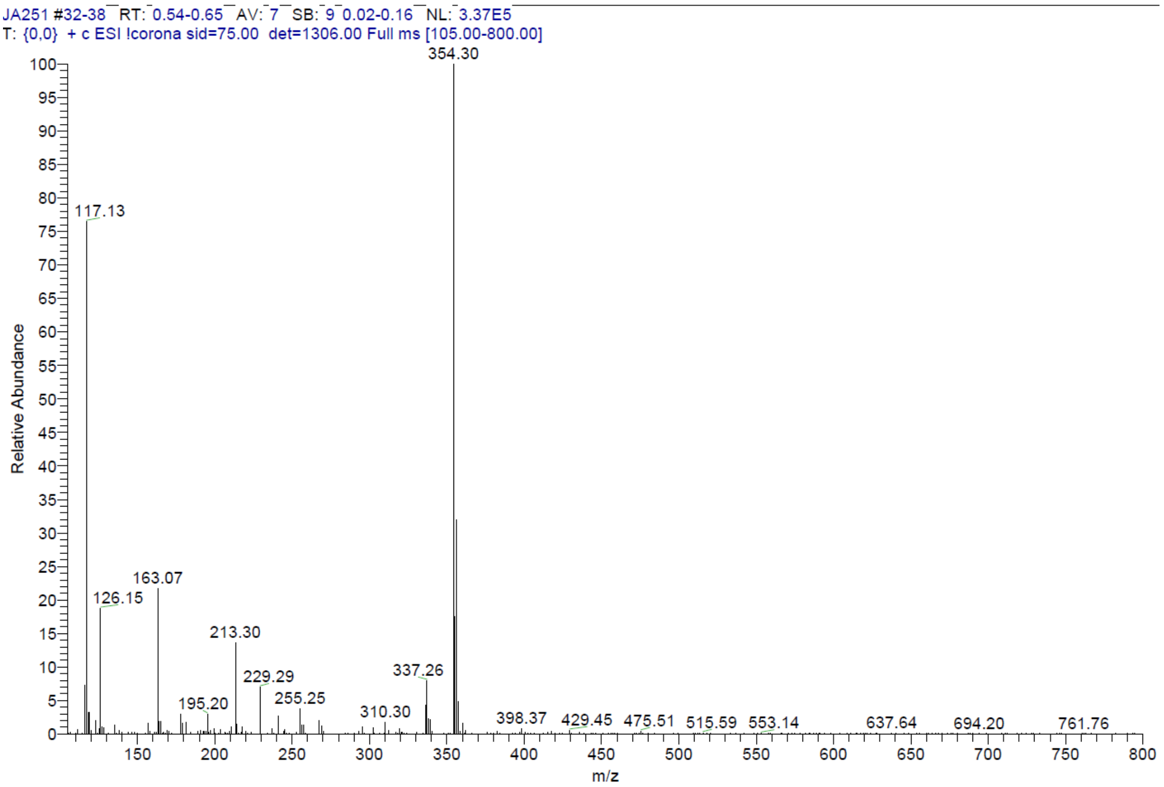

ESI data of compound **17a**.

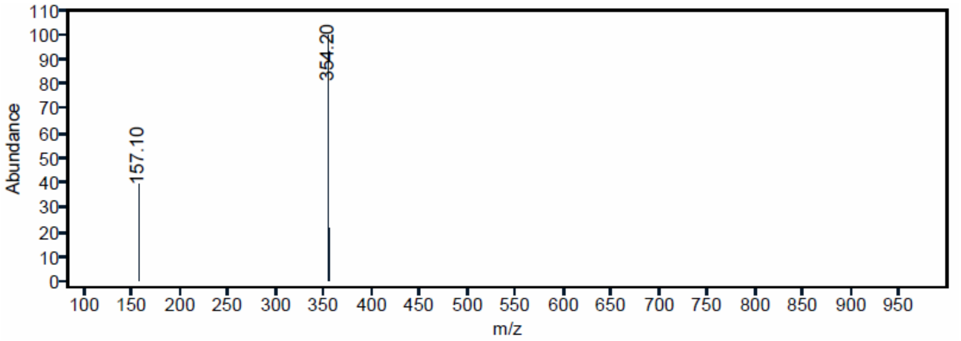


ESI data of compound **18a**.

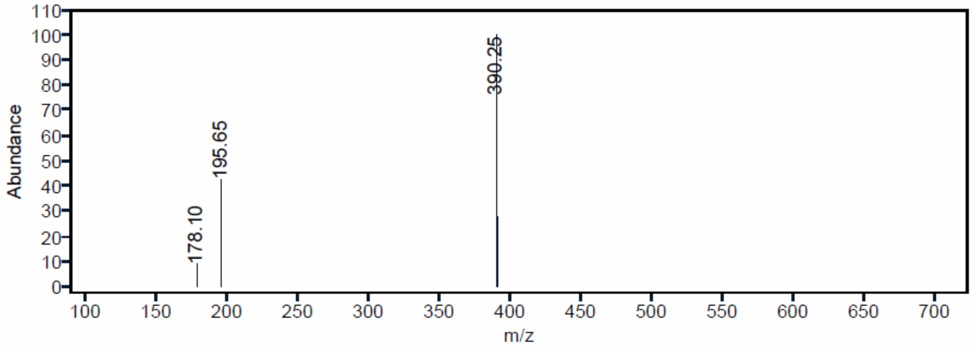


ESI data of compound **18b**.

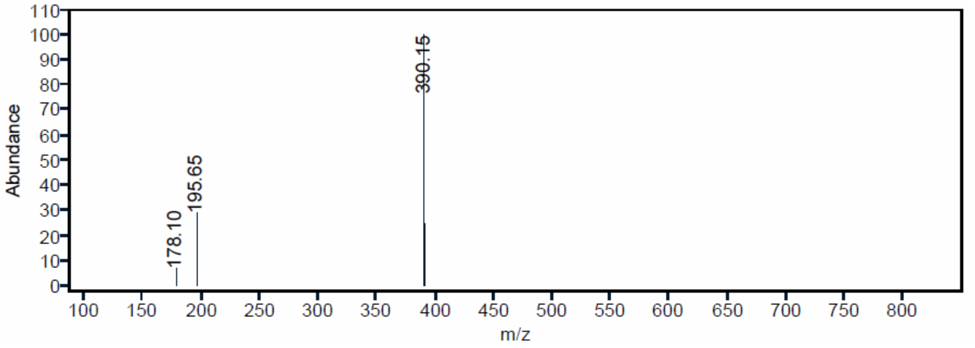

ESI data of compound **18c**.

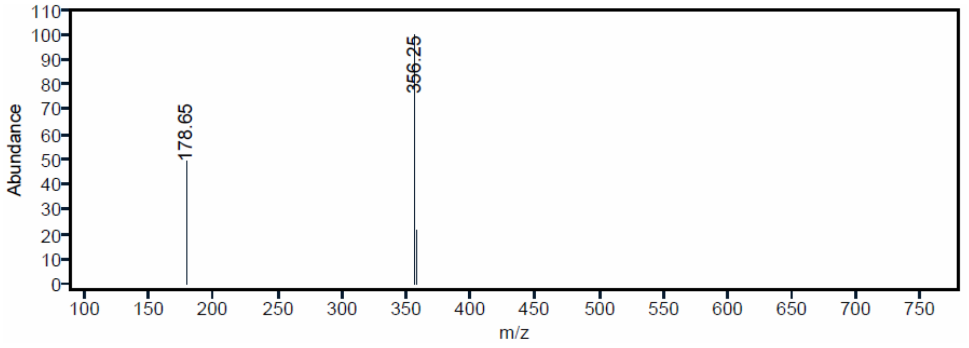


ESI data of compound **18d**.

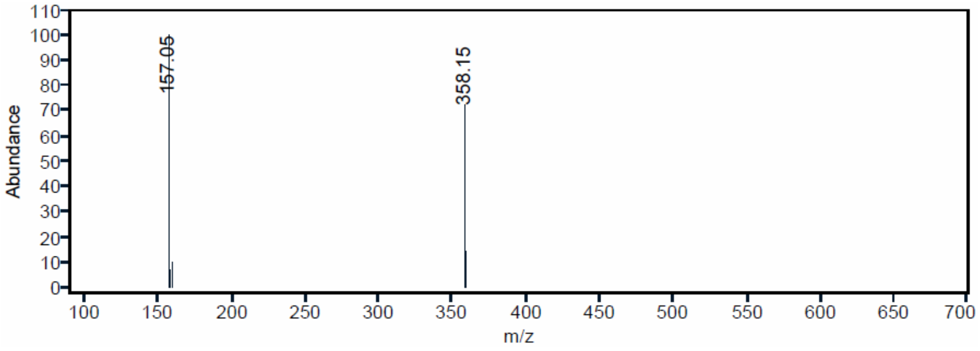


ESI data of compound **18e**.

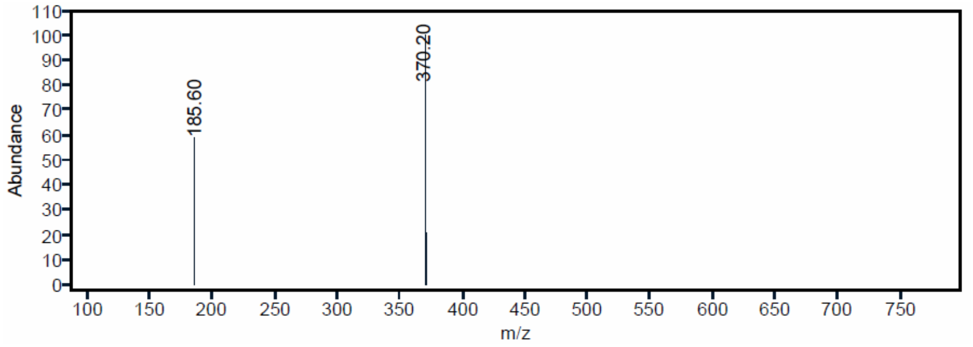

ESI, HRMS, $^{1}H$, $^{13}C$ NMR and HPLC data of compound **19a**.

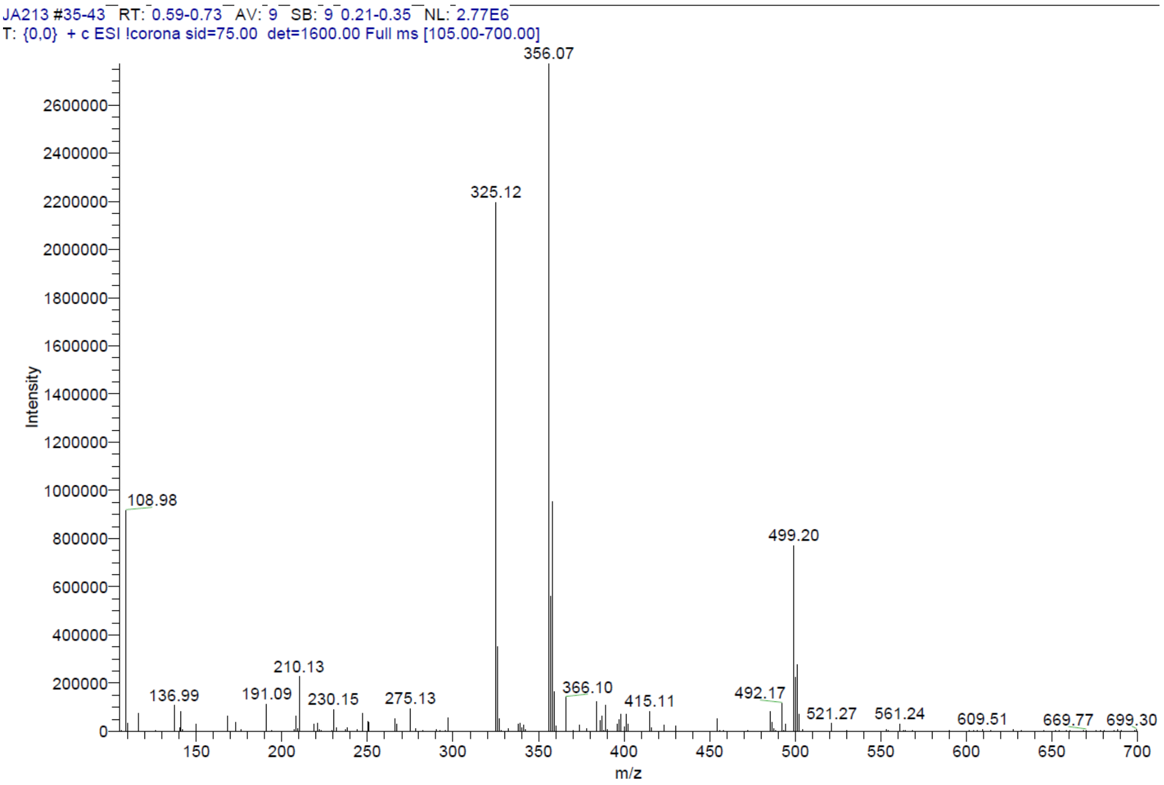


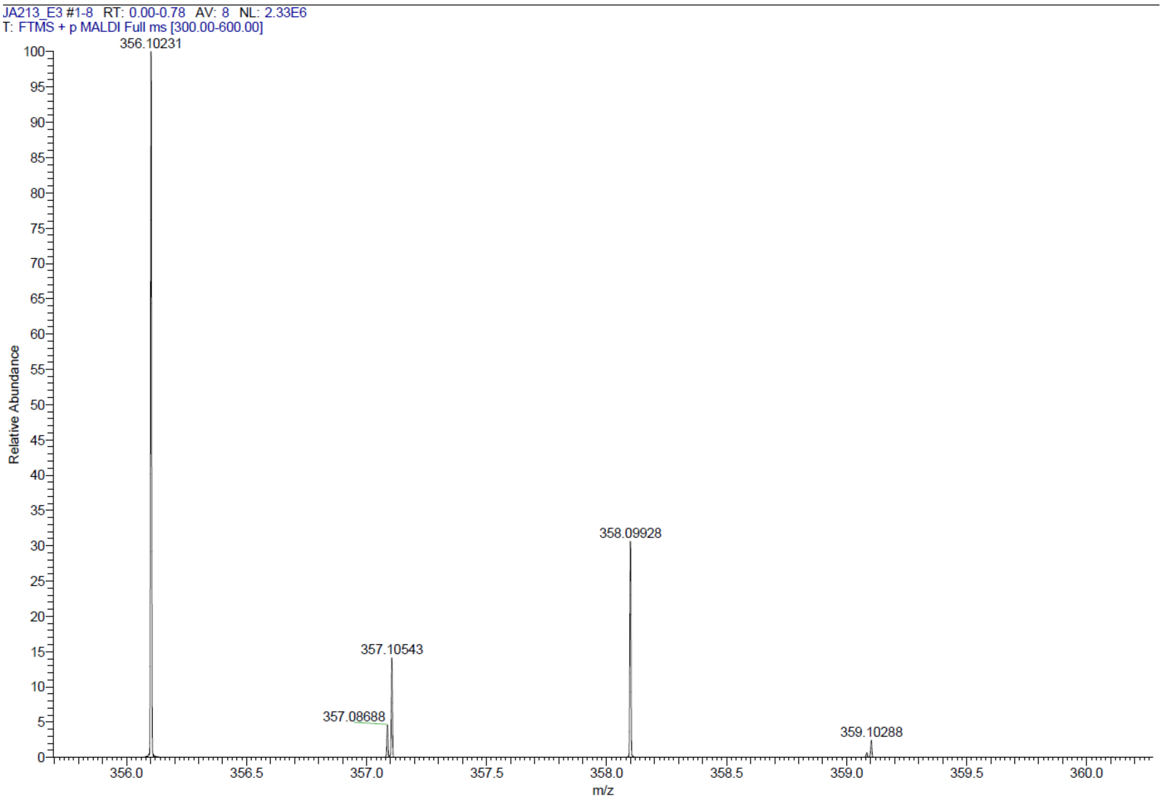

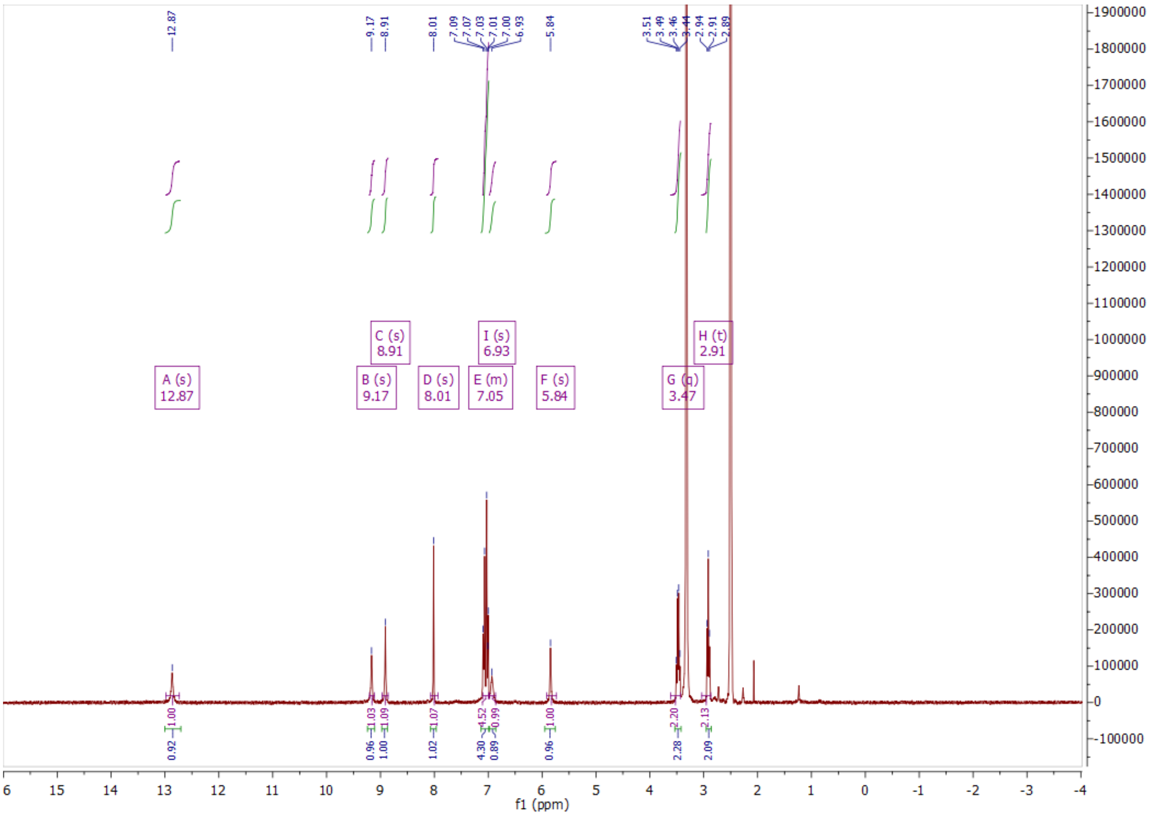
A (s)
12.87
B (s)
9.17
C (s)
8.91
D (s)
8.01
E (m)
7.05
I (s)
6.93
F (s)
5.84
G (q)
3.47
H (t)
2.91
f1 (ppm)

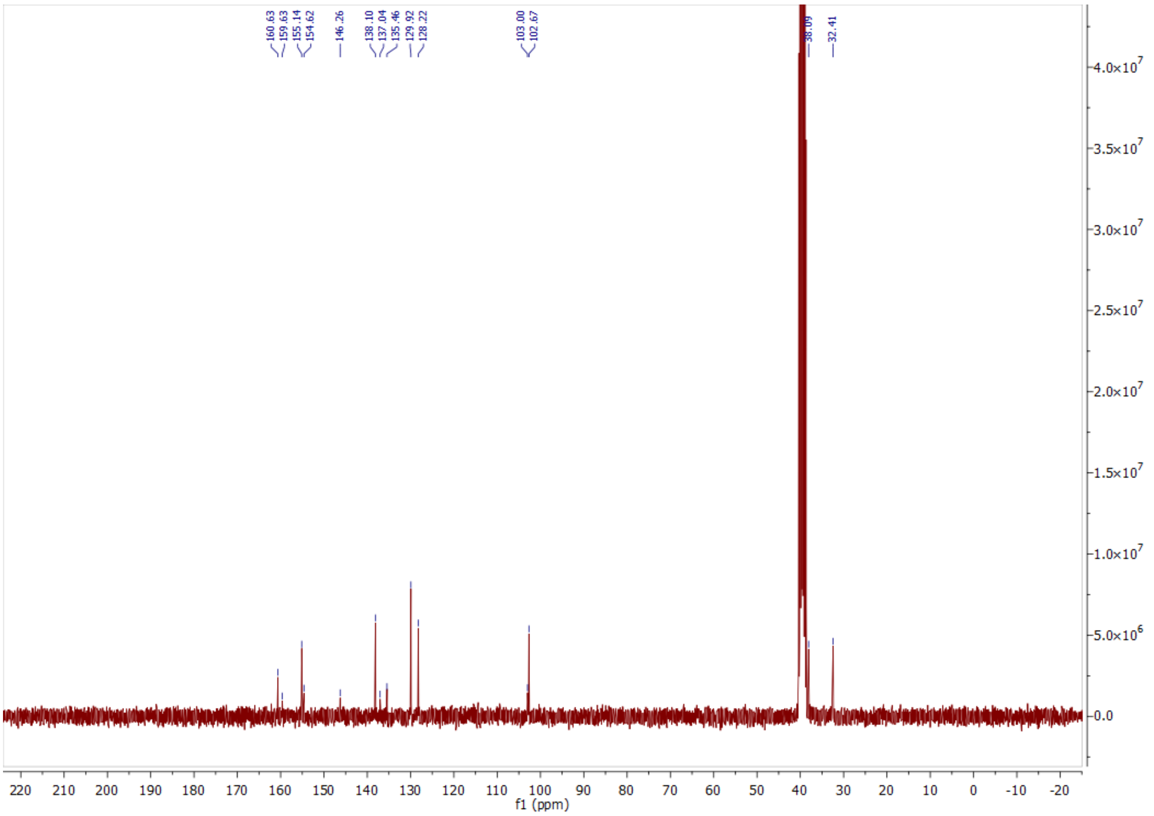
160.63
159.63
155.14
154.62
146.26
138.10
137.04
135.46
129.92
128.22
103.00
102.67
38.09
32.41
f1 (ppm)

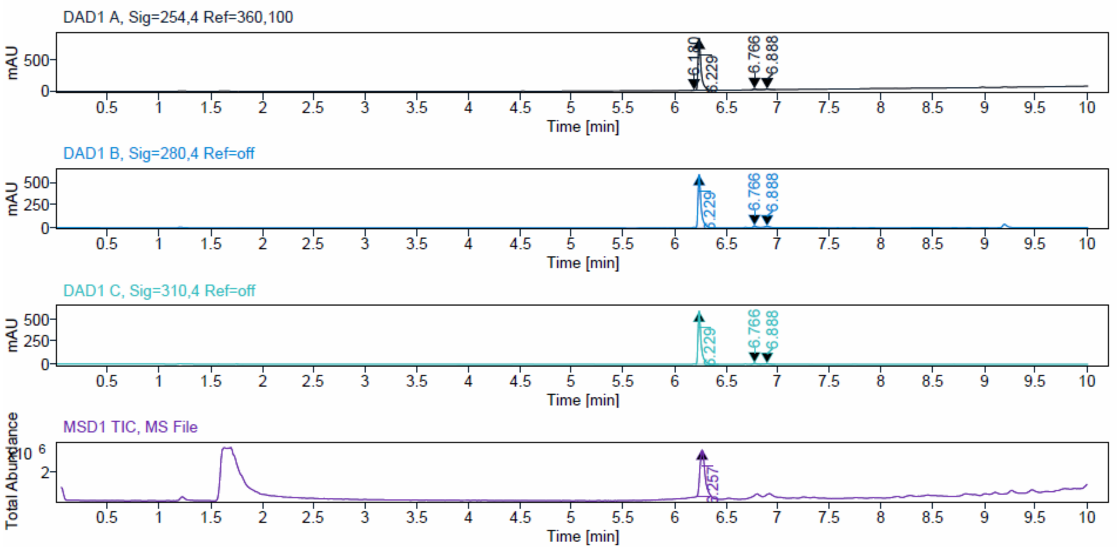


**Sample Purity**

Signal Description DAD1 A, Sig=254,4 Ref=360,100

| Sample Name | Name | RT | Width | Area | Area% | Height |
|---|---|---|---|---|---|---|
| JA213 | | 6.180 | 0.025 | 5.6884 | 0.32 | 3.6250 |
| JA213 | | 6.229 | 0.028 | 1720.6082 | 96.00 | 838.1130 |
| JA213 | | 6.766 | 0.029 | 37.2645 | 2.08 | 19.1771 |
| JA213 | | 6.888 | 0.042 | 28.7154 | 1.60 | 9.6482 |

Max Area% 96.001

UV Signal Purity>95% Pass

Signal Description DAD1 B, Sig=280,4 Ref=off

| Sample Name | Name | RT | Width | Area | Area% | Height |
|---|---|---|---|---|---|---|
| JA213 | | 6.229 | 0.028 | 1280.3722 | 95.72 | 614.5184 |
| JA213 | | 6.766 | 0.027 | 31.5171 | 2.36 | 17.7402 |
| JA213 | | 6.888 | 0.037 | 25.7224 | 1.92 | 11.4198 |

Max Area% 95.721

UV Signal Purity>95% Pass

Signal Description DAD1 C, Sig=310,4 Ref=off

| Sample Name | Name | RT | Width | Area | Area% | Height |
|---|---|---|---|---|---|---|
| JA213 | | 6.229 | 0.028 | 1258.0895 | 96.74 | 610.9068 |
| JA213 | | 6.766 | 0.029 | 25.4599 | 1.96 | 13.0771 |
| JA213 | | 6.888 | 0.042 | 16.8756 | 1.30 | 5.6521 |

Max Area% 96.744

UV Signal Purity>95% Pass

ESI, HRMS, $^{1}$H and HPLC data of compound **19b**.

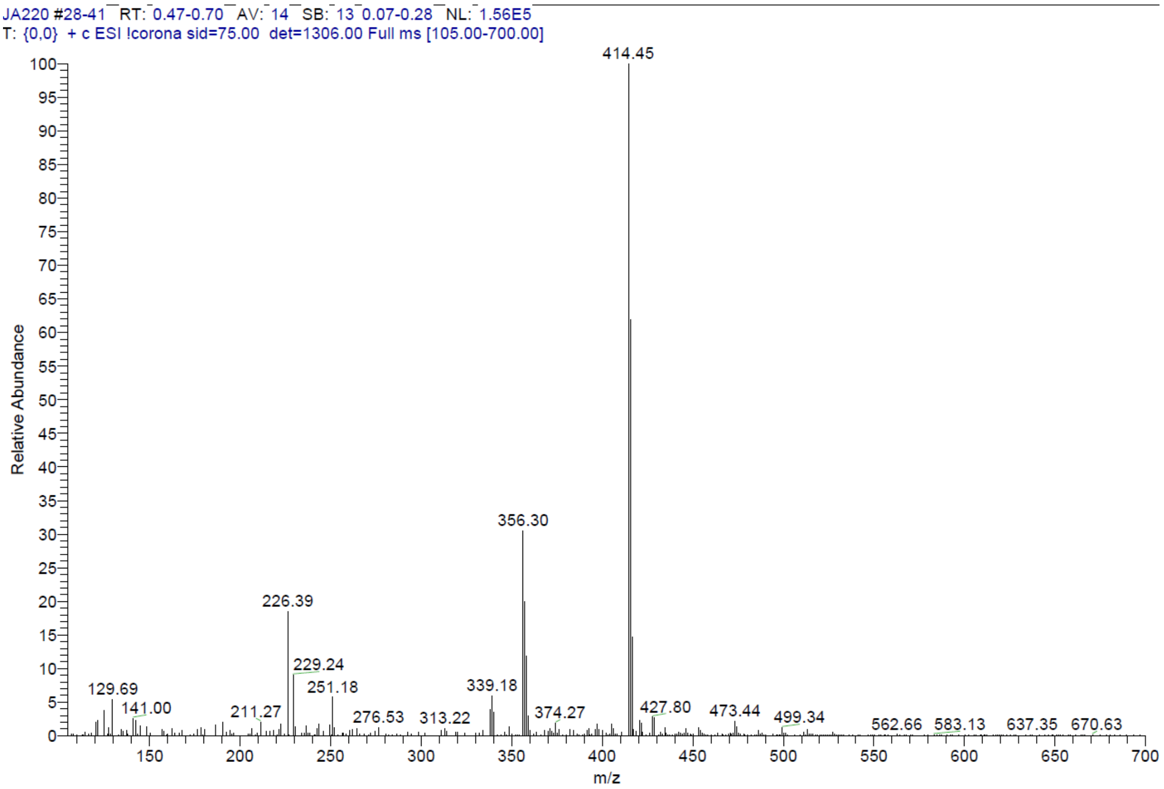


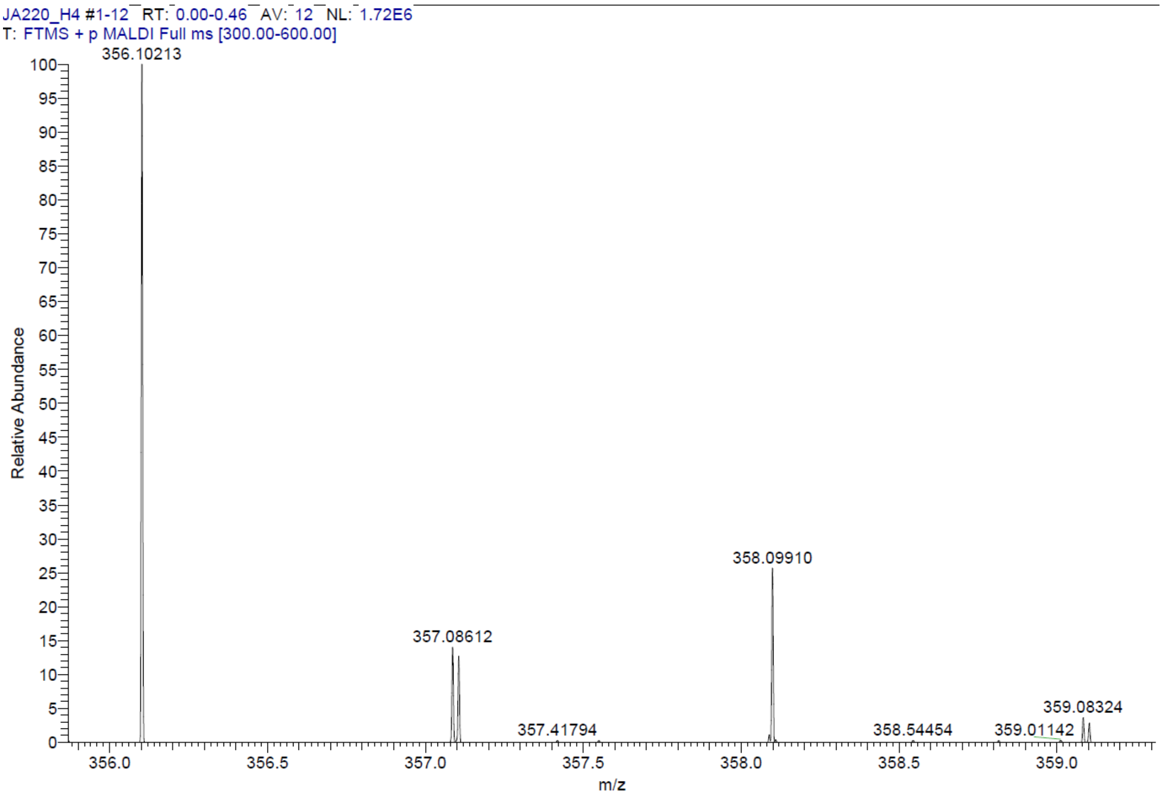

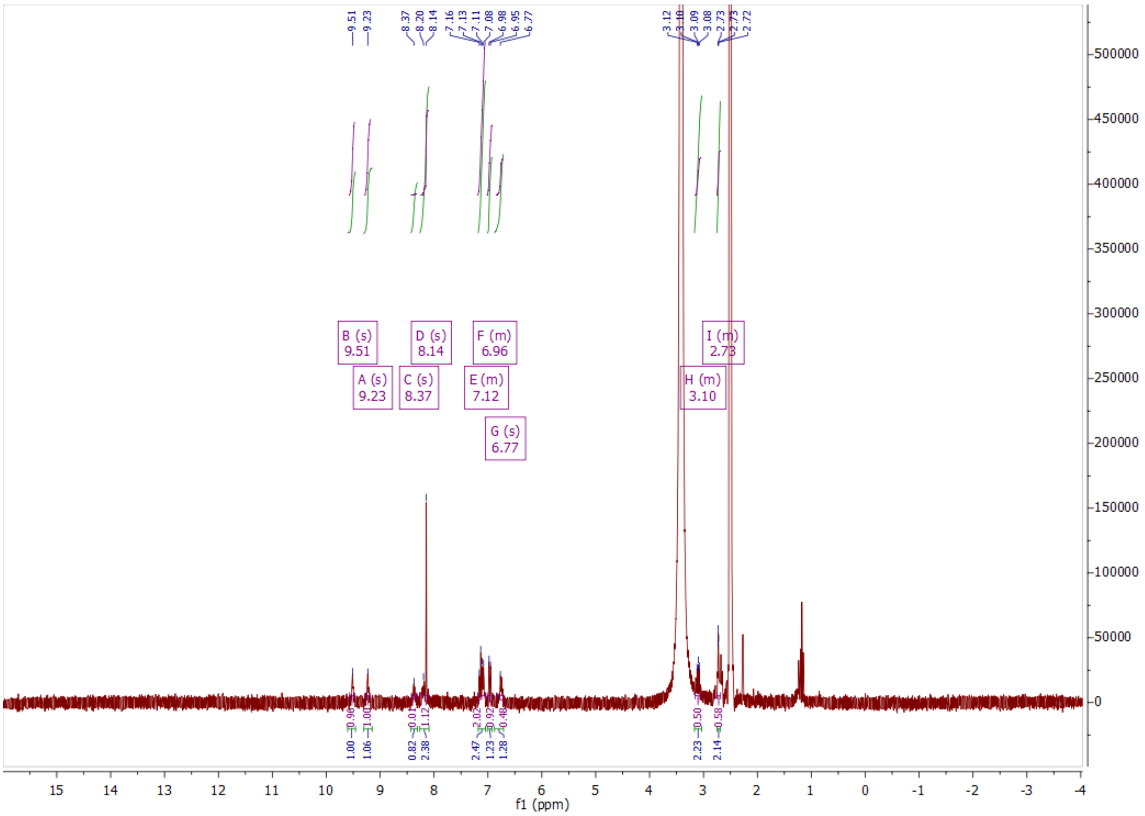
B (s)
9.51
A (s)
9.23
C (s)
8.37
D (s)
8.14
E (m)
7.12
F (m)
6.96
G (s)
6.77
H (m)
3.10
I (m)
2.73
f1 (ppm)

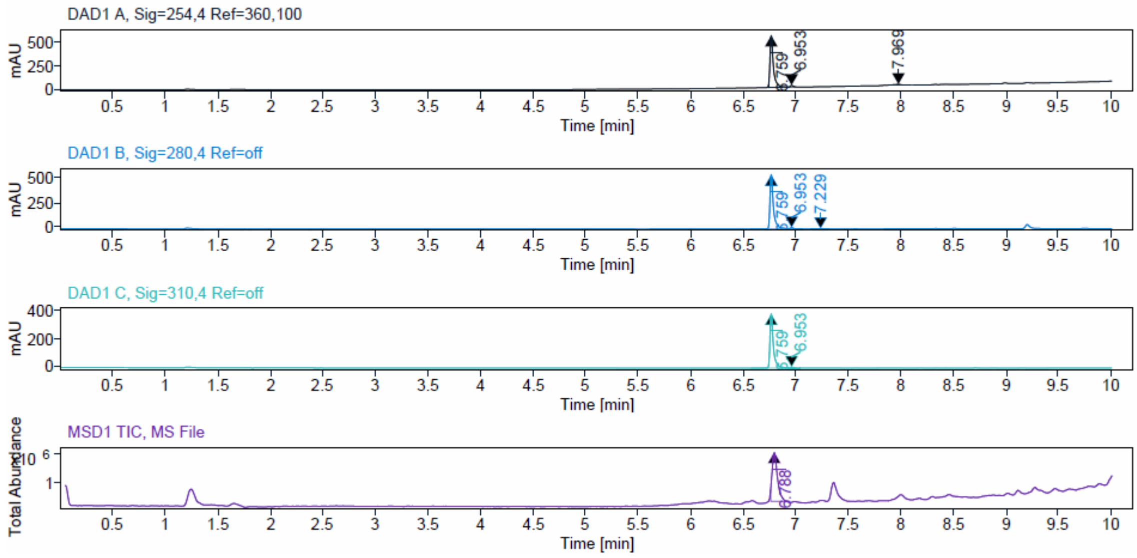
DAD1 A, Sig=254,4 Ref=360,100
DAD1 B, Sig=280,4 Ref=off
DAD1 C, Sig=310,4 Ref=off
MSD1 TIC, MS File
mAU
Total Abundance
Time [min]

Signal Description DAD1 A, Sig=254,4 Ref=360,100

| Sample Name | Name | RT | Width | Area | Area% | Height |
|---|---|---|---|---|---|---|
| JA220_prep_F9 | | 6.759 | 0.030 | 1222.3514 | 95.83 | 560.7394 |
| JA220_prep_F9 | | 6.953 | 0.029 | 34.1272 | 2.68 | 18.7784 |
| JA220_prep_F9 | | 7.969 | 0.050 | 19.0541 | 1.49 | 7.2376 |

Max Area% 95.831

UV Signal Purity>95% Pass

Signal Description DAD1 B, Sig=280,4 Ref=off

| Sample Name | Name | RT | Width | Area | Area% | Height |
|---|---|---|---|---|---|---|
| JA220_prep_F9 | | 6.759 | 0.030 | 1249.2893 | 97.16 | 552.7388 |
| JA220_prep_F9 | | 6.953 | 0.032 | 30.1885 | 2.35 | 12.0851 |
| JA220_prep_F9 | | 7.229 | 0.079 | 6.3587 | 0.49 | 1.2462 |

Max Area% 97.158

UV Signal Purity>95% Pass

Signal Description DAD1 C, Sig=310,4 Ref=off

| Sample Name | Name | RT | Width | Area | Area% | Height |
|---|---|---|---|---|---|---|
| JA220_prep_F9 | | 6.759 | 0.030 | 860.9752 | 98.11 | 383.5234 |
| JA220_prep_F9 | | 6.953 | 0.032 | 16.6036 | 1.89 | 6.7376 |

Max Area% 98.108

UV Signal Purity>95% Pass

ESI, HRMS, [1]H and HPLC data of compound **19c**.

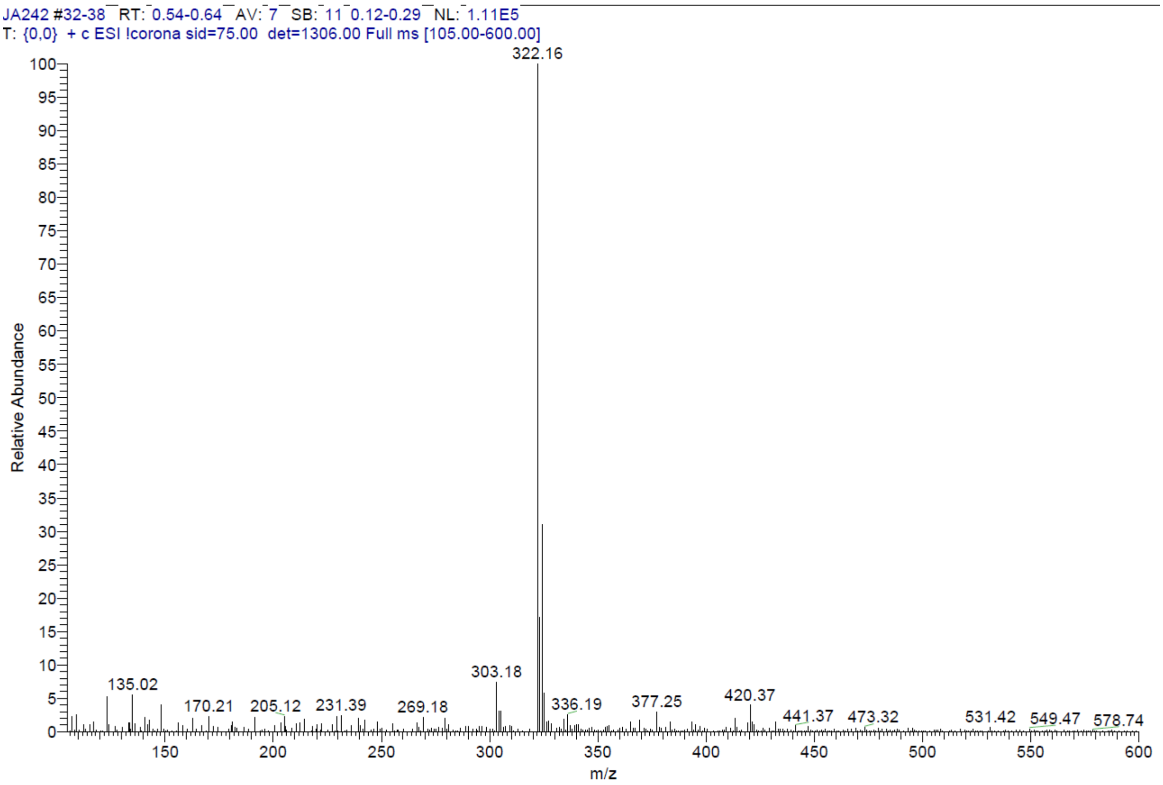


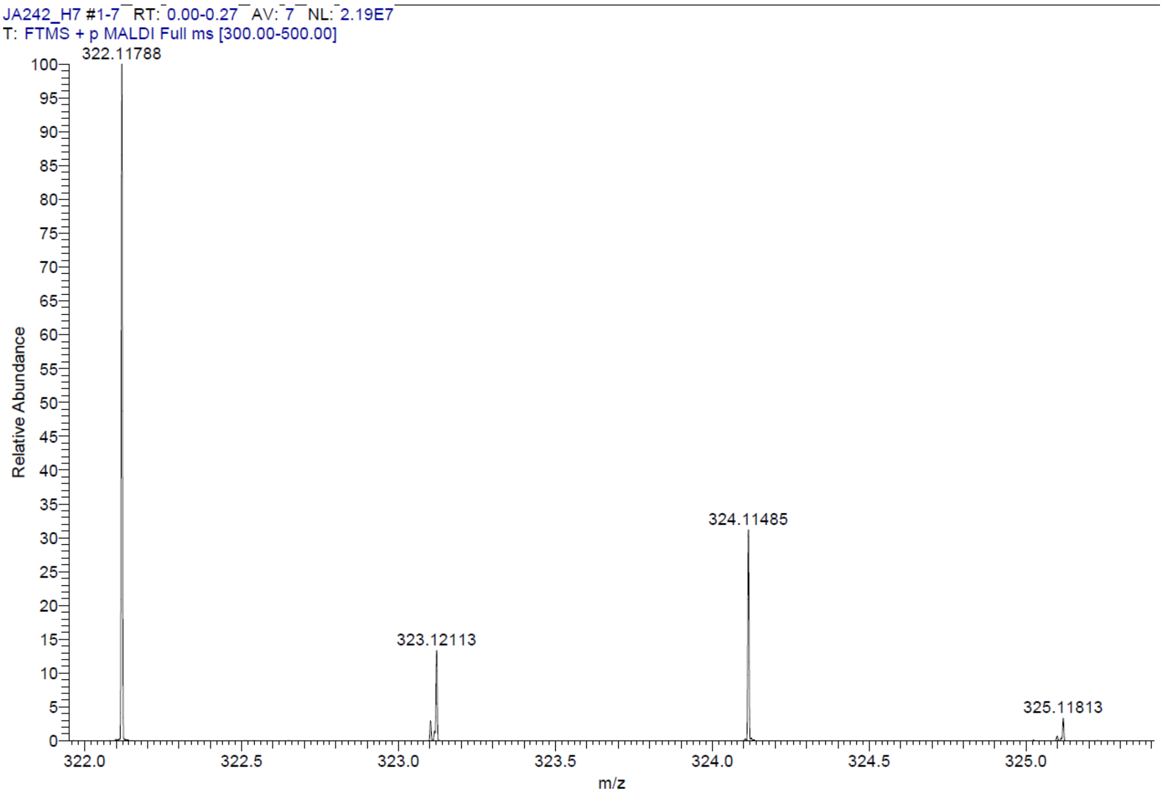

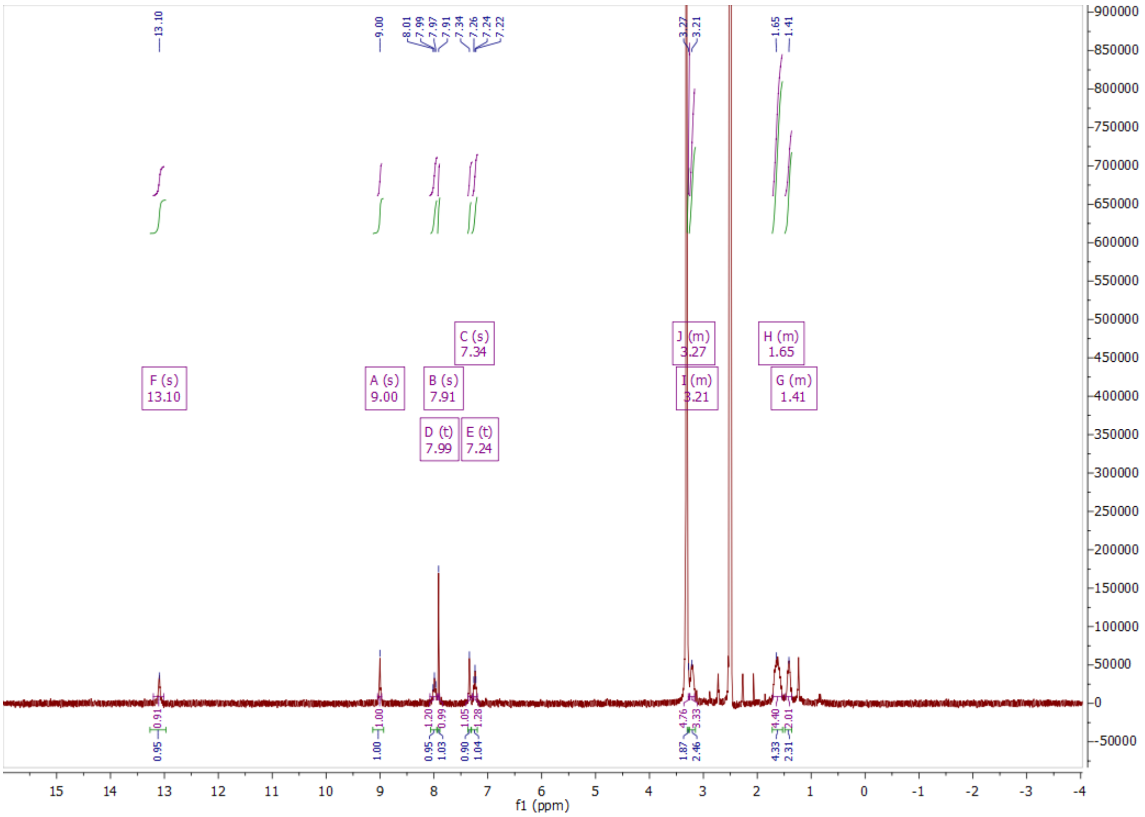

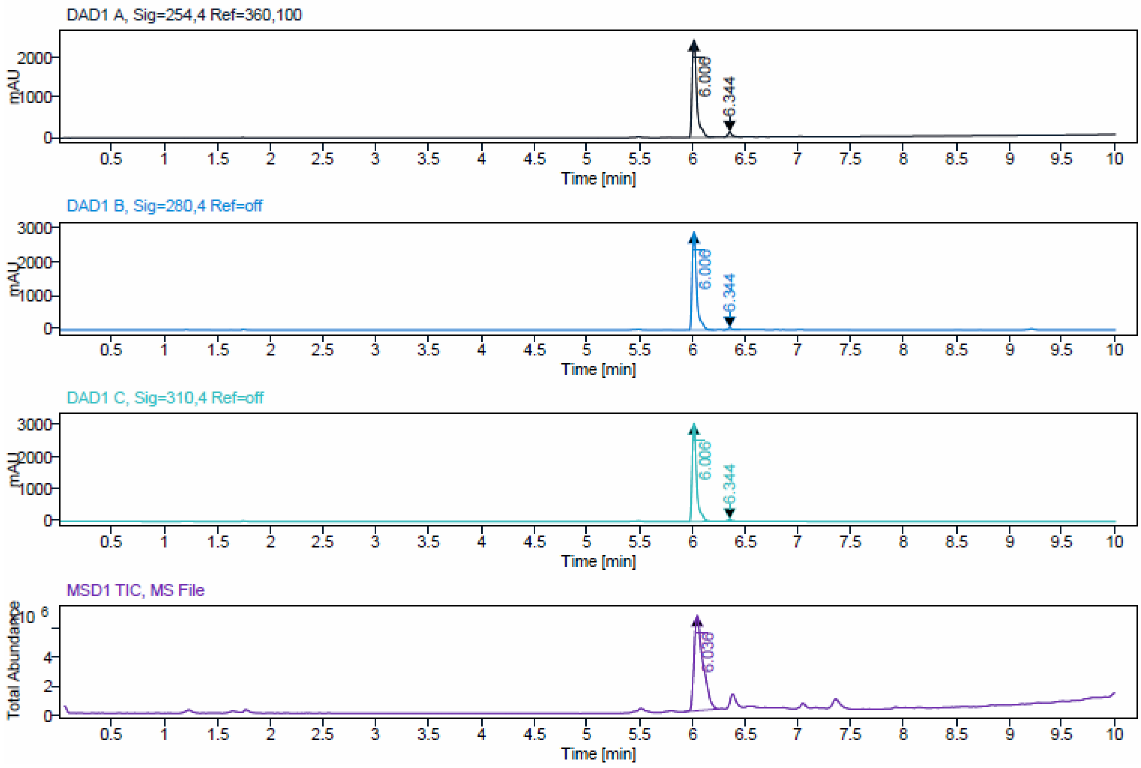

| Signal Description | DAD1 A, Sig=254,4 Ref=360,100 | | | | | |
|---|---|---|---|---|---|---|
| **Sample Name** | **Name** | **RT** | **Width** | **Area** | **Area%** | **Height** |
| JA242_F23 | | 6.006 | 0.043 | 7399.1284 | 95.60 | 2435.1013 |
| JA242_F23 | | 6.344 | 0.036 | 340.4313 | 4.40 | 127.2999 |

Max Area% 95.601

UV Signal Purity>95% Pass

| Signal Description | DAD1 B, Sig=280,4 Ref=off | | | | | |
|---|---|---|---|---|---|---|
| **Sample Name** | **Name** | **RT** | **Width** | **Area** | **Area%** | **Height** |
| JA242_F23 | | 6.006 | 0.041 | 8543.2344 | 98.20 | 2912.8694 |
| JA242_F23 | | 6.344 | 0.036 | 156.2793 | 1.80 | 56.4538 |

Max Area% 98.204

UV Signal Purity>95% Pass

| Signal Description | DAD1 C, Sig=310,4 Ref=off | | | | | |
|---|---|---|---|---|---|---|
| **Sample Name** | **Name** | **RT** | **Width** | **Area** | **Area%** | **Height** |
| JA242_F23 | | 6.006 | 0.040 | 8798.6826 | 98.20 | 3064.7908 |
| JA242_F23 | | 6.344 | 0.035 | 160.8911 | 1.80 | 58.4175 |

Max Area% 98.204

UV Signal Purity>95% Pass

ESI, HRMS, $^{1}$H and HPLC data of compound **19d**.

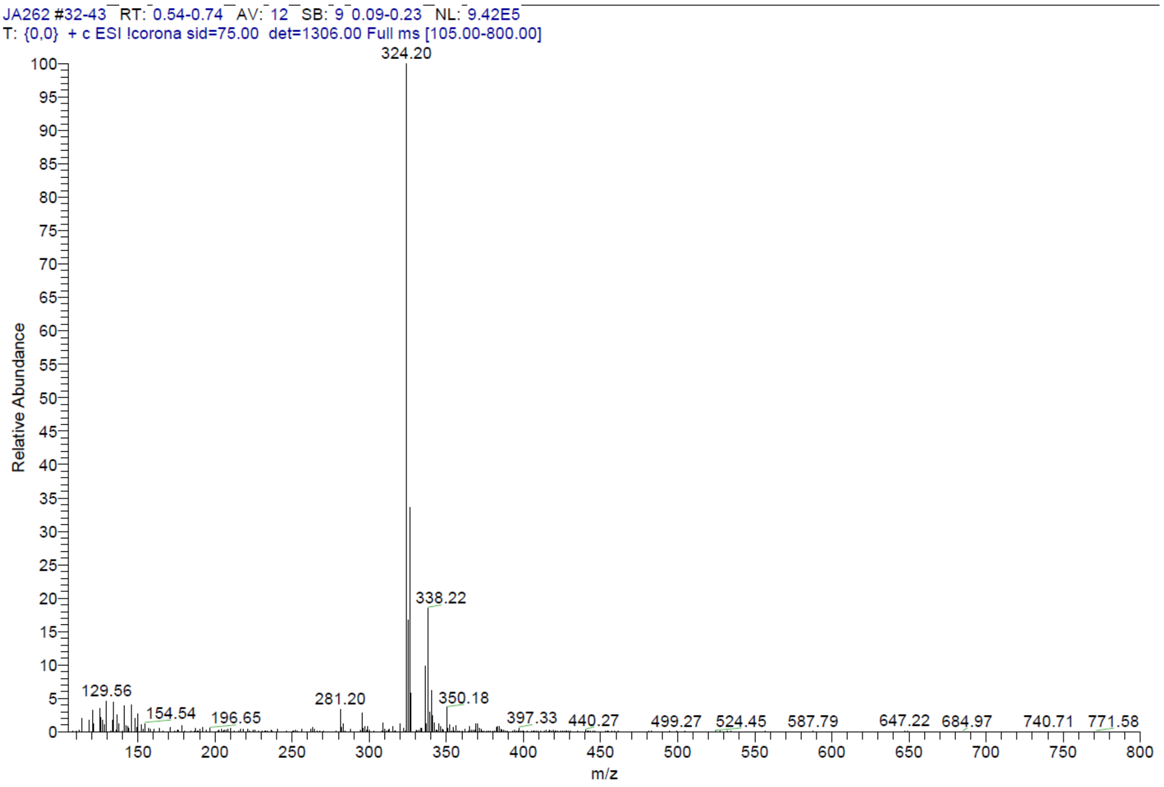

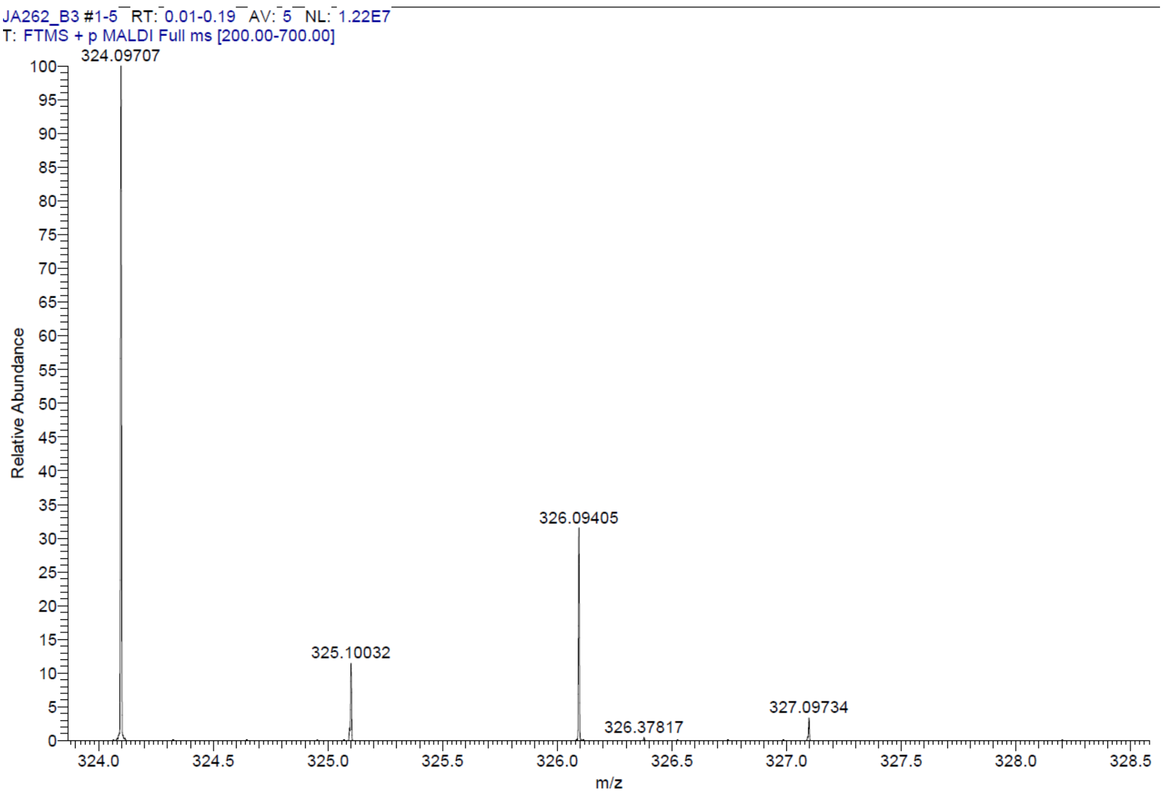

JA262_B3 #1-5 RT: 0.01-0.19 AV: 5 NL: 1.22E7
T: FTMS + p MALDI Full ms [200.00-700.00]
324.09707
325.10032
326.09405
326.37817
327.09734
Relative Abundance
m/z


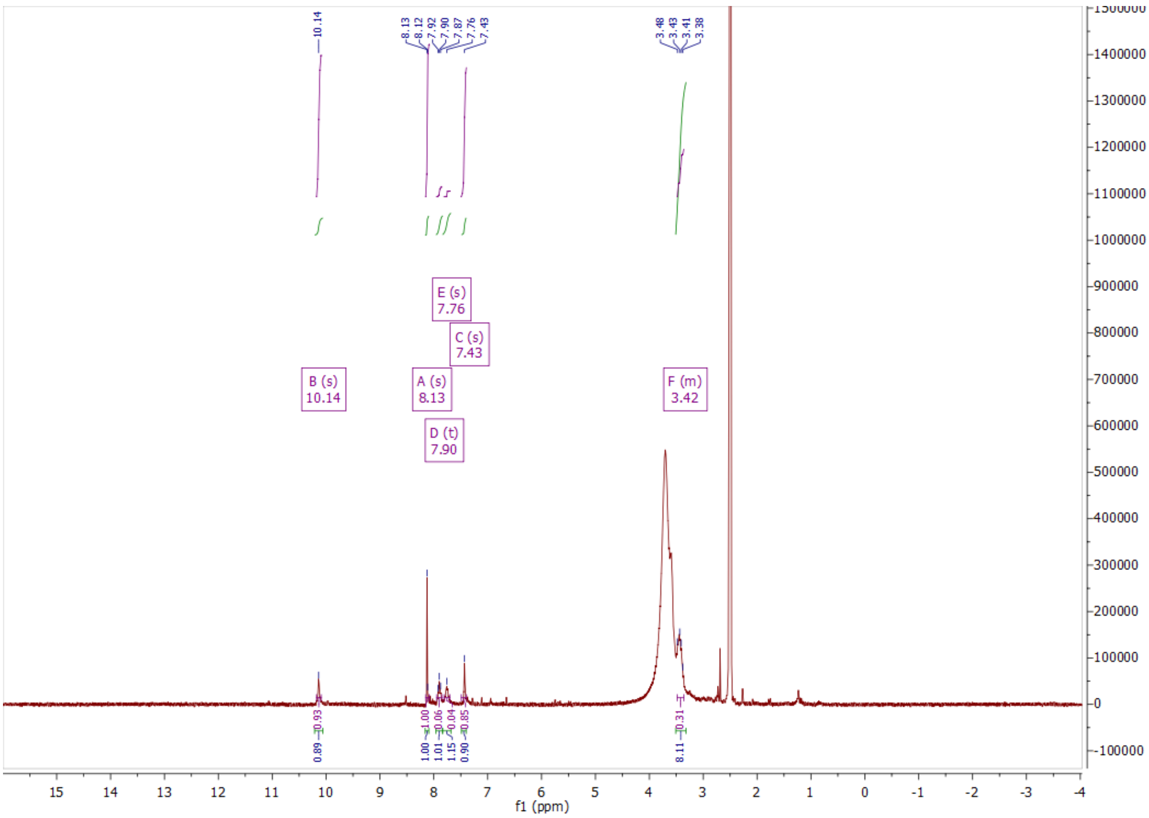

B (s)
10.14
A (s)
8.13
E (s)
7.76
C (s)
7.43
D (t)
7.90
F (m)
3.42
f1 (ppm)

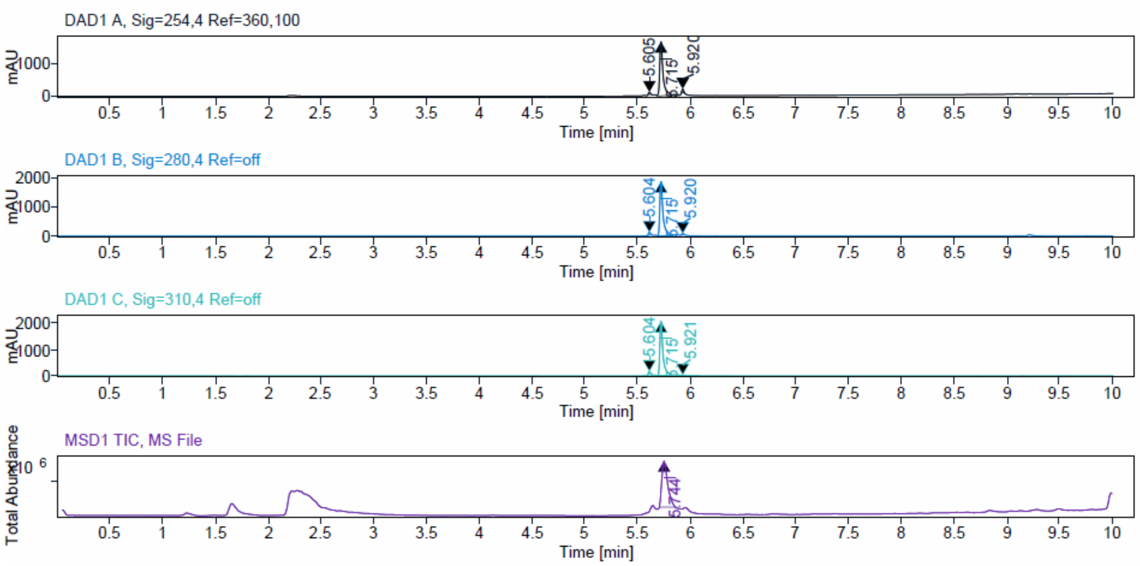


| Signal Description | DAD1 A, Sig=254,4 Ref=360,100 |
|---|---|

| Sample Name | Name | RT | Width | Area | Area% | Height |
|---|---|---|---|---|---|---|
| JA262 | | 5.605 | 0.020 | 79.2127 | 1.94 | 70.8270 |
| JA262 | | 5.715 | 0.029 | 3876.5098 | 95.10 | 1737.8448 |
| JA262 | | 5.920 | 0.018 | 120.6121 | 2.96 | 114.9335 |

| | |
|---|---|
| Max Area% | 95.098 |
| UV Signal Purity>95% | Pass |

| Signal Description | DAD1 B, Sig=280,4 Ref=off |
|---|---|

| Sample Name | Name | RT | Width | Area | Area% | Height |
|---|---|---|---|---|---|---|
| JA262 | | 5.604 | 0.023 | 132.5787 | 3.16 | 102.2514 |
| JA262 | | 5.715 | 0.028 | 3994.9648 | 95.28 | 1899.8936 |
| JA262 | | 5.920 | 0.021 | 65.3224 | 1.56 | 53.7192 |

| | |
|---|---|
| Max Area% | 95.280 |
| UV Signal Purity>95% | Pass |

| Signal Description | DAD1 C, Sig=310,4 Ref=off |
|---|---|

| Sample Name | Name | RT | Width | Area | Area% | Height |
|---|---|---|---|---|---|---|
| JA262 | | 5.604 | 0.022 | 177.5710 | 3.94 | 133.4612 |
| JA262 | | 5.715 | 0.028 | 4306.3735 | 95.64 | 2093.3418 |
| JA262 | | 5.921 | 0.022 | 18.7303 | 0.42 | 13.4436 |

| | |
|---|---|
| Max Area% | 95.640 |
| UV Signal Purity>95% | Pass |

ESI, HRMS, $^{1}H$, $^{13}C$ NMR and HPLC data of compound **19e**.

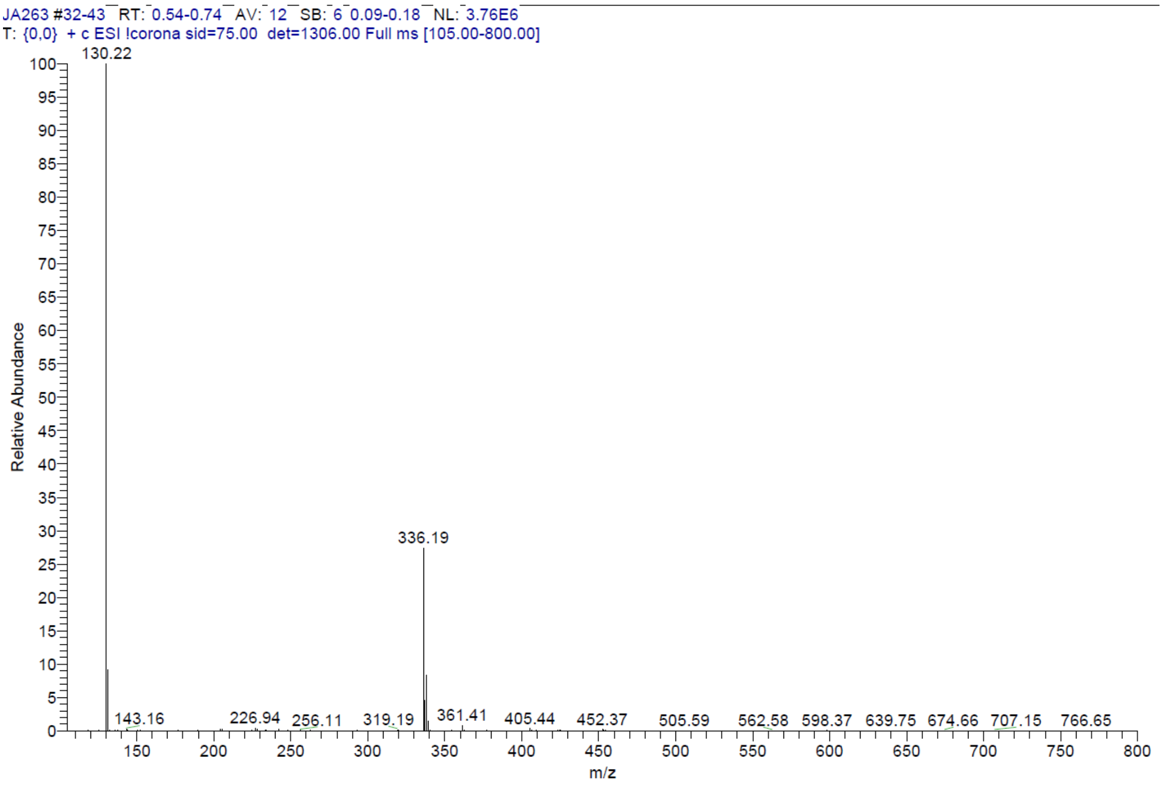


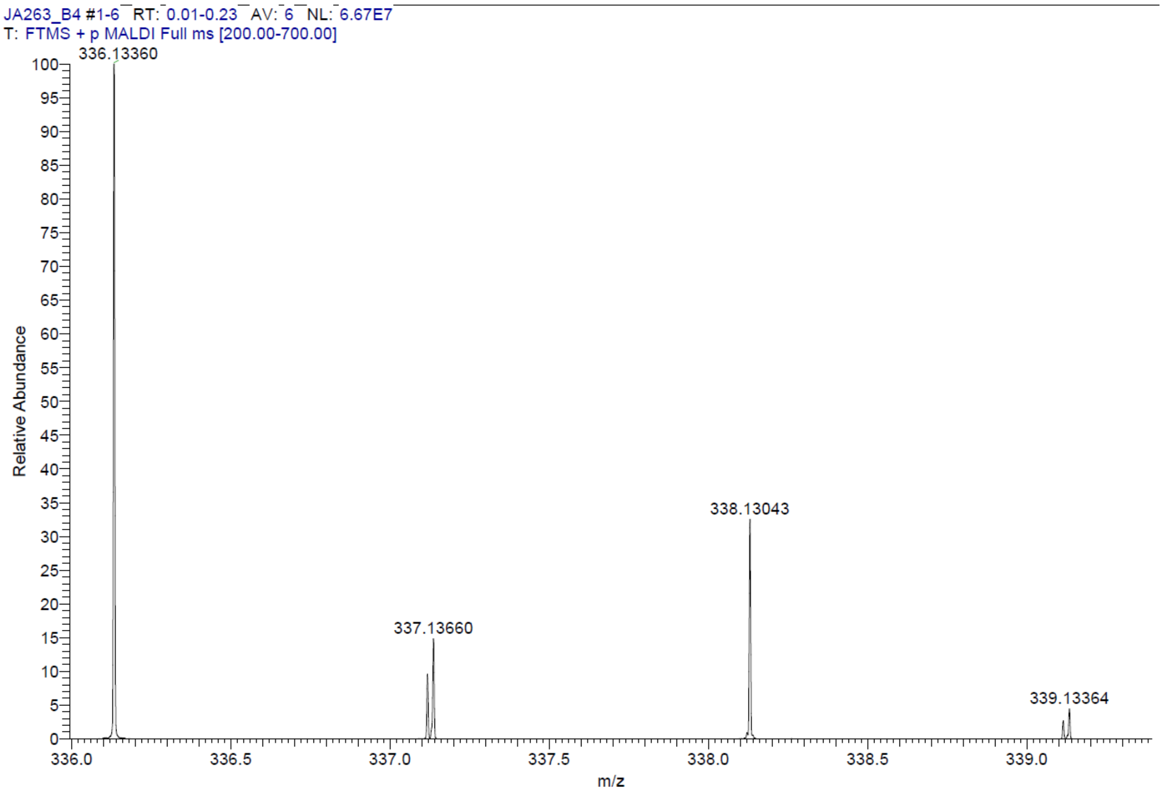

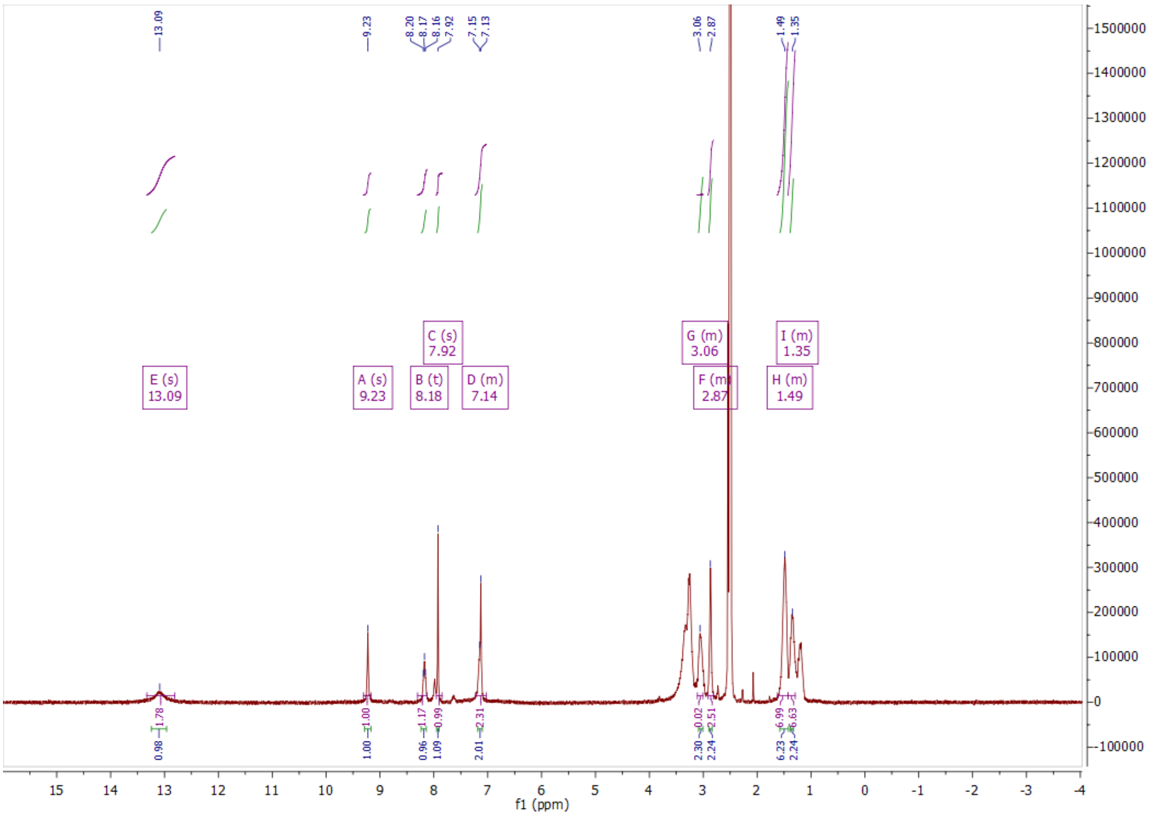
E (s)
13.09
A (s)
9.23
B (t)
8.18
C (s)
7.92
D (m)
7.14
G (m)
3.06
F (m)
2.87
I (m)
1.35
H (m)
1.49
f1 (ppm)

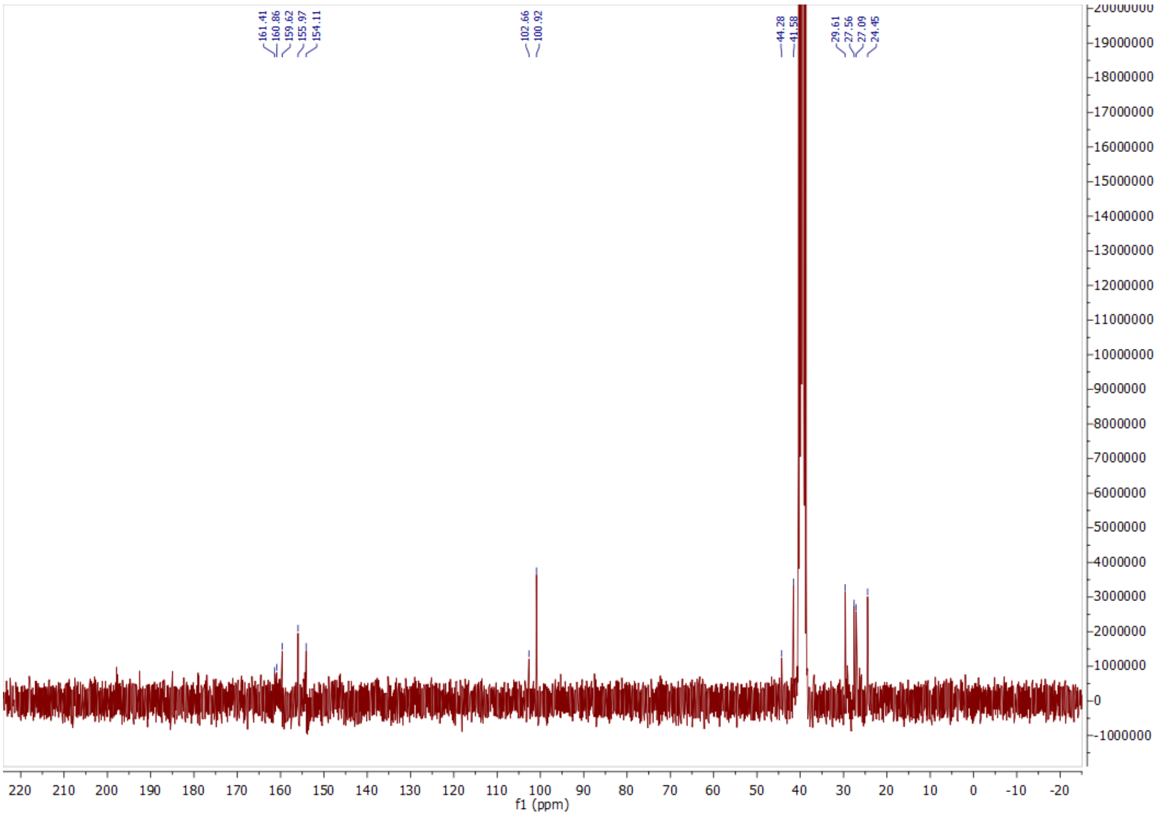
161.41
160.86
159.62
155.97
154.11
102.66
100.92
44.28
41.58
29.61
27.56
27.09
24.45
f1 (ppm)

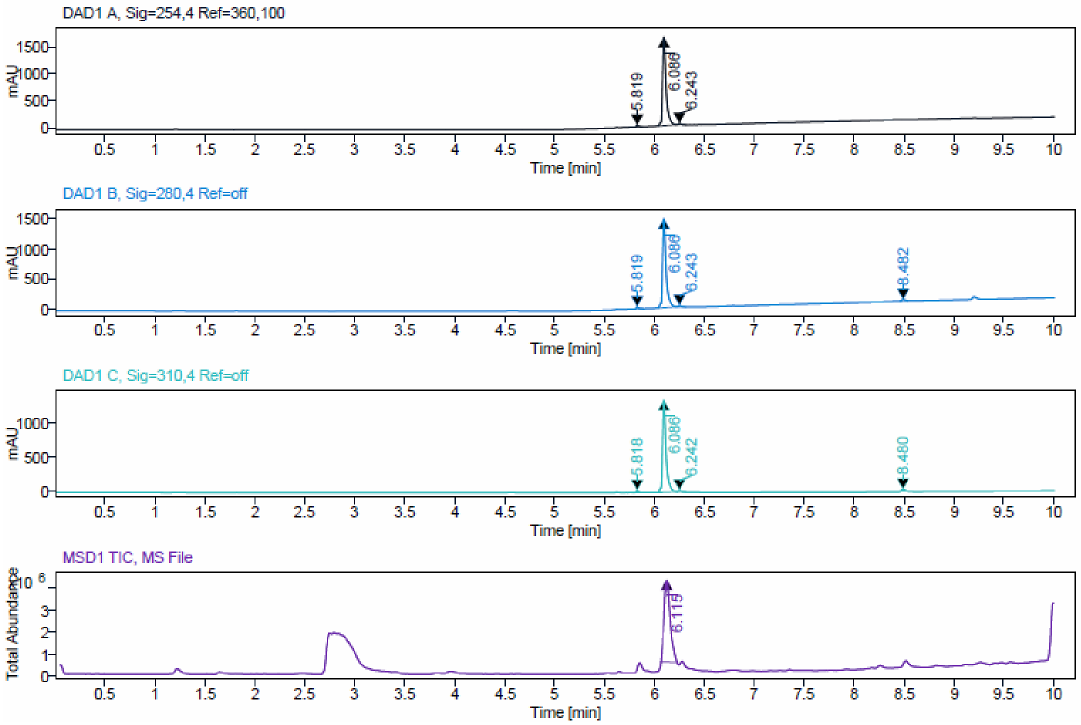


Signal Description DAD1 A, Sig=254,4 Ref=360,100

| Sample Name | Name | RT | Width | Area | Area% | Height |
|---|---|---|---|---|---|---|
| JA263_F13-15 | | 5.819 | 0.026 | 50.6392 | 1.21 | 29.6551 |
| JA263_F13-15 | | 6.086 | 0.035 | 4080.2395 | 97.60 | 1636.6354 |
| JA263_F13-15 | | 6.243 | 0.028 | 49.5384 | 1.19 | 25.7739 |

Max Area% 97.604

UV Signal Purity>95% Pass

Signal Description DAD1 B, Sig=280,4 Ref=off

| Sample Name | Name | RT | Width | Area | Area% | Height |
|---|---|---|---|---|---|---|
| JA263_F13-15 | | 5.819 | 0.026 | 56.9175 | 1.46 | 32.4069 |
| JA263_F13-15 | | 6.086 | 0.035 | 3702.4114 | 95.21 | 1493.6373 |
| JA263_F13-15 | | 6.243 | 0.029 | 42.9040 | 1.10 | 22.2771 |
| JA263_F13-15 | | 8.482 | 0.050 | 86.4594 | 2.22 | 30.5377 |

Max Area% 95.210

UV Signal Purity>95% Pass

Signal Description DAD1 C, Sig=310,4 Ref=off

| Sample Name | Name | RT | Width | Area | Area% | Height |
|---|---|---|---|---|---|---|
| JA263_F13-15 | | 5.818 | 0.026 | 57.3672 | 1.65 | 28.8671 |
| JA263_F13-15 | | 6.086 | 0.035 | 3308.8438 | 95.22 | 1334.3612 |
| JA263_F13-15 | | 6.242 | 0.027 | 38.7551 | 1.12 | 20.4256 |
| JA263_F13-15 | | 8.480 | 0.037 | 70.0378 | 2.02 | 30.1895 |

Max Area% 95.218

UV Signal Purity>95% Pass

ESI, HRMS, [1]H and HPLC data of compound **20a**.

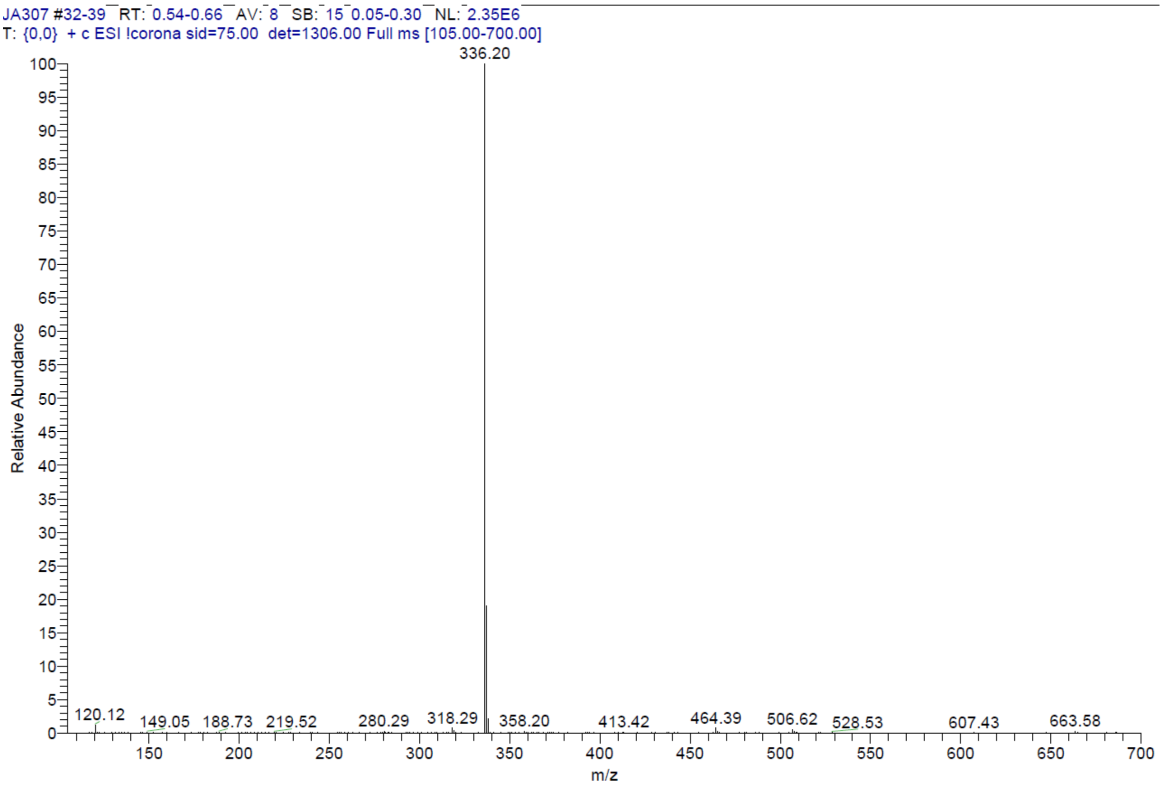


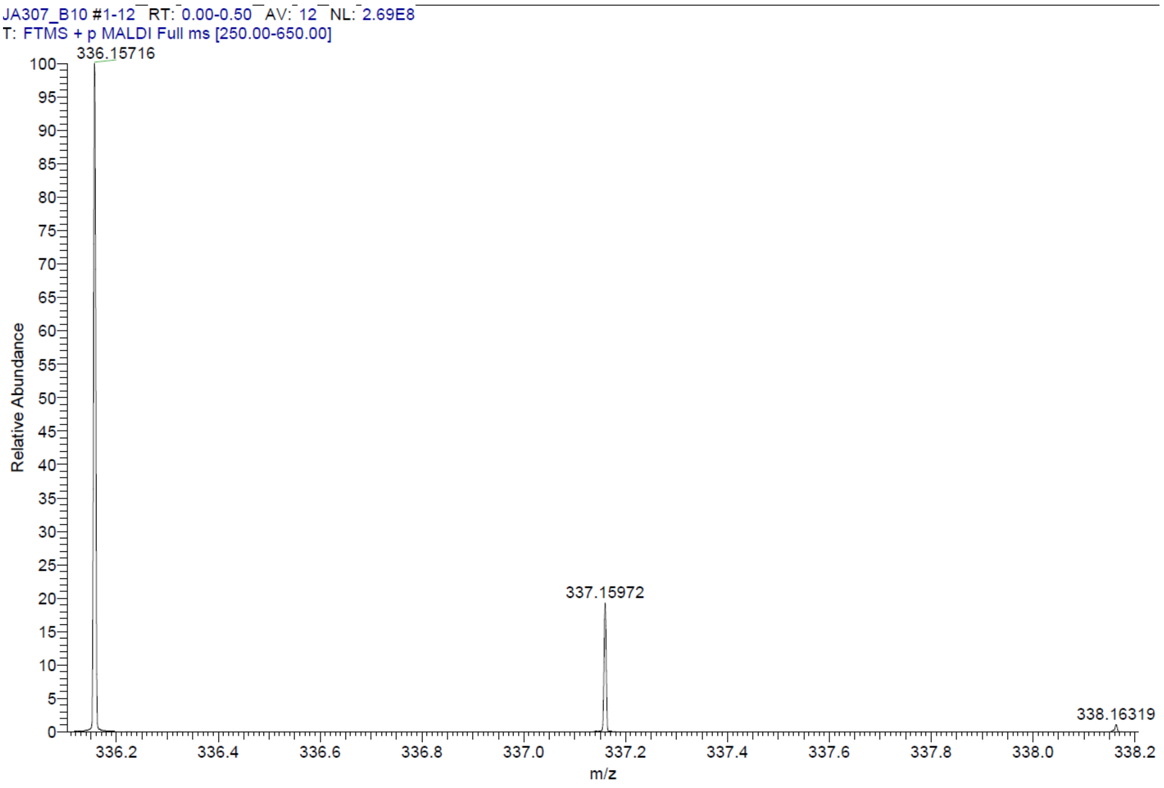

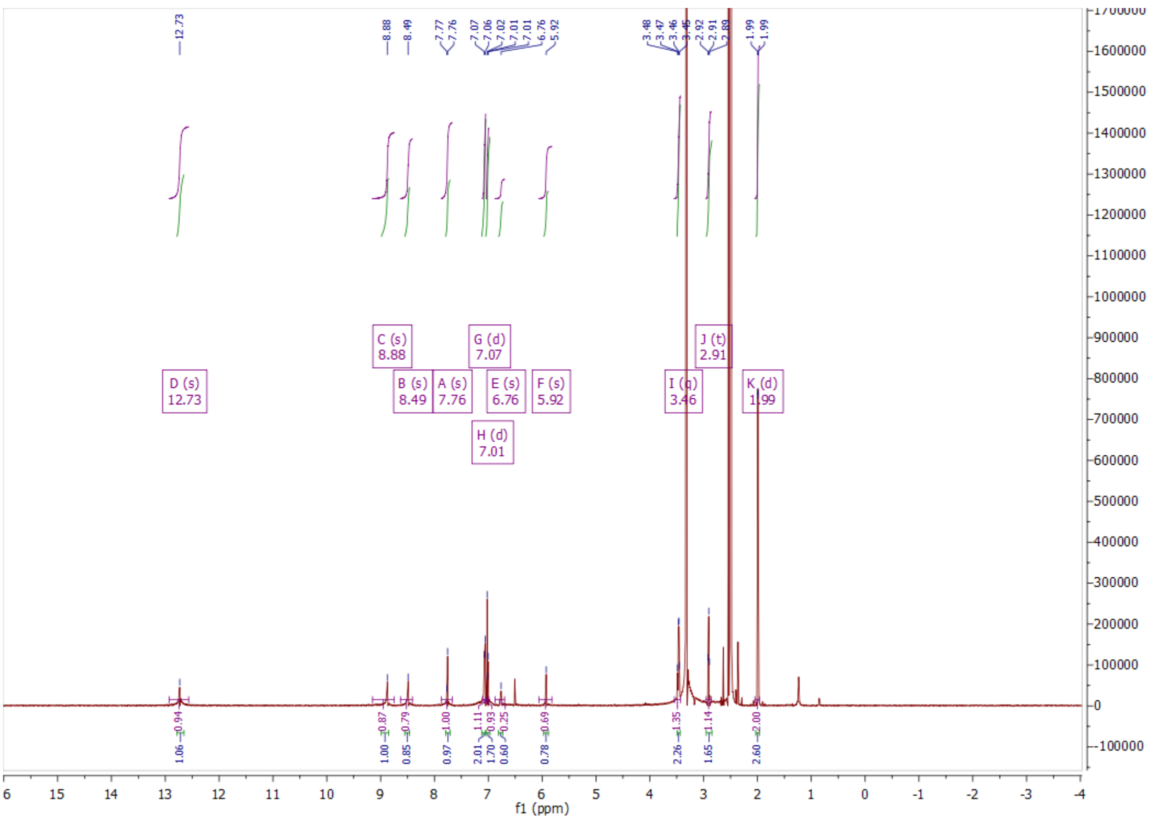
D (s)
12.73
C (s)
8.88
B (s)
8.49
A (s)
7.76
G (d)
7.07
H (d)
7.01
E (s)
6.76
F (s)
5.92
I (q)
3.46
J (t)
2.91
K (d)
1.99
f1 (ppm)

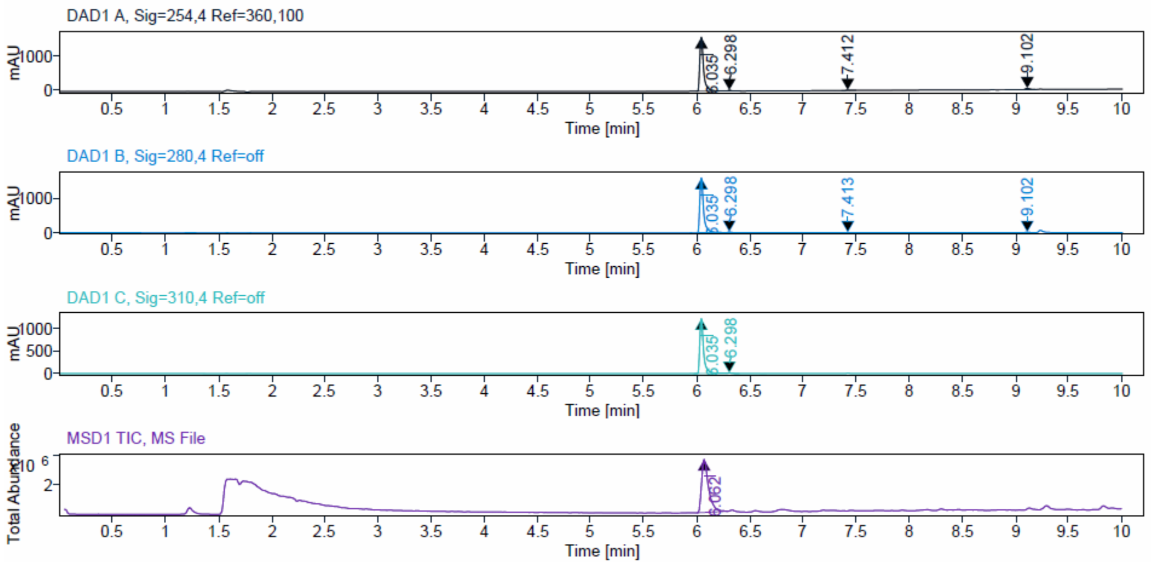
DAD1 A, Sig=254,4 Ref=360,100
DAD1 B, Sig=280,4 Ref=off
DAD1 C, Sig=310,4 Ref=off
MSD1 TIC, MS File
mAU
Total Abundance
Time [min]
6.035
6.298
7.412
7.413
9.102
6.062

Signal Description DAD1 A, Sig=254,4 Ref=360,100

| Sample Name | Name | RT | Width | Area | Area% | Height |
|---|---|---|---|---|---|---|
| JA307_DMSO_gew_Feststoff | | 6.035 | 0.034 | 3817.8962 | 95.47 | 1576.0287 |
| JA307_DMSO_gew_Feststoff | | 6.298 | 0.027 | 42.2660 | 1.06 | 22.3454 |
| JA307_DMSO_gew_Feststoff | | 7.412 | 0.045 | 27.5445 | 0.69 | 10.2990 |
| JA307_DMSO_gew_Feststoff | | 9.102 | 0.045 | 111.3475 | 2.78 | 38.2120 |

Max Area% 95.470
UV Signal Purity>95% Pass

Signal Description DAD1 B, Sig=280,4 Ref=off

| Sample Name | Name | RT | Width | Area | Area% | Height |
|---|---|---|---|---|---|---|
| JA307_DMSO_gew_Feststoff | | 6.035 | 0.034 | 3999.1626 | 97.75 | 1647.3435 |
| JA307_DMSO_gew_Feststoff | | 6.298 | 0.026 | 37.0720 | 0.91 | 19.5302 |
| JA307_DMSO_gew_Feststoff | | 7.413 | 0.049 | 25.1452 | 0.61 | 6.1407 |
| JA307_DMSO_gew_Feststoff | | 9.102 | 0.043 | 29.7854 | 0.73 | 10.4764 |

Max Area% 97.751
UV Signal Purity>95% Pass

Signal Description DAD1 C, Sig=310,4 Ref=off

| Sample Name | Name | RT | Width | Area | Area% | Height |
|---|---|---|---|---|---|---|
| JA307_DMSO_gew_Feststoff | | 6.035 | 0.034 | 3021.7954 | 99.35 | 1241.6019 |
| JA307_DMSO_gew_Feststoff | | 6.298 | 0.027 | 19.8662 | 0.65 | 10.3310 |

Max Area% 99.347
UV Signal Purity>95% Pass

ESI, HRMS, $^{1}H$, $^{13}C$ NMR and HPLC data of compound **21a**.

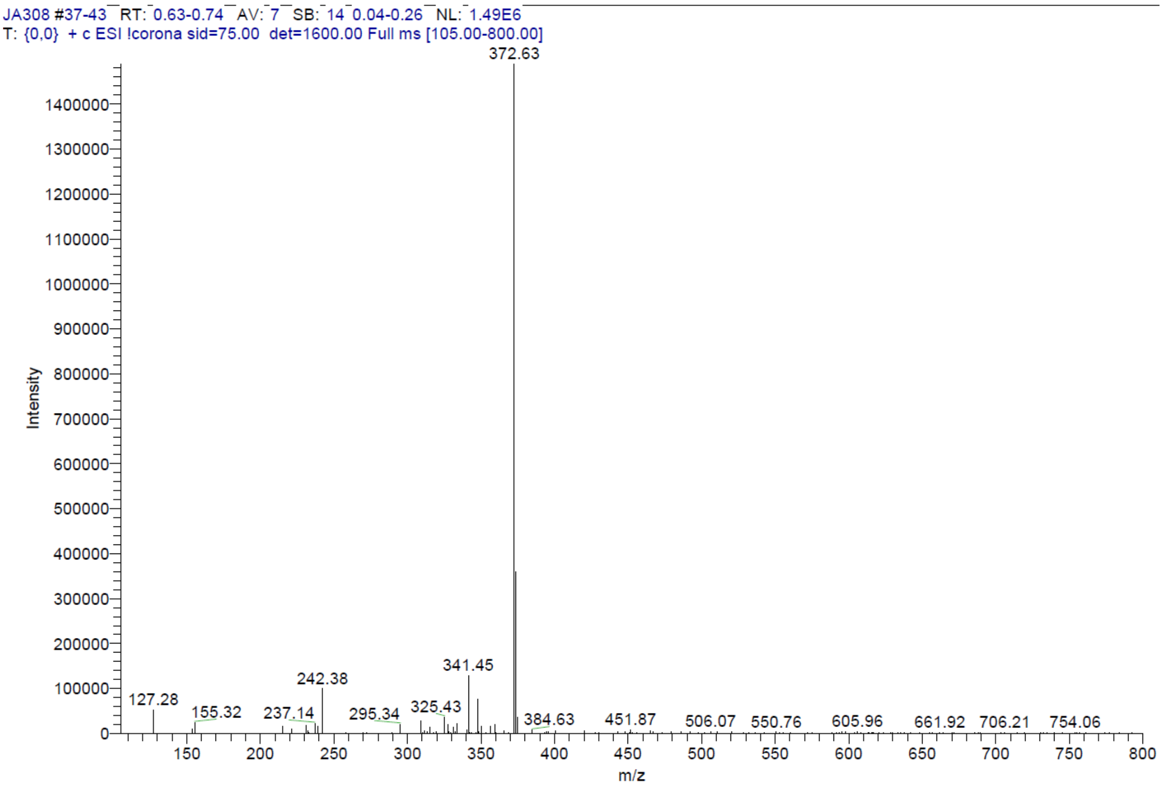


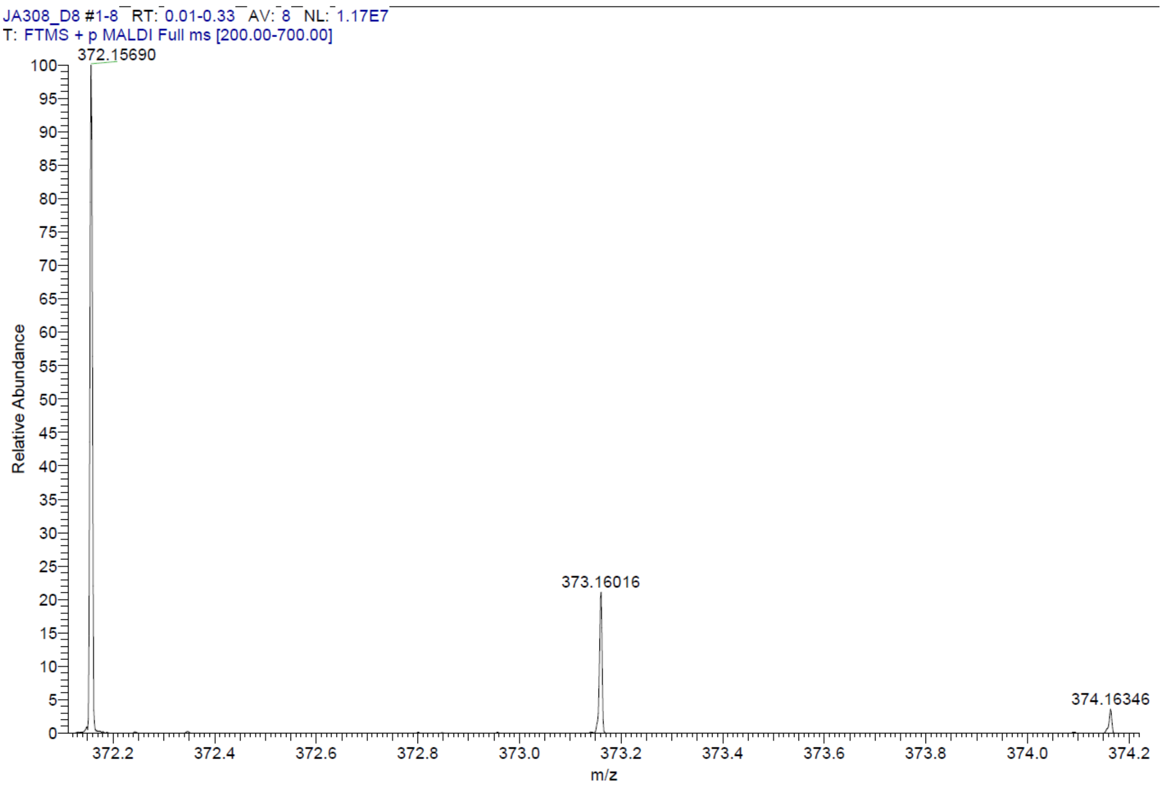

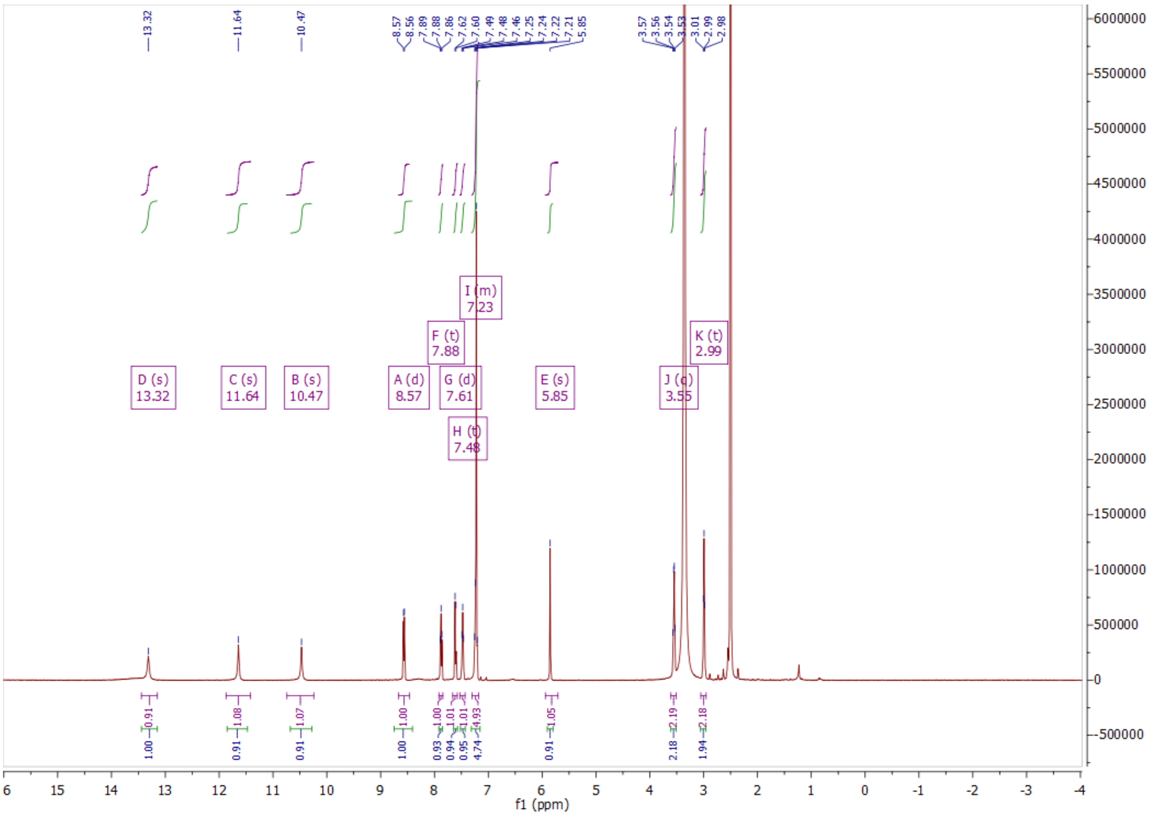
13.32
11.64
10.47
8.57 8.56 7.89 7.88 7.86 7.62 7.60 7.49 7.48 7.46 7.25 7.24 7.22 7.21 5.85
3.57 3.56 3.54 3.53 3.01 2.99 2.98
D (s) 13.32
C (s) 11.64
B (s) 10.47
A (d) 8.57
F (t) 7.88
G (d) 7.61
H (t) 7.48
I (m) 7.23
E (s) 5.85
J (q) 3.55
K (t) 2.99
1.00 0.91 0.91 1.00 0.93 0.94 0.95 4.74 0.91 2.18 1.94
0.91 1.08 1.07 1.00 1.00 1.01 1.01 4.93 1.05 2.19 2.18
f1 (ppm)

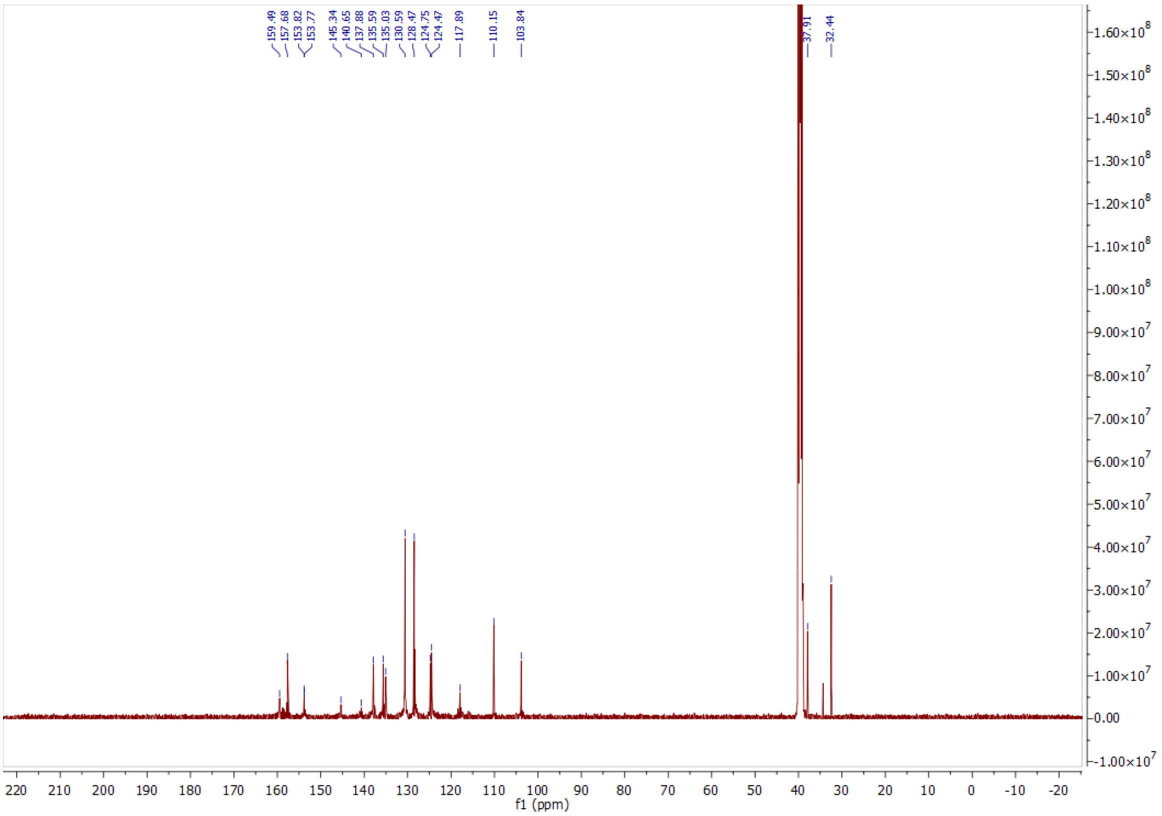
159.49 157.68 153.82 153.77 145.34 140.65 137.88 135.59 135.03 130.59 128.47 124.75 124.47 117.89 110.15 103.84
37.91 32.44
f1 (ppm)

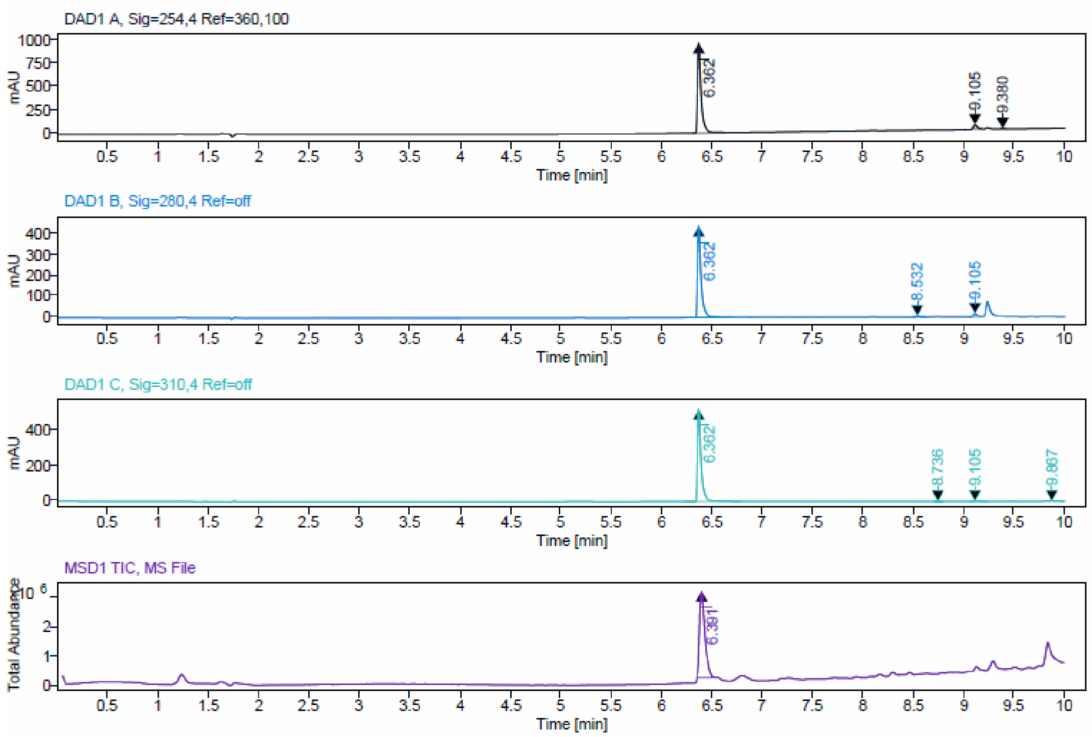


| Signal Description | DAD1 A, Sig=254,4 Ref=360,100 |
|---|---|

| Sample Name | Name | RT | Width | Area | Area% | Height |
|---|---|---|---|---|---|---|
| JA308 | | 6.362 | 0.035 | 2460.7002 | 95.55 | 970.4872 |
| JA308 | | 9.105 | 0.040 | 107.7534 | 4.18 | 42.3996 |
| JA308 | | 9.380 | 0.029 | 6.8039 | 0.26 | 3.7424 |

| Max Area% | 95.552 |
|---|---|
| UV Signal Purity>95% | Pass |

| Signal Description | DAD1 B, Sig=280,4 Ref=off |
|---|---|

| Sample Name | Name | RT | Width | Area | Area% | Height |
|---|---|---|---|---|---|---|
| JA308 | | 6.362 | 0.034 | 1098.9163 | 95.41 | 440.6306 |
| JA308 | | 8.532 | 0.035 | 14.2090 | 1.23 | 5.6370 |
| JA308 | | 9.105 | 0.044 | 38.6599 | 3.36 | 13.4714 |

| Max Area% | 95.410 |
|---|---|
| UV Signal Purity>95% | Pass |

| Signal Description | DAD1 C, Sig=310,4 Ref=off |
|---|---|

| Sample Name | Name | RT | Width | Area | Area% | Height |
|---|---|---|---|---|---|---|
| JA308 | | 6.362 | 0.034 | 1309.4125 | 98.57 | 520.4860 |
| JA308 | | 8.736 | 0.060 | 7.4379 | 0.56 | 2.4104 |
| JA308 | | 9.105 | 0.046 | 5.9130 | 0.45 | 1.7108 |
| JA308 | | 9.867 | 0.034 | 5.6593 | 0.43 | 2.3037 |

| Max Area% | 98.569 |
|---|---|
| UV Signal Purity>95% | Pass |

ESI, HRMS, $^{1}H$, $^{13}C$ NMR and HPLC data of compound **21b**.

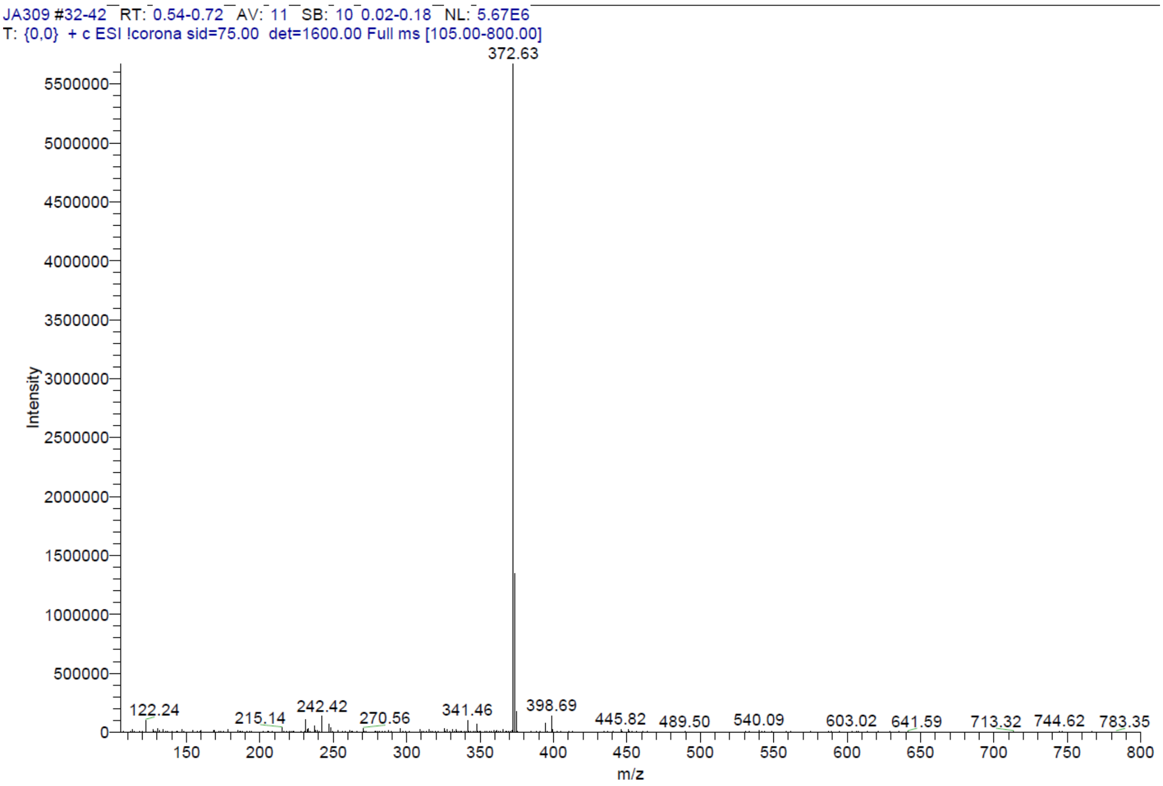


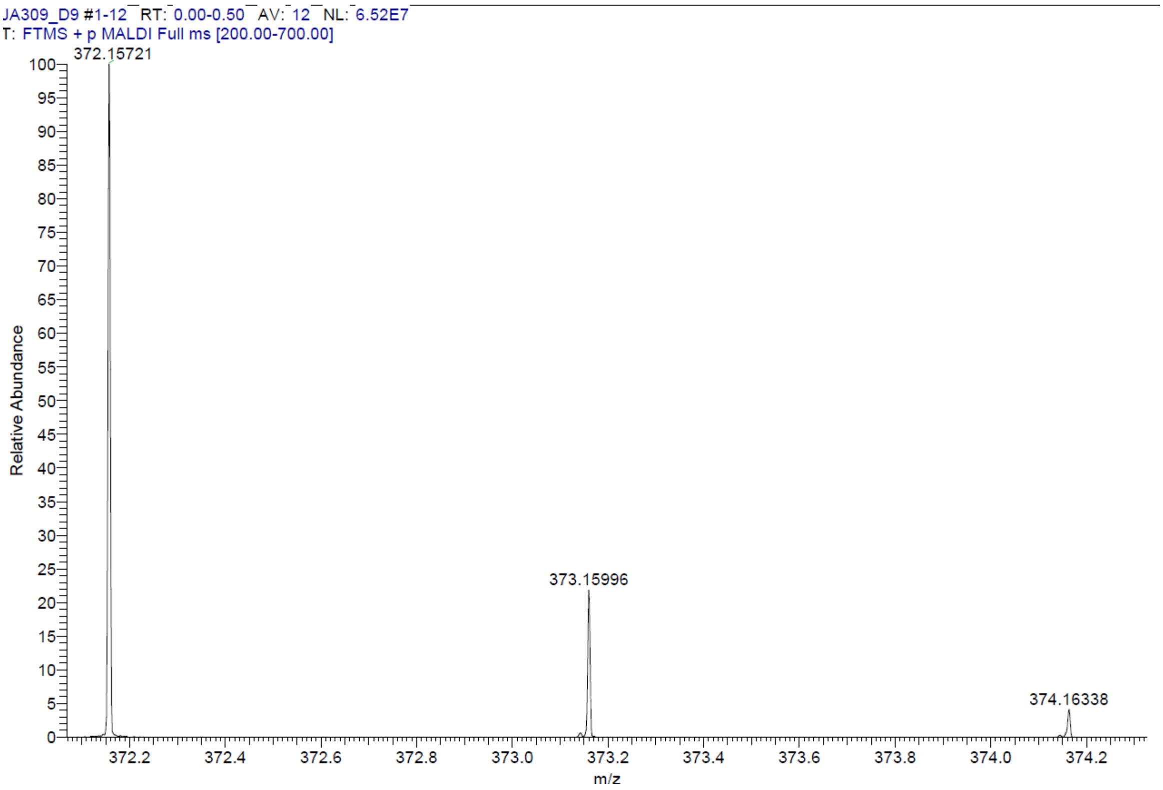

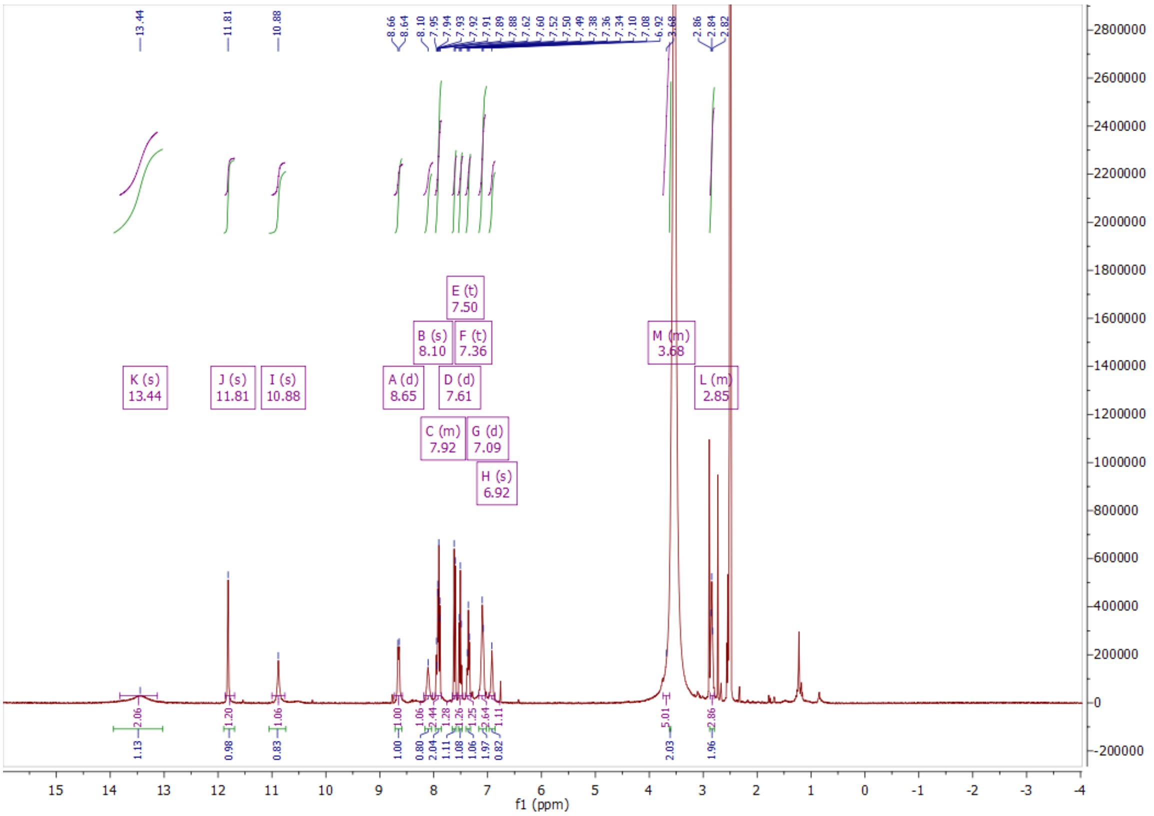

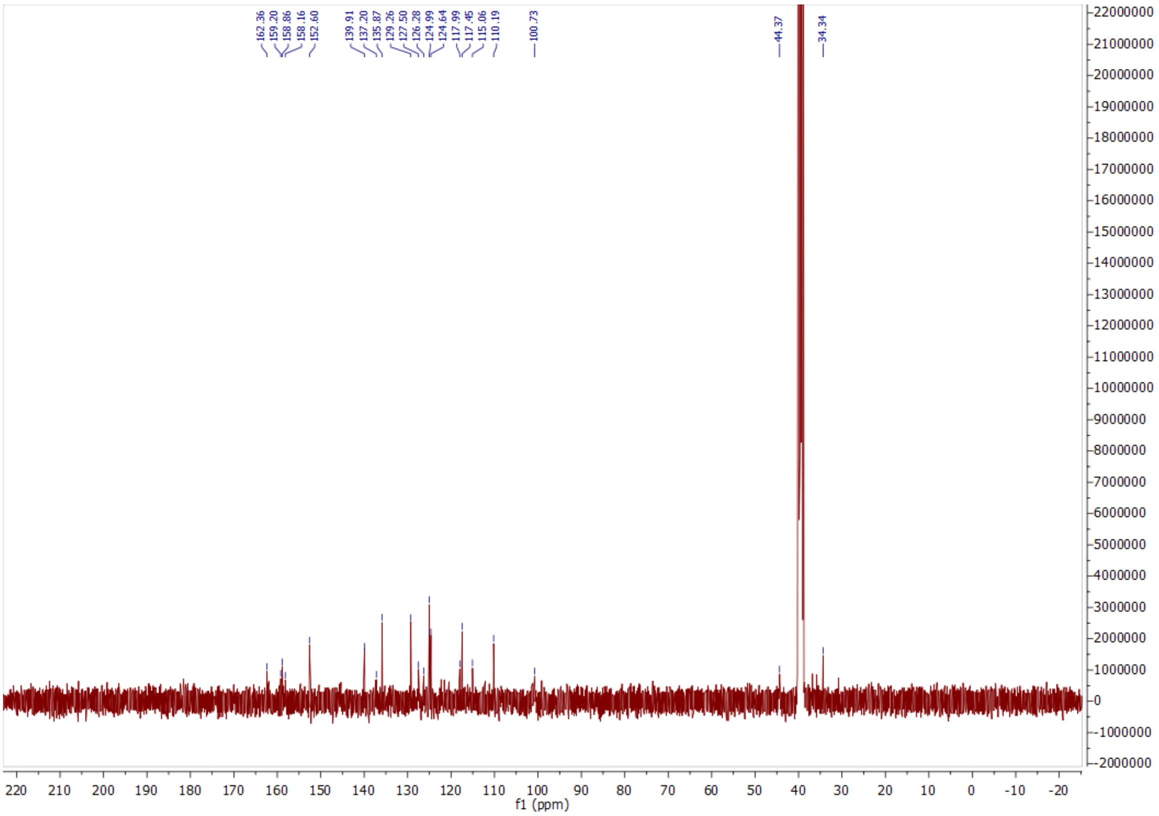

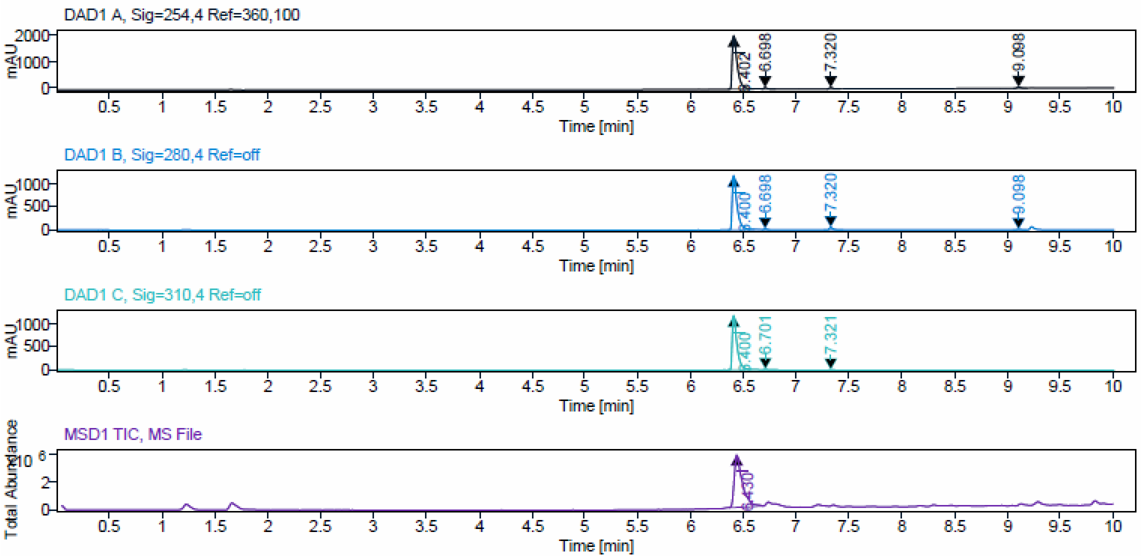


Signal Description DAD1 A, Sig=254,4 Ref=360,100

| Sample Name | Name | RT | Width | Area | Area% | Height |
|---|---|---|---|---|---|---|
| JA309_prep1_F6 | | 6.402 | 0.047 | 6587.8296 | 95.06 | 1991.6217 |
| JA309_prep1_F6 | | 6.698 | 0.027 | 103.2575 | 1.49 | 54.3592 |
| JA309_prep1_F6 | | 7.320 | 0.032 | 127.5994 | 1.84 | 56.7479 |
| JA309_prep1_F6 | | 9.098 | 0.044 | 111.5866 | 1.61 | 38.3908 |

Max Area% 95.059

UV Signal Purity>95% Pass

Signal Description DAD1 B, Sig=280,4 Ref=off

| Sample Name | Name | RT | Width | Area | Area% | Height |
|---|---|---|---|---|---|---|
| JA309_prep1_F6 | | 6.400 | 0.045 | 3711.9941 | 95.10 | 1186.5934 |
| JA309_prep1_F6 | | 6.698 | 0.027 | 31.4653 | 0.81 | 18.2172 |
| JA309_prep1_F6 | | 7.320 | 0.031 | 129.4886 | 3.32 | 56.7950 |
| JA309_prep1_F6 | | 9.098 | 0.043 | 30.1413 | 0.77 | 10.6235 |

Max Area% 95.104

UV Signal Purity>95% Pass

Signal Description DAD1 C, Sig=310,4 Ref=off

| Sample Name | Name | RT | Width | Area | Area% | Height |
|---|---|---|---|---|---|---|
| JA309_prep1_F6 | | 6.400 | 0.045 | 3668.7888 | 98.91 | 1186.0647 |
| JA309_prep1_F6 | | 6.701 | 0.032 | 26.1911 | 0.71 | 11.9842 |
| JA309_prep1_F6 | | 7.321 | 0.032 | 14.2413 | 0.38 | 5.7937 |

Max Area% 98.910

UV Signal Purity>95% Pass

ESI, HRMS, $^{1}H$, $^{13}C$ NMR and HPLC data of compound **21c**.

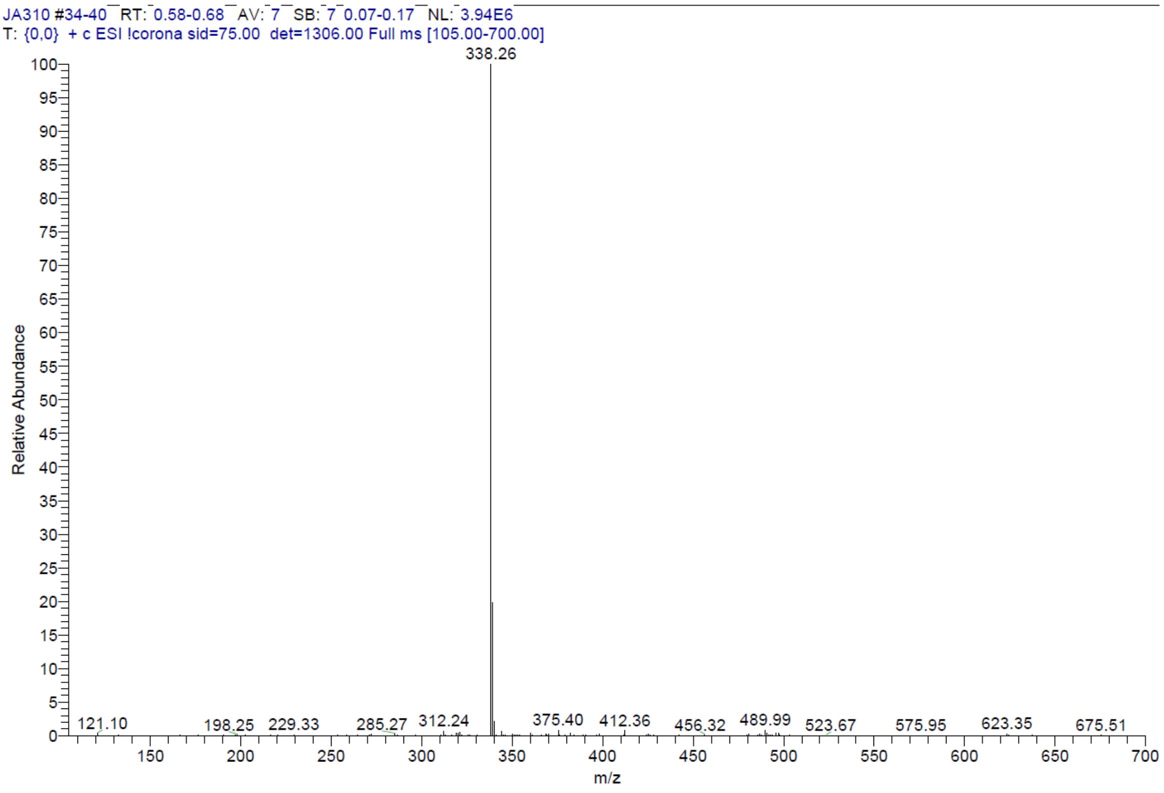


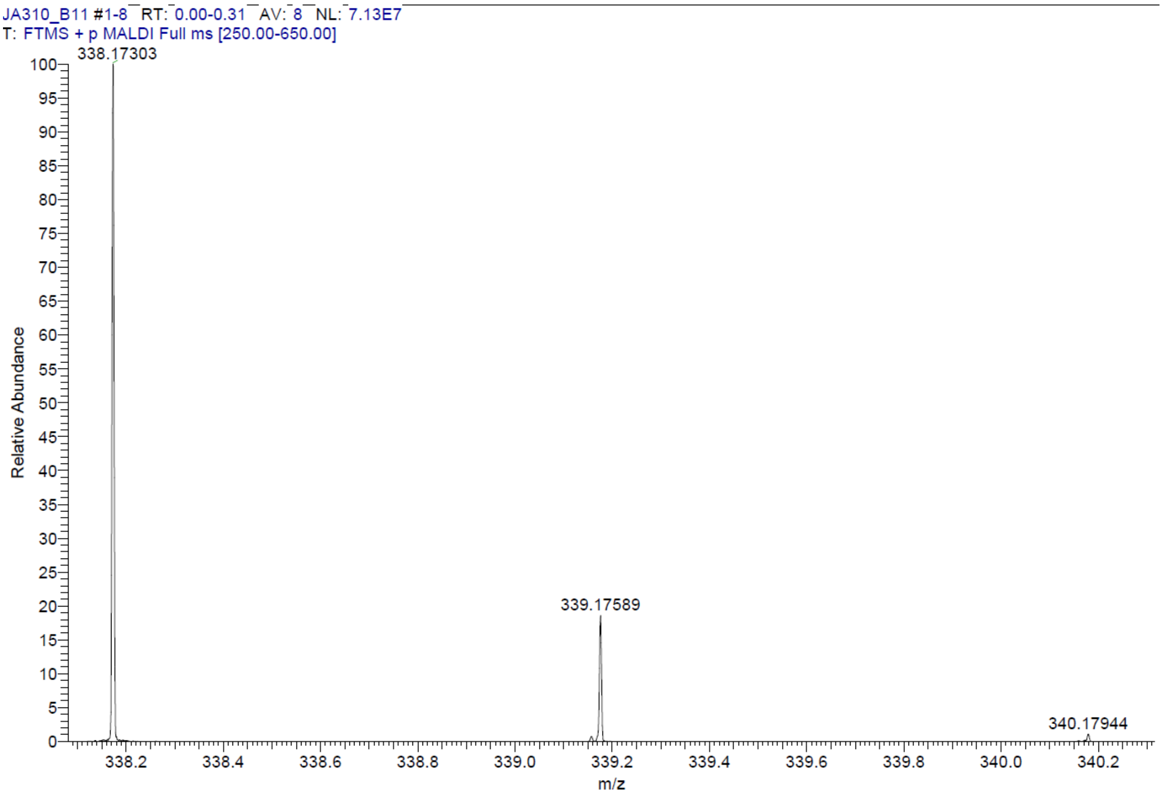

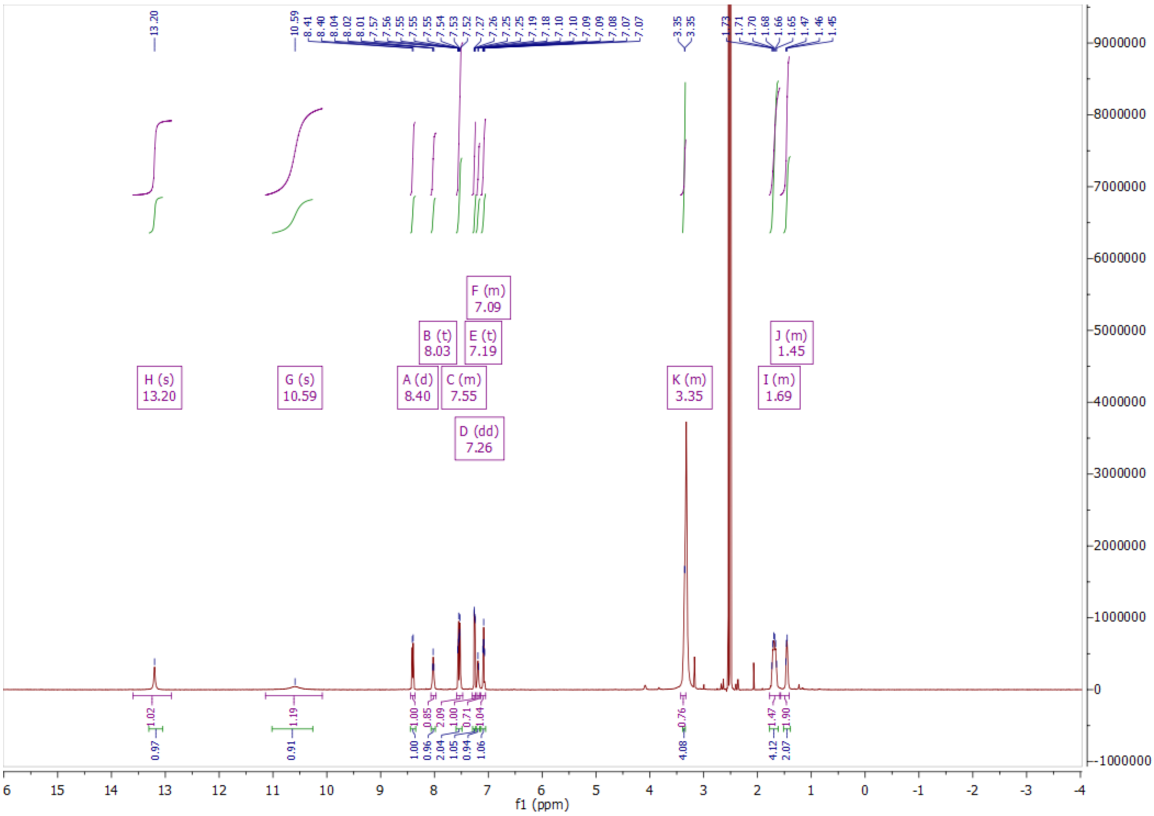
H (s) 13.20
G (s) 10.59
A (d) 8.40
B (t) 8.03
C (m) 7.55
D (dd) 7.26
E (t) 7.19
F (m) 7.09
K (m) 3.35
I (m) 1.69
J (m) 1.45
f1 (ppm)

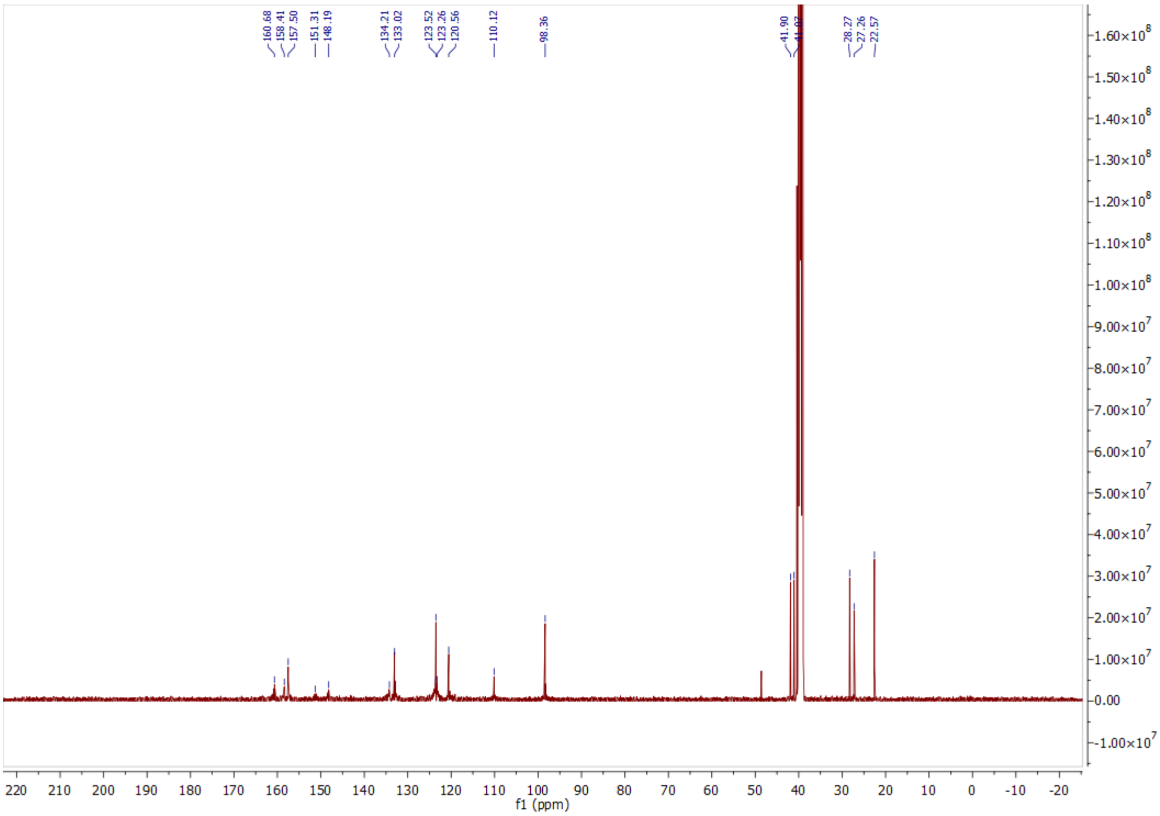
160.68
158.41
157.50
151.31
148.19
134.21
133.02
123.52
123.26
120.56
110.12
98.36
41.90
28.27
27.26
22.57
f1 (ppm)

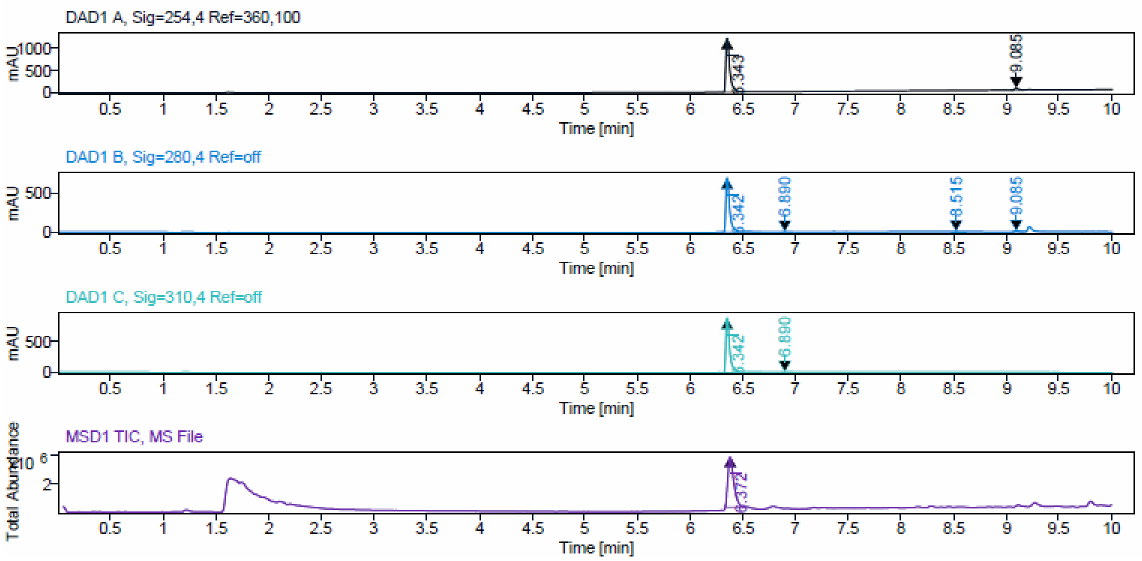


Signal Description DAD1 A, Sig=254,4 Ref=360,100

| Sample Name | Name | RT | Width | Area | Area% | Height |
|---|---|---|---|---|---|---|
| JA310_feststoff_verd | | 6.343 | 0.036 | 3023.5352 | 96.15 | 1210.3330 |
| JA310_feststoff_verd | | 9.085 | 0.044 | 121.0906 | 3.85 | 41.5302 |

Max Area% 96.149

UV Signal Purity>95% Pass

Signal Description DAD1 B, Sig=280,4 Ref=off

| Sample Name | Name | RT | Width | Area | Area% | Height |
|---|---|---|---|---|---|---|
| JA310_feststoff_verd | | 6.342 | 0.035 | 1793.2151 | 97.11 | 719.7684 |
| JA310_feststoff_verd | | 6.890 | 0.030 | 7.1919 | 0.39 | 3.3742 |
| JA310_feststoff_verd | | 8.515 | 0.035 | 13.6499 | 0.74 | 5.4570 |
| JA310_feststoff_verd | | 9.085 | 0.043 | 32.4976 | 1.76 | 11.4949 |

Max Area% 97.111

UV Signal Purity>95% Pass

Signal Description DAD1 C, Sig=310,4 Ref=off

| Sample Name | Name | RT | Width | Area | Area% | Height |
|---|---|---|---|---|---|---|
| JA310_feststoff_verd | | 6.342 | 0.035 | 2214.8662 | 99.59 | 887.2886 |
| JA310_feststoff_verd | | 6.890 | 0.030 | 9.0342 | 0.41 | 4.2442 |

Max Area% 99.594

UV Signal Purity>95% Pass

ESI, HRMS, [1]H, [13]C NMR and HPLC data of compound **21d**.

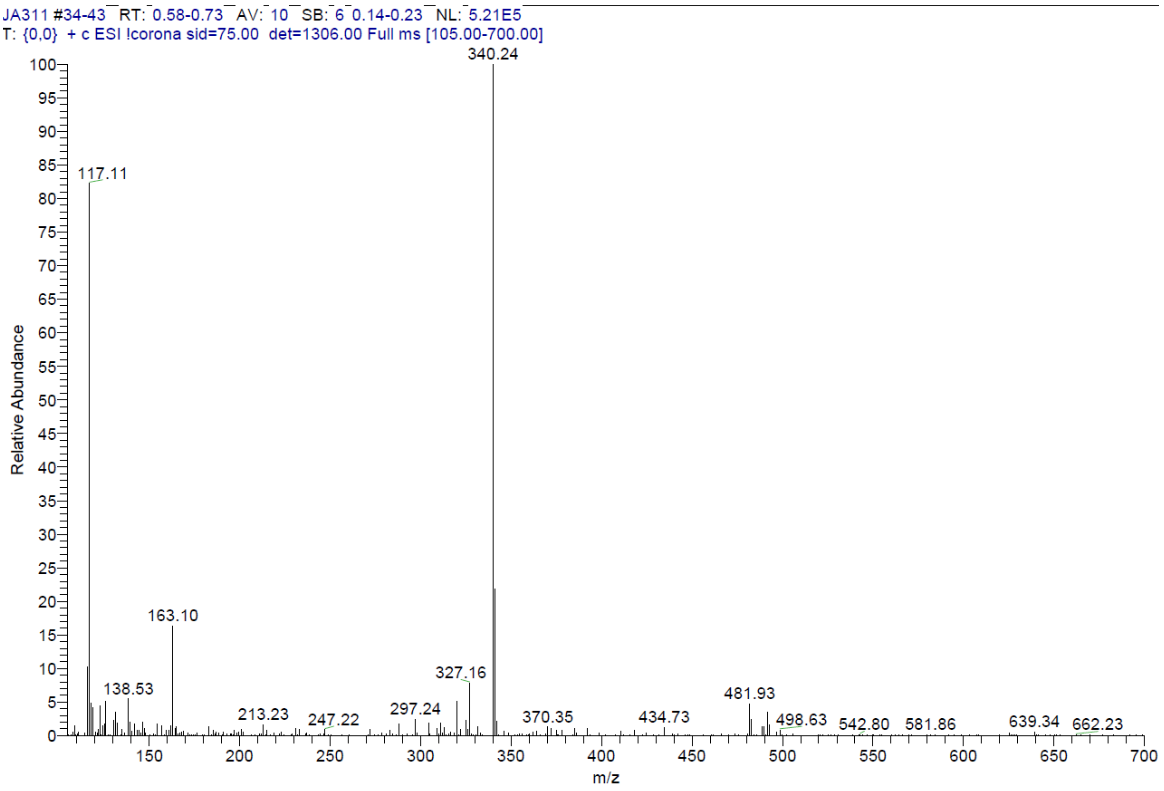


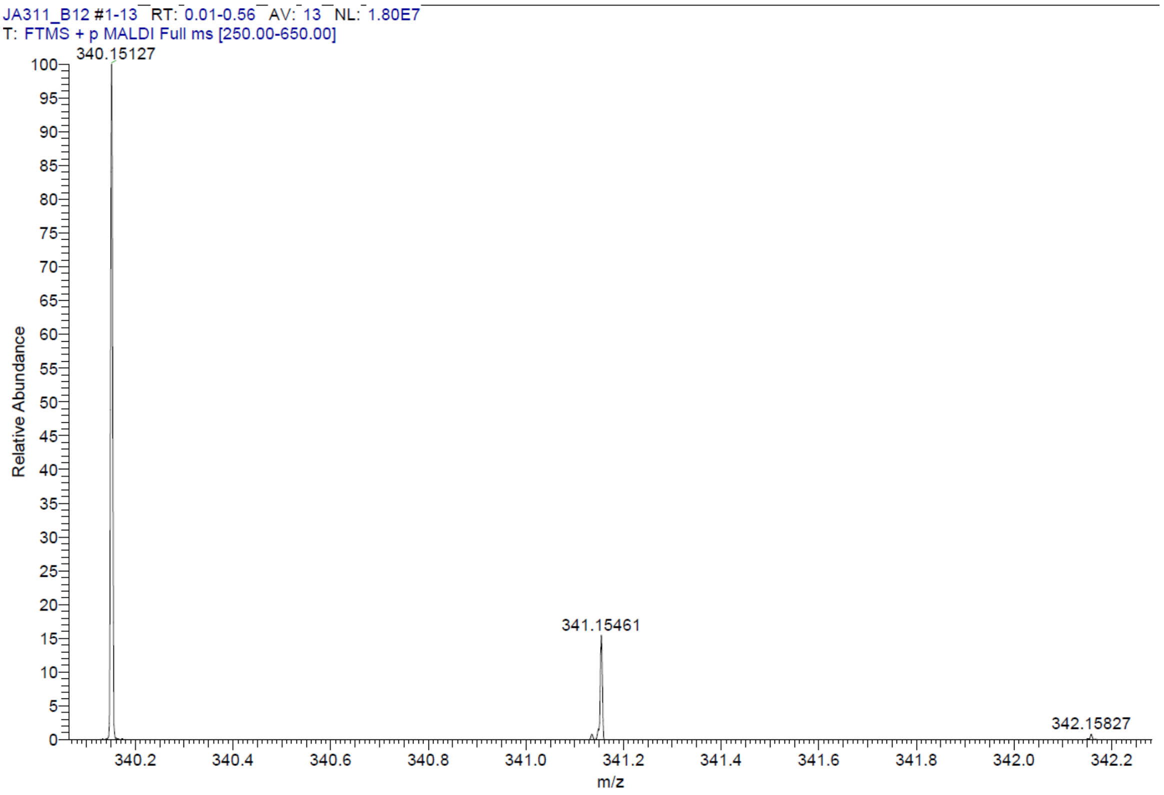

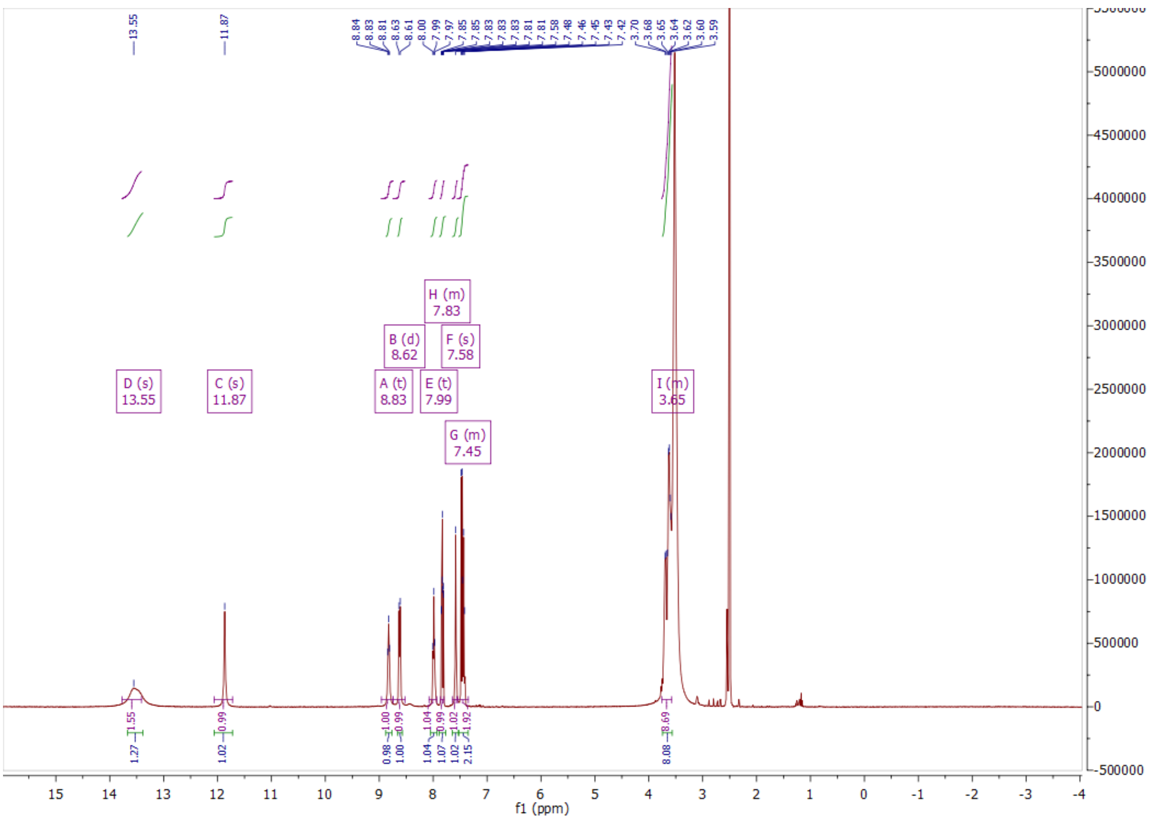
D (s)
13.55
C (s)
11.87
A (t)
8.83
B (d)
8.62
E (t)
7.99
H (m)
7.83
F (s)
7.58
G (m)
7.45
I (m)
3.65
f1 (ppm)

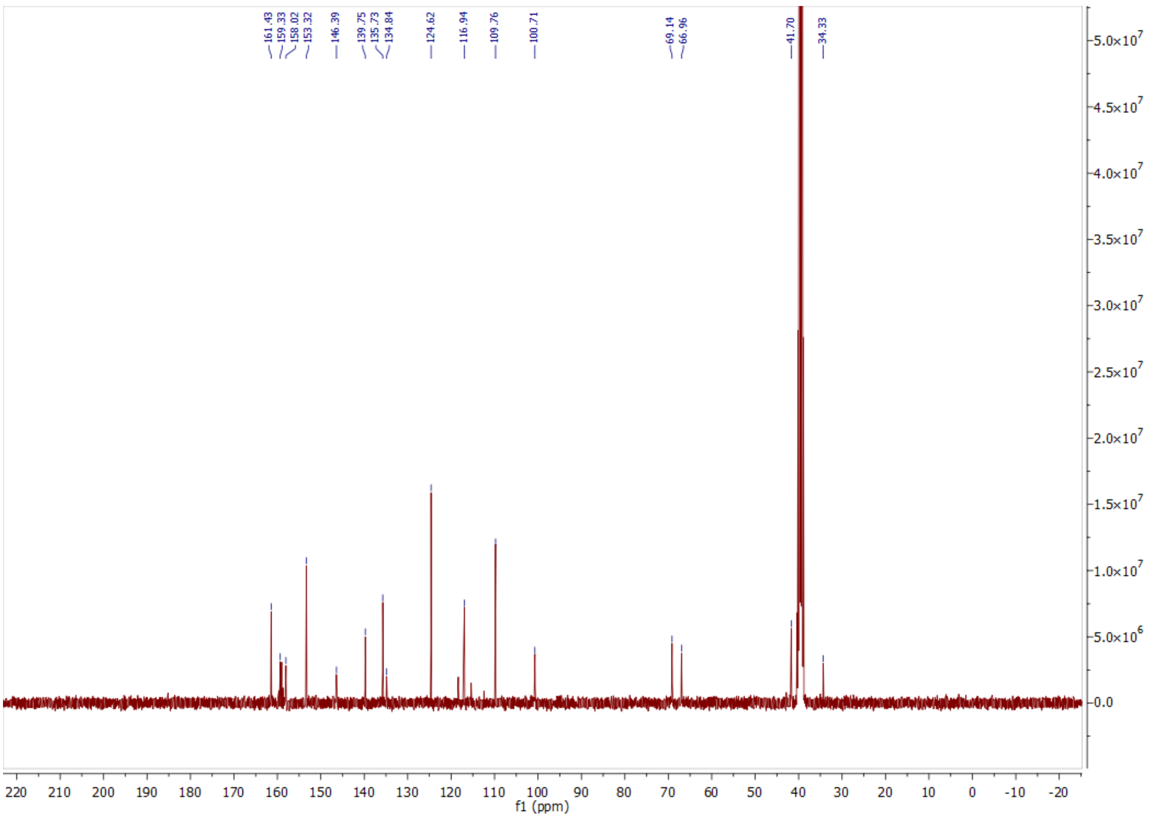
161.43
159.33
158.02
153.32
146.39
139.75
135.73
134.84
124.62
116.94
109.76
100.71
69.14
66.96
41.70
34.33
f1 (ppm)

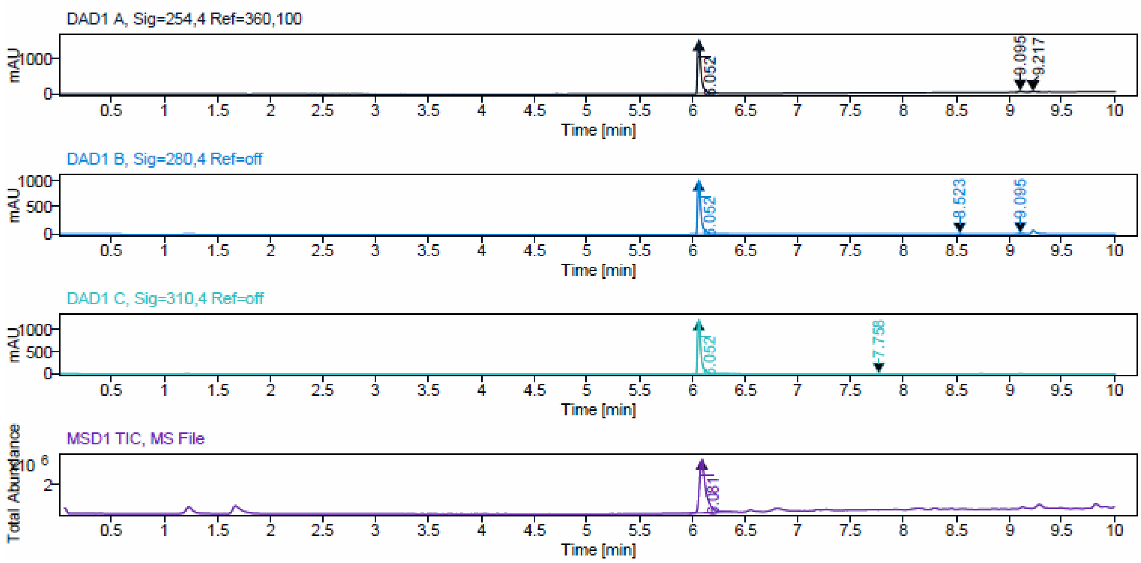


Signal Description DAD1 A, Sig=254,4 Ref=360,100

| Sample Name | Name | RT | Width | Area | Area% | Height |
|---|---|---|---|---|---|---|
| JA311_prepF10 | | 6.052 | 0.031 | 3462.2913 | 96.08 | 1557.5930 |
| JA311_prepF10 | | 9.095 | 0.044 | 109.1214 | 3.03 | 38.2731 |
| JA311_prepF10 | | 9.217 | 0.037 | 32.2700 | 0.90 | 13.3005 |

Max Area% 96.076

UV Signal Purity>95% Pass

Signal Description DAD1 B, Sig=280,4 Ref=off

| Sample Name | Name | RT | Width | Area | Area% | Height |
|---|---|---|---|---|---|---|
| JA311_prepF10 | | 6.052 | 0.029 | 2280.1594 | 98.19 | 1056.1769 |
| JA311_prepF10 | | 8.523 | 0.037 | 12.4935 | 0.54 | 5.4809 |
| JA311_prepF10 | | 9.095 | 0.043 | 29.4275 | 1.27 | 10.5853 |

Max Area% 98.195

UV Signal Purity>95% Pass

Signal Description DAD1 C, Sig=310,4 Ref=off

| Sample Name | Name | RT | Width | Area | Area% | Height |
|---|---|---|---|---|---|---|
| JA311_prepF10 | | 6.052 | 0.030 | 2717.5564 | 99.49 | 1252.4003 |
| JA311_prepF10 | | 7.758 | 0.088 | 13.8282 | 0.51 | 2.9480 |

Max Area% 99.494

UV Signal Purity>95% Pass

ESI, HRMS, $^{1}H$, $^{13}C$ NMR and HPLC data of compound **21e**.

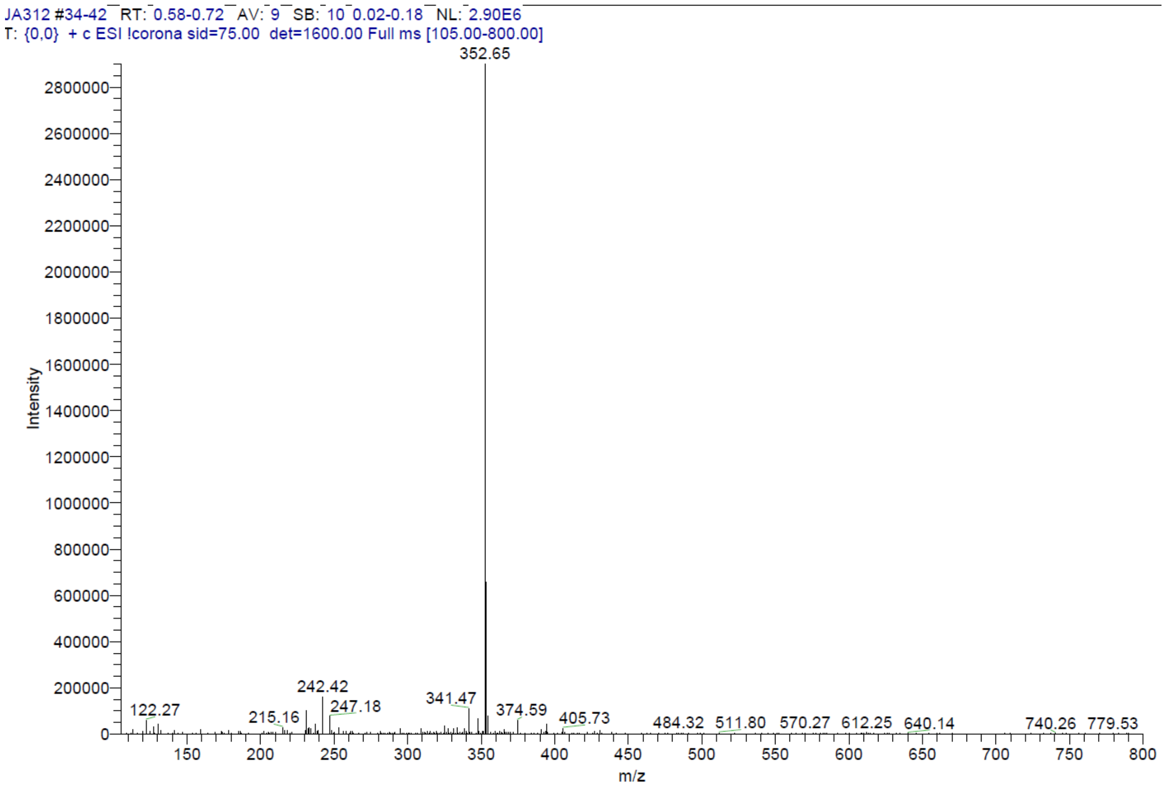


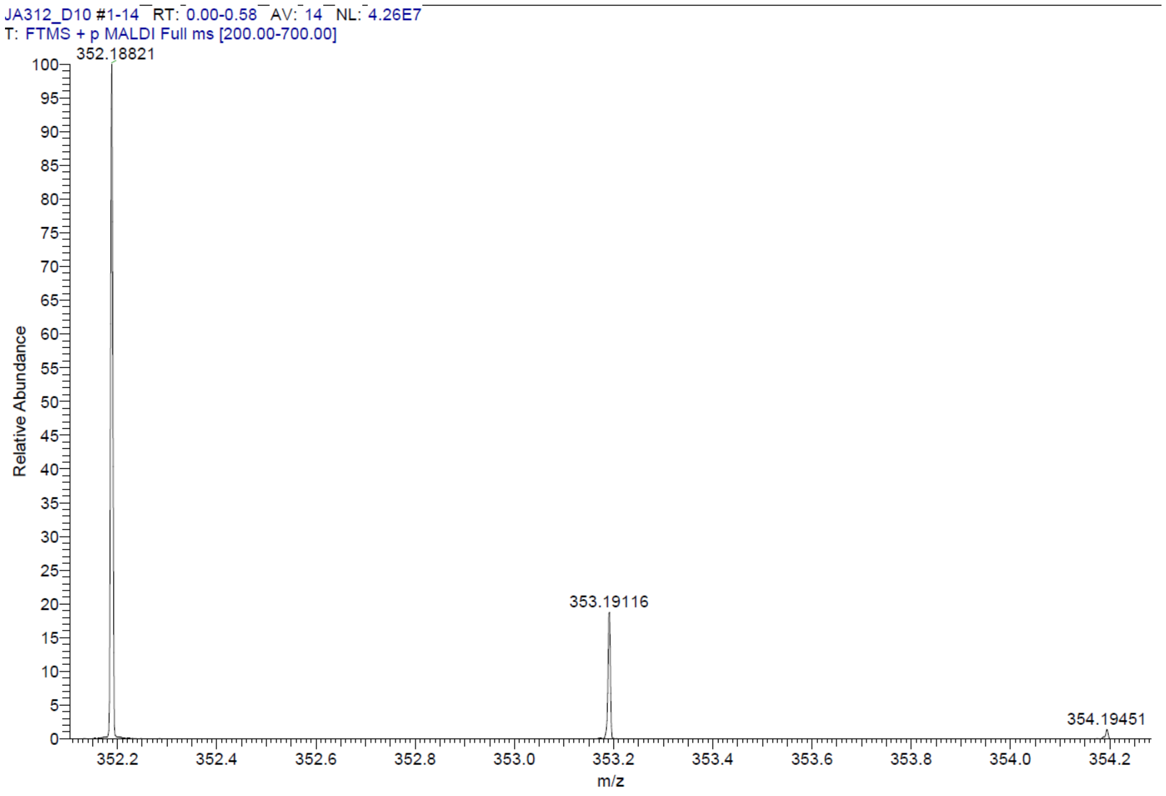

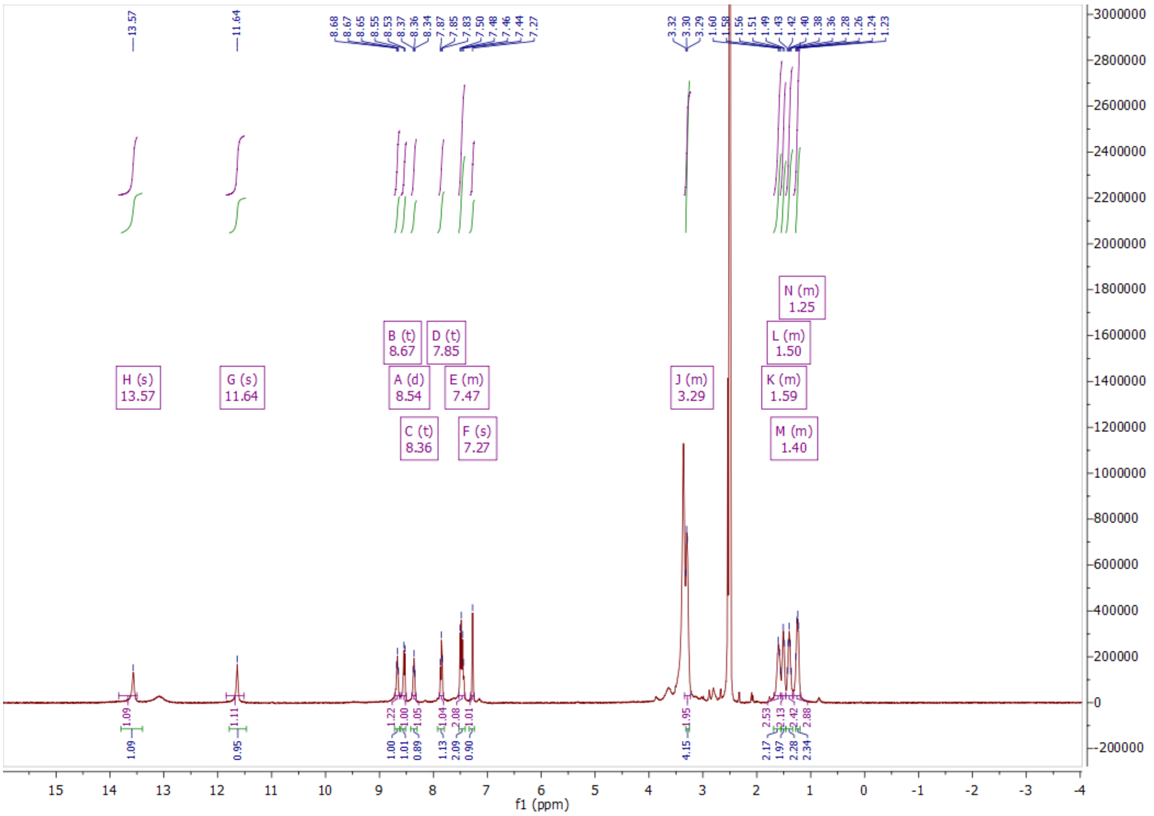

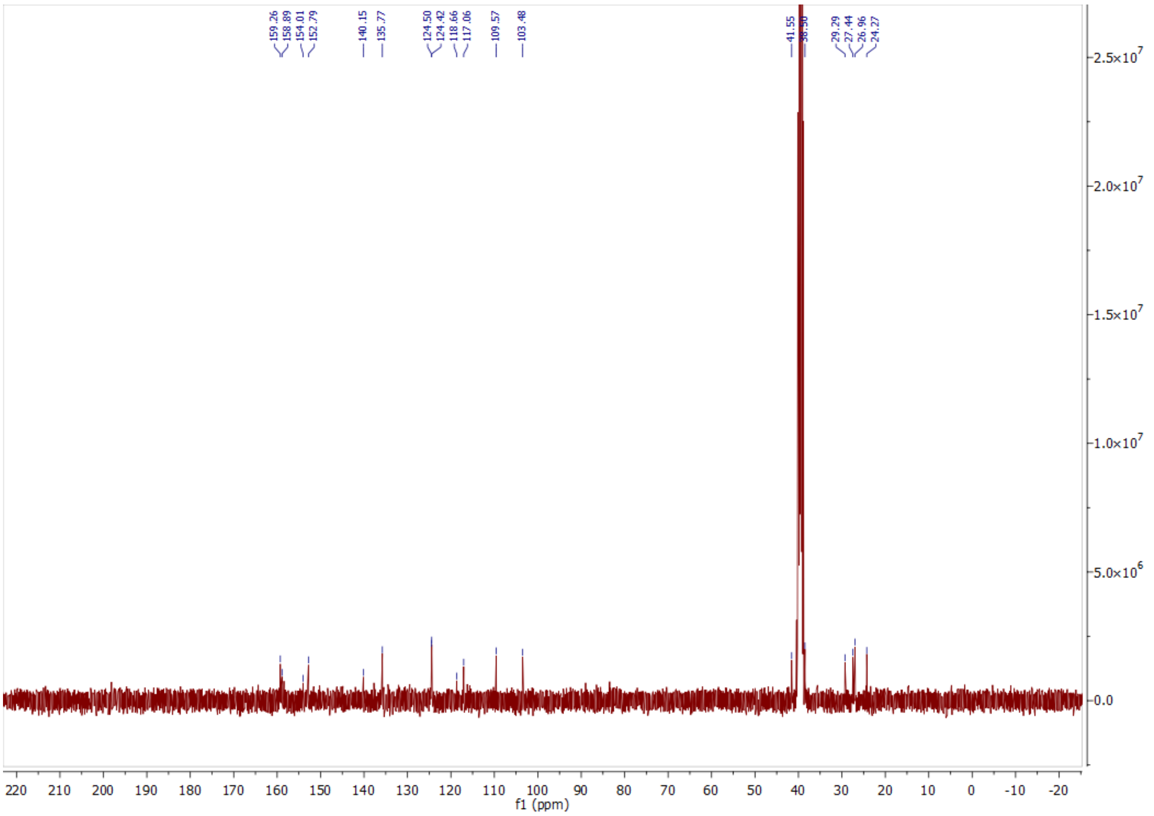

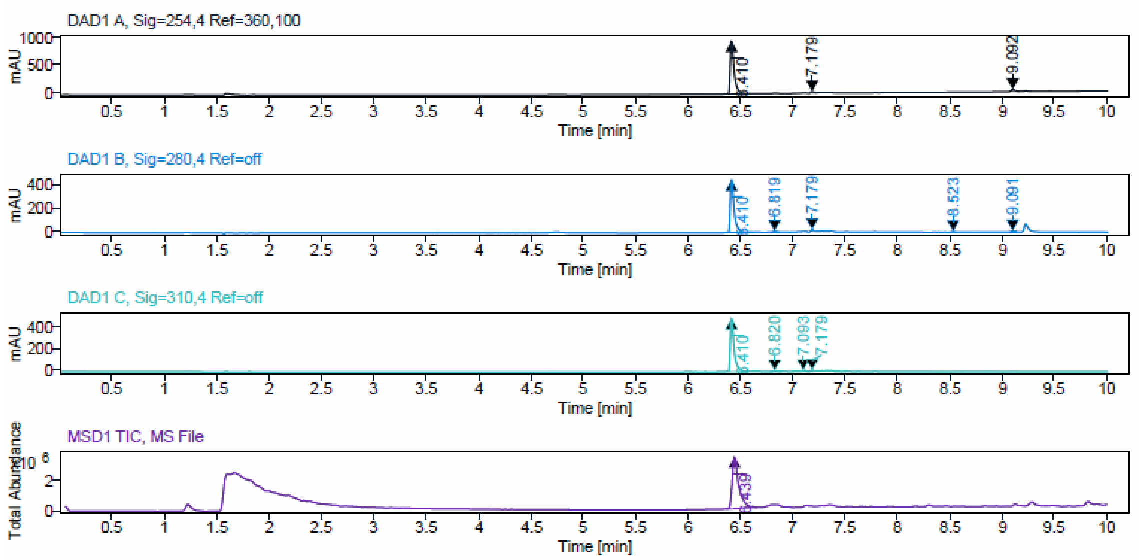


Signal Description DAD1 A, Sig=254,4 Ref=360,100

| Sample Name | Name | RT | Width | Area | Area% | Height |
|---|---|---|---|---|---|---|
| JA312_F25-27_festst_gew_neu | | 6.410 | 0.036 | 2342.4419 | 95.03 | 925.2548 |
| JA312_F25-27_festst_gew_neu | | 7.179 | 0.027 | 37.2181 | 1.51 | 22.8373 |
| JA312_F25-27_festst_gew_neu | | 9.092 | 0.040 | 85.2498 | 3.46 | 34.0602 |

Max Area% 95.032

UV Signal Purity>95% Pass

Signal Description DAD1 B, Sig=280,4 Ref=off

| Sample Name | Name | RT | Width | Area | Area% | Height |
|---|---|---|---|---|---|---|
| JA312_F25-27_festst_gew_neu | | 6.410 | 0.035 | 1138.9548 | 95.07 | 445.1817 |
| JA312_F25-27_festst_gew_neu | | 6.819 | 0.042 | 16.9998 | 1.42 | 7.0401 |
| JA312_F25-27_festst_gew_neu | | 7.179 | 0.025 | 19.5159 | 1.63 | 14.2377 |
| JA312_F25-27_festst_gew_neu | | 8.523 | 0.026 | 6.1444 | 0.51 | 3.4809 |
| JA312_F25-27_festst_gew_neu | | 9.091 | 0.035 | 16.4315 | 1.37 | 8.1017 |

Max Area% 95.068

UV Signal Purity>95% Pass

Signal Description DAD1 C, Sig=310,4 Ref=off

| Sample Name | Name | RT | Width | Area | Area% | Height |
|---|---|---|---|---|---|---|
| JA312_F25-27_festst_gew_neu | | 6.410 | 0.035 | 1186.4512 | 95.02 | 475.2104 |
| JA312_F25-27_festst_gew_neu | | 6.820 | 0.048 | 24.1487 | 1.93 | 8.5165 |
| JA312_F25-27_festst_gew_neu | | 7.093 | 0.045 | 18.9328 | 1.52 | 7.0253 |
| JA312_F25-27_festst_gew_neu | | 7.179 | 0.028 | 19.1242 | 1.53 | 11.2304 |

Max Area% 95.018

UV Signal Purity>95% Pass
